# Insights into dispersed duplications and complex structural mutations from whole genome sequencing 706 families

**DOI:** 10.1101/2020.08.03.235358

**Authors:** Christopher W. Whelan, Robert E. Handsaker, Giulio Genovese, Seva Kashin, Monkol Lek, Jason Hughes, Joshua McElwee, Michael Lenardo, Daniel MacArthur, Steven A. McCarroll

## Abstract

Two intriguing forms of genome structural variation (SV) – dispersed duplications, and de novo rearrangements of complex, multi-allelic loci – have long escaped genomic analysis. We describe a new way to find and characterize such variation by utilizing identity-by-descent (IBD) relationships between siblings together with high-precision measurements of segmental copy number. Analyzing whole-genome sequence data from 706 families, we find hundreds of “IBD-discordant” (IBDD) CNVs: loci at which siblings’ CNV measurements and IBD states are mathematically inconsistent. We found that commonly-IBDD CNVs identify dispersed duplications; we mapped 95 of these common dispersed duplications to their true genomic locations through family-based linkage and population linkage disequilibrium (LD), and found several to be in strong LD with genome-wide association (GWAS) signals for common diseases or gene expression variation at their revealed genomic locations. Other CNVs that were IBDD in a single family appear to involve *de novo* mutations in complex and multi-allelic loci; we identified 26 *de novo* structural mutations that had not been previously detected in earlier analyses of the same families by diverse SV analysis methods. These included a *de novo* mutation of the amylase gene locus and multiple *de novo* mutations at chromosome 15q14. Combining these complex mutations with more-conventional CNVs, we estimate that segmental mutations larger than 1kb arise in about one per 22 human meioses. These methods are complementary to previous techniques in that they interrogate genomic regions that are home to segmental duplication, high CNV allele frequencies, and multi-allelic CNVs.

**Author Summary:** Copy number variation is an important form of genetic variation in which individuals differ in the number of copies of segments of their genomes. Certain aspects of copy number variation have traditionally been difficult to study using short-read sequencing data. For example, standard analyses often cannot tell whether the duplicated copies of a segment are located near the original copy or are dispersed to other regions of the genome. Another aspect of copy number variation that has been difficult to study is the detection of mutations in the copy number of DNA segments passed down from parents to their children, particularly when the mutations affect genome segments which already display common copy number variation in the population. We develop an analytical approach to solving these problems when sequencing data is available for all members of families with at least two children. This method is based on determining the number of parental haplotypes the two siblings share at each location in their genome, and using that information to determine the possible inheritance patterns that might explain the copy numbers we observe in each family member. We show that dispersed duplications and mutations can be identified by looking for copy number variants that do not follow these expected inheritance patterns. We use this approach to determine the location of 95 common duplications which are dispersed to distant regions of the genome, and demonstrate that these duplications are linked to genetic variants that affect disease risk or gene expression levels. We also identify a set of copy number mutations not detected by previous analyses of sequencing data from a large cohort of families, and show that repetitive and complex regions of the genome undergo frequent mutations in copy number.

## Introduction

Genome structural variation involves the deletion, duplication, or rearrangement of genomic segments. Though the identification of deletions and duplications is today an important part of genome analysis, more complex features of genome structural variation remain invisible to routine analyses. These features include the structure of dispersed duplications and the presence of *de novo* structural rearrangements at complex loci. These questions are even more difficult to study when copy number variants (CNVs) occur at genomic loci which are part of segmental duplications or multi-allelic CNVs, which contribute most genetic variation in gene dosage across the genome [1].

Dispersed duplications, in which a duplicated segment is inserted into a genomic locus distant from the original copy, have been a particularly challenging form of copy number variation to analyze. Some categories of dispersed duplication formation have been described and catalogued, including those created by LINE transduction [2] and the retrotransposition of pseudogenes [3]. However, due to the enrichment of CNVs in segmental duplications and other repetitive sequences, it is often difficult to determine where duplicate copies of a genomic segment reside; most CNV detection tools report duplications at the location of the “known” sequence copy. This is often assumed to mean that the CNV is a tandem duplication, although the true variant might be far from the original reference segment. Such an assumption can confound downstream analyses of genomic variation; for example, a dispersed duplication called as a CNV at the original segment site will not be in linkage disequilibrium (LD) with nearby variants, possibly causing disease association signals to be discarded or obscuring functional effects of dispersed duplication insertions. Previous studies have attempted to map dispersed duplication insertion locations using LD signal with nearby SNPs [4] or by analyzing and assembling read pair signals [5], but in many cases neither of these signals can be measured unambiguously with short-read sequencing data due to repetitive sequence at or near the insertion point.

Though *de novo* mutations have a large impact on neurodevelopmental and other disorders, analysis of *de novo* copy-number mutations has largely been limited to simple, single-copy regions of the genome. Previous studies have identified large *de novo* copy number changes in families and shown that they contribute to risk for diseases including autism and schizophrenia [6,7]. More-recent studies have used whole genome sequencing data from families, including an autism cohort used in this study, to identify smaller *de novo* CNVs [8–10]. All of these analyses, however, have largely depended on measuring an increase or decrease in the total array intensity, read depth, or paired-end and split read mapping signals in a proband as compared to the same signals in both parents or the entire cohort. Such approaches therefore do not generally detect *de novo* copy number changes at sites of common or multi-allelic CNVs (mCNVs), since doing so comprehensively requires at least partial decomposition of the copy-number alleles transmitted within a family. Palta et al. [11] devised a strategy for this problem based on combining CNVs called from array genotyping data, B-allele frequency, and pedigrees, but were limited by the resolution and accuracy of array-based data and could solve only some cases of CNV transmission. DNA data from families has long been used to reveal and apply principles of molecular inheritance. Family data provides powerful ways to critically evaluate genotype or sequence data quality, as high-quality data should have few inconsistencies with Mendelian inheritance. Rare exceptions to Mendelian inheritance are today used to find *de novo* mutations in father-mother-offspring trios [12,13]. Apparent Mendelian inconsistencies, when genomically clustered, revealed deletion polymorphisms segregating in the human genome [14,15]. Data from more family members allows more-complex analyses. The sharing of long genomic segments among relatives in three-generation pedigrees was used to create the first linkage maps of the human genome [16]. The boundaries of shared segments correspond to crossover events, which have been inferred from family-level single nucleotide polymorphism (SNP) and sequence data [17,18].

Sequencing data from quartets allows the mapping, along each chromosome, of siblings’ identity-by-descent (IBD) state – the number of parental haplotypes (ranging from zero to two) shared by the siblings at every genomic locus. Determination of IBD state in quartets has previously been used to identify crossovers, control genotyping error rates, improve phasing, and accurately estimate mutation rates for SNPs and small indels [18–20].

Here, we show that IBD state also makes strong, testable predictions about the mathematical relationships among copy-number measurements for parents and children – and that violations of these relationships can be leveraged to conduct previously impossible analyses of some kinds of copy number variation (CNV). We describe a new approach based on the combination of IBD analysis and accurate copy number quantitation from whole genome sequencing data, finding that this approach allows insights about dispersed duplications, even at genomic loci with complex and multiallelic copy number variation, and can be used to identify previously hidden *de novo* copy number changes.

## Results

Our approach to analyzing family-level sequence data is to find genomic regions in which segmental copy number measurements and sibling IBD states initially appear to be incompatible. IBD state makes strong, testable predictions about the mathematical relationships among copy-number measurements for parents and children (**Fig 1A**), provided that segmental copy number can be measured precisely (rather than just approximately). For example, if siblings are IBD-2 at a locus, then all segmental copy-number measurements should agree precisely; if they are IBD-0, then their copy number measurements should sum to the same integer as the sum of their parents. A more-complex mathematical relationship governs IBD-1 regions (**Fig 1A**). These rules are akin to determining whether a set of SNP genotypes in a pedigree are concordant with Mendelian transmission of alleles, but they generalize this approach to the case in which a genotype can be any of many different possible integers. Violations of these relationships can be detected and further analyzed. An overview of our workflow is given in **Fig 1B**. We first use whole genome sequencing (WGS) data aligned to GRCh37 to call SNPs and indels; these data inform the inference of IBD state across the genome for each quartet. In addition to using fine-scale variants to analyze IBD, we also use read depth signal to identify CNVs, and, critically, to determine integer copy number at every CNV site, in each genome in the cohort. We refer to the copy number state of all four members of a quartet at a CNV locus as a *qCNV*. We combine IBD state information with qCNVs to evaluate whether the segregation of copy numbers in the family is concordant with the IBD state at the putative locus of the CNV. We then further evaluate IBD-discordant (IBDD) qCNVs to determine the cause of discordance.

**Fig 1.**
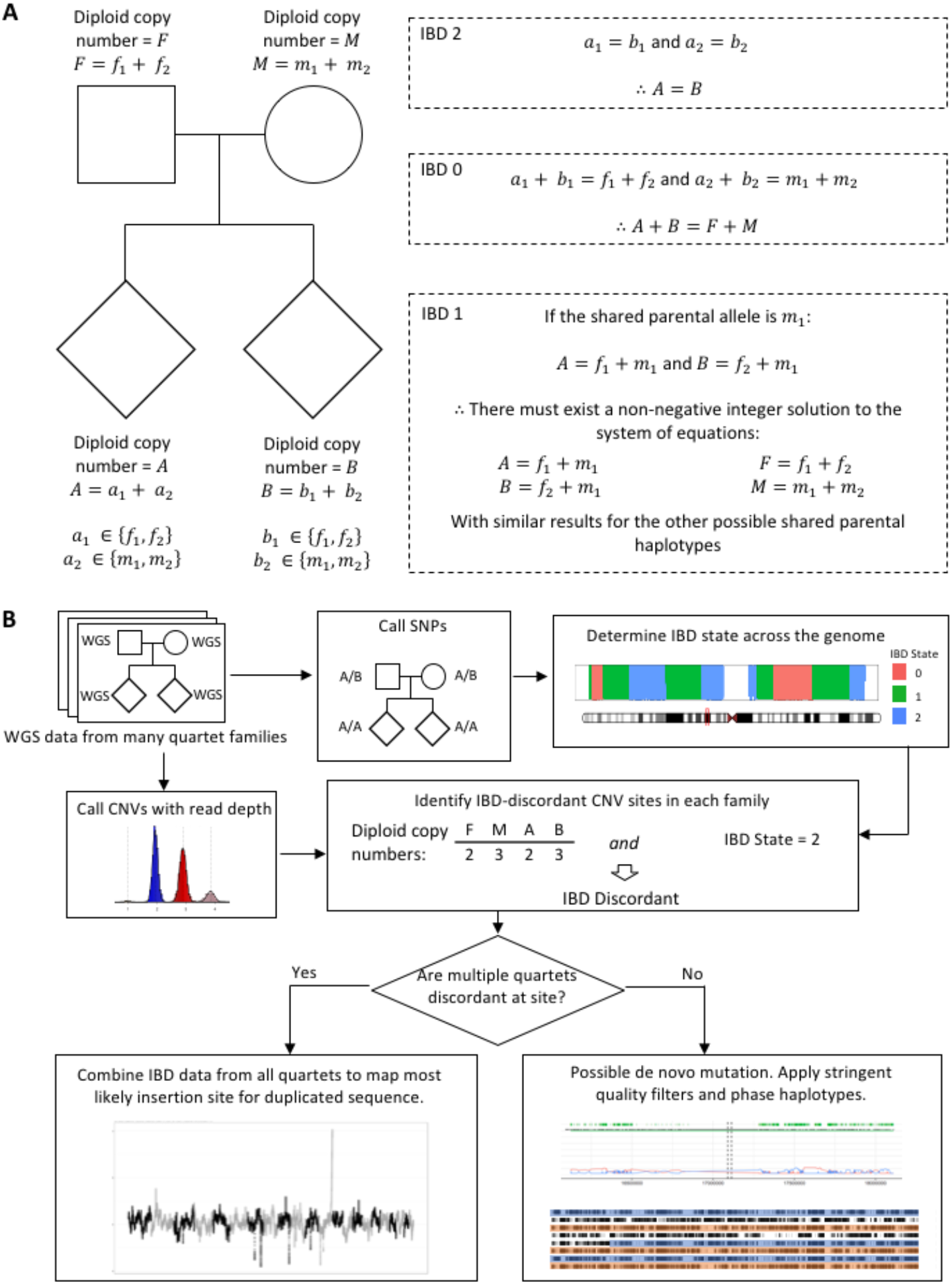
Identifying genomic loci at which sibling identity-by-descent (IBD) and copy number measurements are inconsistent. **(A)** The level of IBD sharing between two siblings at a locus prescribes a mathematical relationship among the copy number states in their family for that locus. At IBD-2 sites, the diploid copy numbers (DCNs) of the two siblings must be equal. At IBD-0 sites, the sum of the siblings’ DCNs must be equal to the sum of their parents’ DCNs. At IBD-1 sites, there must be an integer solution to a system of equations that models the haploid copy numbers with a shared parental allele. We identify CNVs for which copy-number measurements violate these mathematical relationships. (**B)** Analysis workflow. We begin by calling SNP genotypes and copy number states at CNV sites. Based on the SNP genotypes we determine the IBD state of each pair of siblings at every locus in the genome. We then examine the copy number state of each quartet at each CNV site (qCNVs) to determine whether it is concordant with the sibling IBD state. If many families have IBD-discordant (IBDD) qCNVs, we combine IBD information from all quartets to determine the most likely insertion site for the duplication in the genome. If only one or a small number of families have IBDD qCNVs at the site, we further evaluate whether the discordance is due to a *de novo* mutation.

Our methods identified a median of 167 IBD blocks, with each block corresponding to a specific number of parental haplotypes shared (IBD-0, IBD-1, or IBD-2), per sibling pair. These represent 72 autosomal recombinations per child in the cohort, after removing blocks beginning or ending at telomeres (**S1 Fig**). IBD block sizes had median and mean lengths of 9.8MB and 16.1MB, respectively. (Although our IBD analysis method does not explicitly distinguish paternal from maternal sharing of IBD-1 blocks, we found that this could be resolved in most cases given SNP data from the parents: among those IBD-1 blocks with at least 50 SNP sites that were informative for parental haplotype sharing, 97% had agreement among at least 95% of the informative sites about the parent of origin of the shared haplotype (**S2 Fig**).)

To precisely measure integer copy number at CNV loci, we used methods described in Handsaker et al. and implemented in the Genome STRiP software package. We used three different pipelines based on Genome STRiP algorithms to identify CNV sites (more detail in **Methods**). The first pipeline discovers CNV sites based on changes in the read depth signal at uniquely mappable regions of the genome; in our cohorts, this detected a median of 1,722 copy-number-variant sites, with deletions and mixed sites of at least 1Kb and duplications of at least 1.5Kb in length per sample (**S3 Fig**), prior to merging calls together across the cohort. The second pipeline measures read depth within pairs of annotated segmental duplications on the reference genome (**Methods**). Finally, we looked for retroposed pseudogenes by copy number genotyping sets of intervals corresponding to the exonic regions of all annotated transcripts. For each method, we merged all calls into a single call set and cross-genotyped every call in each of our cohorts.

As described above, by examining the integer copy numbers of all members of a quartet at a CNV site, we can determine whether or not the copy number state of the quartet is *concordant* or *discordant* with the called IBD state (**Fig 1A**). Wherever a qCNV is discordant with IBD state (“IBD-discordant”, or IBDD), the discordance must arise from one of the following possibilities: (i) an error in calling the copy number of one or more of the family members; (ii) an error in determining the IBD state at the locus; (iii) the CNV could be a dispersed duplication in which the duplicated copies are inserted at a location in the genome where the siblings are in a different IBD state; or (iv) the IBD discordance could be due to a *de novo* mutation. In the following sections we explore the latter two possibilities while attempting to control for or model sources of error.

## Dispersed Duplications

For CNV loci at which multiple quartets had copy-number measurements that were IBDD, we tested the possibility that the discordances were due to the presence of a dispersed duplication. We developed a method to localize the dispersed insertion based on linkage mapping, drawing upon the IBD information and copy number measurements from all members of the 763 pairs of siblings and their parents in the data set. Briefly, we evaluate every possible location on the autosomes as a potential insertion site and determine the log odds (LOD) of observing the copy numbers of all samples for the CNV given their IBD states at that location as compared to the original CNV site. We calculated likelihoods using a probabilistic model that accounts for potential errors in copy number measurement and IBD state calling. By normalizing and integrating LOD scores at each location, we were able to compute an estimated 95% confidence region (CR) – analogous to a linkage peak – for the genomic location at which the duplicate sequence is located. **Fig 2** shows an example highlighting a 2.6kbp duplication in an intronic region of the *NEGR1* (neuronal growth regulator 1) gene. We conducted this analysis for duplications observed at a single interval in the reference genome, duplications of repeated sequence represented by segmental duplications in the reference genome, and sets of intervals representing cDNA sequences of annotated transcripts – the latter being an attempt to find insertions of retroposed pseudogenes. Our method identified 95 dispersed duplications for which an insertion location (distinct from the source locus) was confidently predicted by IBD analysis (**S1 Table**; plots similar to **Fig 2** for each dispersed duplication are given in **S1 Appendix**). In addition, for two dispersed duplications for which our method could not find a confident insertion location prediction in the autosome, we noticed that copy number status correlated with having a Y chromosome, and were thus able to predict that their duplicate segments are inserted on the Y chromosome.

**Fig 2.**
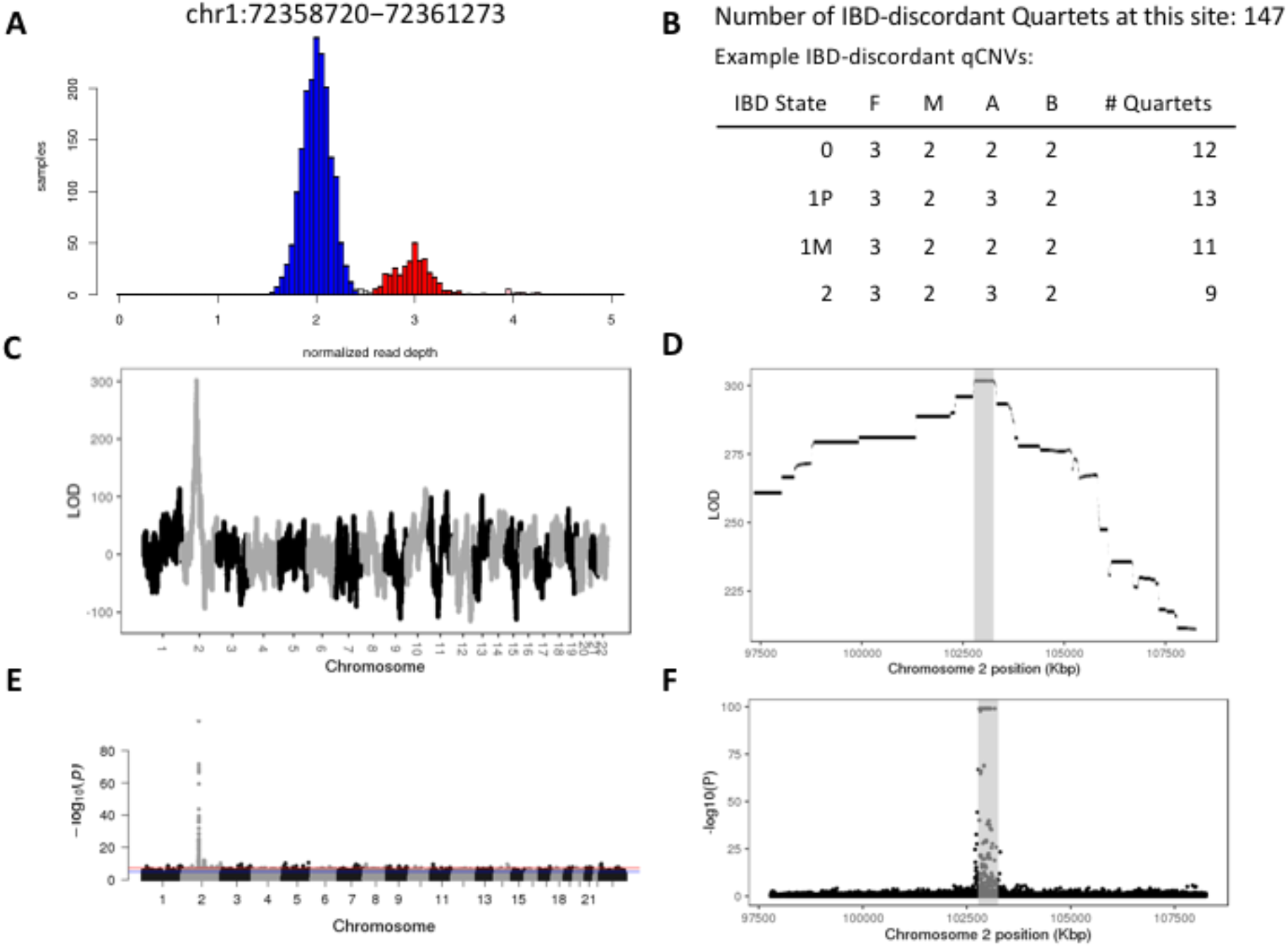
Localization of a dispersed duplication by linkage and association analysis. **(A)** Distribution of copy number measurements from 2076 samples for a 2.6kb duplication annotated as residing within an intron of *NEGR1*. (**B)** Copy number distributions of example IBD-discordant quartets at this site. 147 quartets exhibited IBDD copy number measurements at this locus. The most frequent configuration of copy numbers (qCNVs) within discordant quartets in each IBD state is shown. **(C)** LOD scores, plotted across the entire genome, of the likelihood of each tested segment containing the insertion of the duplicated sequence. **(D)** Zoomed-in plot of the data from **(C)** in a 10Mbp window centered on the 95% confidence interval of the insertion locus predicted from the LOD scores. **(E)** Manhattan plot of association between SNP genotype and copy number state in the parents. **(F)** Data from **(E)** zoomed-in to the same 10Mbp window shown in **(D)** shows that the linkage and association analyses are concordant, and that the association analysis further localizes the insertion. All coordinates refer to GRCh37.

Linkage disequilibrium provides an additional source of information for localizing dispersed duplications [4]. We conducted LD mapping for our identified dispersed duplications by performing a series of genome-wide association mappings using the copy number of each variant as the associated trait (**Fig 2D** and **2E**) within a subset of parent samples from our cohort. Of 95 dispersed duplications analyzed, we found that 84 of the events had an LD peak within the 95% confidence interval for the insertion location computed by IBD-based localization; an additional 4 had a peak within the 99% confidence interval. For 26 of these 84 sites with LD peaks within their IBD-based mapping confidence intervals, this was the only genome-wide significant association peak. The remaining 58 of these sites had more than one LD peak — potentially reflecting mismapped SNPs — illustrating cases in which IBD-based mapping provided uniquely useful mapping information. Of these, many had additional peaks of association near the annotated CNV site. Although we masked markers within the duplicated segment which we were studying to prevent paralog-specific variants (PSVs) on the inserted copy from providing a spurious LD signal, it is possible that the actual duplicated region included additional sequence that is paralogous to the surrounding region that might harbor PSVs. The remaining 11 sites did not have an LD peak that agreed with their IBD-based mapping.

64 of the regions duplicated by dispersed duplication events in our call set intersected with annotated genes. These included 6 retroposed pseudogenes in which a gene’s exons (but not its introns) were duplicated. 65 of the duplicated regions intersected annotated segmental duplications, and 6 of those were identified only by analyzing copy numbers across both regions of a segmental duplication. Allele frequencies of the dispersed duplications ranged from 1% to events for which almost every genome appeared to have at least one dispersed copy of the duplicated segment. 31 of the 95 events were interchromosomal, involving the migration of genomic sequences to other chromosomes; in the rest the dispersed duplication was localized to the same chromosome as the duplicated segment. We observed relatively little overlap with previous lists of dispersed duplications; only 6 of our variants matched events described by Conrad *et al.* [4] which to the best of our knowledge is the largest earlier catalogue of dispersed duplications based on population-scale data. The low overlap between the two sets of results can be explained by differences in the data types — the prior study used SNP genotyping array data rather than WGS — and in the analytical approaches: as we describe below, the LD-based mapping approach used in the prior study is complementary to our IBD-based method and can be used to confirm its results, but it identifies different sets of duplications when used as the primary mapping technique. The six retroposed pseudogenes were fewer in number than those discovered in previous studies that used read pair and split read mappings [21,22], indicating that methods that look for breakpoints (rather than measure the copy number of genomic segments) may provide a more sensitive screen for these events, likely due to the non-repetitive nature of the duplicated exonic sequence.

Given the high rate of overlap of dispersed duplications with annotated segmental duplications, we speculated that segmental duplications may play an important role in dispersal. For 31 of the 65 events for which the observed CNV region overlapped with one paralog of an SD, at least 50% of the 95% confidence region (CR) for the insertion location of the duplicate sequence also lay within segmental duplications. Further investigation revealed that in 37 of the original 65 events, the 95% CR for the duplication locus at least partially overlapped an annotated segmental duplication that was paralogous to the source locus. This leads us to hypothesize a mechanism for the formation of events that appear as dispersed duplications in read depth-based analyses, but are the result of polymorphic deletions in one copy of a segmentally duplicated region (**Fig 3**). For these loci (about 40% of the total), the dispersed duplication is an older event that may well be fixed (monomorphic) in human populations; polymorphism arises from a subsequent deletion within one of the paralogs. If the human reference genome is constructed from individuals who carry the haplotype with the deletion (or if the duplicated nature of the sequence is unrecognized during genome assembly), sequencing data from individuals who carry one or more haplotypes without the deletion appear to indicate a duplication relative to the reference genome.

**Fig 3.**
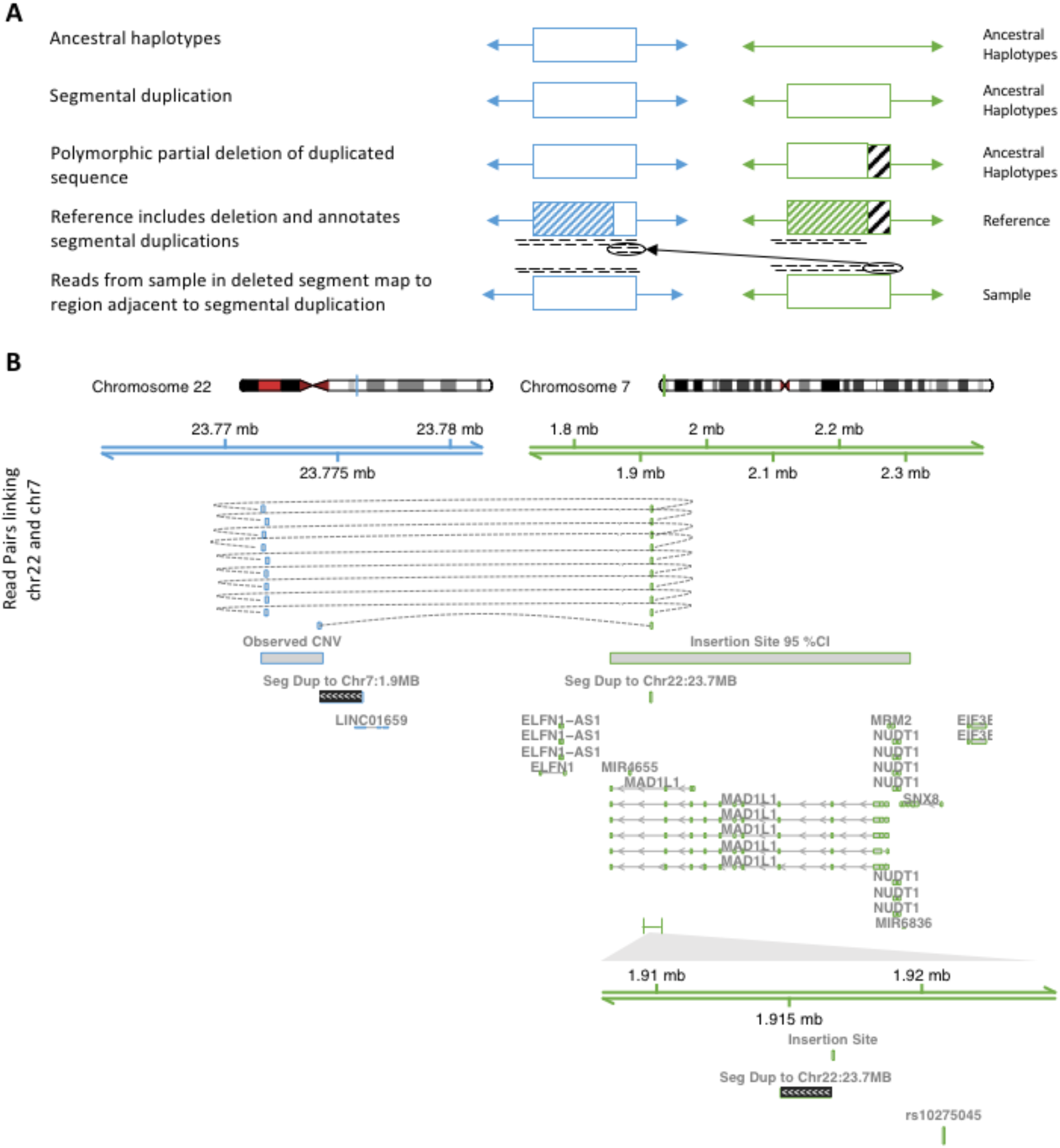
Dispersed duplications caused by segmental duplication (SD) followed by partial deletion. **(A)** A model of dispersed duplication formation through SD and deletion. First, a segment on an ancestral haplotype is duplicated to another locus. In a later generation, part of one of the duplicate segments is deleted; the deletion remains polymorphic in the population. The human genome reference is built from a haplotype that carries the deletion; a region smaller than the original duplication is annotated as the SD. Finally, when a genome which does not carry the deletion is sequenced, reads from the region matching the deleted segment map to the area adjacent to the annotated SD. **(B)** An example of an observed duplication that matches this pattern. The dispersed duplication is located on chromosome 22, flanking an annotated SD with chromosome 7. In this case, we observe read pairs with mates mapped to chromosome 7, allowing us to pinpoint the exact insertion location of the duplication. Read pairs from a randomly chosen sample are depicted. The bounds of the duplication and the 95% confidence interval for the insertion location based on IBD mapping are highlighted. Gene annotations and segmental duplications in the region of the CNV and near the insertion location at the *MAD1L1* locus on chromosome 7 are shown. **C)** A close up of the insertion region as identified by read pair mapping. The insertion lies adjacent to the other side of the SD from chromosome 22, as predicted by the model in **(A)**. This insertion is in strong LD (*R*^*2*^ = 0.93) with a nearby SNP associated with schizophrenia, rs10275045. All coordinates refer to GRCh37.

A few dispersed duplications have been previously characterized and served as positive controls. For example, our method identified the duplication of a large segment that includes *DUSP22* from chromosome 6 to a centromeric region of chromosome 16. A duplication between these two regions was one of the earliest observed dispersed duplications in humans [23], and has been further elucidated by multiple subsequent studies [1,24–26]. Our method also identified a portion of the well-characterized segmental duplication of the *HYDIN* locus from chromosome 16 to chromosome 1 [27,28].

In addition to these known large events, some of the dispersed duplications we identified occurred in linked clusters, with a group of multiple nearby source CNVs for which IBD mapping predicted insertion locations clustered nearby one another in a distal genomic location. In such cases it was difficult to determine whether the clustered CNVs represented a single large event or multiple similar events; therefore we have left them as separate events in this analysis. For example, we detected seven duplicated segments in a span of over 1MB along 1q21.2. While the IBD analysis-based insertion region confidence interval for these events is large (**S4 Fig**), it includes 1p11.2, corresponding to a previously described segmental duplication that includes the *NBPF* (neuroblastoma breakpoint family) genes [29]. This segmental duplication is represented by new sequence in the GRCh38 reference. Because our study was conducted with data aligned to GRCh37, we observe a duplication, whereas in GRCh38-aligned data one would observe a deletion of the new sequence; the presence of the event in this scan highlights that it is polymorphic in copy number. In this region, the correlations between the copy numbers observed in different samples at different sites in the cluster is high (**S5 Fig**), indicating that these duplications are mostly part of the same event. In contrast, we also detected a cluster of dispersed duplications in annotated segmental duplications in 8p23.1. IBD mapping placed their insertion locations approximately 4MB away (**S6 Fig**), in a region on the other side of a polymorphic 4.2MB inversion which is known to harbor copy number variability associated with the beta defensin gene family [30]. In this cluster of CNVs, correlation between copy numbers at different sites is lower (**S7 Fig**), indicating that there may be multiple dispersed duplication events carried on different haplotypes at this region. An additional cluster of dispersed duplications can be found linking the segmental duplications on 16p13 and 16p12 which flank the *XYLT1* locus, and another cluster is located within SDs in 17p11.2 which are associated with the CNVs which cause Smith-Magenis and Potocki-Lupski syndromes [31].

We found 3 cases in which a dispersed duplication polymorphism was in LD with a genome-wide significant GWAS SNP [32] at the dispersed duplication’s predicted destination locus (**Table 1**). (All three of the correlated SNPs were within 200kb of the maximum LOD score IBD mapping site for the dispersed duplication predicted by our methods, supporting the accuracy of the mapping approach.)

**Table 1:** Dispersed duplications in LD with significant GWAS SNPs.

| Disease or Trait | GWAS index SNP ID and location | CNV Source Location (s) | Dispersed Duplication Location (MAX LOD score) | LD (R<sup>2</sup>) of dispersed duplication to GWAS index SNP | Genes Near Source CNV | Genes Near Duplication |
| --- | --- | --- | --- | --- | --- | --- |
| Lipoprotein (a) - cholesterol levels [33] | rs1620921<br>6:161,197,087 | 6:153,742,231-153,747,450<br>6:153,748,600-153,752,000 | 6:160,998,148-160,999,191 | 0.516 |  | *LPA*,<br>*PLG* |
| Autism spectrum disorder, attention deficit-hyperactivity disorder, bipolar disorder, major depressive disorder, and schizophrenia (combined) [34] | rs8321<br>6:30,032,522 | 12:133,085,650-133,088,265 | 6:29,958,526-29,958,531 | 0.964 | *FBRSL1* | *HLA-H*<br>*HLA-G*<br>*HLA-J* |
| Schizophrenia or bipolar disorder [35] | rs10275045<br>7:1,920,826 | 22:23,771,600-23,774,339 | 7:1,886,333-1,886,388 | 0.934 |  | *MAD1L1* |

All coordinates refer to GRCh37.

Two of the linkages between GWAS SNPs and dispersed duplications are most likely due to the presence of long LD blocks in the regions in which they occur. For example, in the HLA region on chromosome 6, the LD between the duplication of sequence from chromosome 12 and a GWAS SNP associated with psychiatric diseases including ASD and schizophrenia is likely simply due to the presence of both alleles on a long haplotype. Similarly, in an event that duplicates sequence from 6q25 to 6q26, annotations of the duplicated region show that much of the duplicated sequence content is made up of human endogenous retrovirus, and therefore matches a known HERV insertion in this region [36]. The insertion site is upstream of the *LPA* and *PLG* genes, and likely tags a haplotype containing *LPA* variation that is causal for changes in lipoprotein(a) levels.

Of more potential specificity is the duplication from chromosome 22 to chromosome 7, for which copy number is strongly correlated with the index SNP for one of the strongest GWAS signals in schizophrenia (**Fig 3**). This duplication, previously identified by Conrad et al. [4], is inserted into an intronic region of *MAD1L1*, adjacent to an annotated segmental duplication between 7:1,914,665-1,916,576 and 22:23,774,213-23,776,121. The SNP with which it is in high LD, rs10275045, is approximately 5kb upstream on chromosome 22; this SNP is also listed as an eQTL for the gene *FTSJ2* in GTEx v7 [37]. The duplicated segment on chromosome 22 contains a ChromHMM-annotated strong enhancer in the K562 cell line, according to data generated by the ENCODE project [38], raising the possibility that the duplicated segment may contain regulatory elements affecting the expression of genes at its destination locus, or alternatively that the haplotype represented in the reference genome represents the deletion of a set of regulatory elements.

Based on the observation that many of the identified dispersed duplications overlap with annotated regulatory elements, H3K27Ac marks, or DNase hypersensitivity clusters, we tested whether copy number of dispersed duplications associated with changes in gene expression using publicly available data. To do so, for each dispersed duplication we identified the annotated eQTL from GTEx v7 with which it was in highest LD. For each such CNV-eQTL pair, we determined the copy number state and SNP genotype of each GTEx sample, and then re-analyzed the eQTL’s expression data using copy number status of the duplication as a covariate in the model. We found 166 instances where the copy number of the CNV achieved nominal significance in a tissue data set for a given gene even after controlling for the genotype of the previously reported eQTL. 22 different dispersed duplications passed this threshold for at least one tissue-gene pair. The top results are summarized in **Table 2**, and the full set is presented in **S2 Table**.

**Table 2.** Top associations between dispersed duplication copy number and gene expression after controlling for previously reported eQTL genotype.

| CNV Location | eQTL | R<sup>2</sup> | Gene | Tissue | p-value of copy number in joint model |
| --- | --- | --- | --- | --- | --- |
| 1:149,232,622-149,239,750 | 1:148,546,082 | 0.182 | *HIST2H2BB* | Transformed Fibroblasts | 1.67E-28 |
| 1:149,297,554-149,315,000 | 1:148,546,082 | 0.176 | *FCGR1C* | Lung | 3.41E-17 |
| 2:113,045,432-113,049,031 | 2:87,123,114 | 0.916 | *PLGLB1* | Lung | 5.88E-04 |
| 3:195,399,263-195,403,622 | 3:195,661,089 | 0.378 | *AC024937.4* | Esophagus Mucosa | 5.83E-13 |
| 5:177,217,820-177,221,664 | 5:175,469,223 | 0.857 | *RP11-826N14.2* | Testis | 4.18E-03 |
| 7:5,940,936-5,951,376 | 7:6,774,135 | 0.155 | *GRID2IP* | Caudate Basal Ganglia | 2.18E-02 |
| 7:64,551,729-64,554,250 | 7:55,803,970 | 0.876 | *NUPR1L* | Esophagus Mucosa | 1.05E-02 |
| 7:66,438,701-66,441,000 | 7:72,332,773 | 0.656 | *TRIM50* | Artery (Tibial) | 3.30E-02 |
| 8:7,211,500-7,227,500 | 8:11,985,833 | 0.452 | *RP11-351I21.11* | Skin (Sun exposed) | 2.33E-22 |
| 8:7,746,183-7,750,670 | 8:11,985,833 | 0.347 | *RP11-351I21.11* | Skin (Sun exposed) | 8.03E-16 |
| 8:7,773,888-7,791,373 | 8:12,002,588 | 0.413 | *FAM86B2* | Artery (Aorta) | 4.32E-07 |
| 8:12,235,600-12,248,700 | 8:7,744,937 | 0.287 | *FAM66E* | Brain (Cortex) | 2.05E-04 |
| 12:131,792,472-131,796,941 | 12:132,163,328 | 0.884 | *RP11-495K9.7* | Testis | 3.04E-02 |
| 16:16,629,916-16,637,000 | 16:18,776,381 | 0.267 | *NOMO2* | Brain (Cerebellar) | 1.03E-05 |
| 16:18,615,000-18,635,104 | 16:18,776,381 | 0.120 | *NOMO2* | Brain (Cerebellar) | 4.81E-05 |
| 17:16,656,700-16,709,271 | 17:18,477,409 | 0.984 | *CTD-2303H24.2* | Brain (Hippocampus) | 7.83E-03 |
| 17:16,710,600-16,725,700 | 17:18,477,409 | 0.738 | *KRT16P4* | Testis | 5.22E-05 |
| 17:16,744,849-16,747,500 | 17:18,477,409 | 0.752 | *KRT16P1* | Testis | 1.79E-03 |
| 17:18,624,874-18,633,247 | 17:15,525,044 | 0.428 | *AC005517.3* | Esophagus Muscularis | 2.07E-06 |
| 17:43,655,745-43,662,300 | 17:44,410,368 | 0.618 | *ARL17A* | Skin (Sun exposed) | 1.38E-26 |
| 17:62,915,465-62,918,489 | 17:44,410,368 | 0.662 | *ARL17A* | Skin (Sun exposed) | 2.55E-28 |

P-values were computed by fitting a joint linear model with GTEx covariates and the genotypes of a correlated eQTL SNP as well as CNV genotypes. For each dispersed duplication with a nominally significant result in at least one tissue, the most significant result is reported. *HIST2H2BB* lies within the duplication event for the first two CNVs; the other genes listed here lie outside of the duplicated regions. All coordinates refer to GRCh37.

Dispersed duplications could in principle confound many kinds of genetic analysis, particularly those looking for *trans* effects of quantitative trait loci (QTLs). For example, multiple studies [39,40] have described *trans* expression-QTLs in which SNPs on chromosome 16 associate with expression of the *DUSP22* gene on chromosome 22; this effect may be more parsimoniously explained by LD between these SNPs and the extra (dispersed) copy of *DUSP22* near them on chromosome 16. In another example, a recent survey of methylation QTLs [41] includes 10 *trans* mQTLs that match dispersed duplications in our catalog, in that they can be explained by the insertion of a paralogous copy of the segment containing the target CpG site near the candidate mQTL.

### *De Novo* Mutations

Another potential cause of IBD-CN discordances are *de novo* mutations that occurred within specific families. *De novo* copy number mutations have been profoundly important in genetic studies of neurodevelopmental disorders, but to date their analysis has been largely limited to unique (single-copy) regions of the human genome. We attempted to use IBD-discordancy to identify additional kinds of *de novo* structural mutations. To produce a conservative list of mutation calls, we developed a set of stringent filters and further curated the remaining loci (see **Methods**). Our filters removed loci at which IBDD qCNVs could potentially have arisen from imprecision in copy-number measurement, or from errors in IBD inference. In addition, since the other possible source of IBD discordancy is a dispersed duplication, and our method for identifying dispersed duplications depended on observing the event in a large number of families, we also removed loci with IBDD qCNVs that were observed in a small number of families and in which the copy-number states observed in the family members were compatible with a dispersed duplication event. Among 519 quartets on which we performed this analysis, 41 candidate *de novo* mutations satisfied all of these criteria.

In comparing our findings with those of earlier analytical approaches, we were able to make use of data from three recent and well-validated studies of de novo structural variation events in a subset of the samples used in this study [8–10]. We merged the mutation call sets from these three studies to produce a list of 223 *de novo* SV events. Of these, 68 were non-somatic CNV events of over 1kb on autosomes.

These studies used methods which relied heavily on more traditional SV detection techniques, including the analysis of paired end and split read mappings, and were tuned for sensitivity to smaller events in regions in which CNVs are rare. Our method, in contrast, is designed to detect mutations in regions of common copy number variation that are more difficult to analyze using these approaches. We found that 15 of our filtered CNV mutation calls matched events in this callset; the remainder were heavily enriched for common and multiallelic CNV sites (see below). A list of the 26 remaining mutations not called in the other studies is shown in **Table 3**, with more detail presented in **S3 Table**.

**Table 3.** *De novo* CNV mutations that were not identified in previous analyses of WGS data from the same families [8–10].

| Coordinates | Length (kb) | Type | Genes |
| --- | --- | --- | --- |
| 1:104,161,928-104,210,498 | 48.6 | DEL | *AMY1A, AMY2A* |
| 1:248,645,267-248,647,958 | 2.7 | DUP |  |
| 2:125,462,004-125,465,328 | 3.3 | DUP | *CNTNAP5* |
| 2:220,745,641-220,748,604 | 3.0 | DUP |  |
| 4:159,393,426-159,394,857 | 1.4 | DEL | *RXFP1* |
| 5:697,000-785,600 | 88.6 | DUP | *ZDHHC11, ZDHHC11B* |
| 6:71,975,109-71,977,227 | 2.1 | DUP |  |
| 7:39,821,774-39,823,243 | 1.5 | DEL | *LINC00265* |
| 7:143,950,000-144,050,000 | 100.0 | DUP | *OR2A7, CTAGE8, ARHGEF35, OR2A9P, OR2A1, OR2A20P* |
| 7:144,013,000-144,035,000 | 22.0 | DUP | *OR2A1* |
| 10:81,460,000-81,615,000 | 155.0 | DEL | *FAM22B, FAM22E* |
| 13:52,852,408-52,857,516 | 5.1 | DUP | *THSD1P1, TPTE2P2* |
| 15:34,710,000-34,853,500 | 143.5 | DUP | *GOLGA8A, GOLGA8B* |
| 15:34,710,000-34,853,500 | 143.5 | DUP | *GOLGA8A, GOLGA8B* |
| 15:43,850,000-43,959,000 | 109.0 | DEL | *PPIP5K1, CKMT1B, CKMT1A, STRC, CATSPER2* |
| 16:28,612,500-28,626,300 | 13.8 | DEL | *SULT1A1* |
| 16:74,395,000-74,462,000 | 67.0 | DEL | *NPIPL2, CLEC18B* |
| 17:18,990,000-19,100,100 | 110.1 | DEL | *GRAPL, KYNUP1* |
| 18:14,801,000-14,812,500 | 11.5 | DEL | *ANKRD30B* |
| 18:63,200,000-63,207,000 | 7.0 | DUP |  |
| 19:1,950,289-1,951,768 | 1.5 | DUP | *CSNK1G2* |
| 19:2,346,851-2,350,546 | 3.7 | DEL | *SPPL2B* |
| 19:40,367,999-40,384,255 | 16.3 | DUP | *FCGBP* |
| 19:43,827,760-43,833,148 | 5.4 | DUP |  |
| 22:20,332,500-20,660,000 | 327.5 | DEL | *RIMPB3* |
| 22:24,281,500-24,343,000 | 61.5 | Complex | *GSTT2B, DDTL, DDT, GSTT2, GSTTP1* |

All coordinates refer to GRCh37.

Of the *de novo* structural mutations revealed by our analysis, 29/41 mutations overlapped genes (71%); 21 of those were from the 26 mutations revealed only by our method. Many of the de novo mutations overlapped segmental duplications (22/41; 54%); this is particularly true of the sites that were uniquely called by our method (19/26; 73%). In contrast, the merged call set from the three previous studies contained only six events that overlapped segmental duplications (9%), three of which were also detected by our approach. 18 of the 26 calls unique to our method were at sites with multiallelic copy number variation (mCNVs); all but two of these also overlapped segmental duplications. 5 calls were at sites where the majority of samples in the cohort have a copy number greater than the reference, 17 were at sites where alternate copy numbers are common (MAF>10%), and 13 were at sites at which both deletion and duplication alleles are present in the population. By contrast, none of the sites in our call set that were also detected in the comparison call sets have any of these properties. Of the 53 calls from the merged comparison call set that were not made by our method, 11 occurred at sites or in samples that were removed from our analysis by our QC filters; for the remaining 42 sites our input CNV call sets did not contain a called site at the locus. These results indicate that our approach is able to call mutation events at regions inaccessible to other methods, but that it depends heavily on the input CNV call sets, which had not been tuned for sensitivity to rare events as was the case in the other studies.

Several of the de novo structural mutations occurred in structurally complex genomic regions that have been of interest in disease studies. For example, we detected a large de novo mutation of the amylase gene locus (**Fig 4**), a genomic region with many structural alleles and complex structures whose potential influence on obesity and BMI has been debated [42–44]. Earlier work has suggested, based on the low levels of LD of AMY-locus structural alleles to nearby SNPs, that specific structural alleles may have arisen recurrently on multiple SNP haplotypes [42]. By phasing the SNP haplotypes and copy-number measurements at this region in this family, we infer that the mutation was a deletion of 288kb – likely mediated by non-allelic homologous recombination – that caused one of the common, previously characterized structural haplotypes at this locus (AH7-3) to mutate into a much shorter form that corresponds in structure to a different common allele (AH1) (**Fig 4D**). This provides direct evidence that recurring mutation generates the complex relationships of structural alleles to SNP haplotypes previously documented for the *AMY* locus [42]; such mutational processes are also likely to give rise to the recurring mutations suggested by haplotypes at the complement component 4 (C4) locus [45].

**Fig 4.**
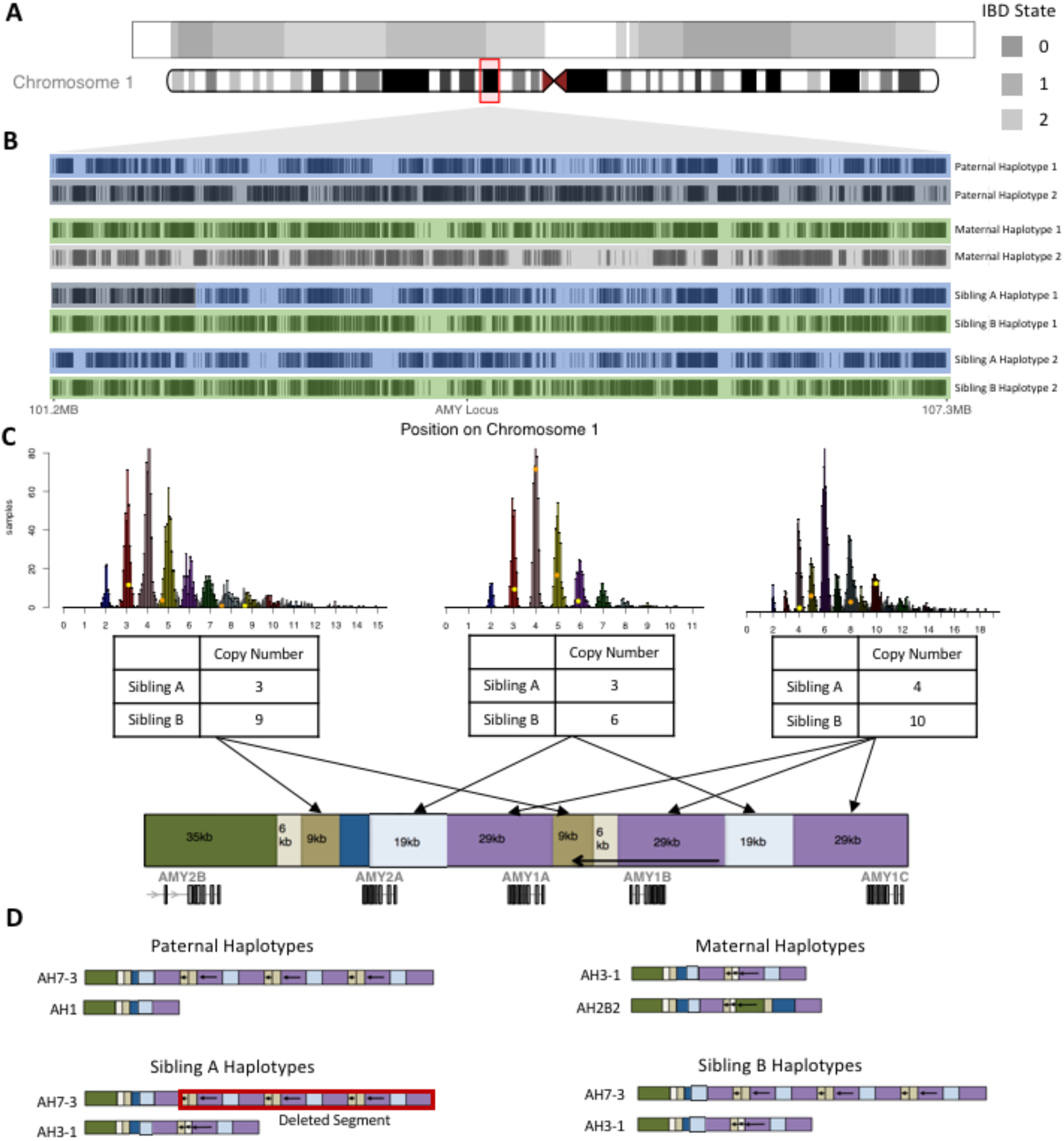
A *de novo* structural mutation at the amylase (*AMY1A/AMY1B/AMY1C, AMY2A/AMY2B*) gene locus on chromosome 1. **(A)** SNPs where IBD state was confidently (phred-scaled quality >= 20) predicted along chromosome 1 are shown for the siblings in a quartet family. **(B)** Inferred SNP haplotypes for family members in a 6-Mbp genomic segment containing the AMY locus. All SNPs informative for IBD state within a 6.13Mbp window centered on the AMY locus are shown. A dark vertical bar indicates that the phased haplotype carries the alternate allele at that site. Position on the Y axis represents order of the SNPs used for the plot rather than genomic coordinates. The paternal and maternal haplotypes transmitted at the AMY locus are highlighted in blue and green; the non-transmitted parental haplotypes are shaded in light and dark gray. Although Sibling A’s paternal chromosome carries a recombination event 3Mbp distal to the *AMY* locus, the two siblings are confidently inferred to be IBD-2 at the *AMY* locus. **(C)** Normalized read depth across three regions within the *AMY* locus for 2,076 samples. The members of the family discussed above are highlighted, with orange points representing the parents and yellow representing the children. Note that despite the siblings’ IBD-2 state for the chromosomal segment containing the AMY genes, measurement of copy number for various genomic segments at the locus are highly discordant between the siblings (e.g. 3 vs. 9 copies of the segment shown on the left) and are extremely surprising given the quantitative resolution/precision of the analysis. The hg19 *AMY* reference haplotype is shown below, with paralogous sequences depicted as blocks of the same color. Read depth of coverage was independently measured for each of the three sequences as indicated by arrows. **(D)** A model of the structural haplotypes and mutation in the family that fits the observed copy number differences. The model is based on the haplotypes described by Usher et al. [42]. In this model, a deletion in Sibling A has caused a haplotype which carried the AH7-3 structure to mutate into a structure that resembles the AH1 structure, with a single copy (instead of 7 copies) of the *AMY1* gene. The mutation likely occurred by non-allelic homologous recombination between paralogous segments shown in purple.

We also detected *de novo* mutations in two different ASD probands at the same 144 kb segment on chromosome 15, between the *GOLGA8A* and *GOLGA8B* genes. An analysis of paralog-specific variants at the locus confirmed that the IBDD qCNVs were indeed due to *de novo* duplications; in both instances a paternal copy of the segment was duplicated in a proband affected by ASD (**S8-9 Fig**). *GOLGA8* core duplicons have been shown to contribute to chromosome 15q13.3 microdeletions – a known cause of ASD, epilepsy and intellectual disability – and to have participated in evolutionary structural rearrangements in humans and apes [46]. The detection of two different *de novo* mutations (among 1,038 offspring) at a single *GOLGA8*-flanked locus suggests that the mutation rate at this locus is appreciable. This locus appears to carry a common deletion allele (9.2% allele frequency in our cohorts) and a rare duplication allele (0.5%).

We also detected an example of complex structural rearrangement involving a deletion and duplication at a site of common copy number variation containing the genes *GSTT2* and *DDTL* (**S10 Fig**). The glutathione S-transferase genes *GSTT1* and *GSTT2*, which exhibit exuberant and common copy number variation, have been the subject of at least 28 genomic studies of their potential associations to drug metabolism and other phenotypes (**S4 Table**).

Combining the set of *de novo* CNV events observed in our analysis of the ASD cohort with the largely complementary sets of CNV calls from the three previous studies [8–10] produced a combined call set containing 94 *de novo* structural mutations, ascertained in 519 quartets with 1,038 total offspring. We thus estimate that *de novo* CNVs of at least 1kb in size arise in 1 per 22 meioses. Of the 94 events, 55 occurred in unaffected siblings of ASD probands, indicating that mutational events of this size (1+ kb) were not significantly more common in subjects with ASD, and that our mutation rate estimate is likely not elevated by the presence of ASD patients in the cohort.

## Discussion

We described an approach for combining IBD analysis in families with precise copy-number measurements to identify dispersed duplications and *de novo* structural mutations of complex sequence. For both dispersed duplications and mutations, our methods appear to be able to access forms of CNV that are inaccessible to standard structural variation analyses based on discordant read pair or split read mappings of short-read Illumina data. In particular, we detect many events in regions of the genome with common and/or multiallelic copy number variation. While many of these events may eventually be detected with long read or linked read sequencing, the ability to detect them in abundant short-read WGS data will allow them to be more conclusively evaluated for relationships to phenotypes.

Many events may have eluded our mapping approach, such that the numbers and frequencies we report should be considered a lower bound. Our approach for identifying and localizing dispersed duplications depend on observing multiple confidently genotyped CNV events at a locus in quartets that have a recombination between the source locus and the location of the insertion (assuming intra-chromosomal events, which make up the majority of those found by this study). Therefore, we are more likely to detect events in regions of the genome with higher recombination rates. Our detection and mapping methods are also likely to be successful only for events present at allele frequencies>~1% in a cohort of 706 families, and were conducted in a sample of families of primarily European ancestry; larger analyses of more families and additional populations will almost certainly uncover and map many more such dispersed duplications.

Some dispersed duplications appear to reflect ancient (perhaps fixed) dispersed duplications for which one genomic copy was subsequently deleted (**Fig 3**). At such loci, it may be beneficial to add the deleted paralog to the human genome reference in order to avoid the misalignment of sequences to its mate, which can cause errors in variant calling, phasing, and other downstream analyses in those regions.

Awareness of dispersed duplications is important for avoiding confounding in association studies and other genome-wide analyses. For example, consider the case of a dispersed duplication in the autosome with the dispersed segment included on the Y chromosome. If PSVs from the dispersed segment are called as SNPs in the autosomal interval, they can generate false associations for phenotypes (such as ASD) with sex-biased incidence. It is therefore important to filter SNPs that are the result of dispersed duplication CNV PSVs from most association analyses. In addition, dispersed variants can cause the false appearance of genomic *trans* effects, as we have highlighted in eQTL and methylation studies.

Highly copy-number-variable regions appear to be a substantial contributor to the overall CNV mutation rate, comprising some 22 of the 94 structural mutations larger than 1kbp that have now been observed in the widely studied families from the Simons Simplex Collection of the Simons Foundation Autism Research Initiative. We note that we almost certainly underestimate the frequency of events in such regions given that that (i) we exclude regions for which copy-number inference is imprecise (e.g. VNTR regions with still-higher copy numbers), and (ii) we exclude events that are so close to a parental haplotype recombination event that we cannot be sure of the exact location of the crossover. We are therefore likely missing many events caused by non-allelic homologous recombination (NAHR) between homologous parental chromosomes, although we may be detecting NAHR events caused by crossing over between sister chromatids in the process of sister chromatid exchange [47].

These results indicate that IBD can provide valuable additional information in structural variation analyses. For example, confidently called IBD-2 blocks provide a wealth of “truth data” that can be used to critically evaluate the calls made by variant calling methods, by evaluating concordance between calls made on sibling-pairs in their IBD-2 regions. We find IBD-guided quality control and validation to be useful in novel methods and pipeline development.

## Methods

### Data used

We analyzed copy number variation in quartet families from three separate cohorts. The first was a collection of families affected by immune deficiency disorders (Cohort “ID”). Usable samples from this cohort included 35 fully sequenced quartets, 5 quintets (consisting of both parents and three children), and 1 sextet (both parents and four children), for a total of 171 total sequenced samples and 56 distinct sibling pairs. Parents and unaffected siblings in this cohort were sequenced to a mean of 30X coverage, while the affected siblings were sequenced to a depth of 60X. The second cohort was composed of 146 quartet or larger families in which a single child exhibited schizophrenia, including 146 quartets, 18 quintets, and 2 sextets, for a total of 689 samples and 188 sibling pairs (Cohort “SCZ”). All samples in this cohort were sequenced to a mean depth of 30X. The final cohort included in the analysis was the Simons Simplex Collection (SSC) made available by the Simons Foundation Autism Research Initiative (SFARI; Cohort “ASD”). This collection consists of quartet families in which one child is affected with autism spectrum disorder, and includes 519 quartets, for a total of 2076 samples sequenced. Combined, the three cohorts include WGS data from 2936 samples, making up 706 families and 763 distinct sibling pairs. All data was aligned to GRCh37 using BWA-MEM. SNPs and indels were called using the GATK best-practices joint calling pipeline [48] independently within each cohort.

### Calling IBD state in sibling pairs

We implemented an HMM-based method for determining the IBD state between siblings across the genome given SNP genotypes of the entire quartet. First, we categorized the possible IBD states present at each biallelic SNP site given the observed genotypes of the quartet as follows:

| Parent1 | Parent2 | Sib1 | Sib2 | Possible IBD States |
| --- | --- | --- | --- | --- |
| A/B | A/B | A/A | A/A | 2 |
| A/B | A/B | A/A | B/B | 0 |
| A/B | A/B | A/B | A/B | 0,2 |
| A/B | A/B | A/B | A/A | 1 |
| A/B | A/A | A/B | A/B | 1,2 |
| A/B | A/A | A/A | A/A | 1,2 |
| A/B | A/A | A/B | A/A | 0,1 |
| A/A | A/A | A/A | A/A | 0,1,2 |

We then analyzed biallelic SNP variants in each quartet using an HMM with hidden states for IBD-0, IBD-1, and IBD-2, setting the observed value at each locus to be the set of possible IBD states given the SNP genotypes in the quartet as described in the table above. For each state we allowed a 3% error probability of observing an incompatible configuration, and divided the remaining emission probability evenly among the valid observation states. In the case of IBD-1, we observed that a common genotyping error mode was for all samples to be heterozygous, and so we allocated a 2% emission probability to that state, dividing up the remaining 1% error probability among the two remaining incompatible observations. Empirically, we found that our algorithm performed best when it was strongly discouraged from switching state, and therefore we set transition probabilities to arbitrarily low values of 10^−16^ for valid recombination possibilities and 10^−18^ for impossible state transitions; despite those values being implausibly low from a biological standpoint, the model was robust to changes in those parameters. Initial state probabilities were distributed uniformly.

Our approach is similar to that taken by ISCA [18,19] with several differences: ISCA allows for the existence of “non-mendelian” states that are inconsistent with mendelian transmission rules for SNPs. Their model add states for ‘compression blocks’, which model CNVs with multiple paralogs not represented in the reference which lead to an overabundance of sites that appear to be heterozygous in all members of the family, and a state that models other forms of mendelian inheritance errors. Rather than explicitly model these error states, we prefer to stringently filter the SNP sites that we use for IBD state determination. We therefore masked all sites that: fell outside the 1000 Genomes Phase 1 strict analysis mask; fell inside a low complexity region; occurred in the 1000 Genomes Hardy-Weinberg failure mask; or overlapped with a set of regions with excess coverage in the 1000 Genomes phase three data set [49]. In addition, given a CNV call set, our tool masks SNP sites on a per sample basis in regions where that sample has been determined to have a copy number different from the reference ploidy.

Our tool emits the most likely boundaries of IBD state blocks for each sibling pair, as computed by executing the Viterbi algorithm on our HMM. In addition, it can optionally compute forward and backward probabilities over the HMM and annotate each SNP site in the VCF with the posterior likelihoods that the quartet being analyzed is in each possible IBD state. While we do not explicitly model IBD-1P and IBD-1M as separate states in the HMM, we track the number of SNP sites that agree with only paternal or maternal inheritance in IBD-1 segments. We then assign paternal or maternal shared inheritance status to IBD-1 segments depending on whether there were more markers that disagree with the former or the latter.

### CNV calling and QC

We used Genome STRiP’s read-depth based CNV calling pipeline to discover CNVs and compute diploid copy numbers in each of the three cohorts separately. Genome STRiP version numbers used for each call set were r1659 for the ID cohort, r1685 for the SCZ cohort, and r1609 for the ASD cohort. The differing versions should have no impact on the output of CNV calling. The ASD cohort was divided into nine separate batches and CNVs were discovered in each independently. Sites discovered independently in different batches were then de-duplicated using the following procedure: first, every site discovered in any batch was copy-number genotyped in all samples. Sites were then annotated using Genome STRiP’s RedundancyAnnotator, and were kept if the duplicate score was set to zero or the site was never listed as preferred to another site with zero discordant genotypes.

In the SCZ cohort, we found that many samples had a much greater than expected number of called CNV variants and failed to give clean estimates of copy number dosage on the X and Y chromosomes. We removed samples that failed a contamination analysis based on allelic depth at common SNPs, as well as samples that had over 19,000 2kb windows for which we estimated a diploid copy number other than two in stage 5 of the Genome STRiP CNV pipeline. Considering only intact families the size of a quartet or greater after filtering, this left 454 samples in 106 quartets and 6 quintets.

To call copy number in paired segmental duplications, we used Genome STRiP’s segmental duplication pipeline [1]. This pipeline estimates read depth at locations that have identical sequence between the paired paralogs that make up the SD to call total copy number across the segmental duplication.

Finally, we produced a last set of copy number calls to try to identify retroposed pseudogene events, in which we measured copy number only on the exonic regions of transcripts. To accomplish this using Genome STRiP’s CNV genotyping pipeline, we created a series of genome masks, one for each transcript in the ENSEMBL transcript database, which masked out all introns in the transcript. We then estimated copy number over the interval spanned by the transcript to produce an estimate of exonic copy number.

For each gene, we chose the transcript copy number estimates that had the greatest separation between estimated copy number clusters. This method proved to have limited utility, only recovering 6 non-redundant dispersed duplication calls, possibly because our current read depth collection methods are not optimized for the short intervals defined by exons; refinement to this method could produce much better results.

### Identification of IBD-Discordant CNVs

Given the IBD state segmentation of the genome and a set of CNV calls that include accurate estimates of diploid copy number in each variable segment, it is possible to identify a set of CNV calls within a quartet that are discordant with the predicted IBD state. First, we exclude all quartet and site combinations where IBD segments were not explicitly called at the locus or not all of the members of the quartet had a confident diploid copy number call. Then, we let *CN*(*S*_1_),*CN*(*S*_2_), *CN(P)*, and *CN(M)* be the observed diploid copy numbers at a given CNV site for the first sibling, second sibling, father, and mother in a quartet, respectively. The CNV is classified as discordant with the IBD state if any of the following conditions hold:

- The IBD state is IBD-0 and *CN*(*S*_1_) + *CN*(*S*_2_) ≠ *CN*(*P*) + *CN*(*M*)
- The IBD state is IBD-1P and there is no integer solution to the system of equations (with a corresponding set of equations for sites in state IBD-1M:

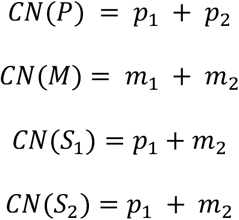
- The IBD state is IBD-2 and *CN*(*S*_1_) ≠ *CN*(*S*_2_)

For any given CNV locus and quartet we used the likelihoods that the sample has a given copy number at that locus (provided by Genome STRiP’s read depth genotyping module [1]) to determine the total likelihood that quartet is discordant at the locus by summing over all possible CNV states for the quartet:

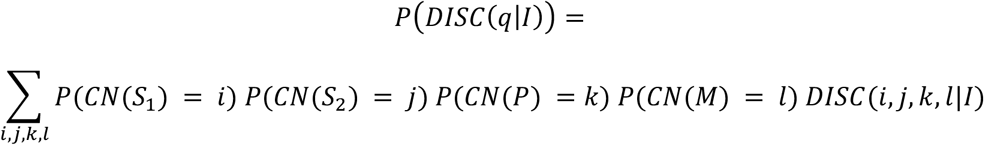

Where *i, j, k*, and *l* are the possible copy number states of each sample, *I* is the inferred IBD state at the locus, and *DISC*(*i*, *j*, *k*, *l*|*I*) is a function which returns zero or one according to the concordance rules listed above given the quartet copy numbers and the chosen IBD state.

For segmental duplication CNVs, which link two separate genomic regions together, we follow the same rules, but in most cases only evaluate IBD-concordancy at segmental duplications where both paralogs of the duplication are in the same IBD state. See the section “Detection of Mutations in Segmental Duplication Paralogs” for an exception.

### Localization of dispersed duplication copies via IBD signal

We attempt to find the most likely locations for the insertion points of dispersed duplications observed across many families as follows. We divided the genome into segments bounded by the SNP markers that were used for IBD segment determination, and for each segment *i*, quartet *q*, and IBD state *I*, store the posterior probability that *IBD*_*q,i*_ = *I*, which we computed using the Forward-Backward algorithm on the IBD state HMM. Since our IBD state detection algorithm does not separately model IBD-1P and IBD-1M but uses a single state to represent IBD1, we approximate the posterior probabilities of the sub-states representing paternal or maternal inheritance. To do so, we use the ratio of SNPs that support paternal (*P*) or maternal (*M*) inheritance in the called IBD-1 segment *s*, adding a pseudocount to each value:

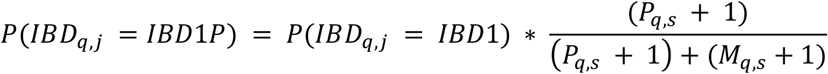

We can then find the likelihood that the duplicate copies of the CNV are actually inserted at segment *i* by computing the value:

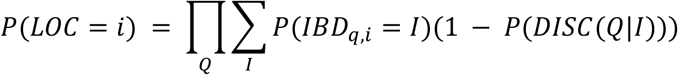

Where *Q* is the set of all quartets in the analysis. We then compute the log odds ratio or LOD score

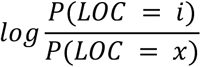

Where *x* is the source locus for the CNV. We found independent peaks in the distribution of LOD scores across the genome using the peak-finding algorithm from the `cardidates` R package [50] and considered alternate loci which have LOD scores greater than 10 and *log*_*10*_ likelihoods in the new location *i* of at least −100 as potential dispersed duplication insertion sites.

### Filtering of dispersed duplication candidates

Using the methods described above, we classified all called quartet genotypes at every CNV site called by Genome STRiP as concordant or discordant. To remove sites with high error rates in copy number genotyping, we filtered out sites with a CNQUAL score of 30 or less, a call rate less than 90% in any of the three sample cohorts, or a cluster separation score (GSCLUSTERSEP) less than 5 in the ASD cohort. After classifying the copy numbers at all remaining CNV sites, we excluded 13 quartets which had over 300 discordant CNV loci with a CNQUAL score of 60 or greater from further analysis. We then ran the localization procedure described above on all CNV sites at which at least two quartets were IBD-discordant and at least 10 quartets were *IBD-informative* at the site, with *IBD-informative* being defined as having a set of diploid copy numbers that would be discordant with at least one IBD state: even if informative quartets are concordant with their called IBD state, they can be useful in localizing the dispersed duplication. After running IBD-mapping, we removed sites with ambiguous mapping results by eliminating those for which the value of the second highest LOD score peak was within 10 of the highest scoring peak. We then attempted to deduplicate sites representing the same copy number variant event by collapsing clusters of sites with start and end coordinates within 25kb of each other and identical distal locations for the highest LOD mapping score. After filtering the remaining clustered sites for LOD scores of at least 10, we were left with 232 candidate sites. We then manually curated this set by removing sites with incorrect copy number genotyping (determined by manually inspecting read depth histograms and binned depth profiles at each site), deduplicating overlapping sites with similar copy number allele profiles, and manually re-segmenting the boundaries of these sites by visually inspecting binned read depth profiles in the surrounding areas and re-genotyping the updated segments.

For mapping of segmental duplication CNVs we followed a similar procedure, omitting manual deduplication and re-segmenting, and manually curating candidate sites based on read-depth genotyping plots to ensure that the majority of samples were genotyped correctly.

### Estimation of dispersed duplication frequency in multi-allelic sites

For multi-allelic CNVs which we determined to be dispersed duplications, the problem of determining whether all or some of the copy number variant alleles were dispersed remained. We attempted to resolve this by systematically excluding samples which clearly carried particular CNV alleles and re-running the localization procedure described above. If either re-analysis failed to confirm the existence and inferred insertion site of the dispersed duplication, the allele was excluded from the reported frequency for the dispersed duplication. For example, for a site at which the mode diploid copy number in the cohort was 4, but some samples were measured at copy number 3 and some at copy number 5, we re-analyzed the cohort twice, first excluding samples at copy number 3, and then excluding samples at copy number 5. In this example, if the reanalysis excluding the samples at copy number 3 did not confirm the existence of the dispersed duplication or agree with its predicted localization, we concluded that the deletion allele implied by the copy number 3 measurements was not actually the absence of a common dispersed allele, and reported only the estimated frequency of the duplication allele. Conversely, if the reanalysis excluding the samples carrying the extra duplication allele confirmed the initial analysis, we concluded that it was actually a dispersed copy which was “deleted”, and therefore that all samples in the cohort carried at least one dispersed copy of the original segment.

### Localization of dispersed duplications with linkage disequilibrium

To determine localization mappings using linkage disequilibrium (LD), we extracted the SNP genotypes and copy number calls of each dispersed duplication for the parents within the ASD cohort. We then imported the SNP genotypes into PLINK [51] and performed the following filtering and QC steps: removal of sites with call rates less than 85%; removal of sites with a minor allele fraction less than 0.5%; and pruning of sites with pairwise LD of greater than 0.5. We then ran a principal components analysis and used the top four principal components as covariates in the following analyses. For each candidate dispersed duplication, we treated the called copy number for each sample as a quantitative phenotype. We then ran a genome-wide association analysis for the dispersed duplication copy number phenotype and plotted the results for both the genome-wide association and for the SNPs within 10MB of the insertion site for the dispersed duplication predicted from the IBD mapping. For each individual association analysis we masked out SNPs that fall within the boundaries of the called CNV.

### Association with GWAS SNPS

We filtered the hg19 GWAS catalog and removed SNPs with a reported association p-value of more than 5E-8. We extracted the genotypes of all parent samples from the quartets in our datasets at those SNP loci and converted their genotypes into dosages of zero, one, or two, excluding sites at which>100 samples did not have a called genotype for the SNP. We then extracted the called integer diploid copy numbers of all predicted dispersed duplication sites for those samples and computed the correlation coefficient of SNP dosage and copy number for all pairs of SNPs and CNV sites.

### Gene expression analysis

We downloaded all eGenes identified by GTEx v7 [37] in all tissues and filtered the results to gene-SNP pairs with a *q*-value of less than 0.05 in any tissue. We then determined the eGene SNP with the highest correlation between SNP dosage and copy number in our samples (subset to the ASD cohort). Because the only sample level data for GTEx samples available to us was from the GTEx v8 release, which is mapped to version hg38 of the human reference genome, we determined SNP genotypes and dispersed duplication copy number in GTEx samples by first lifting over to the hg38 reference the coordinates of each dispersed duplication and eGene SNP identified above. We then extracted SNP genotypes from the GTEx v8 WGS VCF and used Genome STRiP to re-genotype copy number variants at the lifted-over dispersed duplication intervals in GTEx v8 WGS data. For each such eGene-dispersed duplication pair, we then re-computed the eQTL analysis for that SNP in each tissue for which the eGene was reported to be significant in GTEx v7, using the GTEx v7 expression matrix and covariates for that tissue. To do so, we fit three models:

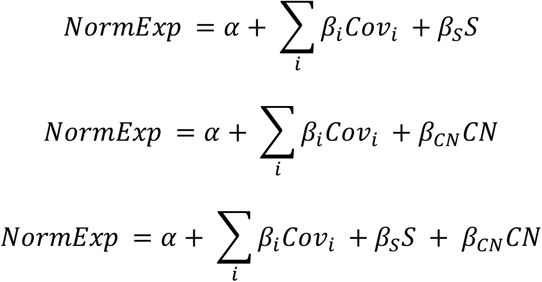

Where *NormExp* is the normalized expression reported by GTEx v7 for that tissue, *Cov*_*i*_ are the GTEx covariates for that tissue, S is the vector of SNP dosages for the eGene SNP, and CN is the vector of copy numbers of the dispersed duplication CNV the GTEx samples. We looked for eGenes in which p-value of *β*_*CN*_ was nominally significant in the joint model.

### Candidate mutation filtering

To produce a list of CNV mutations based on IBD discordancy, we applied a stringent set of filters and manual curation. We began by producing a list of the annotated copy state and IBD status of each quartet in our entire input cohort at every candidate CNV location. First, we excluded all qCNVs at which either the copy number of every family member could not be confidently called or we could not assign an IBD status due to conditions such as a missing IBD state call for the siblings or a partial overlap of the IBD block with the CNV interval. To further ensure that all copy number states were called correctly, we then excluded any qCNVs for which the copy number call quality score for any individual of the quartet was below 40. Next, we filtered out qCNVs which were within 250,000bp of an IBD state change for the quartet, as well as those within IBD blocks within which less than 99% of the SNP markers agreed with the final called IBD state of the block. We then removed all qCNVs at sites which overlapped with any of the dispersed duplications identified in our previous analyses, as well as those at sites which fell within regions of size at least 1MB within the mask that we used to filter out SNPs for IBD analysis. Next, we removed qCNVs from quartets that contained any samples that were flagged in the CNV discovery process as containing a genome-wide excess of depth variability in 2kb windows. We further filtered out qCNVs from families that appeared to be outliers for the number of discordant qCNVs detected at this point in the analysis by removing those families where the number of discordant qCNVs was greater than the median count plus two times the median absolute deviation of the count across all families. We then attempted to remove problematic CNV sites by filtering out all qCNVs from sites at which more than 1% of the qCNVs remaining for the site were IBD discordant.

At this point we subset the remaining qCNVs to those that came from the ASD cohort and had a discordant IDB-CNV status. In this remaining set of discordant qCNVs, we first attempted to merge overlapping and/or fragmented sites which represented the same CNV event by sorting the qCNVs by family and then by genomic coordinates and then joining together pairs of qCNVs at which the start of a site was within 10kb of the second site and the copy numbers called for each family member were consistent. We then turned to manual curation of the remaining mutation candidates. We generated plots showing the histogram of normalized read depth estimates for the cohort at the site, the depth profile in bins tiled across the site, and a visualization of the SNP markers and IBD state posterior probabilities at each marker in the neighborhood of the site (see **S11 Fig** for an example). We then manually curated the remaining sites, removing candidates for the following reasons:

- Poor copy number calls, usually the result of incorrect boundaries for the discovered CNV interval. If a called CNV interval overlaps regions where a sample is in different copy number states, the estimate of total copy number for the sample will be incorrect. We also removed all duplication calls of less than 2500bp due to an observed high rate of erroneous copy number genotyping.
- Poor IBD state calls. We removed qCNVs that had insufficient informative SNP markers between the nearest recombination site and the CNV interval. We also removed any sites for which the Phred-scaled posterior probability of the called IBD state at the closest left- and right-flanking informative SNP markers was less than 10.
- Potential somatic events, at sites where the normalized depth estimate for the samples in the family consistently fell far away from an integer value.
- Potential rare dispersed duplication events, where either multiple discordant qCNVs remained across the cohort, or the observed copy state within a single family was inconsistent with a simple CNV in one of the offspring, and the qCNVs were consistent with a rare dispersed duplication allele.
- Other sites with multiple discordant qCNVs, indicating that the site was problematic for copy number genotyping.

In the remaining set, for sites at multi-allelic CNVs, we conducted an LD analysis for total copy number, as we did for dispersed duplications, to capture further instances of potential rare dispersed duplications that might be tagged by SNPs in other parts of the genome. Finally, we examined the allelic fraction of paralog-specific variants within the CNV interval. In several instances we used this to determine whether the mutation event was a gain in one sibling or a loss in the other, and we excluded sites that contained PSVs with allele fractions incompatible with the proposed IBD state and a simple gain or loss in one of the siblings. An overview of the filtering steps described above and how many qCNVs were filtered at each stage is given in **S5 Table**.

### Detection of mutations in segmental duplication paralogs

As described above, we evaluated the IBD concordancy of copy number state within pairs of segmental duplications by limiting our analysis to pairs of segmental duplications at which both siblings in the quartet shared the same copy number state. To detect and characterize mutations, however, we conducted an additional analysis to determine which paralog in each SD pair was varying in copy number, similar to previous analyses [1]. To do this, we calculated the estimate of copy number using only uniquely mappable loci within each paralog of the SD. Using the copy number likelihoods for each of these paralogs, we limited our analysis to quartets in which the most likely copy number state for one of the paralogs was two for all members of the family, and the other paralog had a copy number estimate other than two for at least one family member. We then plotted these estimates against the copy number estimate derived from the SD as a pair, which uses only positions in the interval with identical sequence between the two paralogs. In cases where it was clear that only one paralog varies in copy number (**S13 Fig**), we conducted our mutation analysis using the IBD state of the quartet at the varying paralog. This produced one additional mutation call not detected by the previous methods, and allowed us to clarify the nature and interval of previously detected mutation candidates.

## Data and Code Availability

Data from the Simons Simplex Collection was obtained through SFARI Base. Approved researchers can obtain the SSC population dataset described in this study (https://www.sfari.org/resource/simons-simplex-collection/) by applying at https://base.sfari.org. The schizophrenia cohort data is in the process of being submitted to an access-controlled public database; this manuscript will be updated with the location and accession number when they become available. Because the immune deficiency cohort data was generated for a research project on immune-deficiency syndromes, it will be published in an access-controlled public database concurrently with that study when it is complete.

The tools used to compute IBD state between siblings, annotate Genome STRiP copy number calls with IBD discordancy information, and localize dispersed duplications are available at https://github.com/cwhelan/quartetibdanalysis.

## Acknowledgements

C.W.W. was funded through NIH R01 HG006855 and U01 MH105641. R.E.H. and S.K. were funded through NIH R01 HG006855 and U01 MH105641. G.G. was funded through NIH R01 HG006855 and U01 MH115727. We are grateful to all of the families that participated in each of the research studies used in this analysis, including those at the participating Simons Simplex Collection (SSC) sites, as well as the principal investigators of that collection (A. Beaudet, R. Bernier, J. Constantino, E. Cook, E. Fombonne, D. Geschwind, R. Goin-Kochel, E. Hanson, D. Grice, A. Klin, D. Ledbetter, C. Lord, C. Martin, D. Martin, R. Maxim, J. Miles, O. Ousley, K. Pelphrey, B. Peterson, J. Piggot, C. Saulnier, M. State, W. Stone, J. Sutcliffe, C. Walsh, Z. Warren, E. Wijsman). We appreciate access to genetic and phenotypic data on SFARI Base. The SCZ cohort samples used in this study were recruited in the Netherlands by the Dutch GROUP Investigators and in the UK by Michael O’Donovan and Michael Owen at the MRC Centre for Neuropsychiatric Genetics and Genomics at Cardiff University and by their collaborators. They were curated by Elliott Rees in Cardiff. We thank them for providing the data for this analysis. Collection and analysis of the immune deficiency cohort was supported in part by the Division of Intramural Research of the National Institute of Allergy and Infectious Diseases. Additional support for sequencing and analysis of the immune deficiency cohort, and for J.H. and J.M., was provided by Merck Research Laboratories, Merck & Co., Boston, MA. Christina Usher provided valuable feedback on the writing of the manuscript. We would also like to thank members of the Broad Institute Data Sciences Platform Methods Group and the BroadSV working group for helpful conversations and feedback, especially Ryan Collins, Harrison Brand, and Michael Talkowski.

## Supporting Information

**S1 Fig. Number of recombinations detected per sibling pair, broken down by cohort used in this study.**

**S2 Fig. Within IBD-1 blocks from all sibling pairs, the ratio of SNP sites supporting a paternal origin of the shared haplotype.**

**S3 Fig. Number of copy number variant sites called in each sample during the first round of CNV calling.**

**S4 Fig. A cluster of dispersed duplications detected in 1q21.** The segment or segments at which variable copy number was detected are plotted in black; the 95% confidence regions for the locations of the dispersed segments are plotted in gray.

**S5 Fig. Correlation of copy number events detected in 1q21.** Pairwise scatter plots of sample copy numbers in the cluster of CNV events detected in 1q21. In this case the variable segments exhibit high correlation, indicating that they are mostly participating in a single large copy number polymorphism.

**S6 Fig. A cluster of dispersed duplications observed at the edges of the polymorphic inversion in 8p23.** The segment or segments at which variable copy number was detected are plotted in black; the 95% confidence regions for the locations of the dispersed segments are plotted in gray.

**S7 Fig. Correlation of copy number events detected in the 8q32 cluster.** Pairwise scatter plots of sample copy numbers in the cluster of CNV events detected in 8p23. The measured copy numbers of pairs of segments have low correlations, indicating that they are capturing different dispersed duplication polymorphism events.

**S8 Fig. Paralog specific variant analysis of the first of two mutations detected at the** *GOLGA8A/B* **locus.** See **S9 Fig** for the other mutation. In this family, the IBD state between siblings at this locus is IBD-0, and the diploid copy numbers are father: 1; mother: 2; sibling 1: 3; sibling 2: 1. Plotted are allele fractions of SNP variants present only in the father’s copy of the segment on the *y* axis, and position on chromosome 15 (within the CNV region) on the *x* axis. Although the father had only a single copy of the segment at this genomic region, the clustering of allele fractions at 66% in the sibling with copy number 3 indicates that two copies of the father’s segment were transmitted to that child −- and therefore a *de novo* duplication of the segment appeared in the father.

**S9 Fig. Paralog specific variant analysis of the second of two mutations detected at the** *GOLGA8A/B* **locus.** See **S8 Fig** for the other mutation. In this family, the IBD state between siblings at this locus is zero, and the diploid copy numbers are father: 2; mother: 1; sibling 1: 3; sibling 2: 1. Plotted on the *y*-axis are the allele fractions of SNP alleles which were found in sibling 1 that were present in only one, but not both, of the parents, and were not found in sibling 2. Position on chromosome 15 (within the CNV region) is plotted on the *x*-axis. Variants which were present only in the father cluster at 66% allele fraction, while variants that were specific to the mother cluster at 33% allele fraction. This indicates that sibling 1 received one copy of the mother’s segment at this locus, and two copies of one of the father’s segments, which was duplicated *de novo*.

**S10 Fig. A complex rearrangement at the** *DDTL* **/** *GSTT2* **locus.** Normalized read depth profiles for the ASD cohort are shown across the locus, where each horizontal line represents the normalized read depth in a single sample in each binned genomic location. Profiles for the two siblings in which the mutation was detected are highlighted as follows: Sibling 1: Red; Sibling 2: Yellow. The siblings are IBD-2 at the region. Divergence of the two sibling’s profiles shows that Sibling 1 has undergone a complex rearrangement involving a duplication of the sequence present at approximate coordinates 24,282,000-24,300,000 on chr22 (highlighted in blue) and a deletion of a segment at approximate coordinates 24,324,000-24,343,000 (highlighted in green).

**S11 Fig. A visualization of the IBD state as supported by SNP markers in the vicinity of a potential CNV mutation.** The top tracks show the location of SNPs with various characteristics used by the IBD calculation algorithm to determine IBD state in the vicinity of a copy number variant -- in this case within segmental duplications at the *AMY* locus, indicated by dotted vertical lines. Top two tracks: MATERNAL and PATERNAL indicate whether the SNP supports inheritance of a maternal or paternal haplotype given that the IBD state is 1. The next six tracks highlight which SNPs could support which IBD states. For example, a SNP marked ONE_OR_TWO could support either IBD-1 or IBD-2 status. Below these all SNPs are shown, colored according to the IBD state with the maximum posterior probability, as well as the mask that was used to filter out SNPs for the purposes of IBD state calculation. At the bottom of the plot, the log10 posterior likelihoods of each IBD state at each SNP are shown. In this case, the quartet is in IBD 2 state at the center of the plot, with a transition to IBD-1M 1.5 Mb upstream.

**S12 Fig. Copy number variation at paralogs of segmental duplications.** Data for two different pairs of annotated segmental duplications are shown. Within each pair, we plot the estimated copy number computed by measuring read depth in only the upstream copy of the SD (Paralog A), the downstream copy (Paralog B), and the sum of the two paralog copy number estimates, on the *y*-axis. On the *x*-axis we plot the estimate of the copy number for the pair of paralogs, based on read depth at pairs of loci that have identical sequence content in both paralogs. In each of these cases it is clear that Paralog A is varying in copy number across the cohort and not Paralog B, so we used only the IBD state for Paralog A in mutation analysis. For each of these examples, the copy number estimates for the four members of a quartet determined to have a mutation are highlighted.

**S1 Table. Dispersed Duplications.**

**S2 Table. Dispersed duplications with significant effects on gene expression.**

**S3 Table. Additional information for** *de novo* **CNV mutations that were not identified in previous analyses of WGS data from the same families**[8–10].Additional information is given about the events listed in Table 3.

**S4 Table. A list of studies associating the copy number of the** *GSTT1* **and** *GSTT2* **genes with various phenotypes.**

**S5 Table. Filtering process for identifying CNV mutations.** We began by considering the copy number state for every family in our input cohorts at every CNV site detected in our input CNV set. We call the grouping of the copy number state of a quartet and the IBD state at a CNV site a qCNV. Beginning with 95.6 million qCNVs, we applied a series of automatic filters and manual curation to identify a set of 41 mutation calls.

**S1 Appendix. Characteristics and localization plots for all dispersed duplications identified in this study.**

## Notes

### Competing Interest Statement

The authors have declared no competing interest.

### Summary of Updates

Fixed formatting issues in the manuscript; added citations to Wong et al. (1990) and Sebat et al. (2004).

## Citations

1. Handsaker RE, Van Doren V, Berman JR, Genovese G, Kashin S, Boettger LM, et al. Large multiallelic copy number variations in humans. Nat Genet. 2015;47: 296–303.

2. Pickeral OK, Makałowski W, Boguski MS, Boeke JD. Frequent human genomic DNA transduction driven by LINE-1 retrotransposition. Genome Res. 2000;10: 411–415.

3. Zhang Y, Li S, Abyzov A, Gerstein MB. Landscape and variation of novel retroduplications in 26 human populations. PLoS Comput Biol. 2017;13: e1005567.

4. Conrad DF, Pinto D, Redon R, Feuk L, Gokcumen O, Zhang Y, et al. Origins and functional impact of copy number variation in the human genome. Nature. 2010;464: 704–712.

5. Kim S, Medvedev P, Paton TA, Bafna V. Reprever: resolving low-copy duplicated sequences using template driven assembly. Nucleic Acids Res. 2013;41: e128.

6. Itsara A, Wu H, Smith JD, Nickerson DA, Romieu I, London SJ, et al. De novo rates and selection of large copy number variation. Genome Res. 2010;20: 1469–1481.

7. Malhotra D, McCarthy S, Michaelson JJ, Vacic V, Burdick KE, Yoon S, et al. High frequencies of de novo CNVs in bipolar disorder and schizophrenia. Neuron. 2011;72: 951–963.

8. Turner TN, Coe BP, Dickel DE, Hoekzema K, Nelson BJ, Zody MC, et al. Genomic Patterns of De Novo Mutation in Simplex Autism. Cell. 2017;171: 710–722.e12.

9. Brandler WM, Antaki D, Gujral M, Kleiber ML, Whitney J, Maile MS, et al. Paternally inherited cis-regulatory structural variants are associated with autism. Science. 2018;360: 327–331.

10. Werling DM, Brand H, An J-Y, Stone MR, Zhu L, Glessner JT, et al. An analytical framework for whole-genome sequence association studies and its implications for autism spectrum disorder. Nat Genet. 2018;50: 727–736.

11. Palta P, Kaplinski L, Nagirnaja L, Veidenberg A, Möls M, Nelis M, et al. Haplotype phasing and inheritance of copy number variants in nuclear families. PLoS One. 2015;10: e0122713.

12. Kong A, Frigge ML, Masson G, Besenbacher S, Sulem P, Magnusson G, et al. Rate of de novo mutations and the importance of father’s age to disease risk. Nature. 2012;488: 471–475.

13. Francioli LC, Polak PP, Koren A, Menelaou A, Chun S, Renkens I, et al. Genome-wide patterns and properties of de novo mutations in humans. Nat Genet. 2015;47: 822–826.

14. McCarroll SA, Hadnott TN, Perry GH, Sabeti PC, Zody MC, Barrett JC, et al. Common deletion polymorphisms in the human genome. Nat Genet. 2006;38: 86–92.

15. Conrad DF, Andrews TD, Carter NP, Hurles ME, Pritchard JK. A high-resolution survey of deletion polymorphism in the human genome. Nat Genet. 2006;38: 75–81.

16. Donis-Keller H, Green P, Helms C, Cartinhour S, Weiffenbach B, Stephens K, et al. A genetic linkage map of the human genome. Cell. 1987;51: 319–337.

17. Coop G, Wen X, Ober C, Pritchard JK, Przeworski M. High-resolution mapping of crossovers reveals extensive variation in fine-scale recombination patterns among humans. Science. 2008;319: 1395–1398.

18. Roach JC, Glusman G, Smit AFA, Huff CD, Hubley R, Shannon PT, et al. Analysis of Genetic Inheritance in a Family Quartet by Whole-Genome Sequencing. Science. 2010;328: 636–639.

19. Roach JC, Glusman G, Hubley R, Montsaroff SZ, Holloway AK, Mauldin DE, et al. Chromosomal haplotypes by genetic phasing of human families. Am J Hum Genet. 2011;89: 382–397.

20. Dewey FE, Chen R, Cordero SP, Ormond KE, Caleshu C, Karczewski KJ, et al. Phased whole-genome genetic risk in a family quartet using a major allele reference sequence. PLoS Genet. 2011;7: e1002280.

21. Abyzov A, Iskow R, Gokcumen O, Radke DW, Balasubramanian S, Pei B, et al. Analysis of variable retroduplications in human populations suggests coupling of retrotransposition to cell division. Genome Res. 2013;23: 2042–2052.

22. Ewing AD, Ballinger TJ, Earl D, Broad Institute Genome Sequencing and Analysis Program and Platform, Harris CC, Ding L, et al. Retrotransposition of gene transcripts leads to structural variation in mammalian genomes. Genome Biol. 2013;14: R22.

23. Wong Z, Royle NJ, Jeffreys AJ. A novel human DNA polymorphism resulting from transfer of DNA from chromosome 6 to chromosome 16. Genomics. 1990;7: 222–234.

24. Genovese G, Handsaker RE, Li H, Kenny EE, McCarroll SA. Mapping the human reference genome’s missing sequence by three-way admixture in Latino genomes. Am J Hum Genet. 2013;93: 411–421.

25. Sebat J, Lakshmi B, Troge J, Alexander J, Young J, Lundin P, et al. Large-scale copy number polymorphism in the human genome. Science. 2004;305: 525–528.

26. Martin J, Han C, Gordon LA, Terry A, Prabhakar S, She X, et al. The sequence and analysis of duplication-rich human chromosome 16. Nature. 2004;432: 988–994.

27. Doggett NA, Xie G, Meincke LJ, Sutherland RD, Mundt MO, Berbari NS, et al. A 360-kb interchromosomal duplication of the human HYDIN locus. Genomics. 2006;88: 762–771.

28. Dougherty ML, Nuttle X, Penn O, Nelson BJ, Huddleston J, Baker C, et al. The birth of a human-specific neural gene by incomplete duplication and gene fusion. Genome Biol. 2017;18: 49.

29. O’Bleness M, Searles VB, Dickens CM, Astling D, Albracht D, Mak ACY, et al. Finished sequence and assembly of the DUF1220-rich 1q21 region using a haploid human genome. BMC Genomics. 2014;15: 387.

30. Mohajeri K, Cantsilieris S, Huddleston J, Nelson BJ, Coe BP, Campbell CD, et al. Interchromosomal core duplicons drive both evolutionary instability and disease susceptibility of the Chromosome 8p23.1 region. Genome Res. 2016;26: 1453–1467.

31. Potocki L, Chen KS, Park SS, Osterholm DE, Withers MA, Kimonis V, et al. Molecular mechanism for duplication 17p11.2-the homologous recombination reciprocal of the Smith-Magenis microdeletion. Nat Genet. 2000;24: 84–87.

32. MacArthur J, Bowler E, Cerezo M, Gil L, Hall P, Hastings E, et al. The new NHGRI-EBI Catalog of published genome-wide association studies (GWAS Catalog). Nucleic Acids Res. 2017;45: D896–D901.

33. Lu W, Cheng Y-C, Chen K, Wang H, Gerhard GS, Still CD, et al. Evidence for several independent genetic variants affecting lipoprotein (a) cholesterol levels. Hum Mol Genet. 2015;24: 2390–2400.

34. Cross-Disorder Group of the Psychiatric Genomics Consortium. Identification of risk loci with shared effects on five major psychiatric disorders: a genome-wide analysis. Lancet. 2013;381: 1371–1379.

35. Ruderfer DM, Fanous AH, Ripke S, McQuillin A, Amdur RL, Schizophrenia Working Group of the Psychiatric Genomics Consortium, et al. Polygenic dissection of diagnosis and clinical dimensions of bipolar disorder and schizophrenia. Mol Psychiatry. 2014;19: 1017–1024.

36. Marchi E, Kanapin A, Magiorkinis G, Belshaw R. Unfixed endogenous retroviral insertions in the human population. J Virol. 2014;88: 9529–9537.

37. eGTEx Project. Enhancing GTEx by bridging the gaps between genotype, gene expression, and disease. Nat Genet. 2017;49: 1664–1670.

38. Ernst J, Kheradpour P, Mikkelsen TS, Shoresh N, Ward LD, Epstein CB, et al. Mapping and analysis of chromatin state dynamics in nine human cell types. Nature. 2011;473: 43–49.

39. Battle A, Mostafavi S, Zhu X, Potash JB, Weissman MM, McCormick C, et al. Characterizing the genetic basis of transcriptome diversity through RNA-sequencing of 922 individuals. Genome Res. 2014;24: 14–24.

40. Bryois J, Buil A, Evans DM, Kemp JP, Montgomery SB, Conrad DF, et al. Cis and trans effects of human genomic variants on gene expression. PLoS Genet. 2014;10: e1004461.

41. McRae AF, Marioni RE, Shah S, Yang J, Powell JE, Harris SE, et al. Identification of 55,000 Replicated DNA Methylation QTL. Sci Rep. 2018;8: 17605.

42. Usher CL, Handsaker RE, Esko T, Tuke MA, Weedon MN, Hastie AR, et al. Structural forms of the human amylase locus and their relationships to SNPs, haplotypes and obesity. Nat Genet. 2015;47: 921–925.

43. Falchi M, El-Sayed Moustafa JS, Takousis P, Pesce F, Bonnefond A, Andersson-Assarsson JC, et al. Low copy number of the salivary amylase gene predisposes to obesity. Nat Genet. 2014;46: 492–497.

44. Carpenter D, Dhar S, Mitchell LM, Fu B, Tyson J, Shwan NAA, et al. Obesity, starch digestion and amylase: association between copy number variants at human salivary (AMY1) and pancreatic (AMY2) amylase genes. Hum Mol Genet. 2015;24: 3472–3480.

45. Sekar A, Bialas AR, de Rivera H, Davis A, Hammond TR, Kamitaki N, et al. Schizophrenia risk from complex variation of complement component 4. Nature. 2016;530: 177–183.

46. Antonacci F, Dennis MY, Huddleston J, Sudmant PH, Steinberg KM, Rosenfeld JA, et al. Palindromic GOLGA8 core duplicons promote chromosome 15q13.3 microdeletion and evolutionary instability. Nat Genet. 2014;46: 1293–1302.

47. Wilson DM 3rd, Thompson LH. Molecular mechanisms of sister-chromatid exchange. Mutat Res. 2007;616: 11–23.

48. Poplin R, Ruano-Rubio V, DePristo MA, Fennell TJ, Carneiro MO, Van der Auwera GA, et al. Scaling accurate genetic variant discovery to tens of thousands of samples. bioRxiv. 2018. p. 201178. doi:10.1101/201178

49. 1000 Genomes Project Consortium, Auton A, Brooks LD, Durbin RM, Garrison EP, Kang HM, et al. A global reference for human genetic variation. Nature. 2015;526: 68–74.

50. Rolinski S, Horn H, Petzoldt T, Paul L. Identifying cardinal dates in phytoplankton time series to enable the analysis of long-term trends. Oecologia. 2007;153: 997–1008.

51. Chang CC, Chow CC, Tellier LC, Vattikuti S, Purcell SM, Lee JJ. Second-generation PLINK: rising to the challenge of larger and richer datasets. GigaScience. 2015;4: 7.

