## Supplementary material for "Insights into dispersed duplications and complex structural mutations from whole genome sequencing 706 families": S1 Appendix

### CNV_1_67452217_67453437

Insertion Location 95% Confidence Region:

16:77705878-78321065

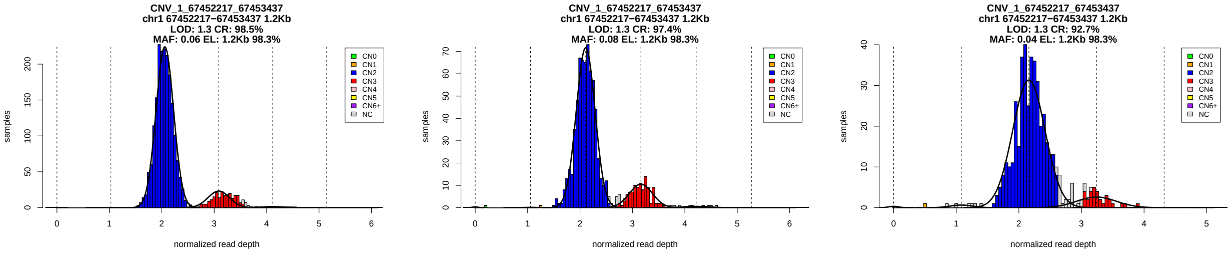


Figure 1: Copy number genotypes in the ASD, ID, and SCZ cohorts.

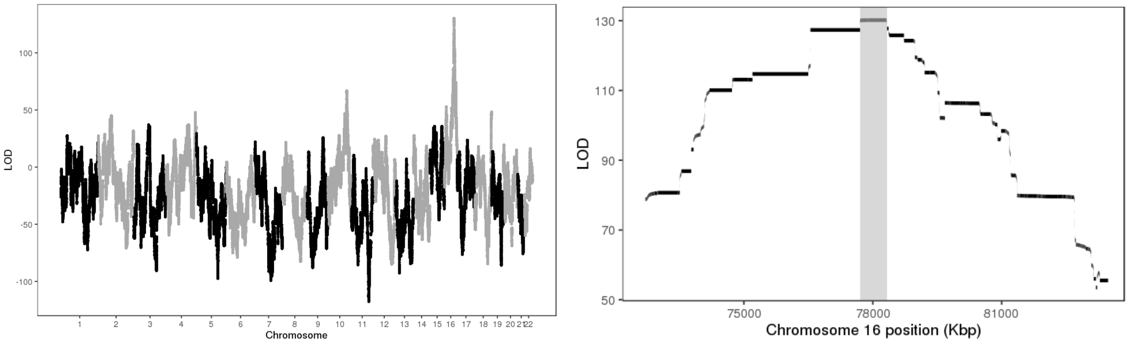


Figure 2: LOD scores of insertion locations across the genome (left) and around the 95% confidence interval (highlighted in gray, right).

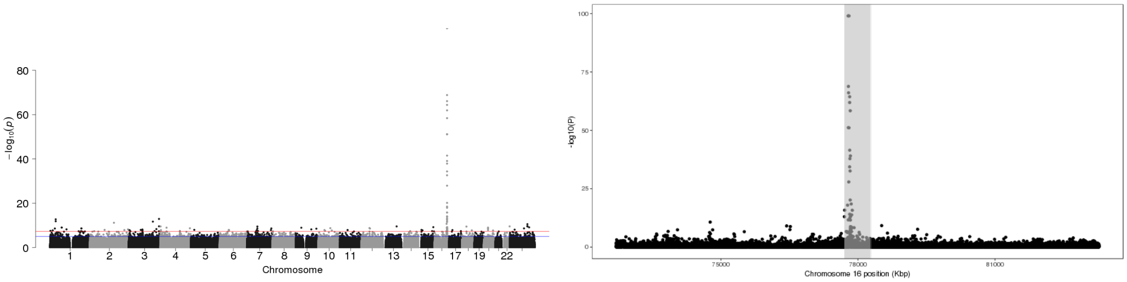


Figure 3: P-values of association between SNP genotypes and duplication copy number across the genome (left), and around the 95% confidence interval (highlighted in gray, right).

All coordinates refer to GRCh37.

### CNV_1_72358720_72361273

Insertion Location 95% Confidence Region:

2:102781911-103257560

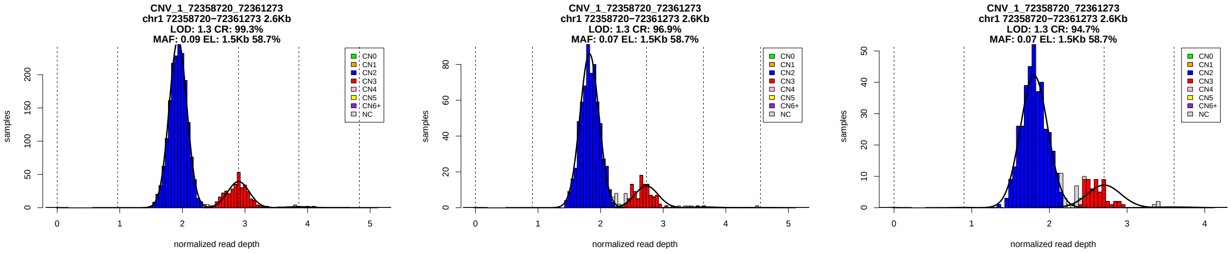


Figure 1: Copy number genotypes in the ASD, ID, and SCZ cohorts.

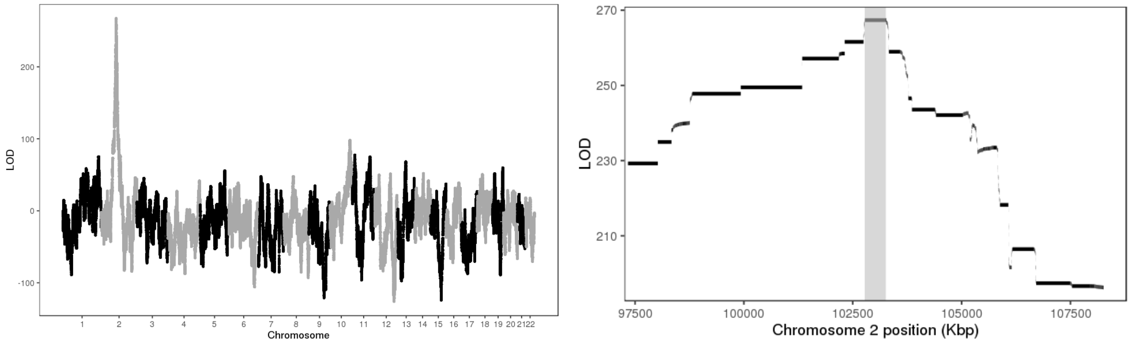


Figure 2: LOD scores of insertion locations across the genome (left) and around the 95% confidence interval (highlighted in gray, right).

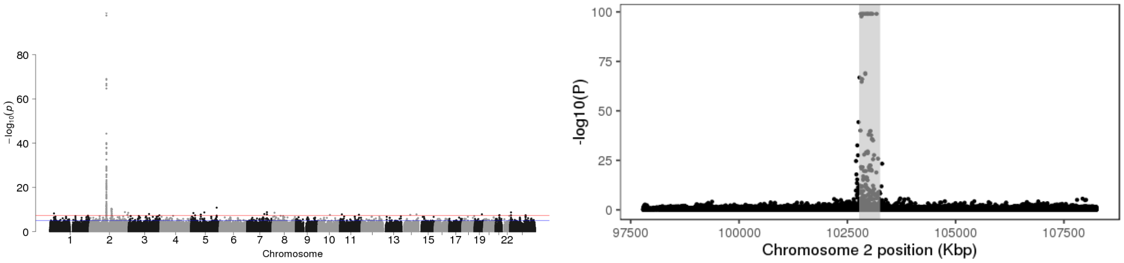


Figure 3: P-values of association between SNP genotypes and duplication copy number across the genome (left), and around the 95% confidence interval (highlighted in gray, right).

All coordinates refer to GRCh37.

### CNV_1_145109500_145118200

Insertion Location 95% Confidence Region:

1:146534681-151499635

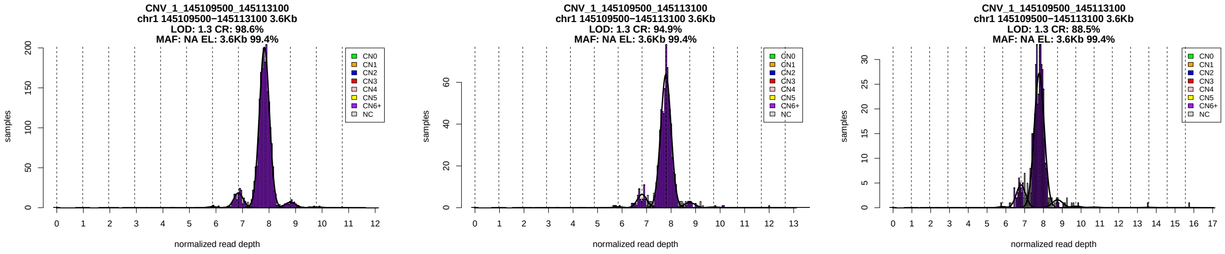


Figure 1: Copy number genotypes in the ASD, ID, and SCZ cohorts.

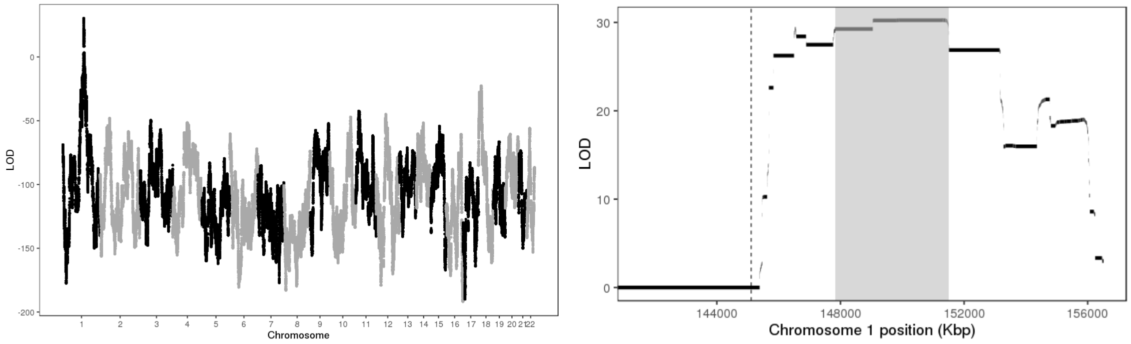


Figure 2: LOD scores of insertion locations across the genome (left) and around the 95% confidence interval (highlighted in gray, right).

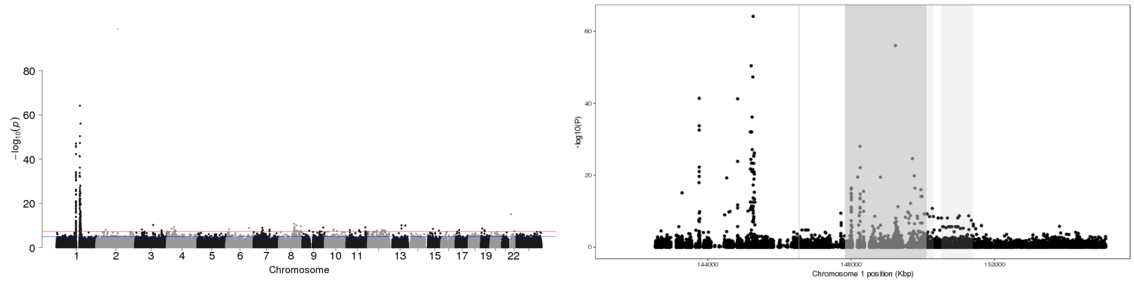


Figure 3: P-values of association between SNP genotypes and duplication copy number across the genome (left), and around the 95% confidence interval (highlighted in gray, right).

All coordinates refer to GRCh37.

### CNV_1_148912285_148953785

Insertion Location 95% Confidence Region:

1:121330376-145383170

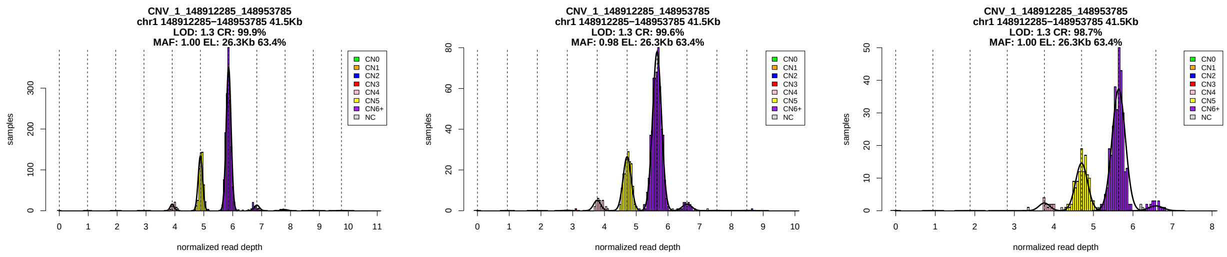


Figure 1: Copy number genotypes in the ASD, ID, and SCZ cohorts.

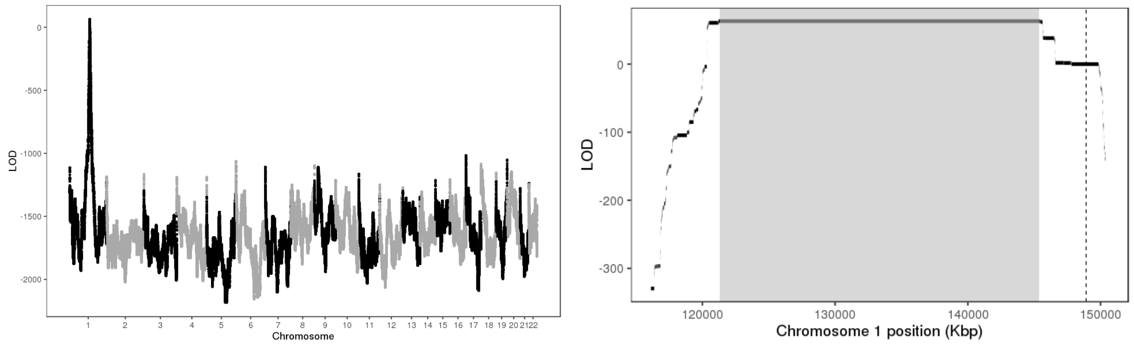


Figure 2: LOD scores of insertion locations across the genome (left) and around the 95% confidence interval (highlighted in gray, right).

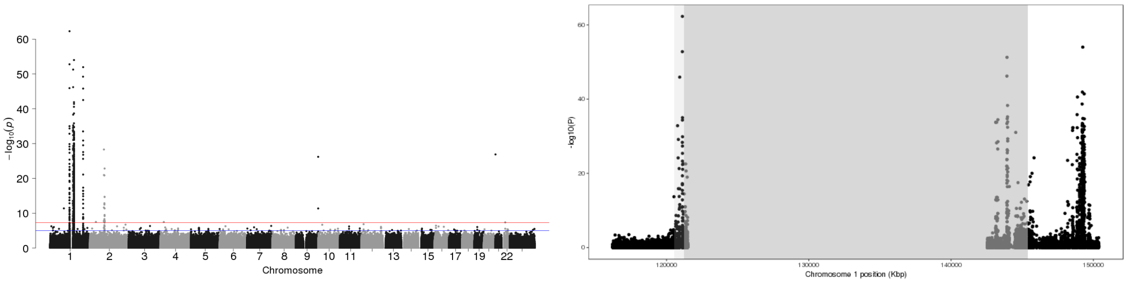


Figure 3: P-values of association between SNP genotypes and duplication copy number across the genome (left), and around the 95% confidence interval (highlighted in gray, right).

All coordinates refer to GRCh37.

# GS_SD_M2_1_148859502_148954460_2_91821870_91916721

Insertion Location 95% Confidence Region:

1:121351198-145383170

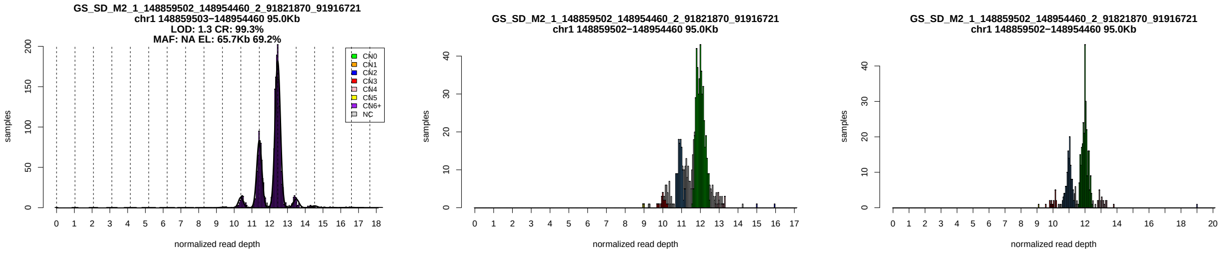


Figure 1: Copy number genotypes in the ASD, ID, and SCZ cohorts.

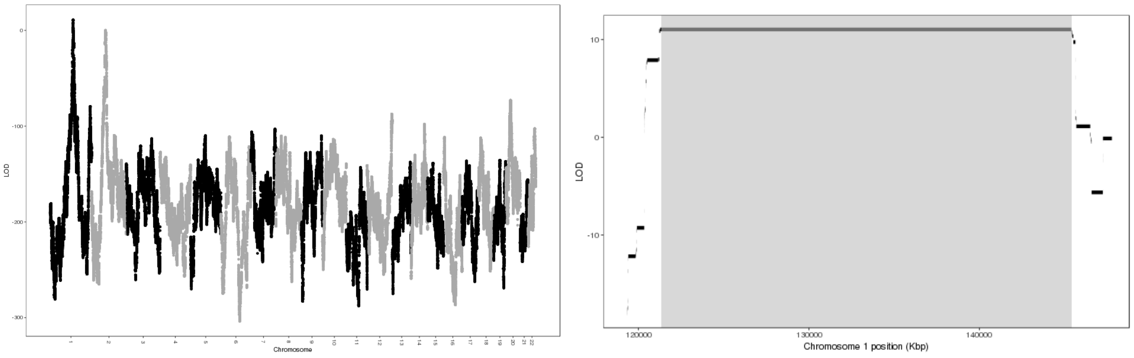


Figure 2: LOD scores of insertion locations across the genome (left) and around the 95% confidence interval (highlighted in gray, right).

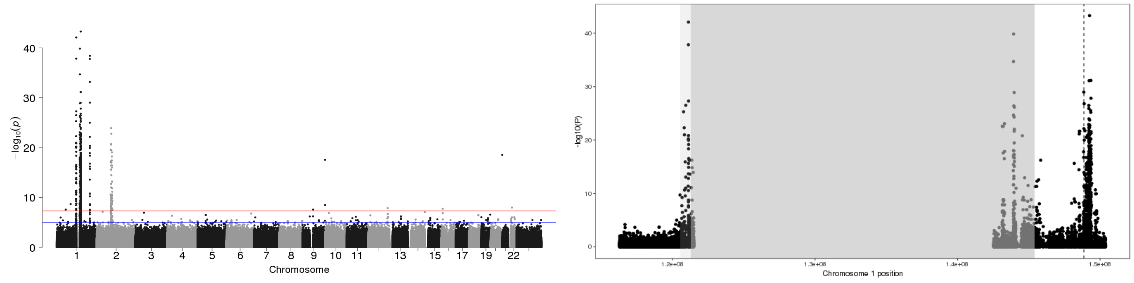


Figure 3: P-values of association between SNP genotypes and duplication copy number across the genome (left), and around the 95% confidence interval (highlighted in gray, right).

All coordinates refer to GRCh37.

### CNV_1_149019900_149024200

Insertion Location 95% Confidence Region:

1:120461536-145383170

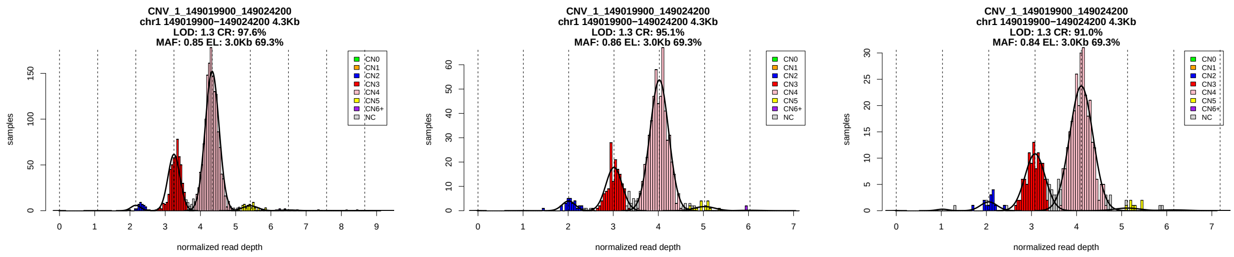


Figure 1: Copy number genotypes in the ASD, ID, and SCZ cohorts.

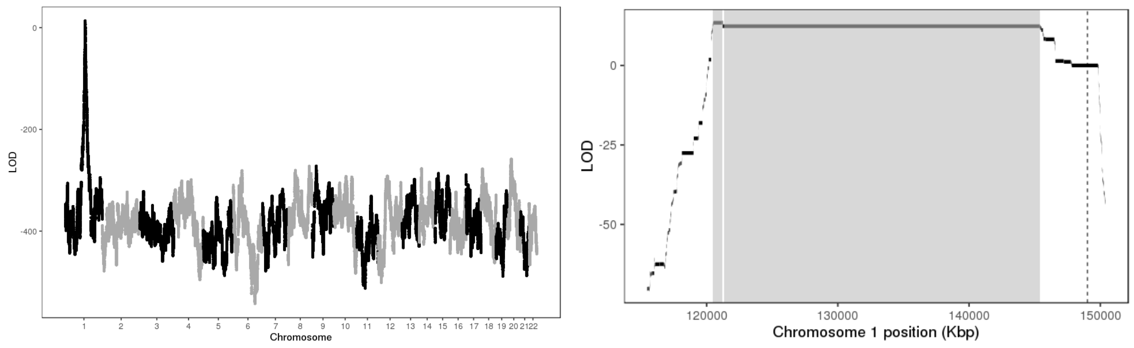


Figure 2: LOD scores of insertion locations across the genome (left) and around the 95% confidence interval (highlighted in gray, right).

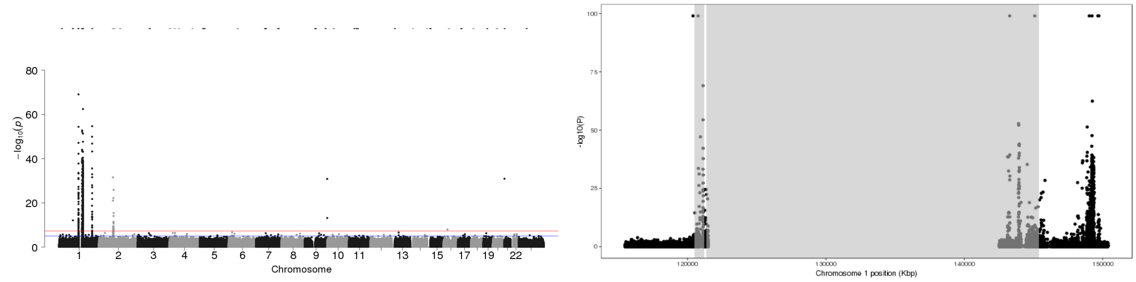


Figure 3: P-values of association between SNP genotypes and duplication copy number across the genome (left), and around the 95% confidence interval (highlighted in gray, right).

All coordinates refer to GRCh37.

### CNV_1_149025500_149031600

Insertion Location 95% Confidence Region:

1:121225332-145383170

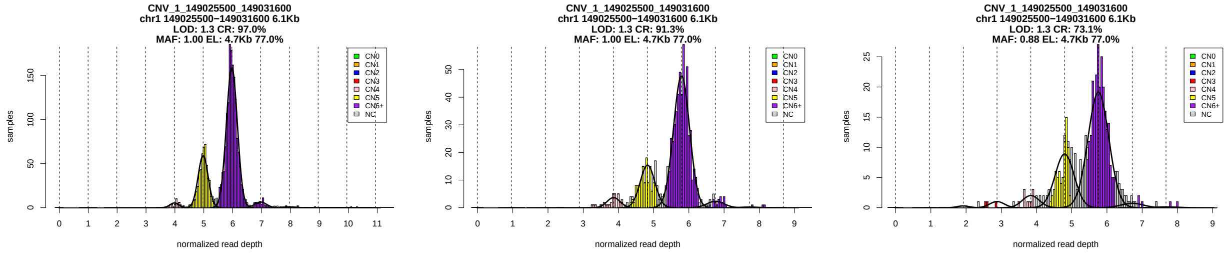


Figure 1: Copy number genotypes in the ASD, ID, and SCZ cohorts.

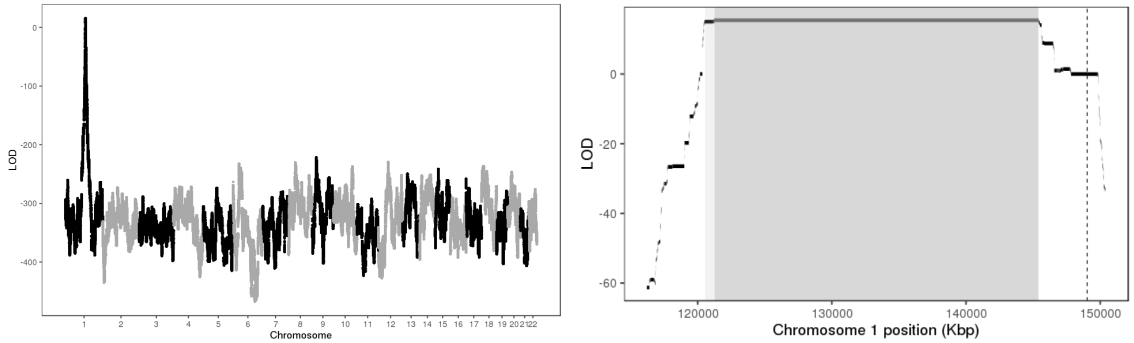


Figure 2: LOD scores of insertion locations across the genome (left) and around the 95% confidence interval (highlighted in gray, right).

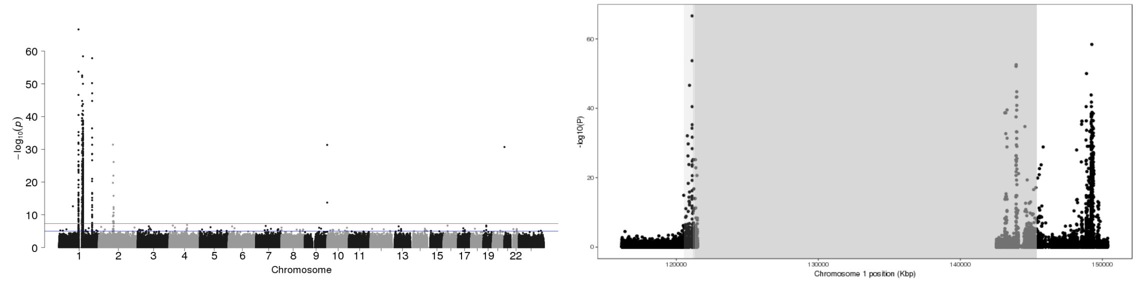


Figure 3: P-values of association between SNP genotypes and duplication copy number across the genome (left), and around the 95% confidence interval (highlighted in gray, right).

All coordinates refer to GRCh37.

### CNV_1_149232622_149239750

Insertion Location 95% Confidence Region:

1:121330376-145383170

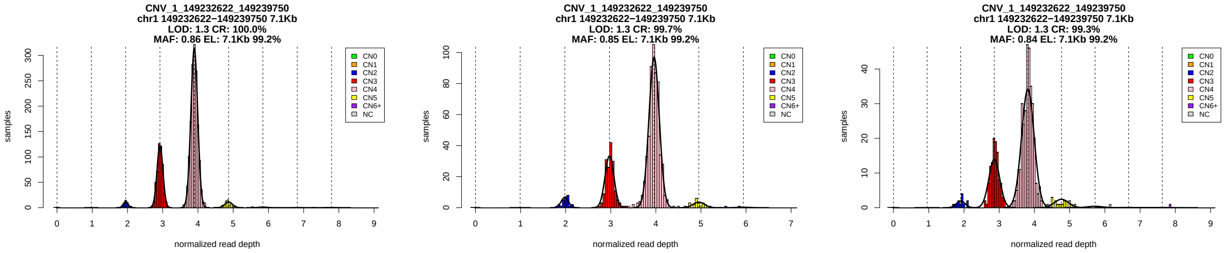


Figure 1: Copy number genotypes in the ASD, ID, and SCZ cohorts.

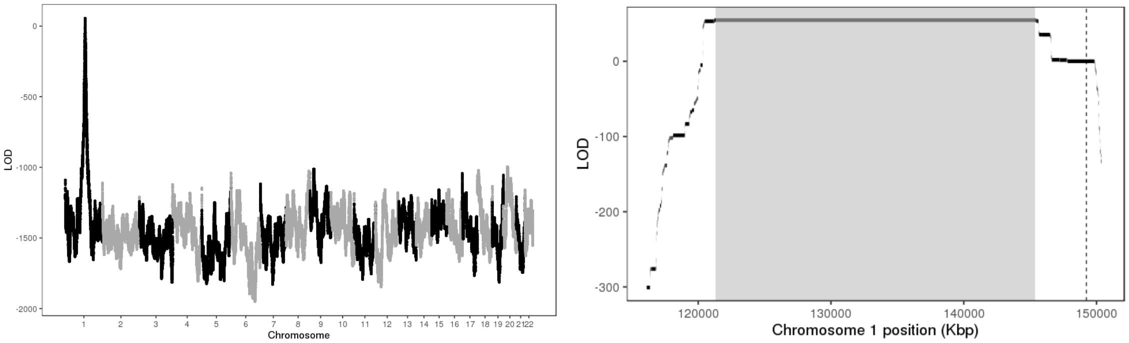


Figure 2: LOD scores of insertion locations across the genome (left) and around the 95% confidence interval (highlighted in gray, right).

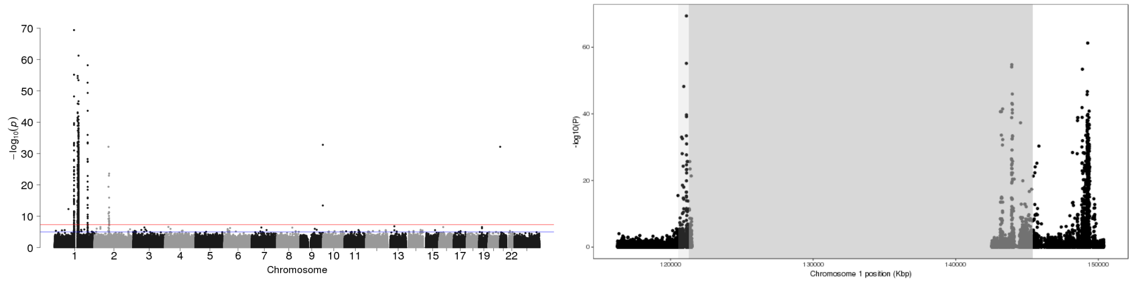


Figure 3: P-values of association between SNP genotypes and duplication copy number across the genome (left), and around the 95% confidence interval (highlighted in gray, right).

All coordinates refer to GRCh37.

# GS_SD_M2_1_149238668_149415268_1_149626126_149799720

Insertion Location 95% Confidence Region:

1:121330376-145383170

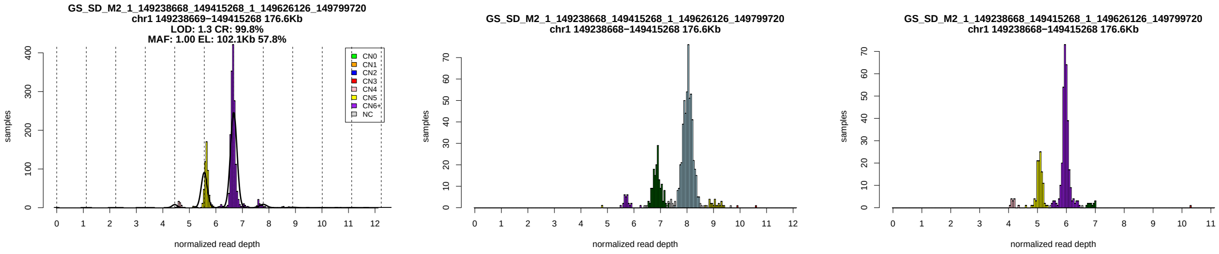


Figure 1: Copy number genotypes in the ASD, ID, and SCZ cohorts.

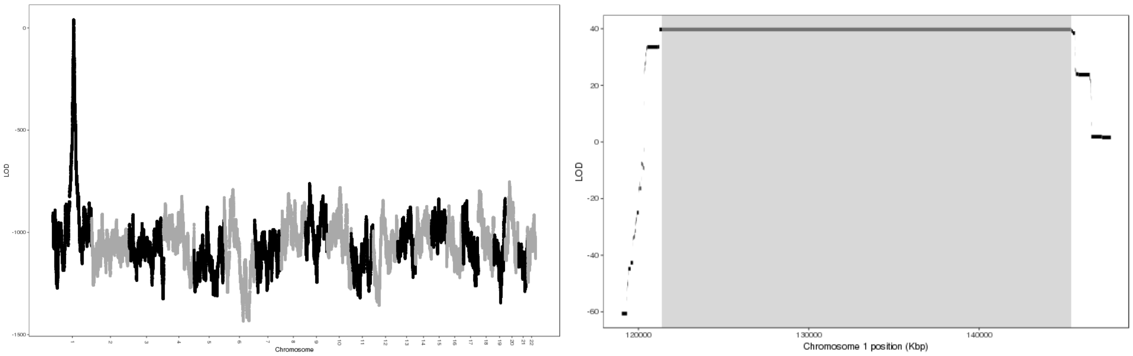


Figure 2: LOD scores of insertion locations across the genome (left) and around the 95% confidence interval (highlighted in gray, right).

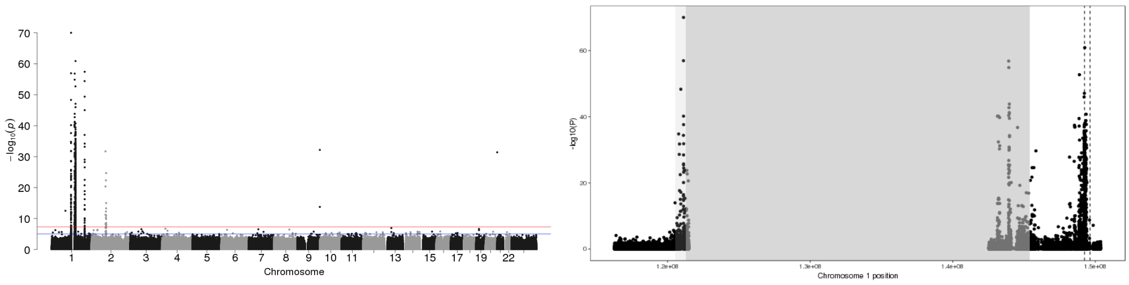


Figure 3: P-values of association between SNP genotypes and duplication copy number across the genome (left), and around the 95% confidence interval (highlighted in gray, right).

All coordinates refer to GRCh37.

### CNV_1_149297554_149315000

Insertion Location 95% Confidence Region:

1:121262781-145383170

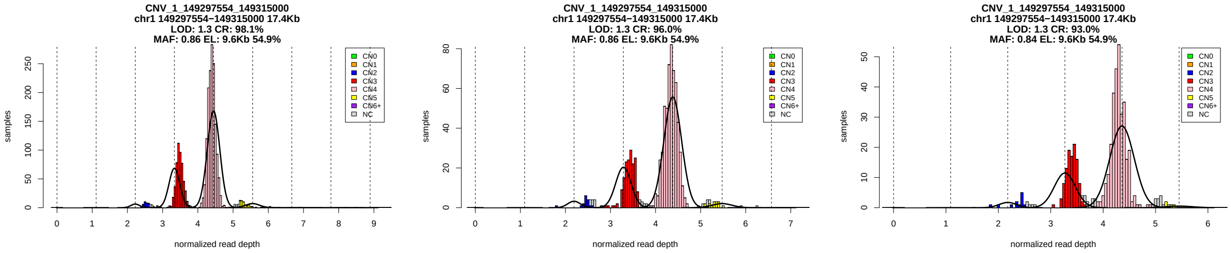


Figure 1: Copy number genotypes in the ASD, ID, and SCZ cohorts.

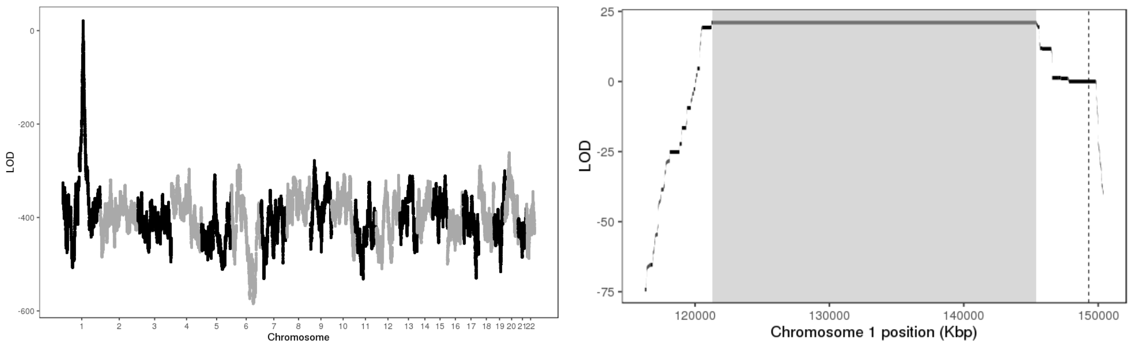


Figure 2: LOD scores of insertion locations across the genome (left) and around the 95% confidence interval (highlighted in gray, right).

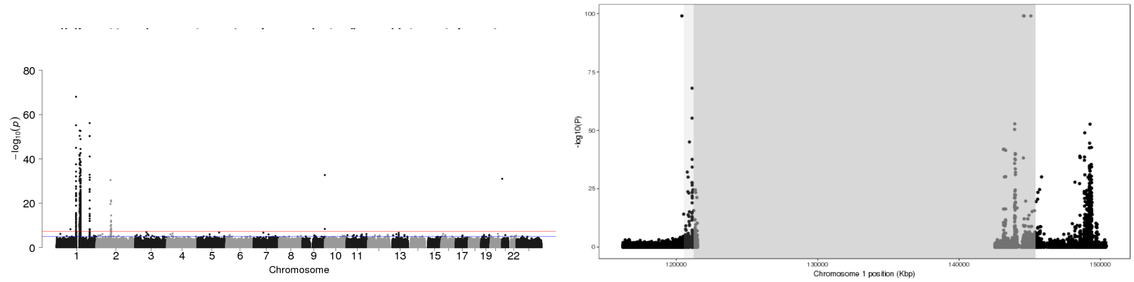


Figure 3: P-values of association between SNP genotypes and duplication copy number across the genome (left), and around the 95% confidence interval (highlighted in gray, right).

All coordinates refer to GRCh37.

### CNV_1_234912161_234919407

Insertion Location 95% Confidence Region:

1:16691022-17337391

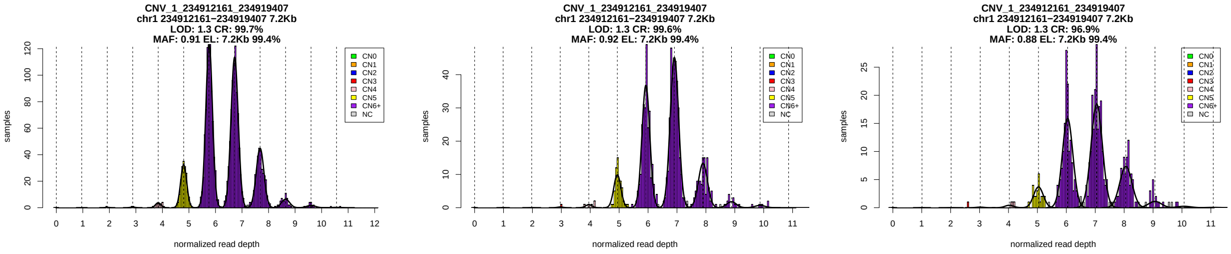


Figure 1: Copy number genotypes in the ASD, ID, and SCZ cohorts.

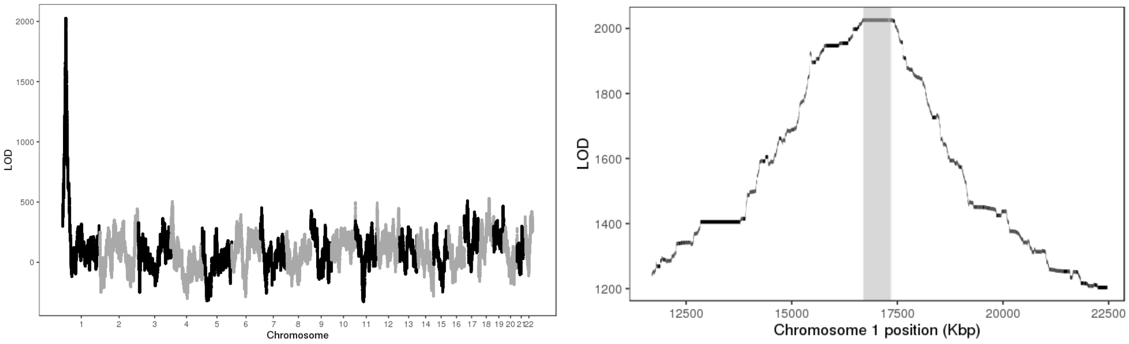


Figure 2: LOD scores of insertion locations across the genome (left) and around the 95% confidence interval (highlighted in gray, right).

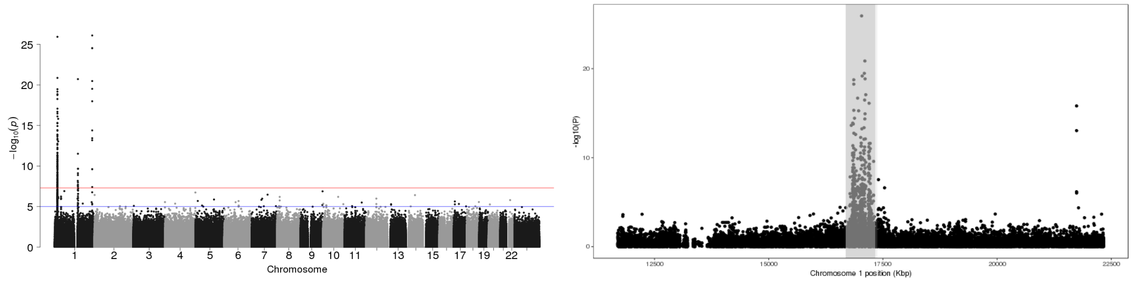


Figure 3: P-values of association between SNP genotypes and duplication copy number across the genome (left), and around the 95% confidence interval (highlighted in gray, right).

All coordinates refer to GRCh37.

### CNV_2_37958100_38002800

Insertion Location 95% Confidence Region:

22:16554852-17608038

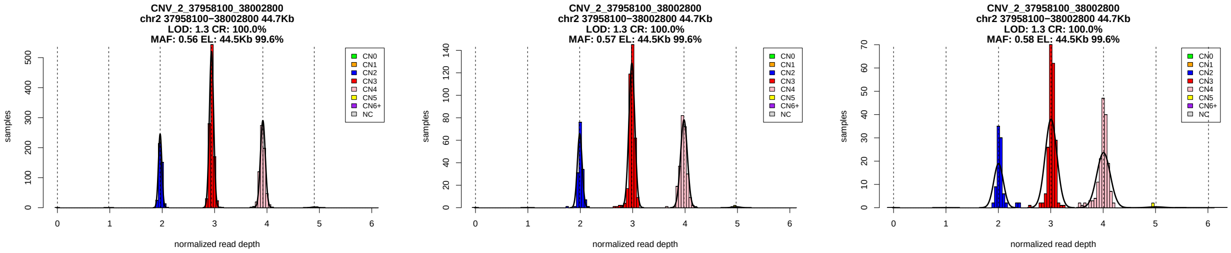


Figure 1: Copy number genotypes in the ASD, ID, and SCZ cohorts.

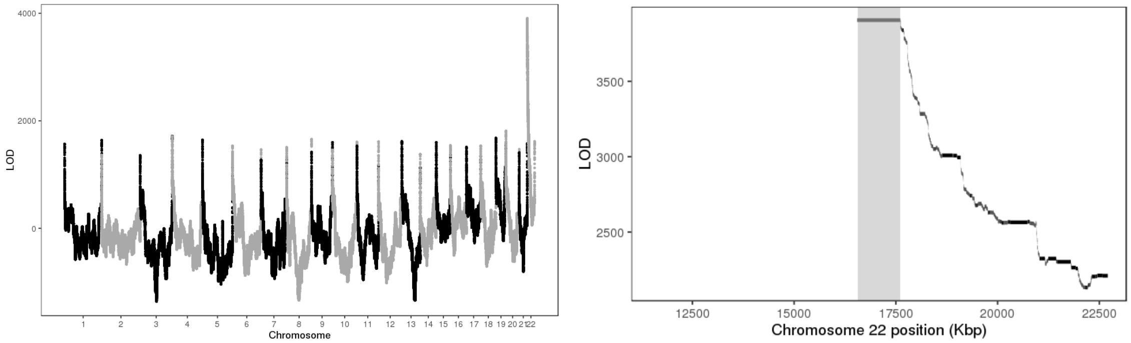


Figure 2: LOD scores of insertion locations across the genome (left) and around the 95% confidence interval (highlighted in gray, right).

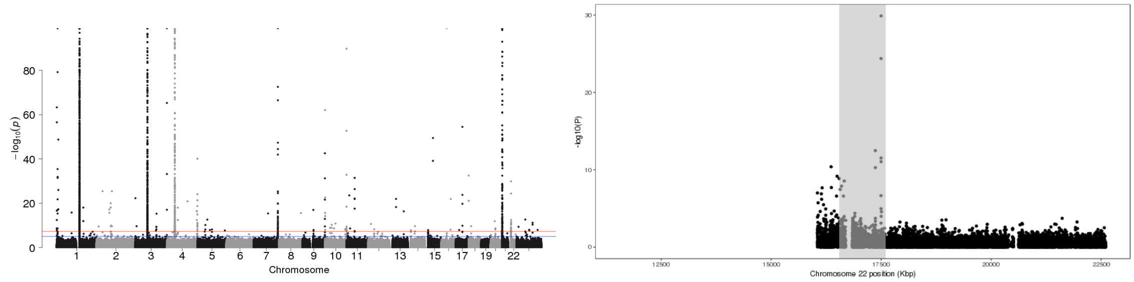


Figure 3: P-values of association between SNP genotypes and duplication copy number across the genome (left), and around the 95% confidence interval (highlighted in gray, right).

All coordinates refer to GRCh37.

### CNV_2_91759300_91821900

Insertion Location 95% Confidence Region:

1:145402354-145626119

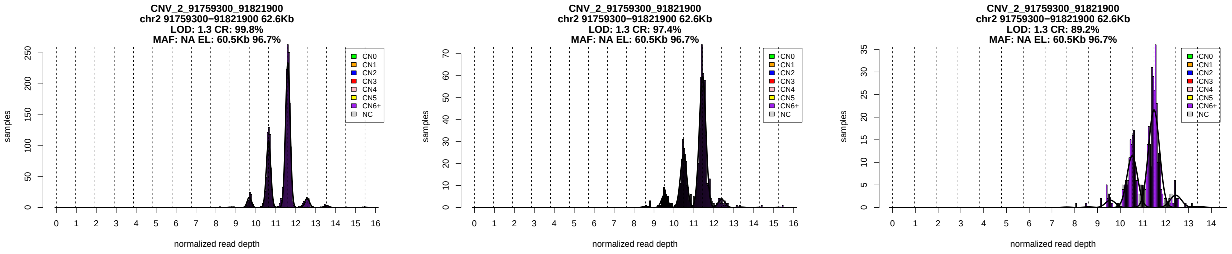


Figure 1: Copy number genotypes in the ASD, ID, and SCZ cohorts.

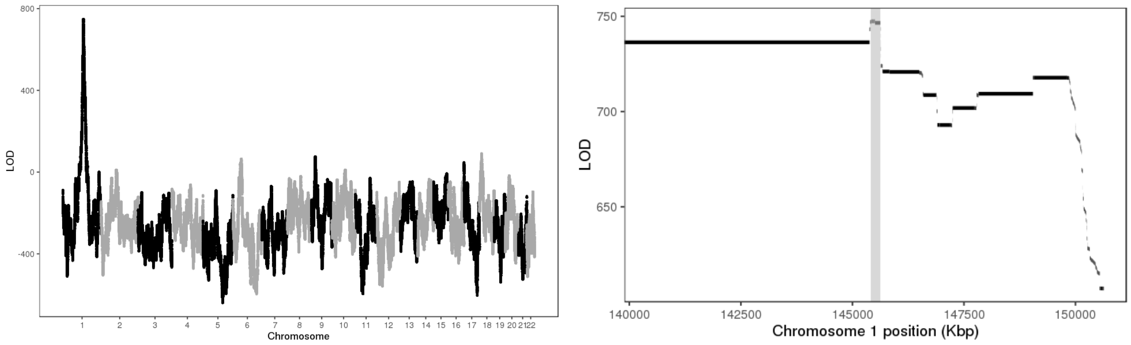


Figure 2: LOD scores of insertion locations across the genome (left) and around the 95% confidence interval (highlighted in gray, right).

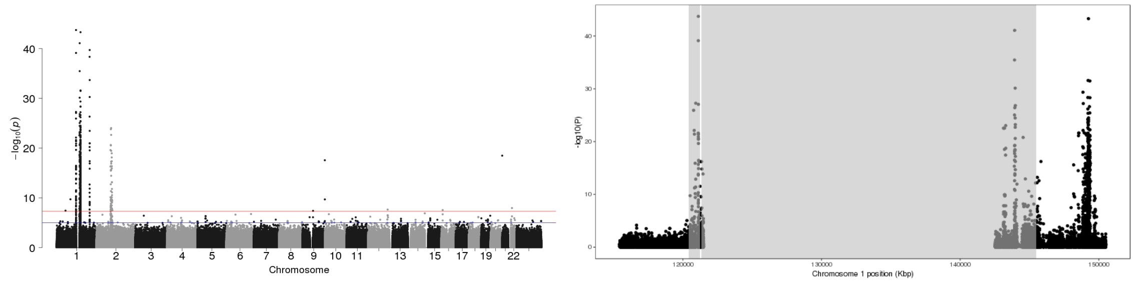


Figure 3: P-values of association between SNP genotypes and duplication copy number across the genome (left), and around the 95% confidence interval (highlighted in gray, right).

All coordinates refer to GRCh37.

### CNV_2_96726300_96735000

Insertion Location 95% Confidence Region:

2:98023626-98307823

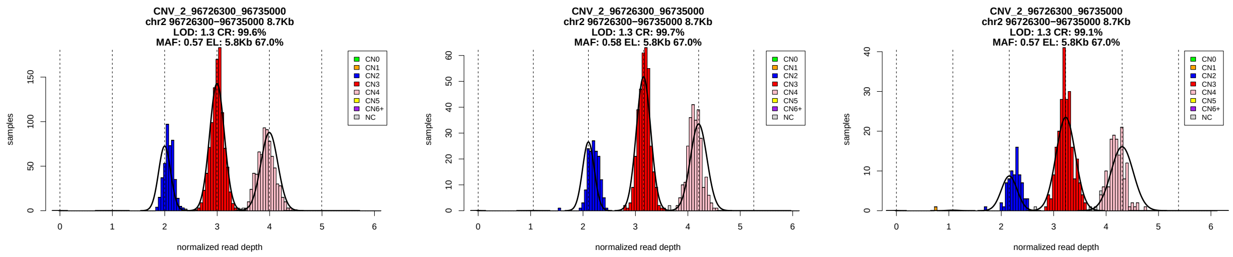


Figure 1: Copy number genotypes in the ASD, ID, and SCZ cohorts.

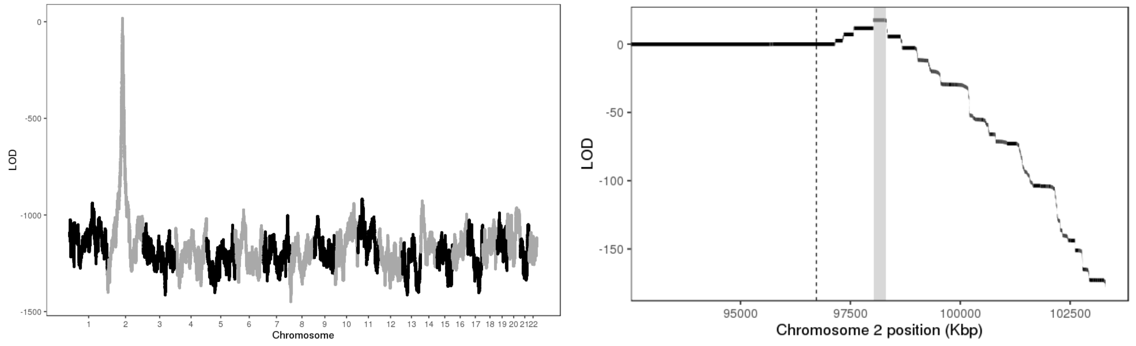


Figure 2: LOD scores of insertion locations across the genome (left) and around the 95% confidence interval (highlighted in gray, right).

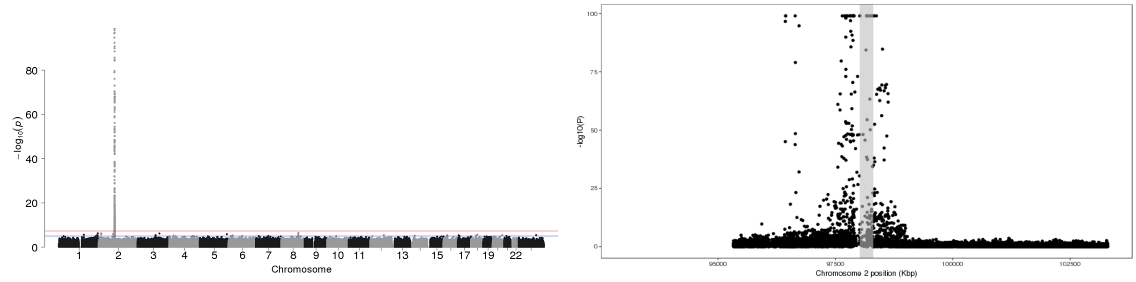


Figure 3: P-values of association between SNP genotypes and duplication copy number across the genome (left), and around the 95% confidence interval (highlighted in gray, right).

All coordinates refer to GRCh37.

### CNV_2_112177604_112185178

Insertion Location 95% Confidence Region:

2:87061188-88295232

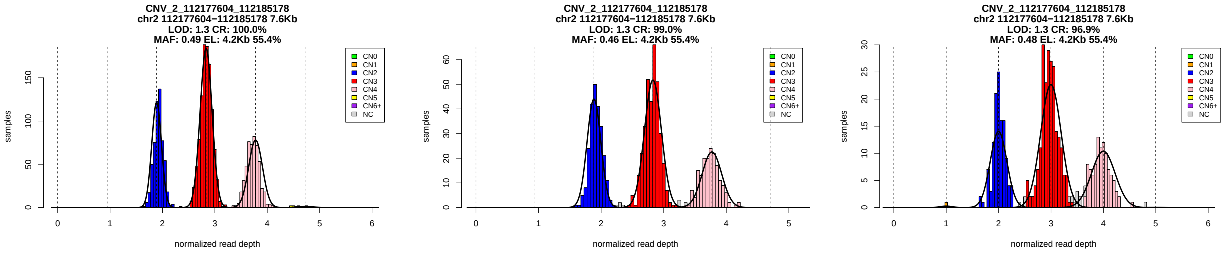


Figure 1: Copy number genotypes in the ASD, ID, and SCZ cohorts.

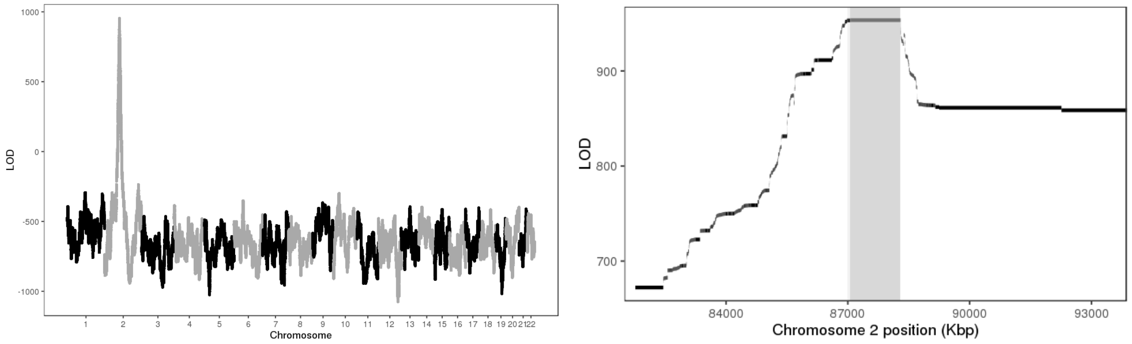


Figure 2: LOD scores of insertion locations across the genome (left) and around the 95% confidence interval (highlighted in gray, right).

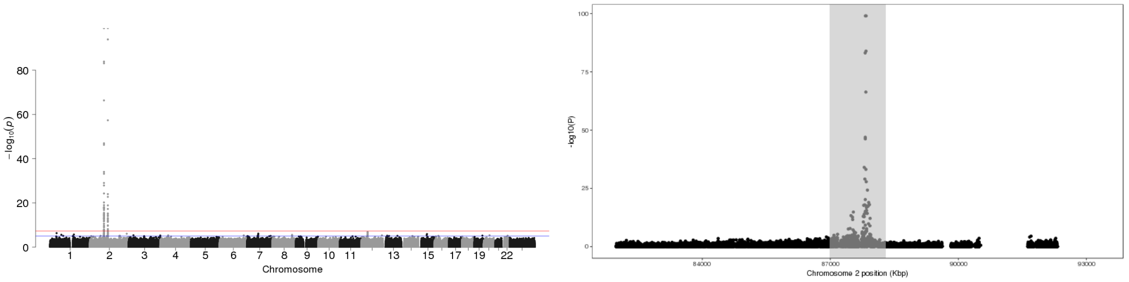


Figure 3: P-values of association between SNP genotypes and duplication copy number across the genome (left), and around the 95% confidence interval (highlighted in gray, right).

All coordinates refer to GRCh37.

### CNV_2_112484464_112494225

Insertion Location 95% Confidence Region:

2:85886667-88295232

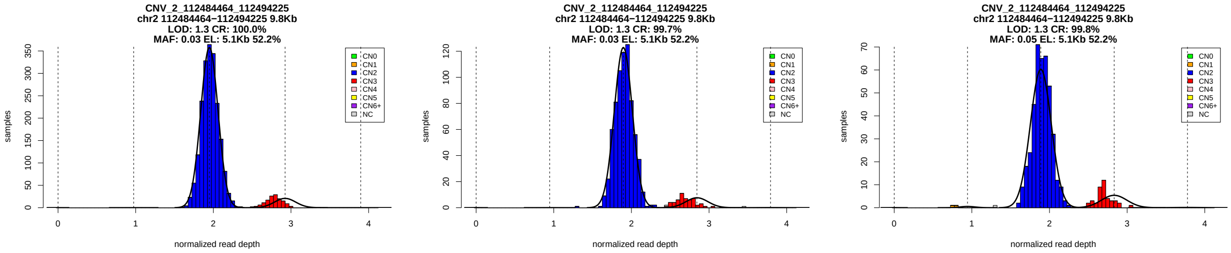


Figure 1: Copy number genotypes in the ASD, ID, and SCZ cohorts.

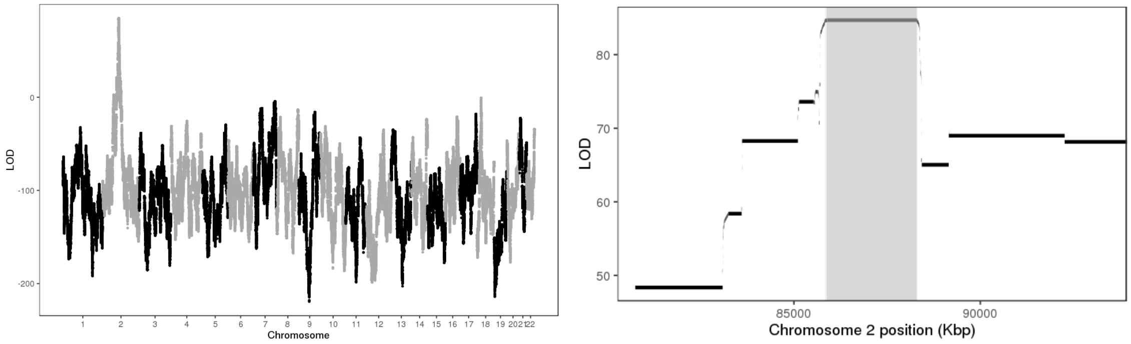


Figure 2: LOD scores of insertion locations across the genome (left) and around the 95% confidence interval (highlighted in gray, right).

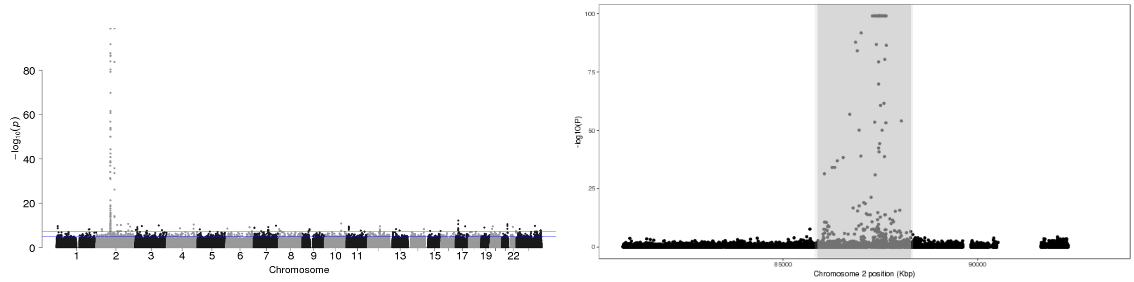


Figure 3: P-values of association between SNP genotypes and duplication copy number across the genome (left), and around the 95% confidence interval (highlighted in gray, right).

All coordinates refer to GRCh37.

### CNV_2_113045432_113049031

Insertion Location 95% Confidence Region:

2:86183095-86897501

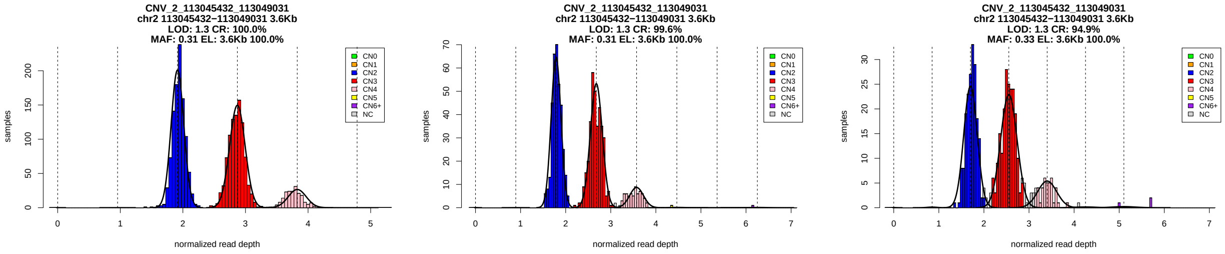


Figure 1: Copy number genotypes in the ASD, ID, and SCZ cohorts.

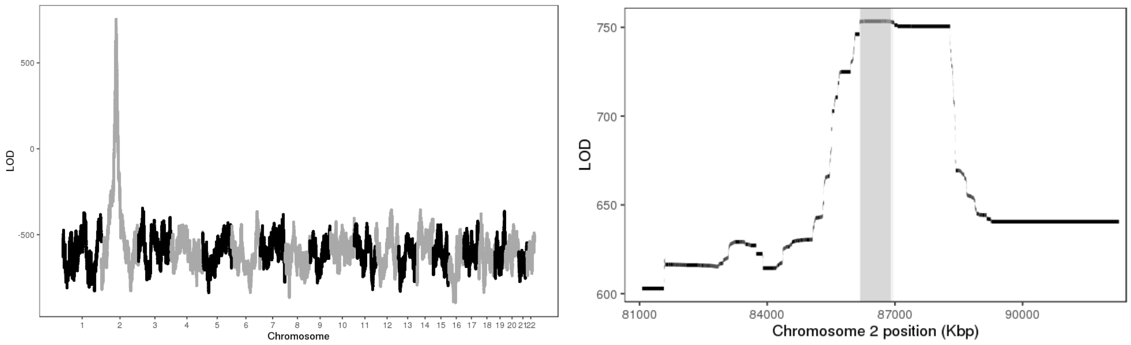


Figure 2: LOD scores of insertion locations across the genome (left) and around the 95% confidence interval (highlighted in gray, right).

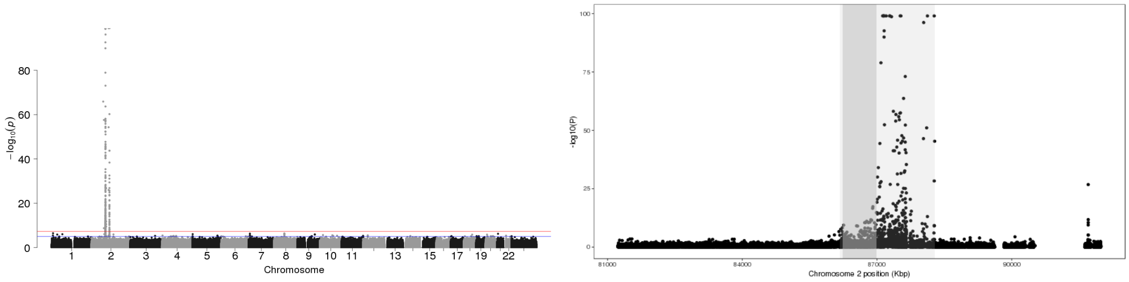


Figure 3: P-values of association between SNP genotypes and duplication copy number across the genome (left), and around the 95% confidence interval (highlighted in gray, right).

All coordinates refer to GRCh37.

### CNV_2_179296140_179315958_ENST00000325748.4

Insertion Location 95% Confidence Region:

6:32350107-32675032

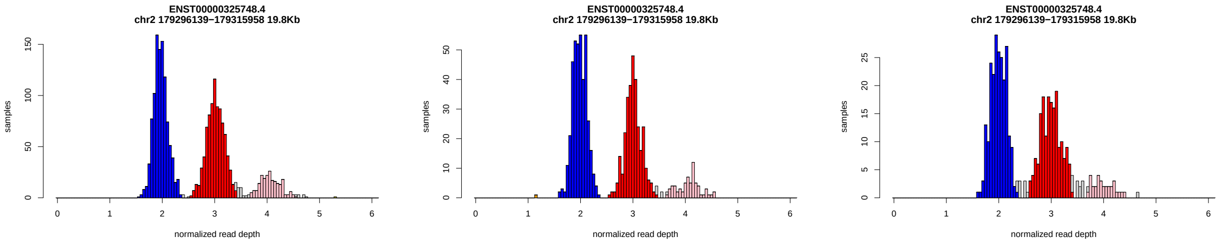


Figure 1: Copy number genotypes in the ASD, ID, and SCZ cohorts.

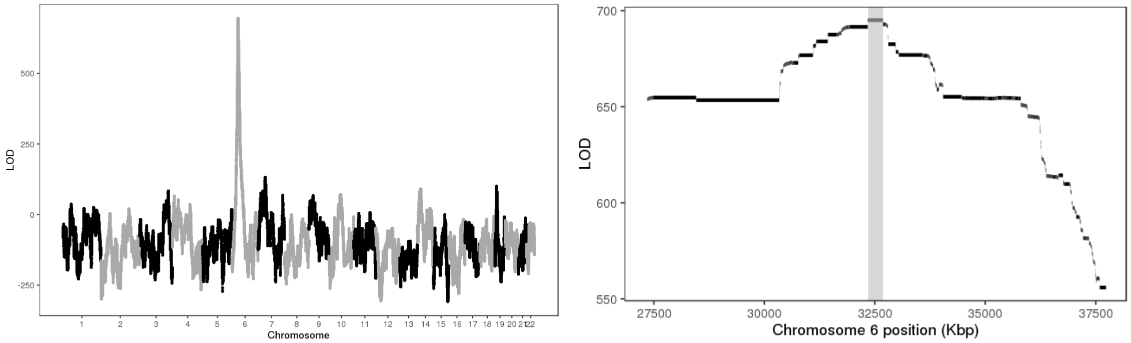


Figure 2: LOD scores of insertion locations across the genome (left) and around the 95% confidence interval (highlighted in gray, right).

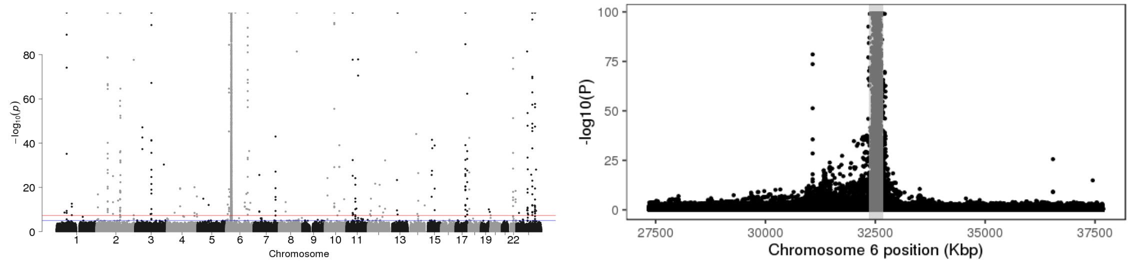


Figure 3: P-values of association between SNP genotypes and duplication copy number across the genome (left), and around the 95% confidence interval (highlighted in gray, right).

All coordinates refer to GRCh37.

### CNV_2_228189884_228222550_ENST00000304593.9

Insertion Location 95% Confidence Region:

15:93718354-93845684

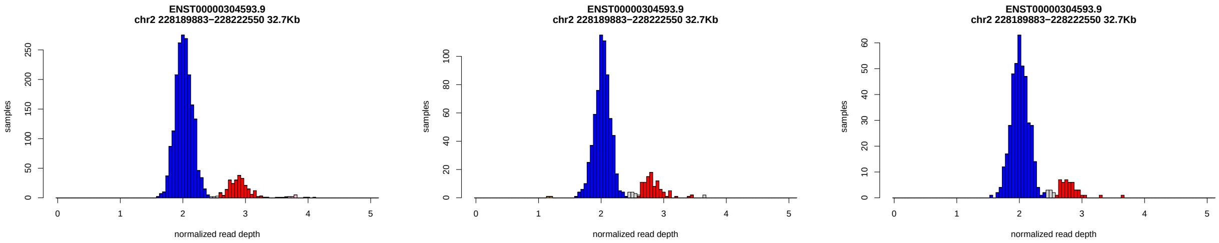


Figure 1: Copy number genotypes in the ASD, ID, and SCZ cohorts.

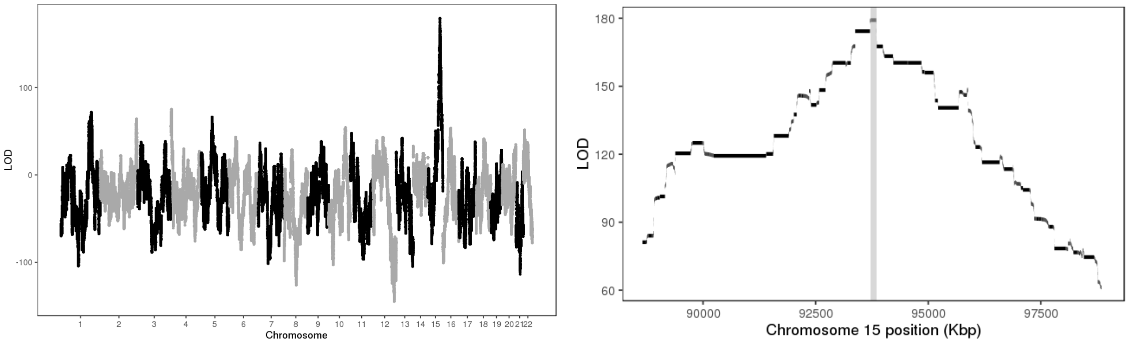


Figure 2: LOD scores of insertion locations across the genome (left) and around the 95% confidence interval (highlighted in gray, right).

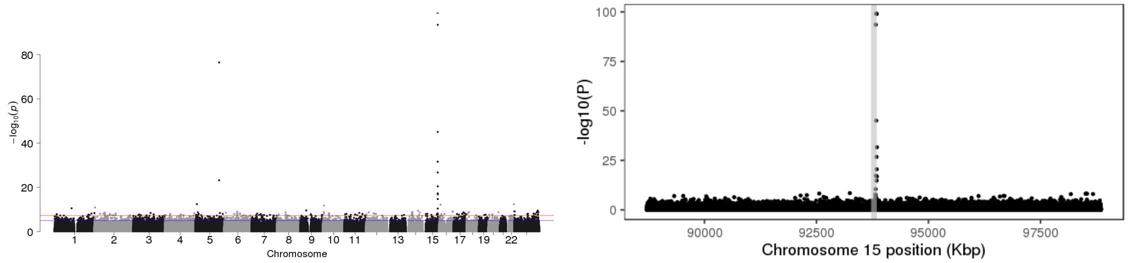


Figure 3: P-values of association between SNP genotypes and duplication copy number across the genome (left), and around the 95% confidence interval (highlighted in gray, right).

All coordinates refer to GRCh37.

### CNV_3_30412801_30413900

Insertion Location 95% Confidence Region:

6:57189829-65243447

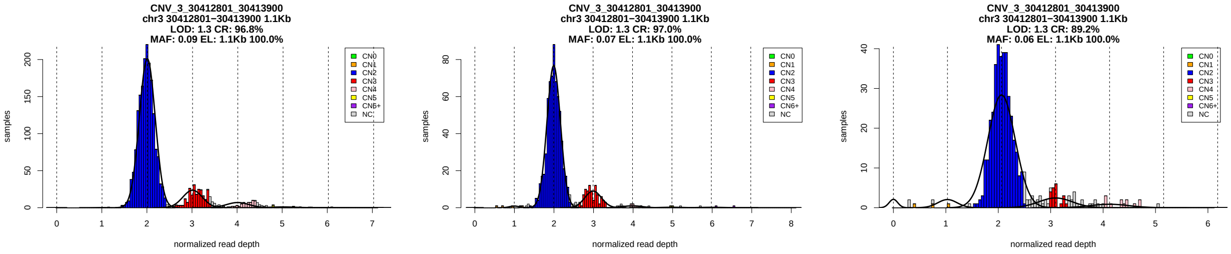


Figure 1: Copy number genotypes in the ASD, ID, and SCZ cohorts.

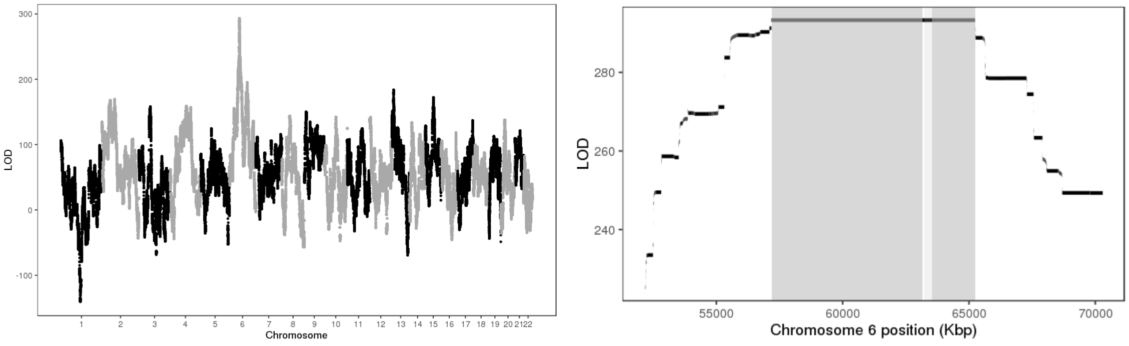


Figure 2: LOD scores of insertion locations across the genome (left) and around the 95% confidence interval (highlighted in gray, right).

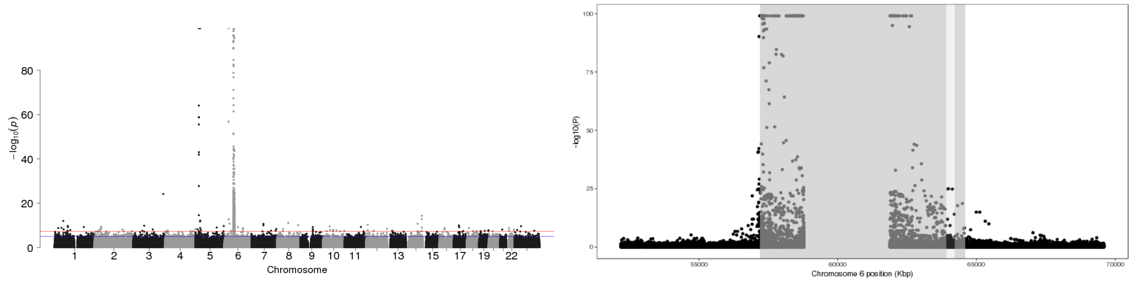


Figure 3: P-values of association between SNP genotypes and duplication copy number across the genome (left), and around the 95% confidence interval (highlighted in gray, right).

All coordinates refer to GRCh37.

### CNV_3_195399263_195403622

Insertion Location 95% Confidence Region:

3:195652899-195857758

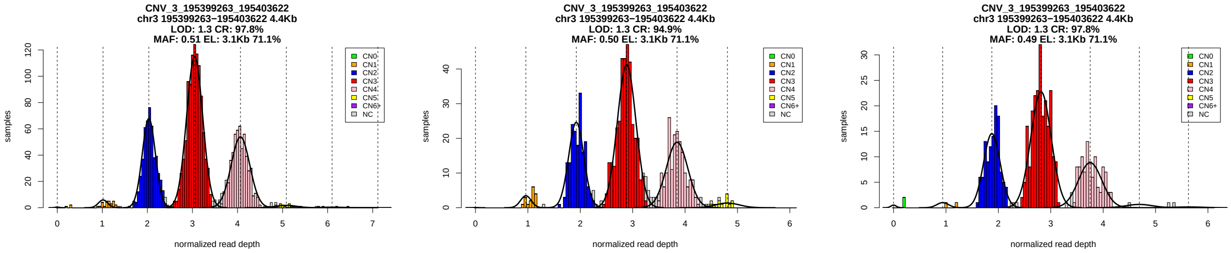


Figure 1: Copy number genotypes in the ASD, ID, and SCZ cohorts.

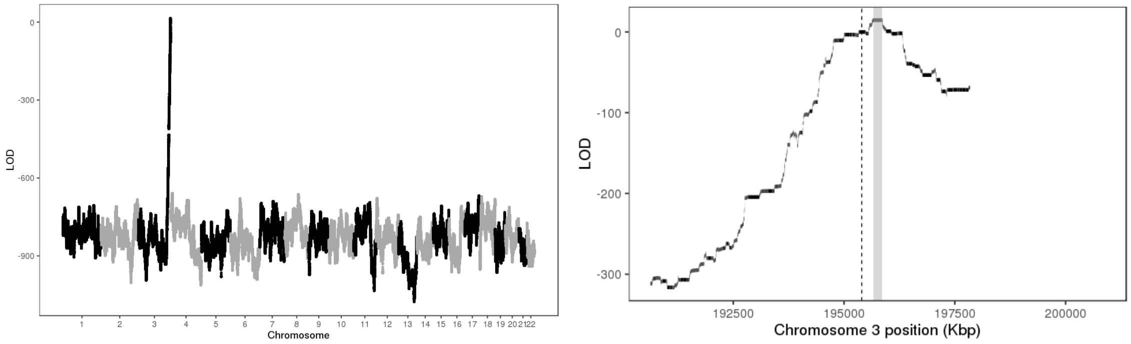


Figure 2: LOD scores of insertion locations across the genome (left) and around the 95% confidence interval (highlighted in gray, right).

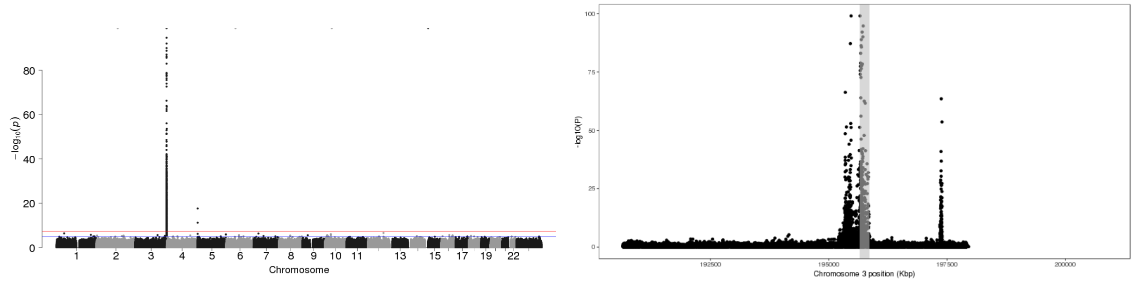


Figure 3: P-values of association between SNP genotypes and duplication copy number across the genome (left), and around the 95% confidence interval (highlighted in gray, right).

All coordinates refer to GRCh37.

### CNV_3_195457200_195471000

Insertion Location 95% Confidence Region:

3:195652899-195857758

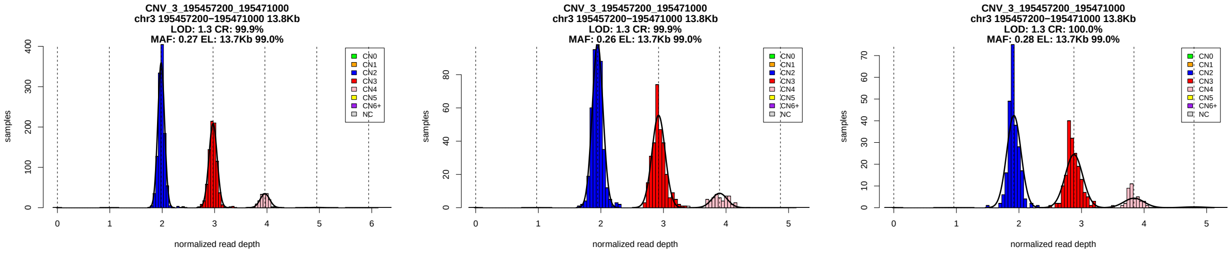


Figure 1: Copy number genotypes in the ASD, ID, and SCZ cohorts.

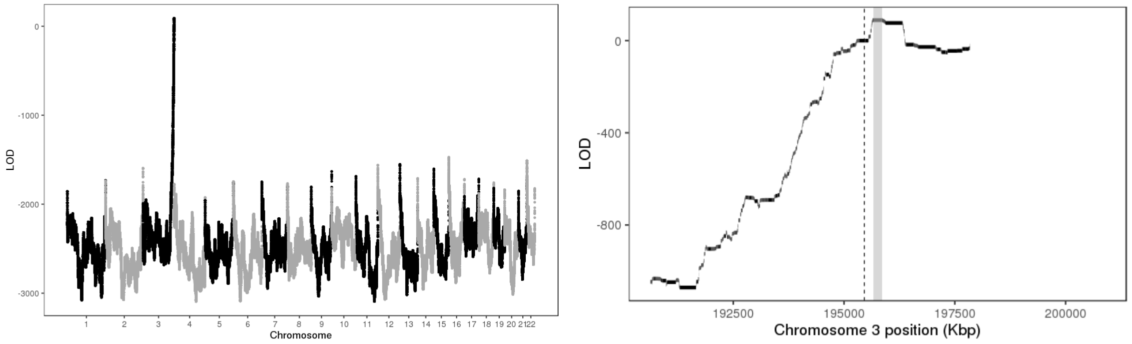


Figure 2: LOD scores of insertion locations across the genome (left) and around the 95% confidence interval (highlighted in gray, right).

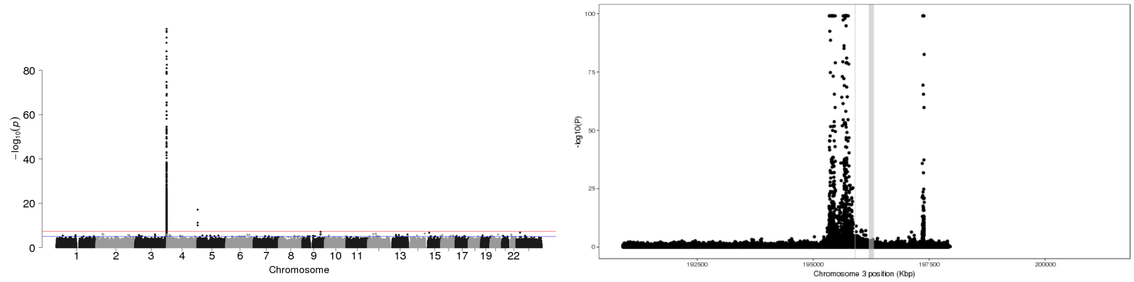


Figure 3: P-values of association between SNP genotypes and duplication copy number across the genome (left), and around the 95% confidence interval (highlighted in gray, right).

All coordinates refer to GRCh37.

### CNV_4_4015000_4021500

Insertion Location 95% Confidence Region:

4:8587465-11156745

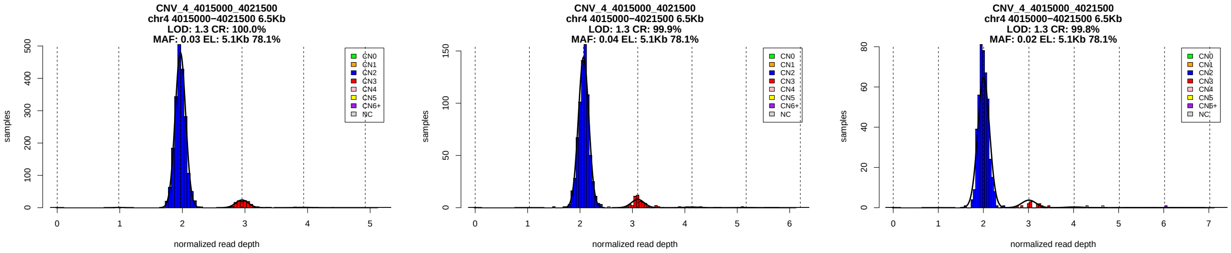


Figure 1: Copy number genotypes in the ASD, ID, and SCZ cohorts.

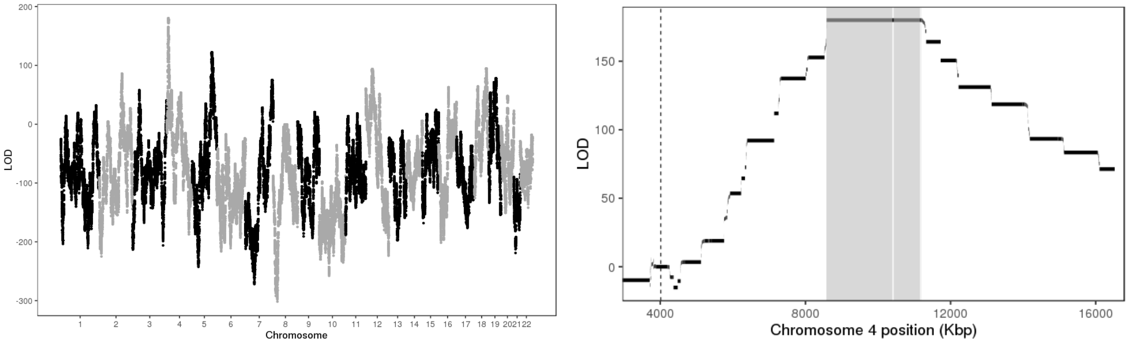


Figure 2: LOD scores of insertion locations across the genome (left) and around the 95% confidence interval (highlighted in gray, right).

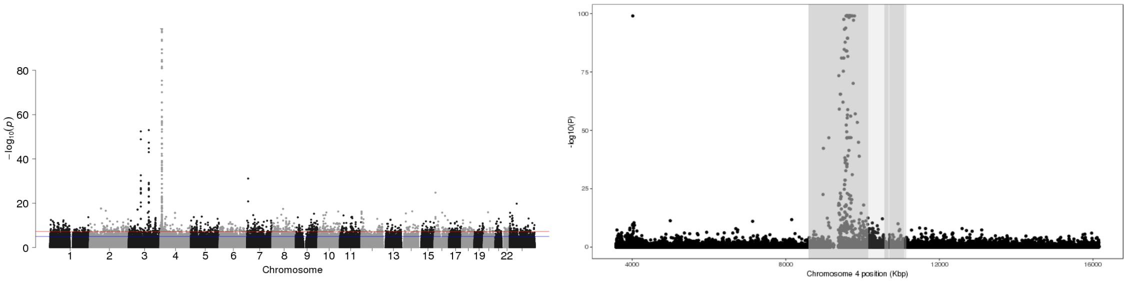


Figure 3: P-values of association between SNP genotypes and duplication copy number across the genome (left), and around the 95% confidence interval (highlighted in gray, right).

All coordinates refer to GRCh37.

### CNV_4_49601000_49631000

Insertion Location 95% Confidence Region:

14:19264866-20475208

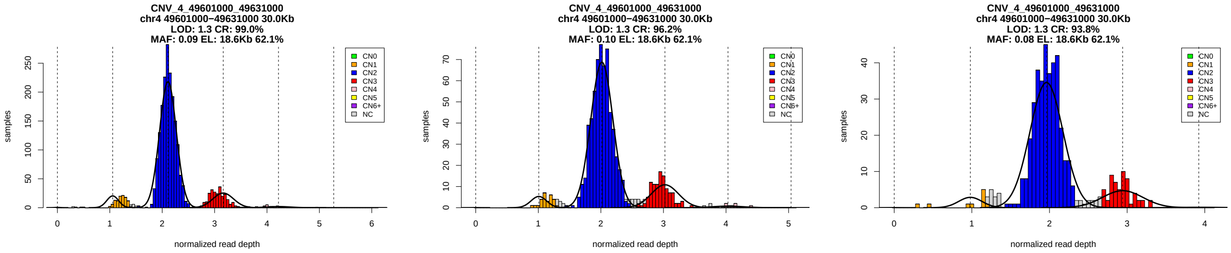


Figure 1: Copy number genotypes in the ASD, ID, and SCZ cohorts.

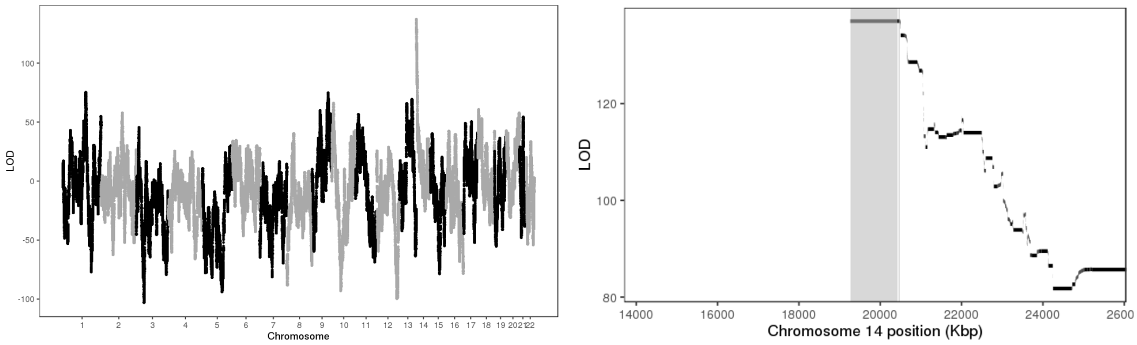


Figure 2: LOD scores of insertion locations across the genome (left) and around the 95% confidence interval (highlighted in gray, right).

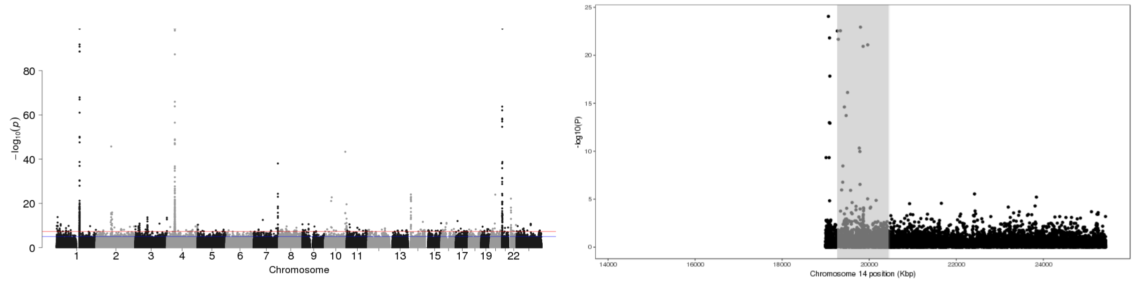


Figure 3: P-values of association between SNP genotypes and duplication copy number across the genome (left), and around the 95% confidence interval (highlighted in gray, right).

All coordinates refer to GRCh37.

### CNV_4_49585000_49600000

Insertion Location 95% Confidence Region:

14:19264866-20475208

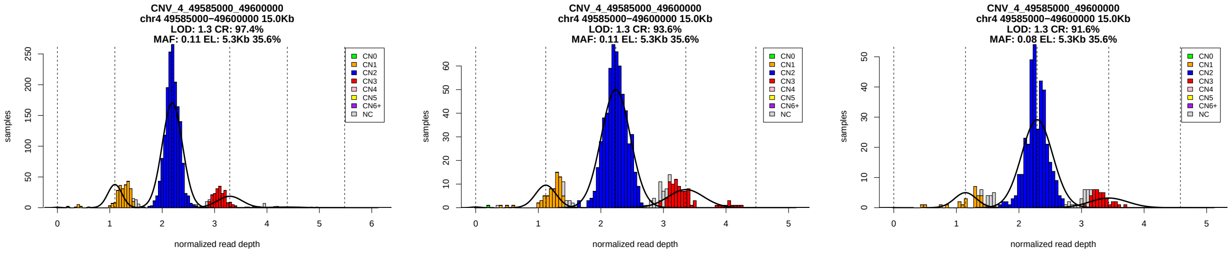


Figure 1: Copy number genotypes in the ASD, ID, and SCZ cohorts.

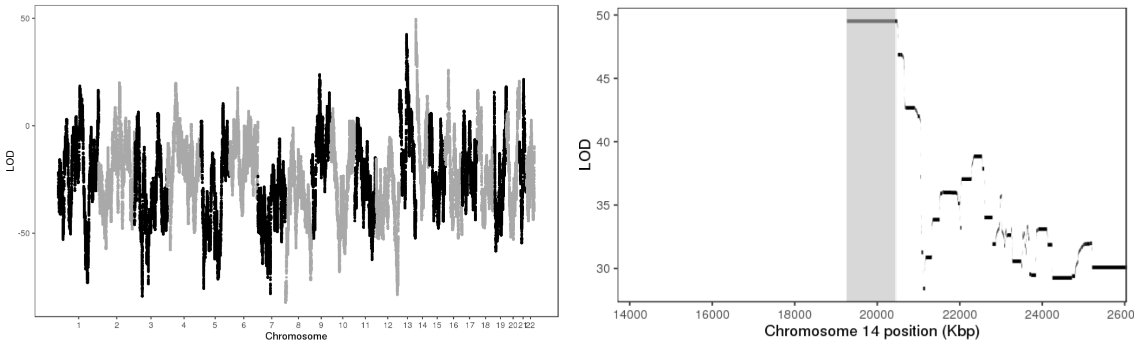


Figure 2: LOD scores of insertion locations across the genome (left) and around the 95% confidence interval (highlighted in gray, right).

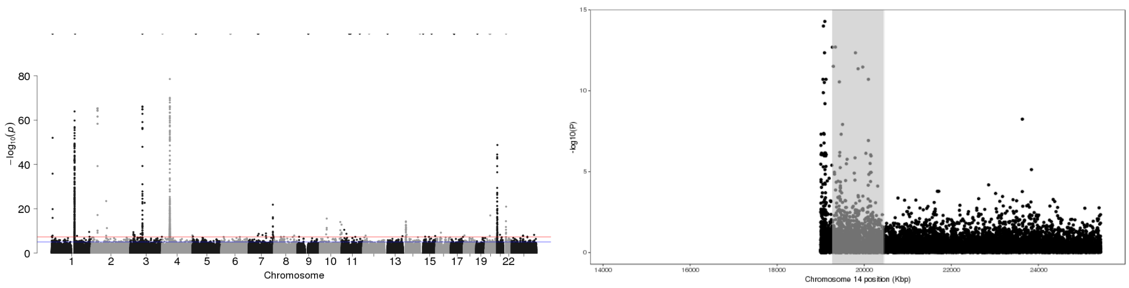


Figure 3: P-values of association between SNP genotypes and duplication copy number across the genome (left), and around the 95% confidence interval (highlighted in gray, right).

All coordinates refer to GRCh37.

### CNV_4_190541000_190680000

Insertion Location 95% Confidence Region:

14:19264866-20426258

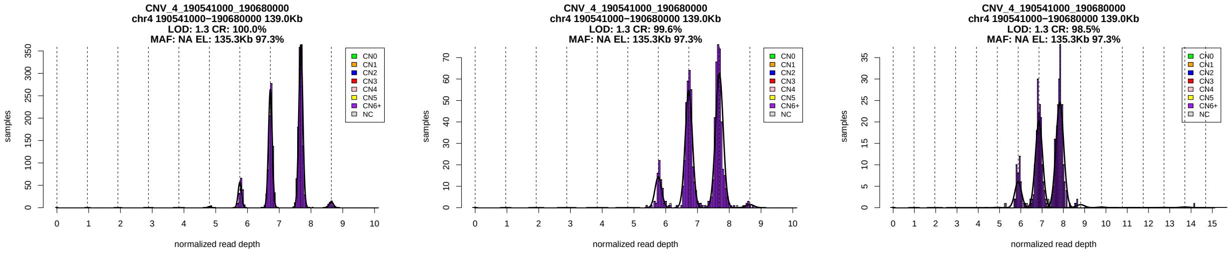


Figure 1: Copy number genotypes in the ASD, ID, and SCZ cohorts.

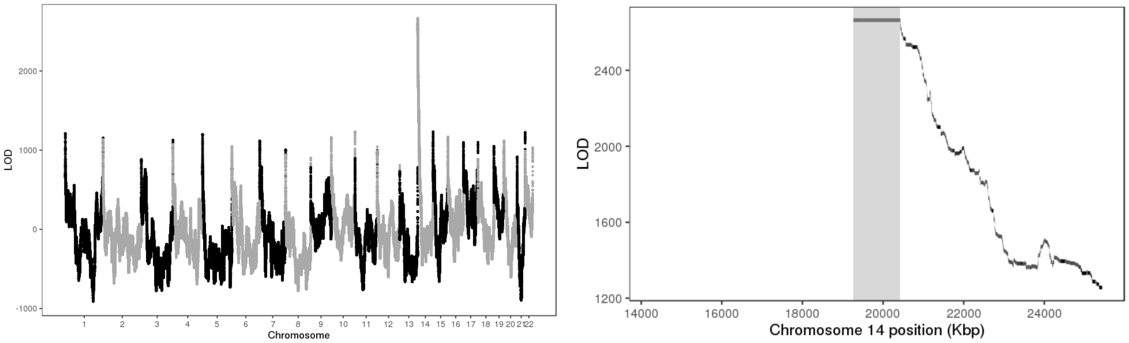


Figure 2: LOD scores of insertion locations across the genome (left) and around the 95% confidence interval (highlighted in gray, right).

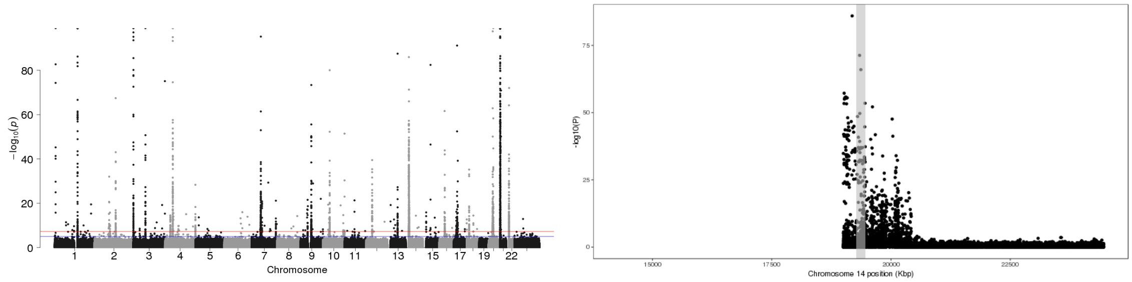


Figure 3: P-values of association between SNP genotypes and duplication copy number across the genome (left), and around the 95% confidence interval (highlighted in gray, right).

All coordinates refer to GRCh37.

### CNV_5_20832555_20834982

Insertion Location 95% Confidence Region:

5:34060053-34490529

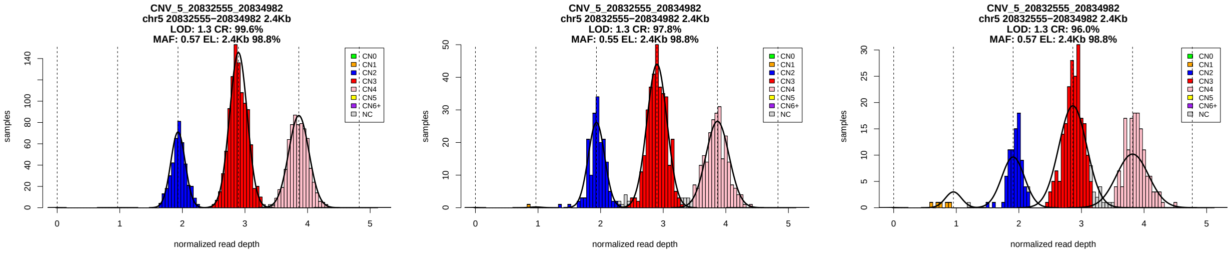


Figure 1: Copy number genotypes in the ASD, ID, and SCZ cohorts.

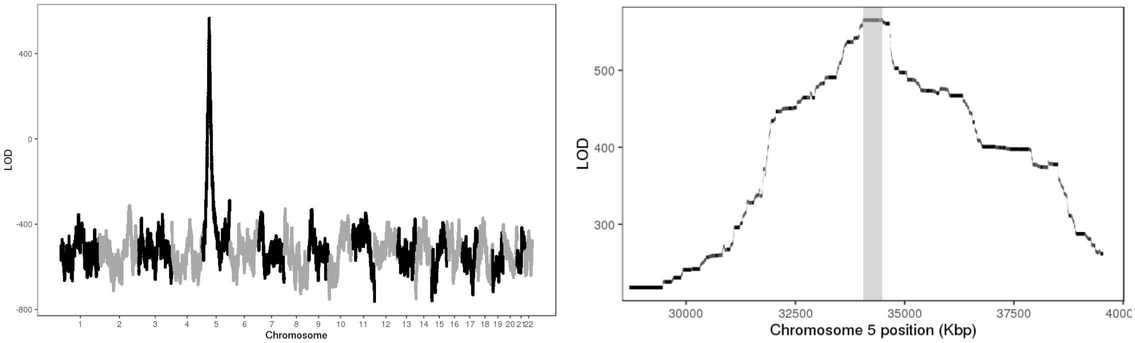


Figure 2: LOD scores of insertion locations across the genome (left) and around the 95% confidence interval (highlighted in gray, right).

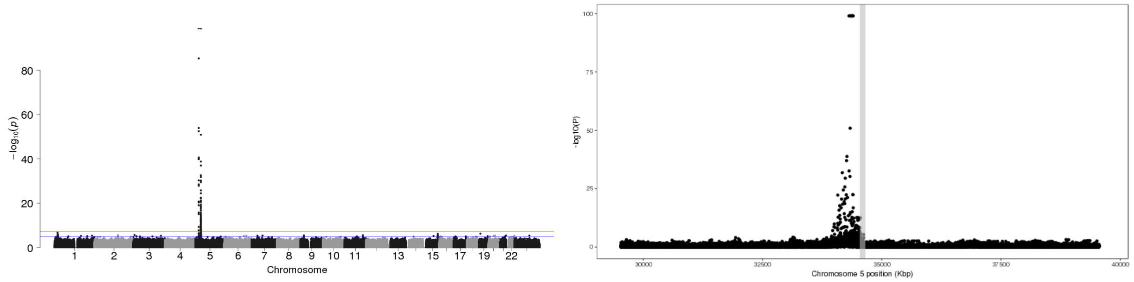


Figure 3: P-values of association between SNP genotypes and duplication copy number across the genome (left), and around the 95% confidence interval (highlighted in gray, right).

All coordinates refer to GRCh37.

### CNV_5_175437826_175440125

Insertion Location 95% Confidence Region:

5:176919656-177408337

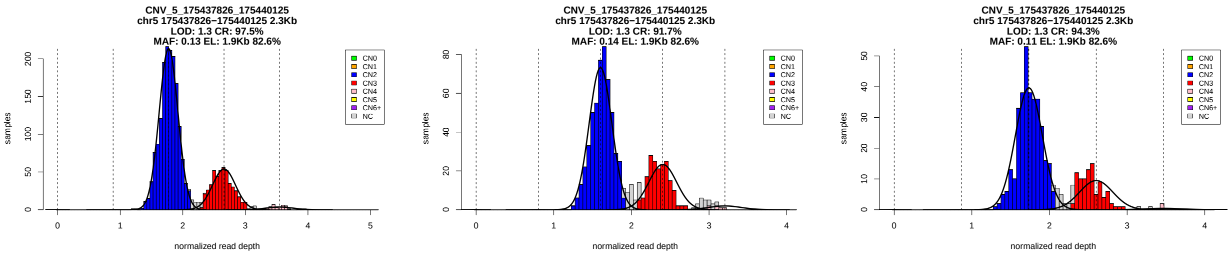


Figure 1: Copy number genotypes in the ASD, ID, and SCZ cohorts.

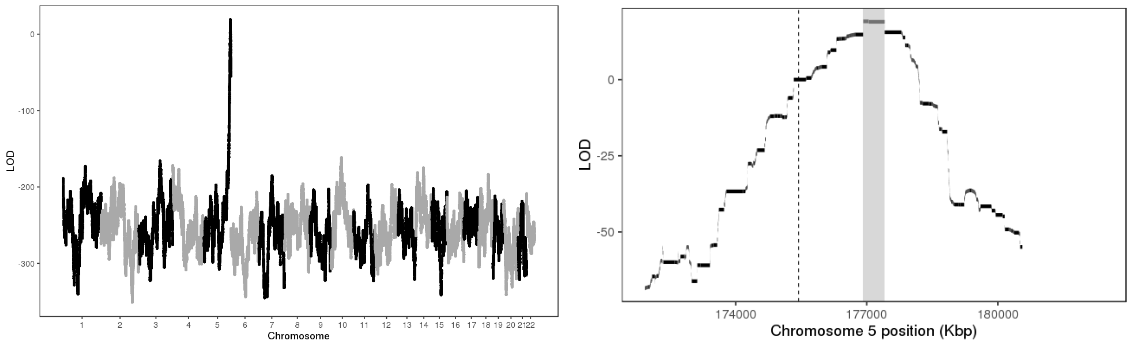


Figure 2: LOD scores of insertion locations across the genome (left) and around the 95% confidence interval (highlighted in gray, right).

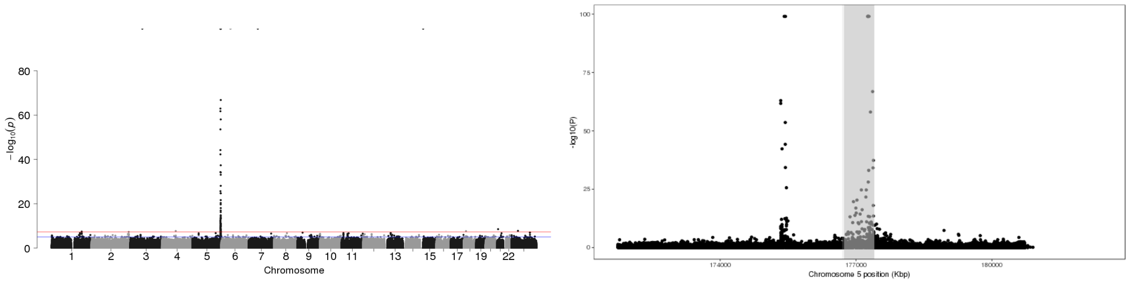


Figure 3: P-values of association between SNP genotypes and duplication copy number across the genome (left), and around the 95% confidence interval (highlighted in gray, right).

All coordinates refer to GRCh37.

### CNV_5_175465359_175471162

Insertion Location 95% Confidence Region:

5:176940274-177786775

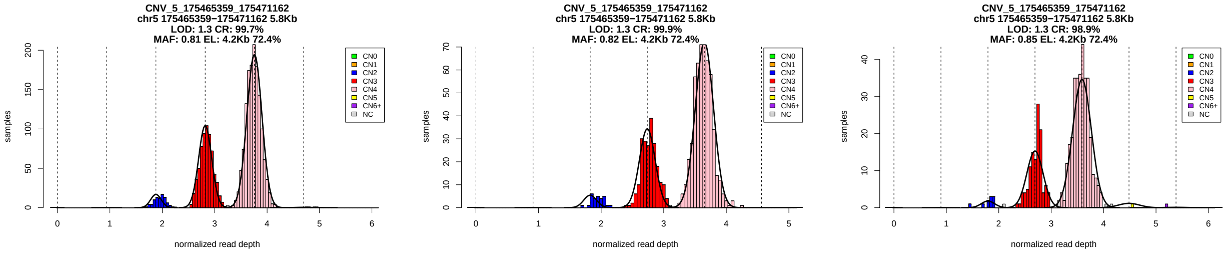


Figure 1: Copy number genotypes in the ASD, ID, and SCZ cohorts.

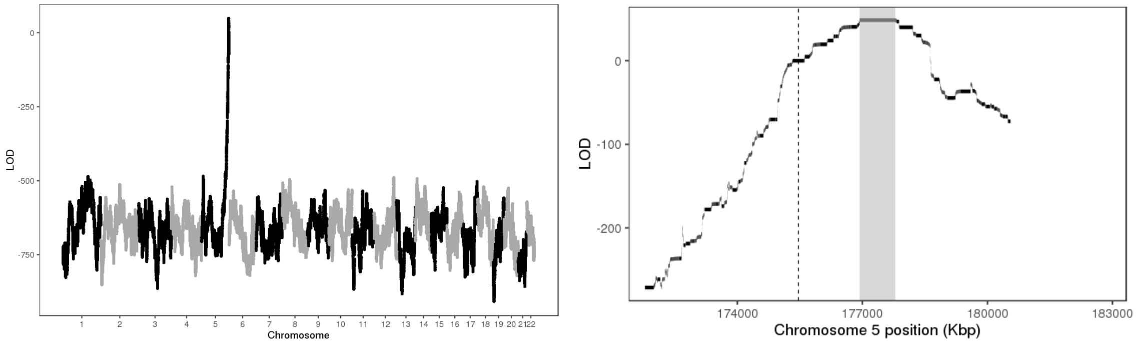


Figure 2: LOD scores of insertion locations across the genome (left) and around the 95% confidence interval (highlighted in gray, right).

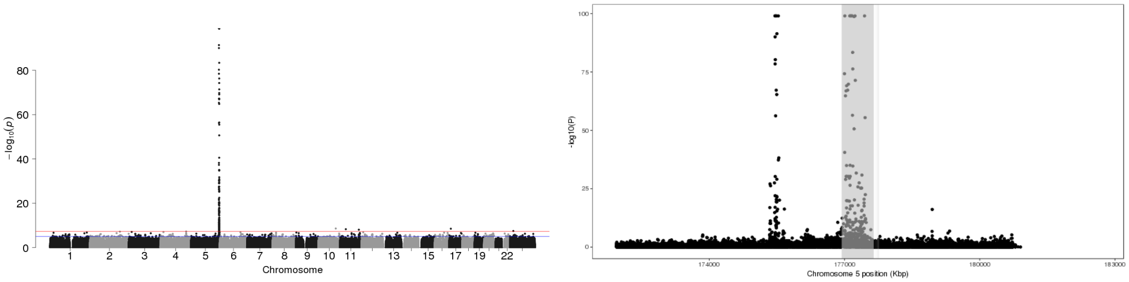


Figure 3: P-values of association between SNP genotypes and duplication copy number across the genome (left), and around the 95% confidence interval (highlighted in gray, right).

All coordinates refer to GRCh37.

### CNV_5_177217820_177221664

Insertion Location 95% Confidence Region:

5:175329033-176045010

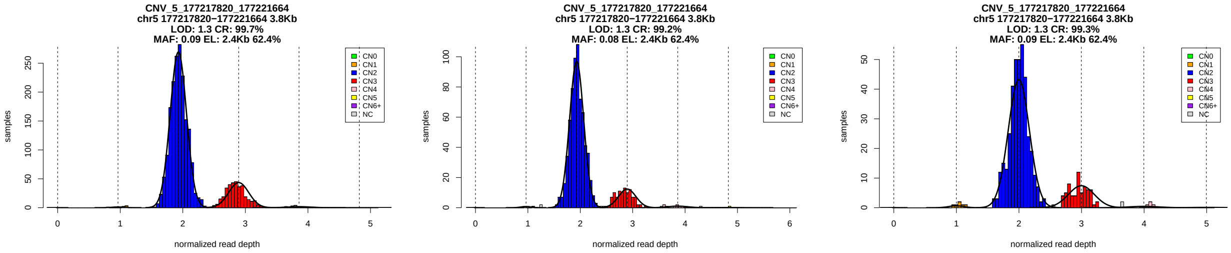


Figure 1: Copy number genotypes in the ASD, ID, and SCZ cohorts.

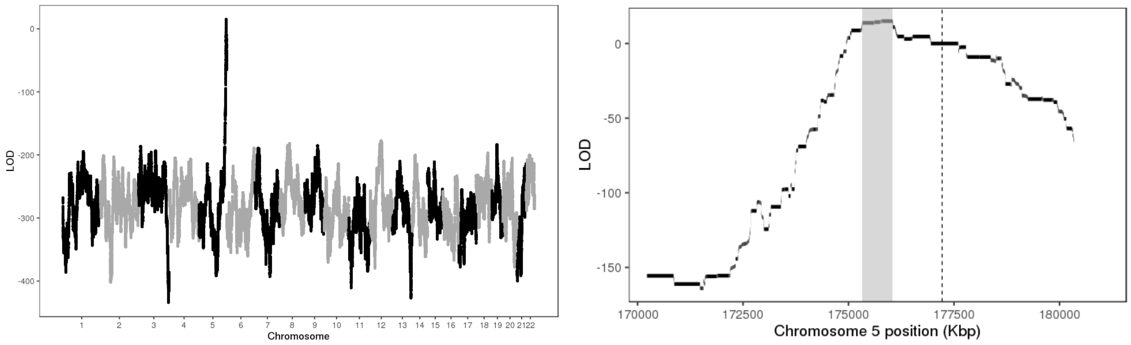


Figure 2: LOD scores of insertion locations across the genome (left) and around the 95% confidence interval (highlighted in gray, right).

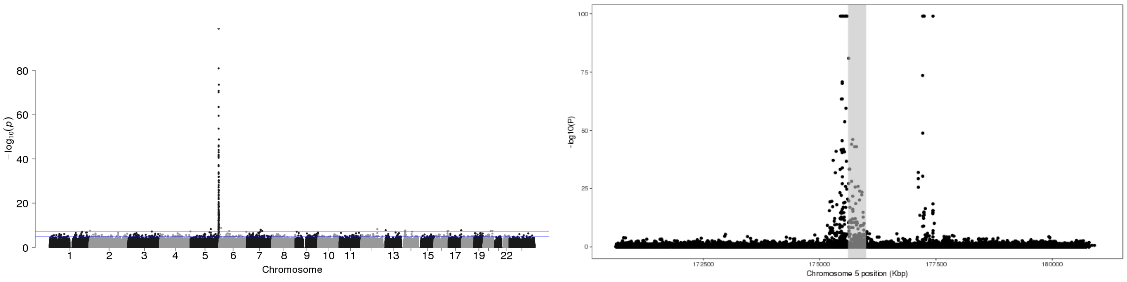


Figure 3: P-values of association between SNP genotypes and duplication copy number across the genome (left), and around the 95% confidence interval (highlighted in gray, right).

All coordinates refer to GRCh37.

### CNV_6_296000_365456

Insertion Location 95% Confidence Region:

16:34202307-46534652

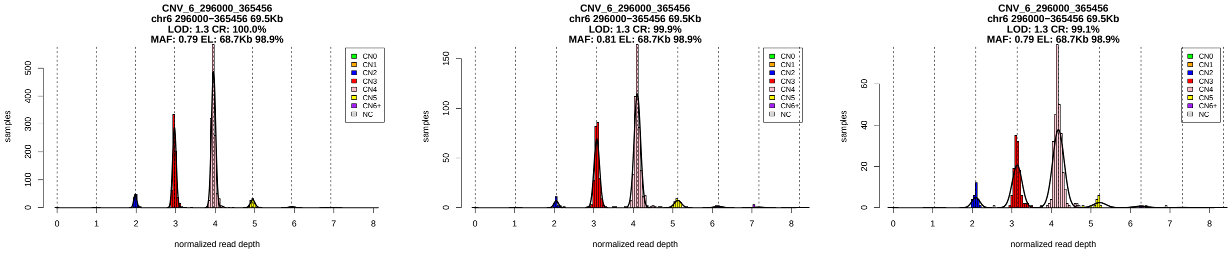


Figure 1: Copy number genotypes in the ASD, ID, and SCZ cohorts.

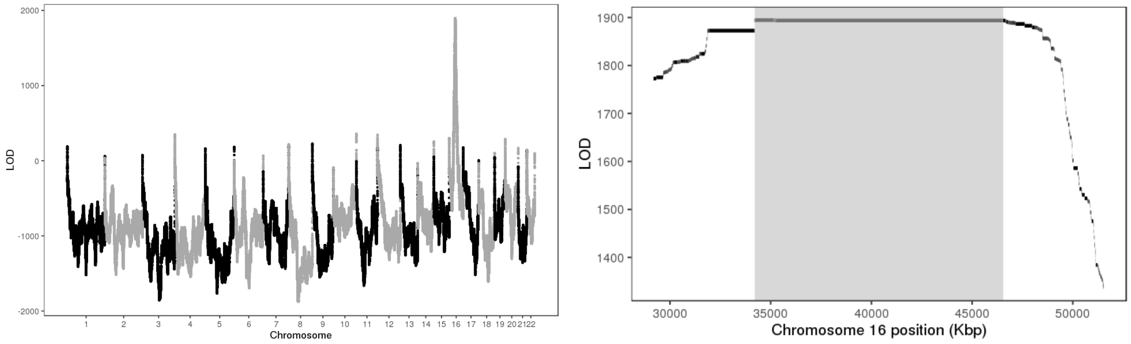


Figure 2: LOD scores of insertion locations across the genome (left) and around the 95% confidence interval (highlighted in gray, right).

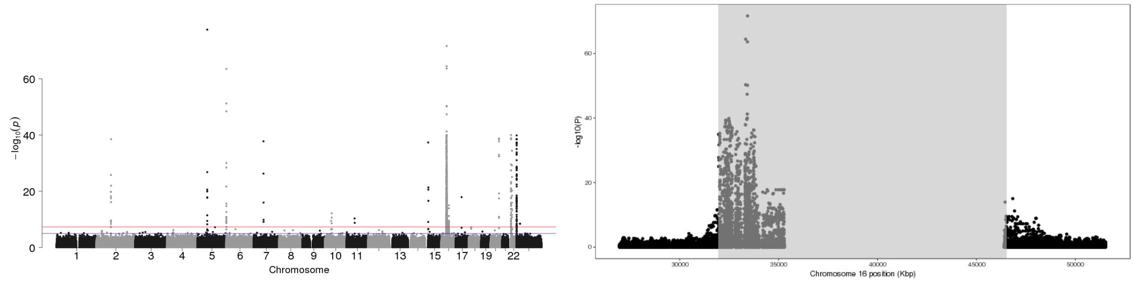


Figure 3: P-values of association between SNP genotypes and duplication copy number across the genome (left), and around the 95% confidence interval (highlighted in gray, right).

All coordinates refer to GRCh37.

### CNV_6_153742231_153747450

Insertion Location 95% Confidence Region:

6:160974428-161315678

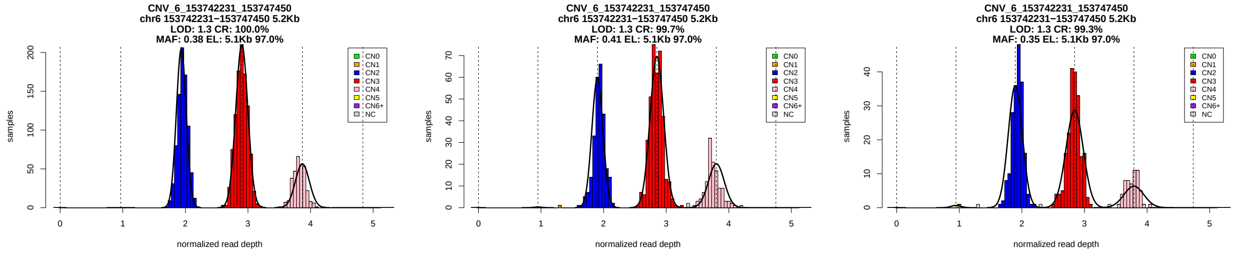


Figure 1: Copy number genotypes in the ASD, ID, and SCZ cohorts.

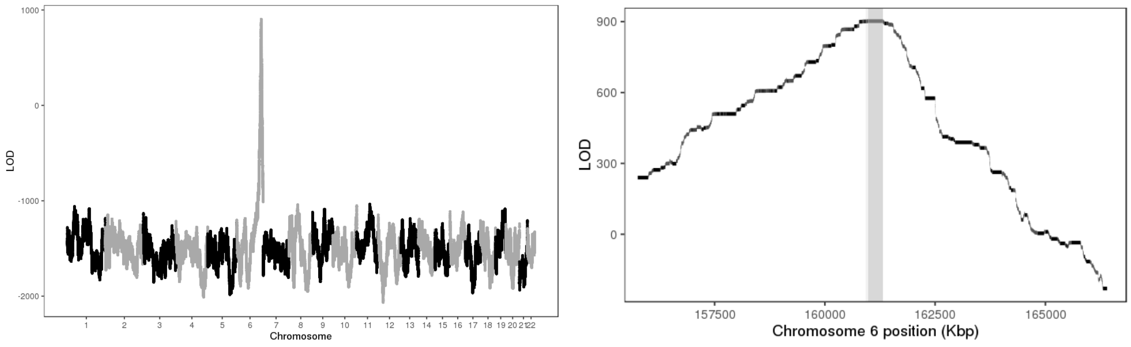


Figure 2: LOD scores of insertion locations across the genome (left) and around the 95% confidence interval (highlighted in gray, right).

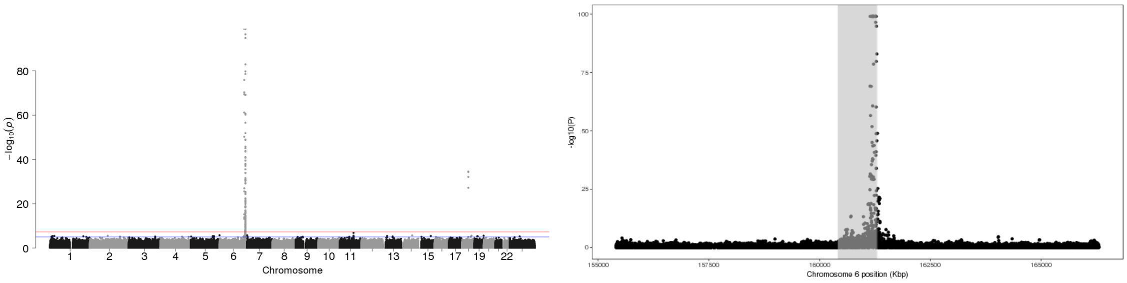


Figure 3: P-values of association between SNP genotypes and duplication copy number across the genome (left), and around the 95% confidence interval (highlighted in gray, right).

All coordinates refer to GRCh37.

### CNV_6_153748600_153752000

Insertion Location 95% Confidence Region:

6:160974428-161315678

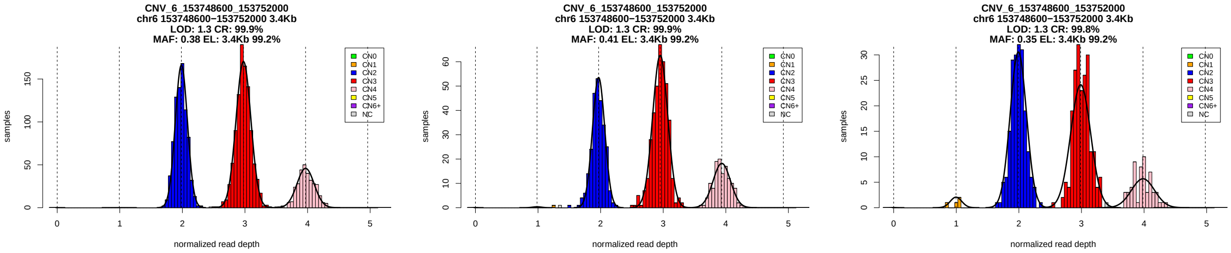


Figure 1: Copy number genotypes in the ASD, ID, and SCZ cohorts.

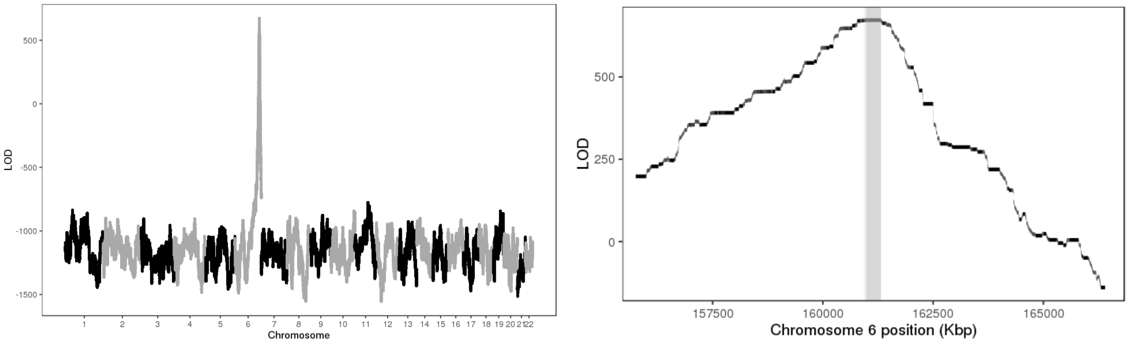


Figure 2: LOD scores of insertion locations across the genome (left) and around the 95% confidence interval (highlighted in gray, right).

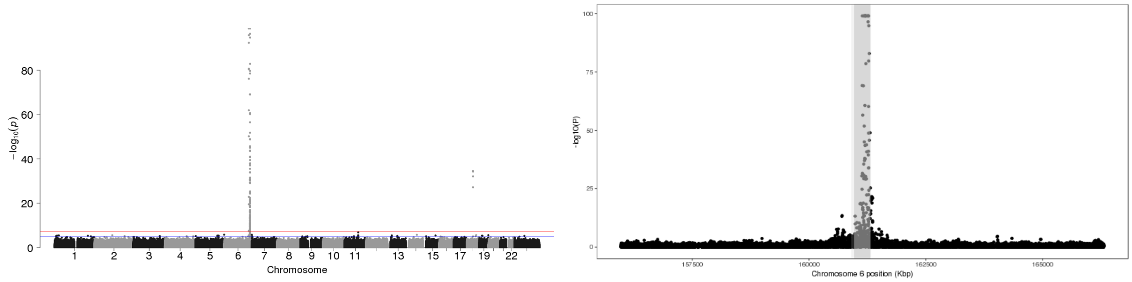


Figure 3: P-values of association between SNP genotypes and duplication copy number across the genome (left), and around the 95% confidence interval (highlighted in gray, right).

All coordinates refer to GRCh37.

### CNV_7_5940936_5951376

Insertion Location 95% Confidence Region:

7:6377836-7143781

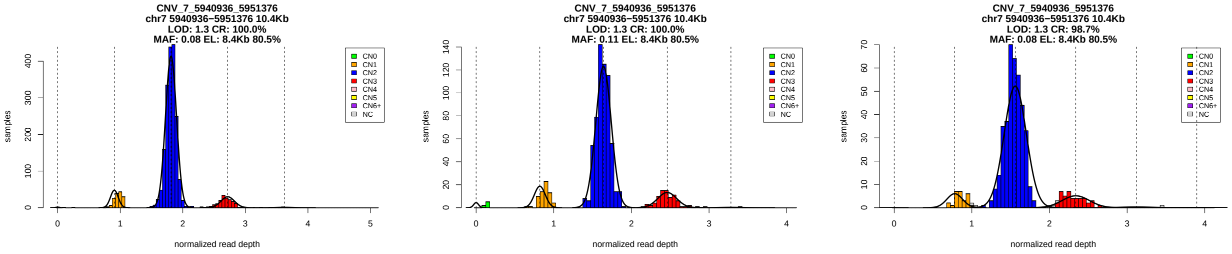


Figure 1: Copy number genotypes in the ASD, ID, and SCZ cohorts.

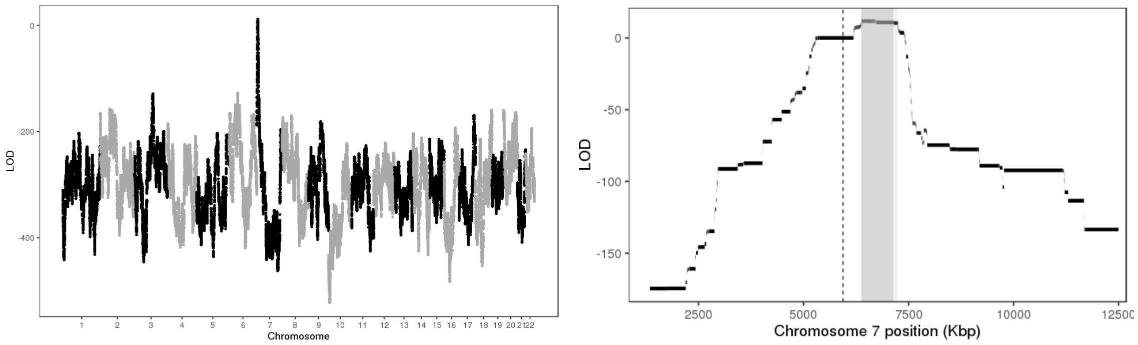


Figure 2: LOD scores of insertion locations across the genome (left) and around the 95% confidence interval (highlighted in gray, right).

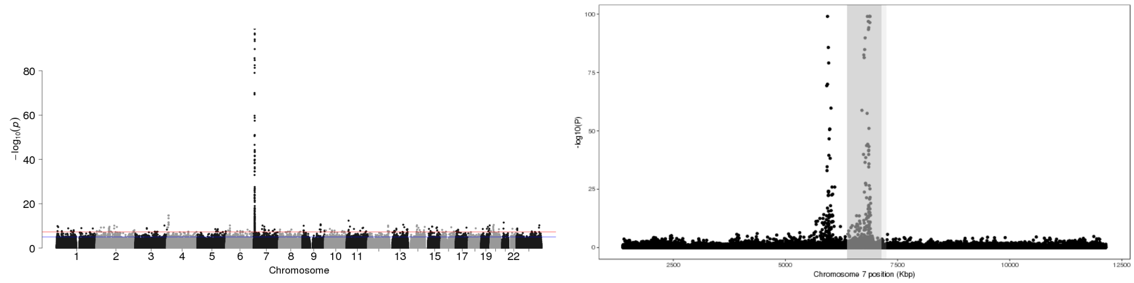


Figure 3: P-values of association between SNP genotypes and duplication copy number across the genome (left), and around the 95% confidence interval (highlighted in gray, right).

All coordinates refer to GRCh37.

### CNV_7_6852447_6863265

Insertion Location 95% Confidence Region:

7:5576680-6416615

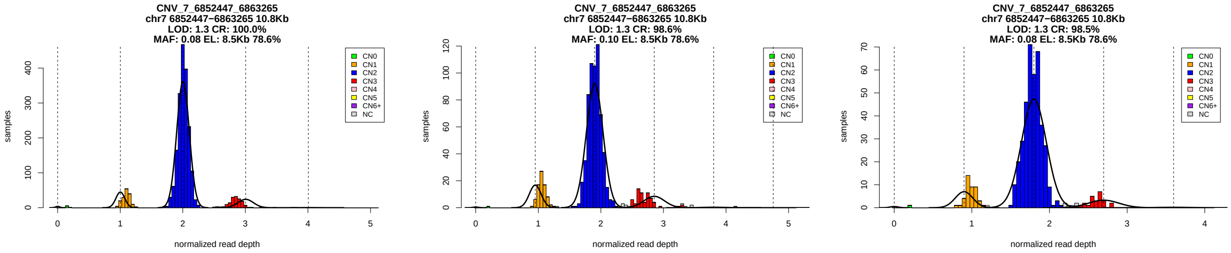


Figure 1: Copy number genotypes in the ASD, ID, and SCZ cohorts.

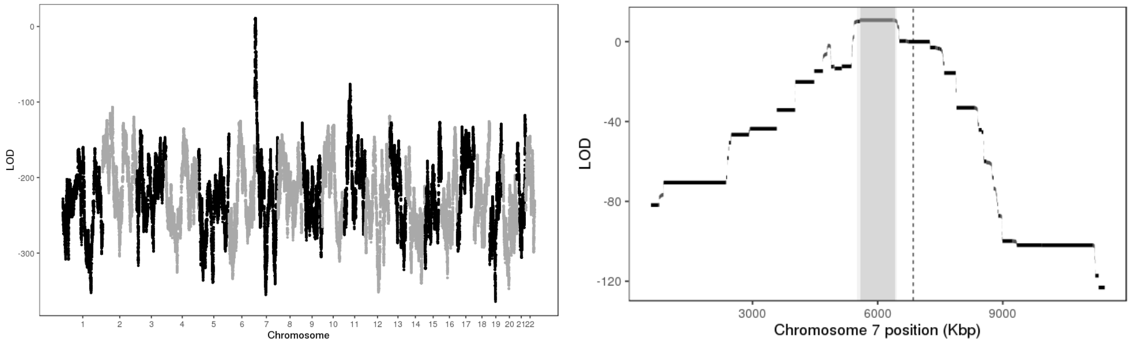


Figure 2: LOD scores of insertion locations across the genome (left) and around the 95% confidence interval (highlighted in gray, right).

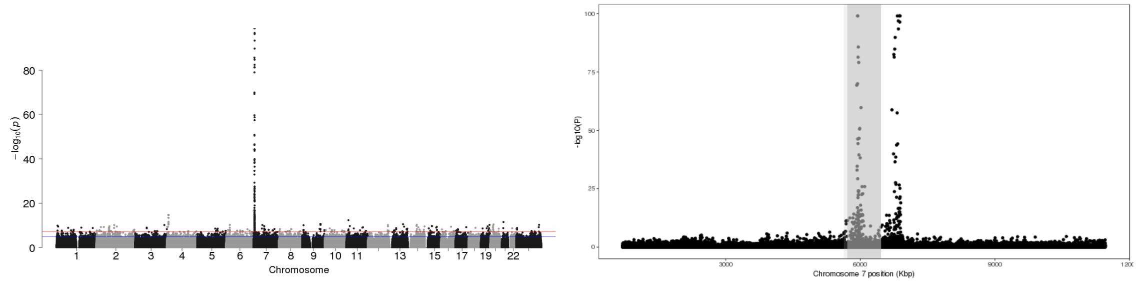


Figure 3: P-values of association between SNP genotypes and duplication copy number across the genome (left), and around the 95% confidence interval (highlighted in gray, right).

All coordinates refer to GRCh37.

### CNV_7_26241384_26252976_ENST00000409747.1

Insertion Location 95% Confidence Region:

15:40637622-40988418

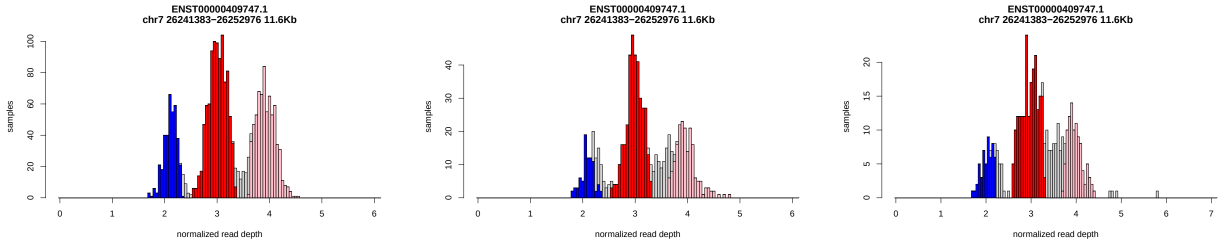


Figure 1: Copy number genotypes in the ASD, ID, and SCZ cohorts.

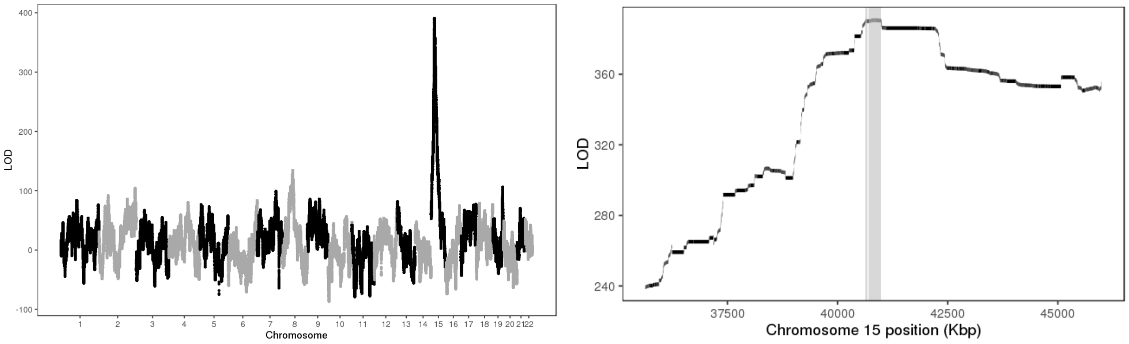


Figure 2: LOD scores of insertion locations across the genome (left) and around the 95% confidence interval (highlighted in gray, right).

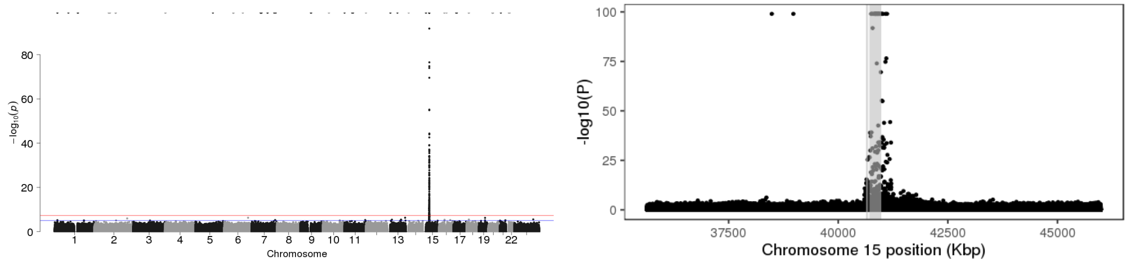


Figure 3: P-values of association between SNP genotypes and duplication copy number across the genome (left), and around the 95% confidence interval (highlighted in gray, right).

All coordinates refer to GRCh37.

### CNV_7_56557397_56564351

Insertion Location 95% Confidence Region:

7:57433043-62504090

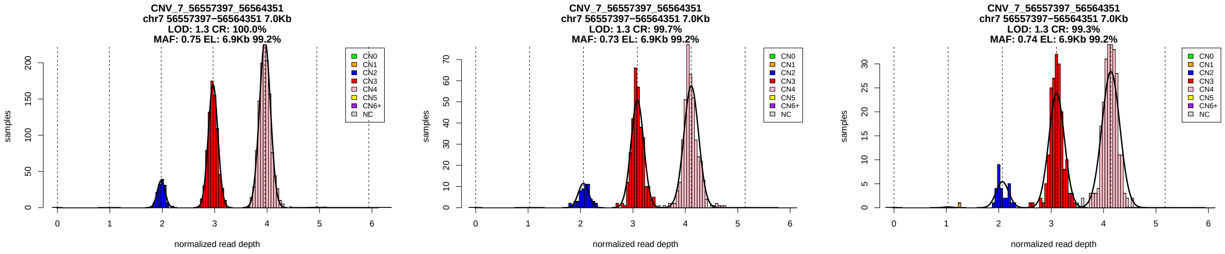


Figure 1: Copy number genotypes in the ASD, ID, and SCZ cohorts.

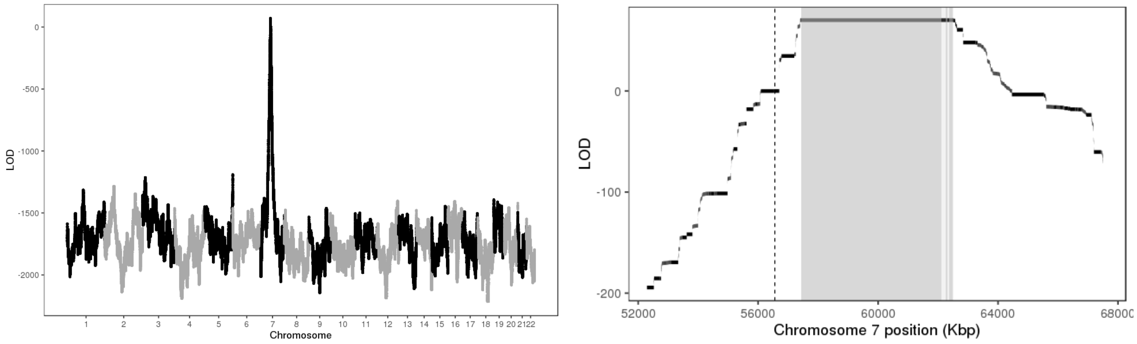


Figure 2: LOD scores of insertion locations across the genome (left) and around the 95% confidence interval (highlighted in gray, right).

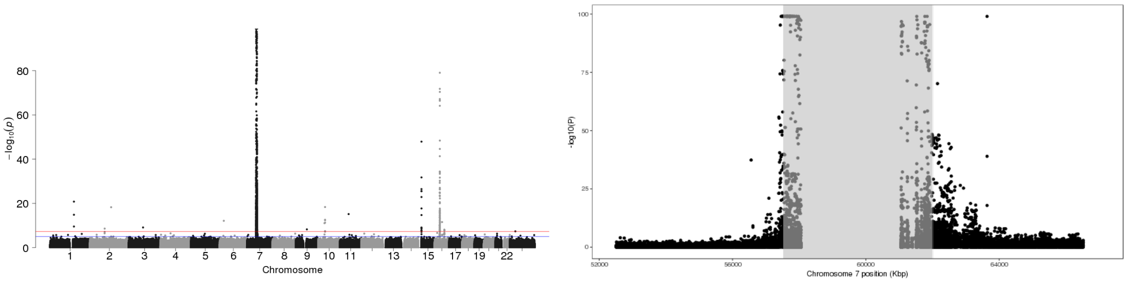


Figure 3: P-values of association between SNP genotypes and duplication copy number across the genome (left), and around the 95% confidence interval (highlighted in gray, right).

All coordinates refer to GRCh37.

### CNV_7_57128276_57131575

Insertion Location 95% Confidence Region:

7:62668534-63313180

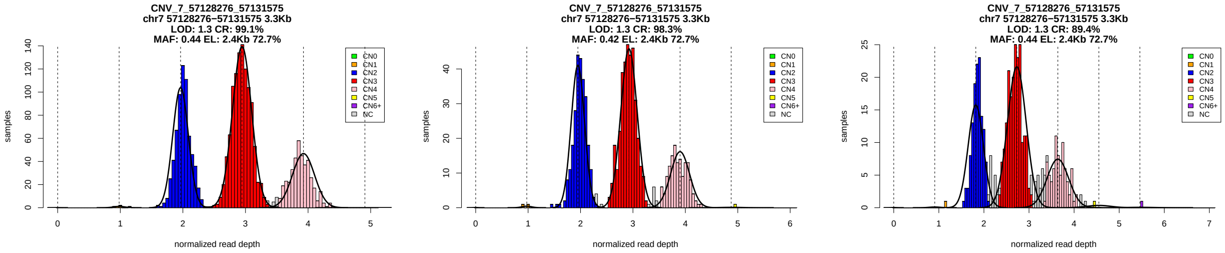


Figure 1: Copy number genotypes in the ASD, ID, and SCZ cohorts.

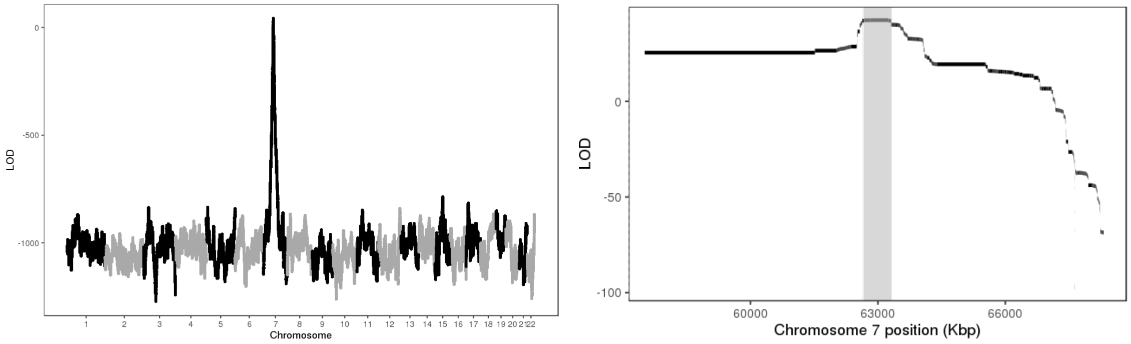


Figure 2: LOD scores of insertion locations across the genome (left) and around the 95% confidence interval (highlighted in gray, right).

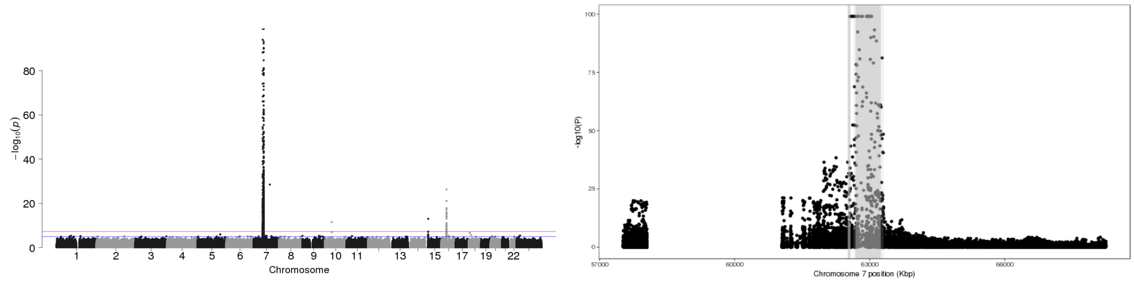


Figure 3: P-values of association between SNP genotypes and duplication copy number across the genome (left), and around the 95% confidence interval (highlighted in gray, right).

All coordinates refer to GRCh37.

### CNV_7_62979567_62986587

Insertion Location 95% Confidence Region:

7:56516052-57350723

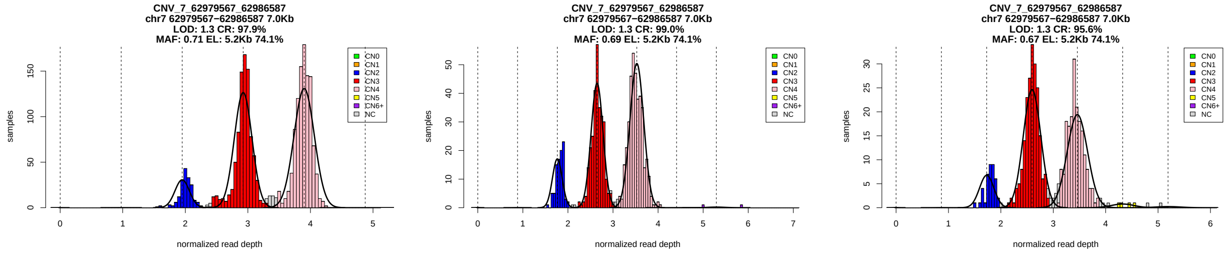


Figure 1: Copy number genotypes in the ASD, ID, and SCZ cohorts.

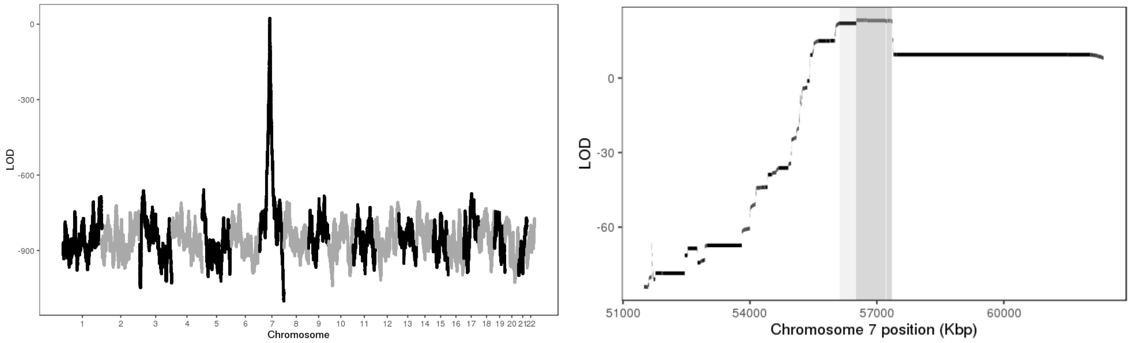


Figure 2: LOD scores of insertion locations across the genome (left) and around the 95% confidence interval (highlighted in gray, right).

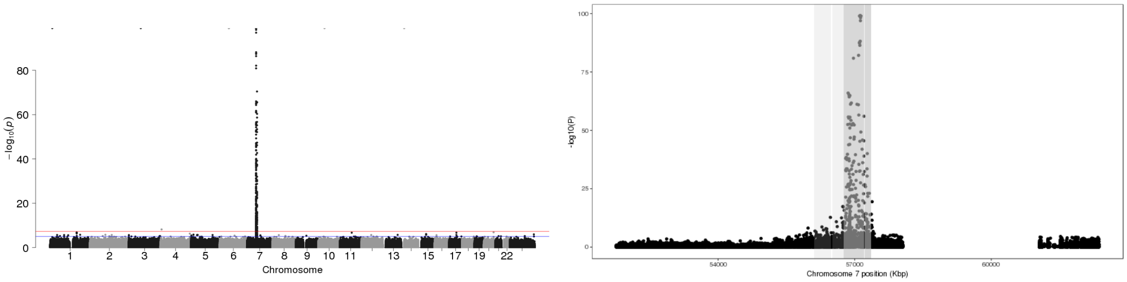


Figure 3: P-values of association between SNP genotypes and duplication copy number across the genome (left), and around the 95% confidence interval (highlighted in gray, right).

All coordinates refer to GRCh37.

### CNV_7_63026573_63041272

Insertion Location 95% Confidence Region:

7:56115321-57353425

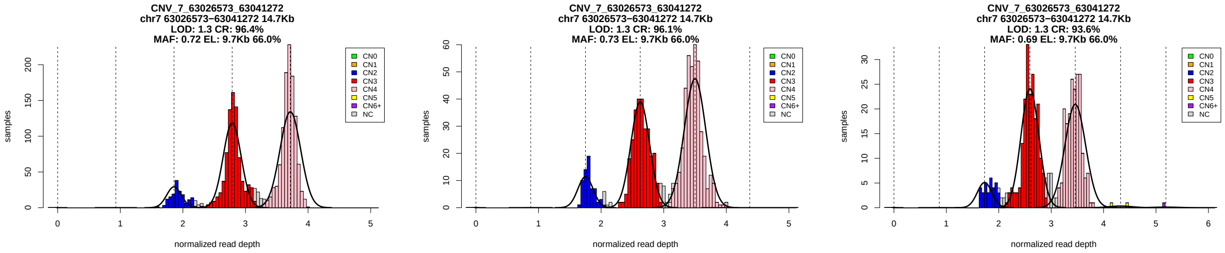


Figure 1: Copy number genotypes in the ASD, ID, and SCZ cohorts.

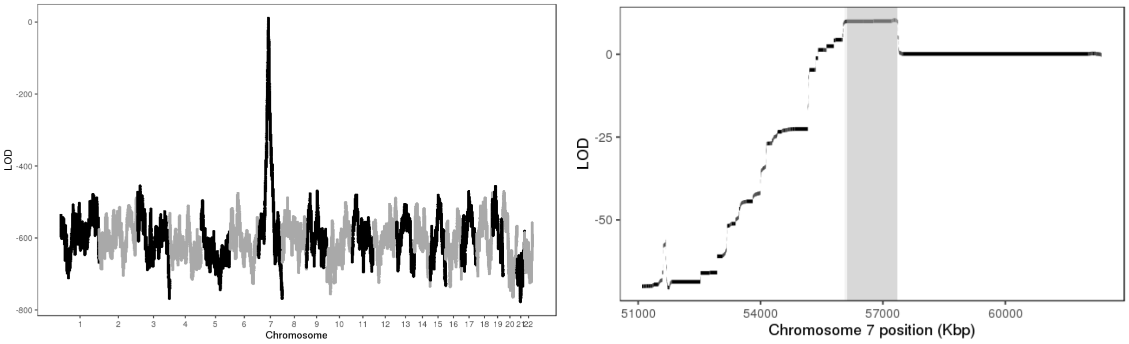


Figure 2: LOD scores of insertion locations across the genome (left) and around the 95% confidence interval (highlighted in gray, right).

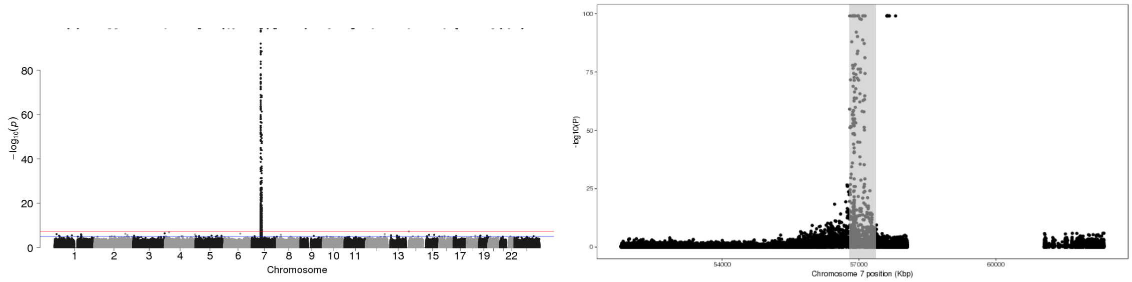


Figure 3: P-values of association between SNP genotypes and duplication copy number across the genome (left), and around the 95% confidence interval (highlighted in gray, right).

All coordinates refer to GRCh37.

### CNV_7_64551729_64554250

Insertion Location 95% Confidence Region:

7:55615076-56513828

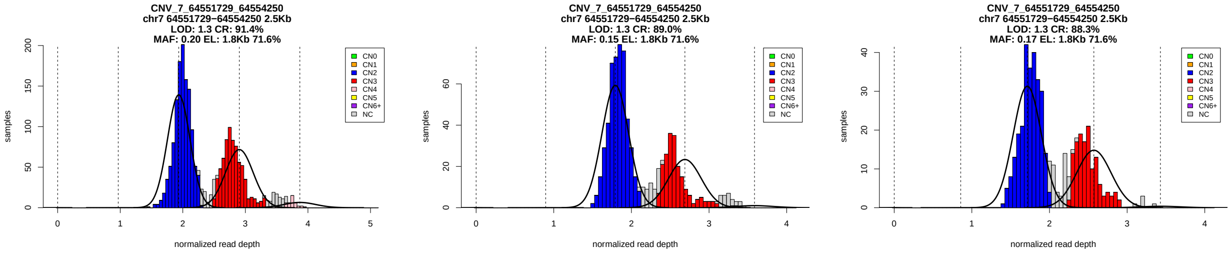


Figure 1: Copy number genotypes in the ASD, ID, and SCZ cohorts.

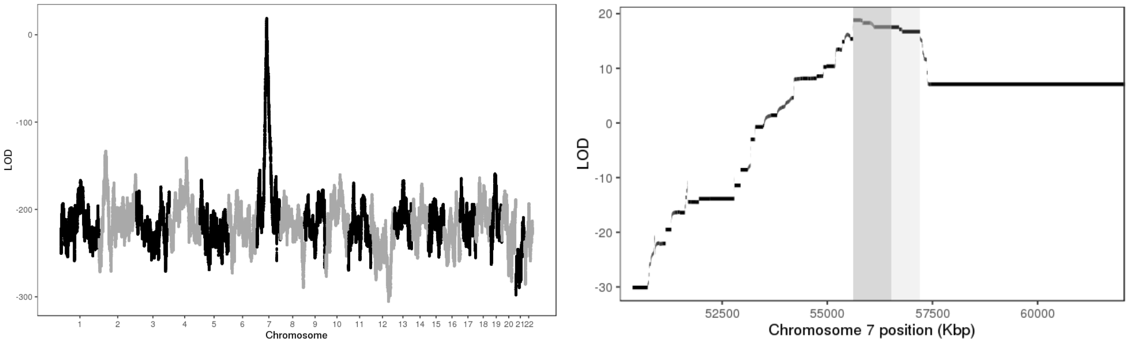


Figure 2: LOD scores of insertion locations across the genome (left) and around the 95% confidence interval (highlighted in gray, right).

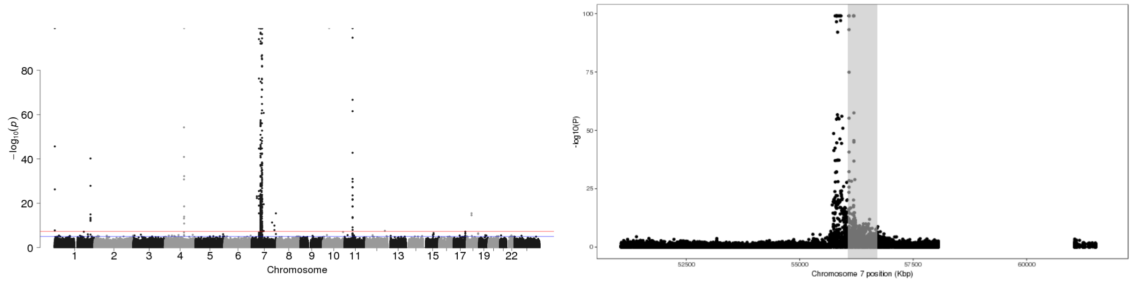


Figure 3: P-values of association between SNP genotypes and duplication copy number across the genome (left), and around the 95% confidence interval (highlighted in gray, right).

All coordinates refer to GRCh37.

### CNV_7_65259784_65287967

Insertion Location 95% Confidence Region:

7:55617815-56704005

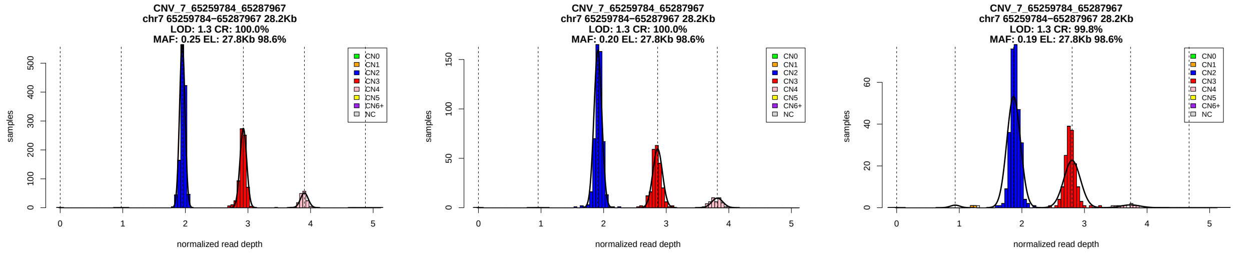


Figure 1: Copy number genotypes in the ASD, ID, and SCZ cohorts.

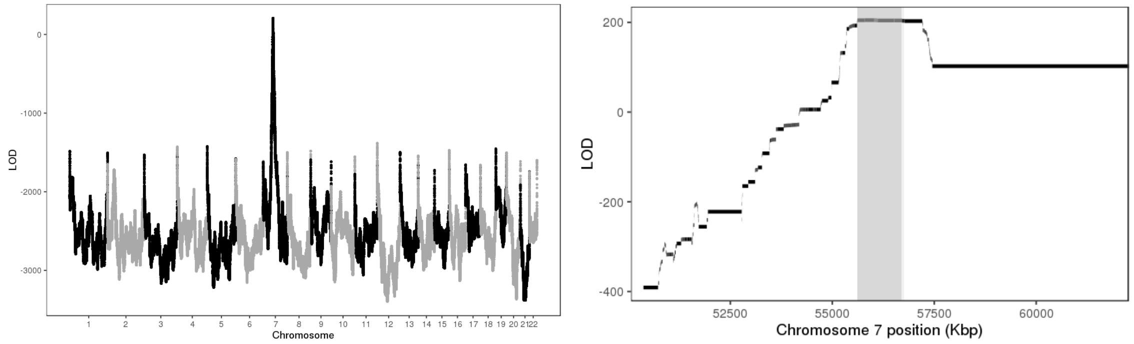


Figure 2: LOD scores of insertion locations across the genome (left) and around the 95% confidence interval (highlighted in gray, right).

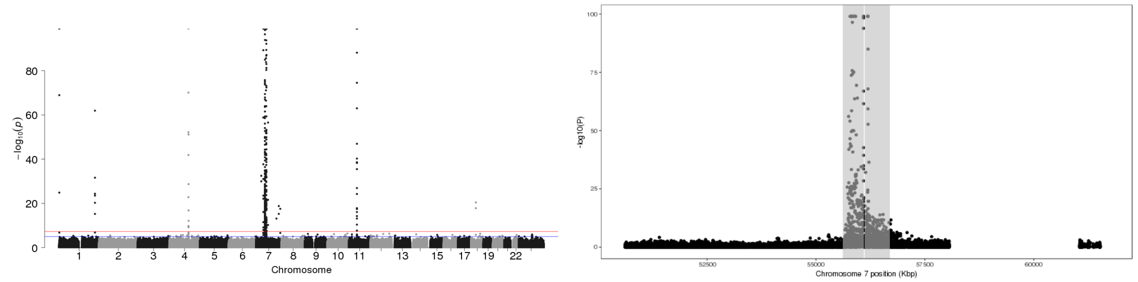


Figure 3: P-values of association between SNP genotypes and duplication copy number across the genome (left), and around the 95% confidence interval (highlighted in gray, right).

All coordinates refer to GRCh37.

### CNV_7_66438701_66441000

Insertion Location 95% Confidence Region:

7:71321075-73590906

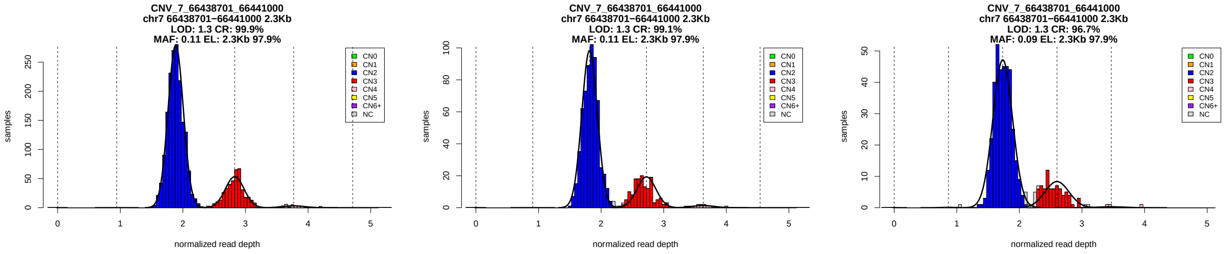


Figure 1: Copy number genotypes in the ASD, ID, and SCZ cohorts.

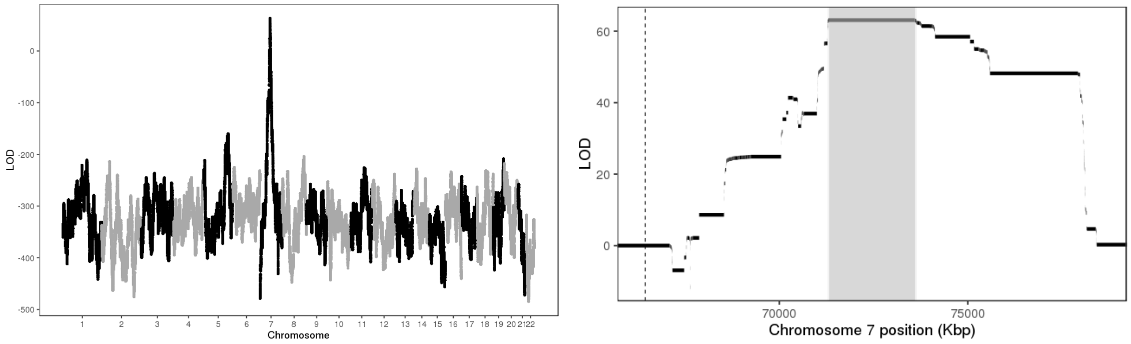


Figure 2: LOD scores of insertion locations across the genome (left) and around the 95% confidence interval (highlighted in gray, right).

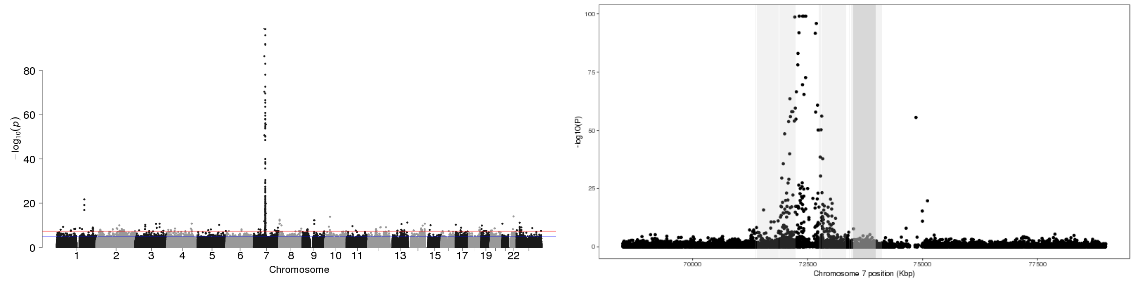


Figure 3: P-values of association between SNP genotypes and duplication copy number across the genome (left), and around the 95% confidence interval (highlighted in gray, right).

All coordinates refer to GRCh37.

### CNV_7_153687900_153705000

Insertion Location 95% Confidence Region:

7:141851540-150205086

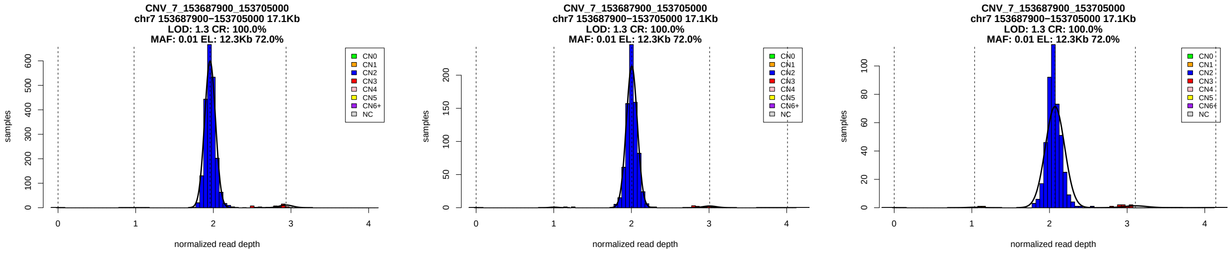


Figure 1: Copy number genotypes in the ASD, ID, and SCZ cohorts.

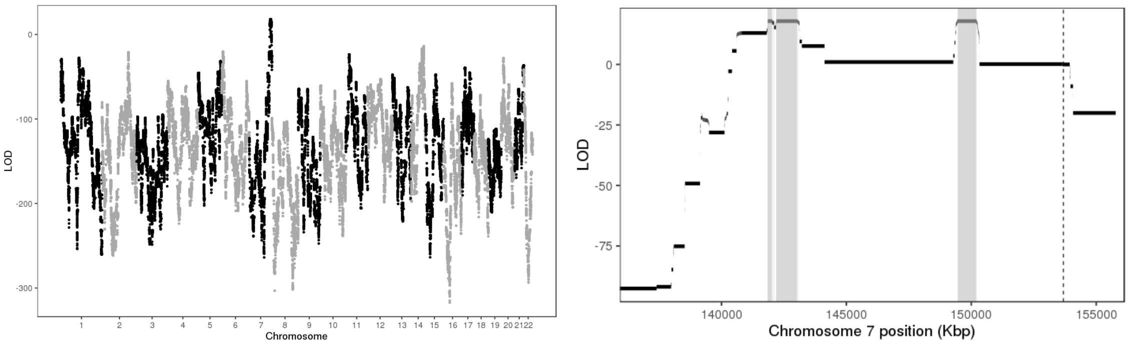


Figure 2: LOD scores of insertion locations across the genome (left) and around the 95% confidence interval (highlighted in gray, right).

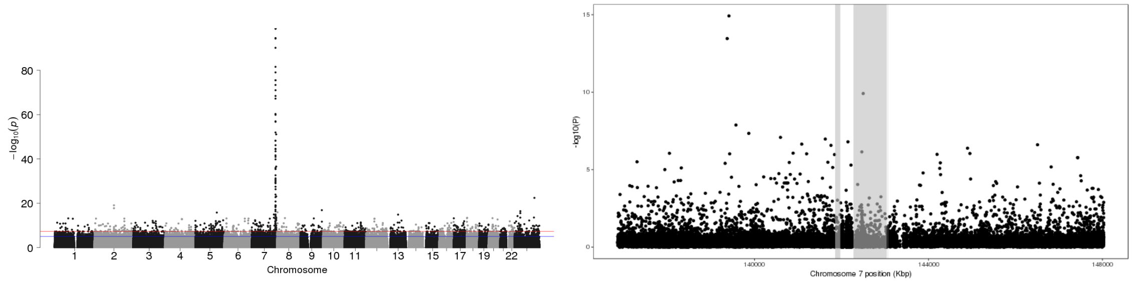


Figure 3: P-values of association between SNP genotypes and duplication copy number across the genome (left), and around the 95% confidence interval (highlighted in gray, right).

All coordinates refer to GRCh37.

### CNV_7_153794100_153796250

Insertion Location 95% Confidence Region:

7:149162885-150004244

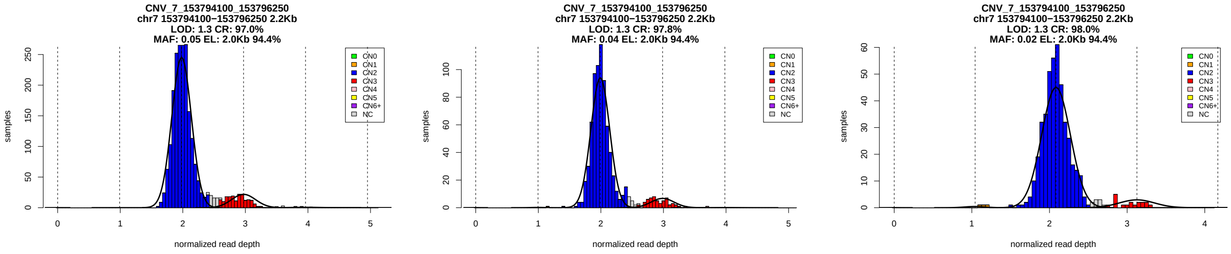


Figure 1: Copy number genotypes in the ASD, ID, and SCZ cohorts.

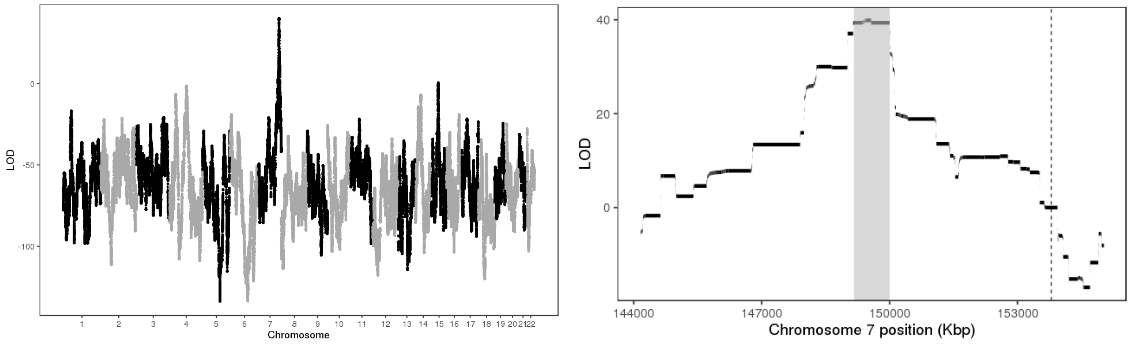


Figure 2: LOD scores of insertion locations across the genome (left) and around the 95% confidence interval (highlighted in gray, right).

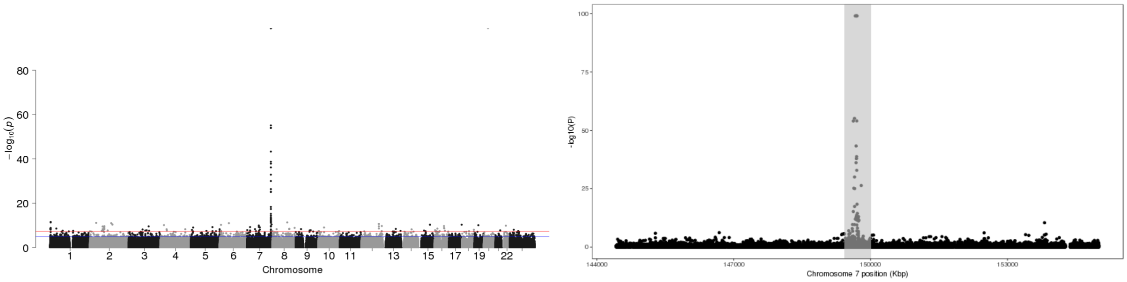


Figure 3: P-values of association between SNP genotypes and duplication copy number across the genome (left), and around the 95% confidence interval (highlighted in gray, right).

All coordinates refer to GRCh37.

### CNV_8_29920641_29940659_ENST00000545648.1

Insertion Location 95% Confidence Region:

1:187663894-193484533

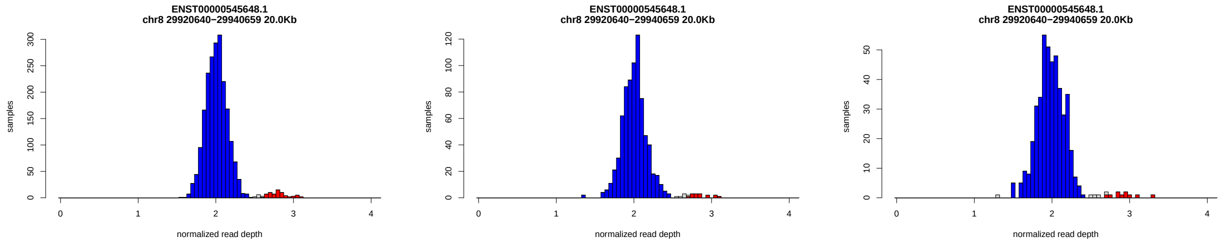


Figure 1: Copy number genotypes in the ASD, ID, and SCZ cohorts.

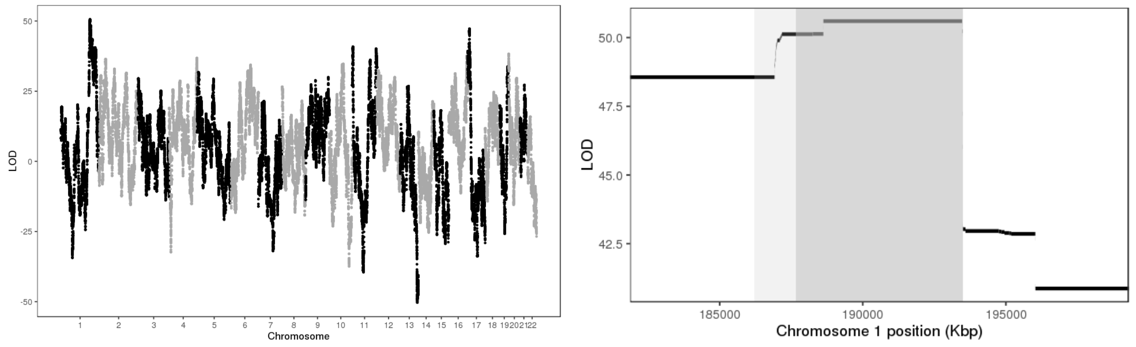


Figure 2: LOD scores of insertion locations across the genome (left) and around the 95% confidence interval (highlighted in gray, right).

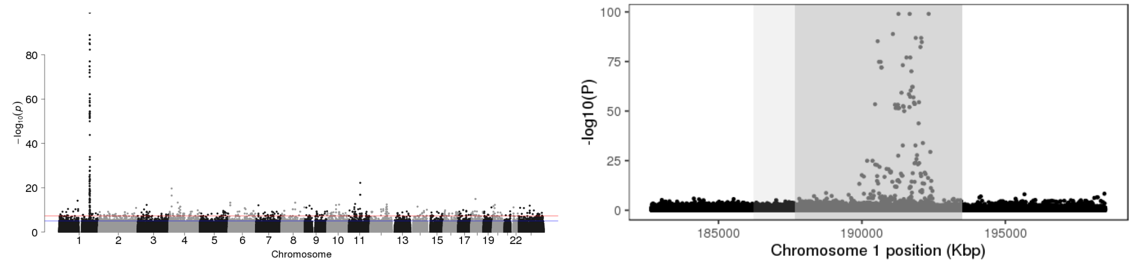


Figure 3: P-values of association between SNP genotypes and duplication copy number across the genome (left), and around the 95% confidence interval (highlighted in gray, right).

All coordinates refer to GRCh37.

# GS_SD_M2_8_7200001_7436083_8_7600001_7825413

Insertion Location 95% Confidence Region:

8:11860259-12578645

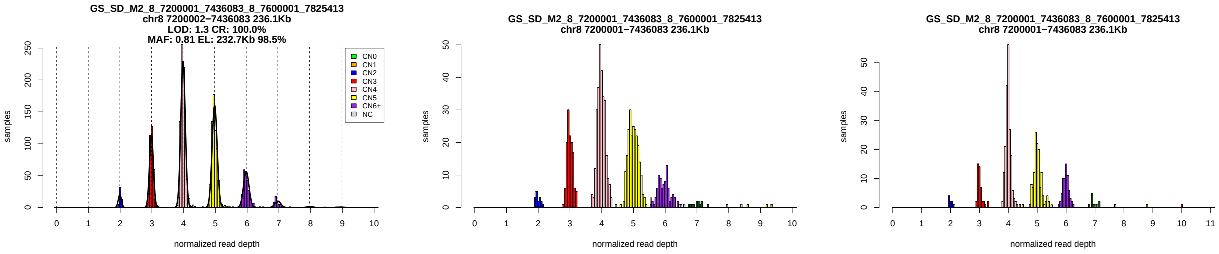


Figure 1: Copy number genotypes in the ASD, ID, and SCZ cohorts.

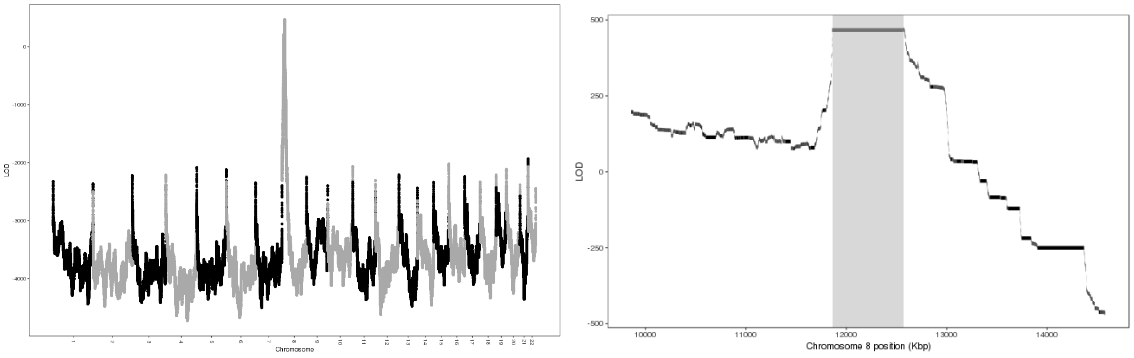


Figure 2: LOD scores of insertion locations across the genome (left) and around the 95% confidence interval (highlighted in gray, right).

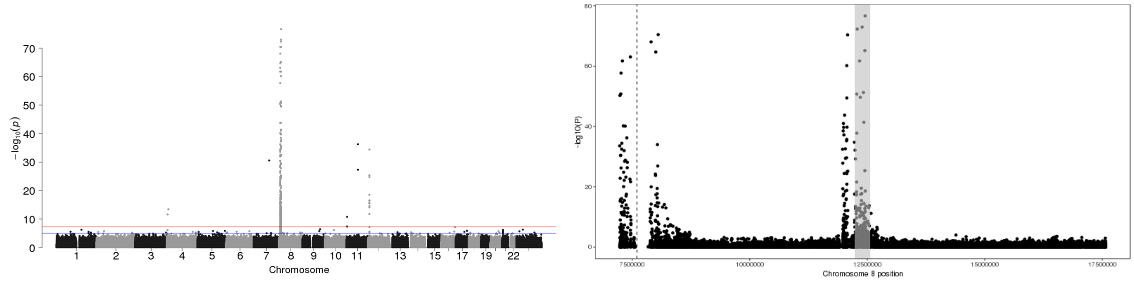


Figure 3: P-values of association between SNP genotypes and duplication copy number across the genome (left), and around the 95% confidence interval (highlighted in gray, right).

All coordinates refer to GRCh37.

### CNV_8_7211500_7227500

Insertion Location 95% Confidence Region:

8:12577657-12578672

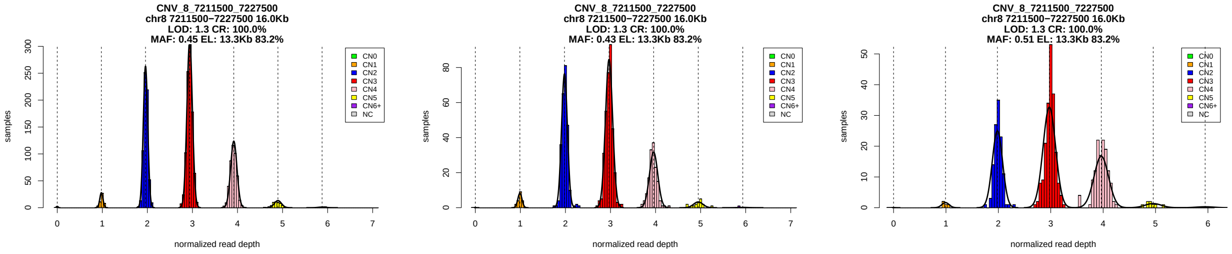


Figure 1: Copy number genotypes in the ASD, ID, and SCZ cohorts.

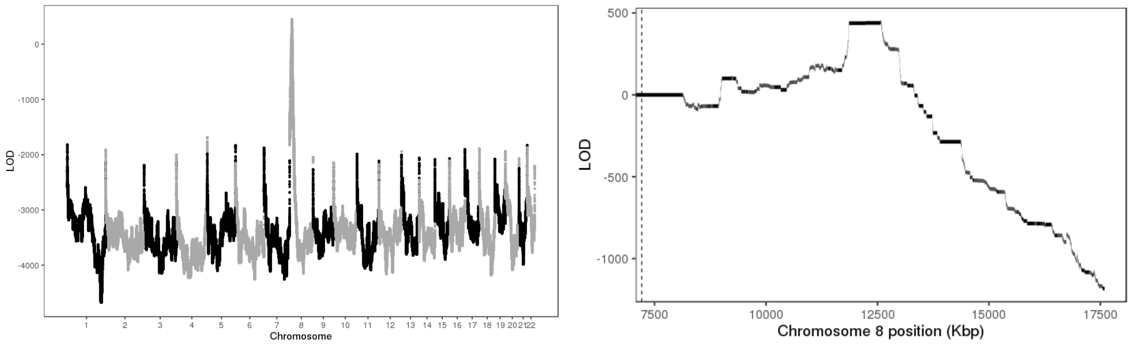


Figure 2: LOD scores of insertion locations across the genome (left) and around the 95% confidence interval (highlighted in gray, right).

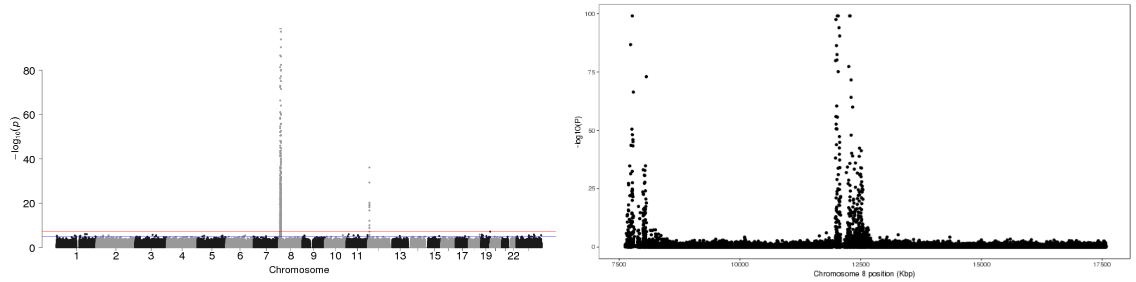


Figure 3: P-values of association between SNP genotypes and duplication copy number across the genome (left), and around the 95% confidence interval (highlighted in gray, right).

All coordinates refer to GRCh37.

### CNV_8_7276000_7280100

Insertion Location 95% Confidence Region:

8:12584092-12585436

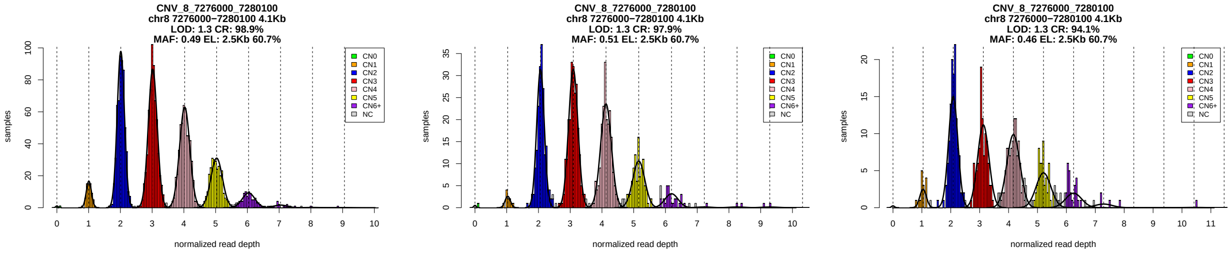


Figure 1: Copy number genotypes in the ASD, ID, and SCZ cohorts.

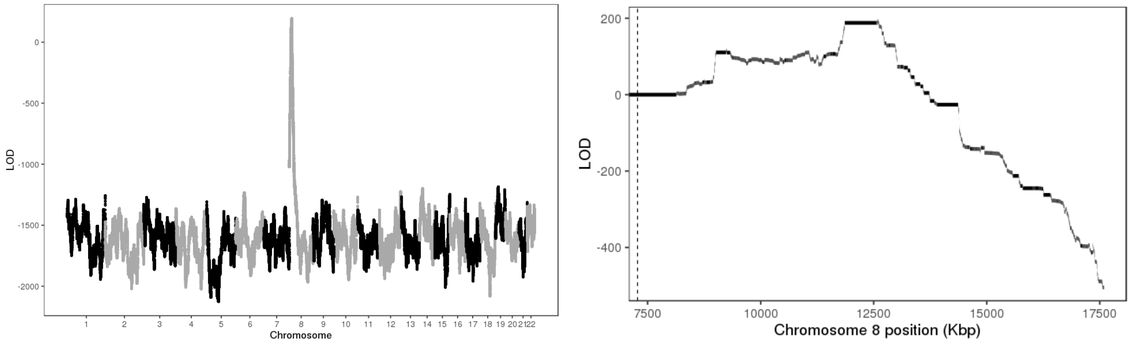


Figure 2: LOD scores of insertion locations across the genome (left) and around the 95% confidence interval (highlighted in gray, right).

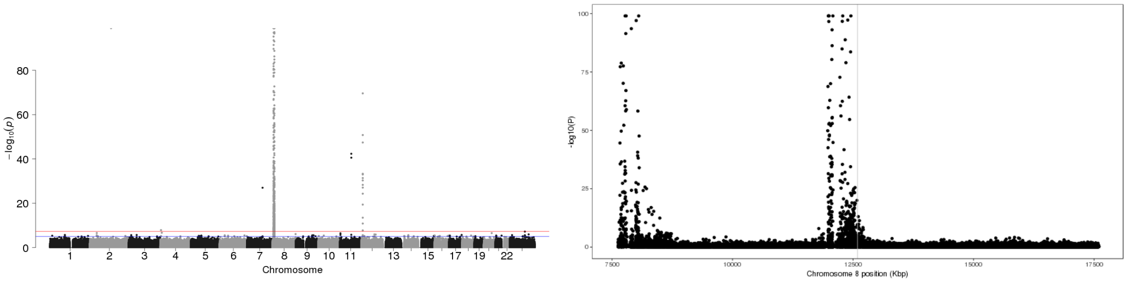


Figure 3: P-values of association between SNP genotypes and duplication copy number across the genome (left), and around the 95% confidence interval (highlighted in gray, right).

All coordinates refer to GRCh37.

### CNV_8_7746183_7750670

Insertion Location 95% Confidence Region:

8:11714941-11828200

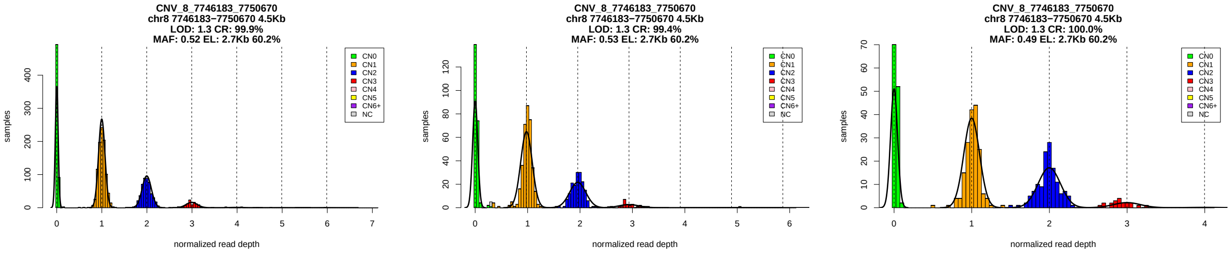


Figure 1: Copy number genotypes in the ASD, ID, and SCZ cohorts.

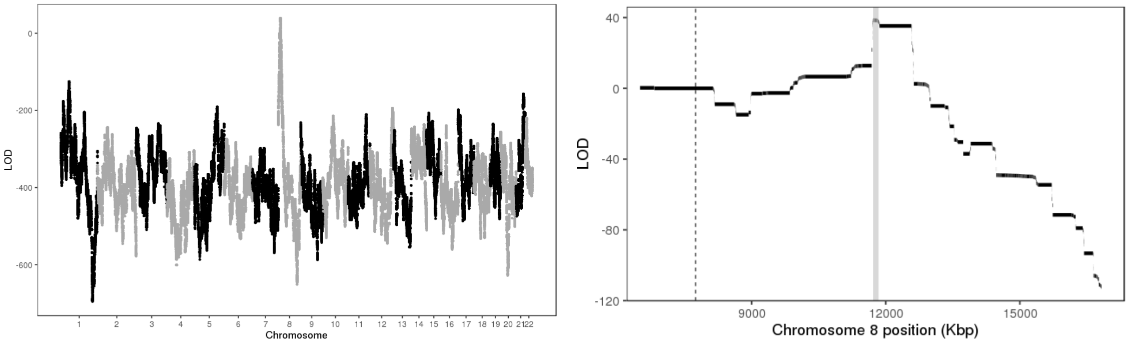


Figure 2: LOD scores of insertion locations across the genome (left) and around the 95% confidence interval (highlighted in gray, right).

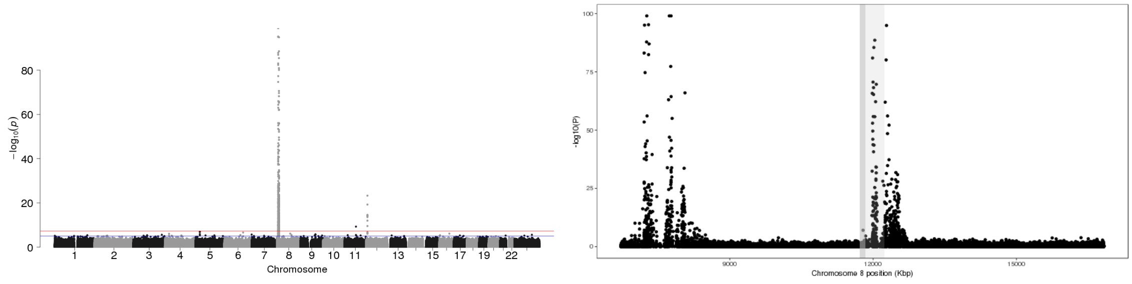


Figure 3: P-values of association between SNP genotypes and duplication copy number across the genome (left), and around the 95% confidence interval (highlighted in gray, right).

All coordinates refer to GRCh37.

### CNV_8_7773888_7791373

Insertion Location 95% Confidence Region:

8:12583654-12586093

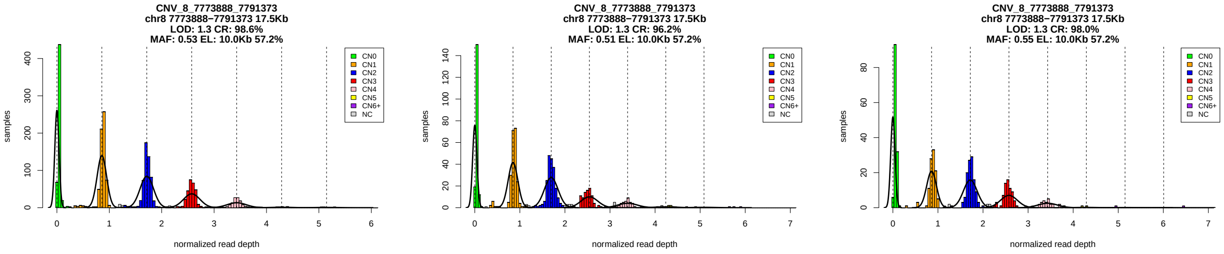


Figure 1: Copy number genotypes in the ASD, ID, and SCZ cohorts.

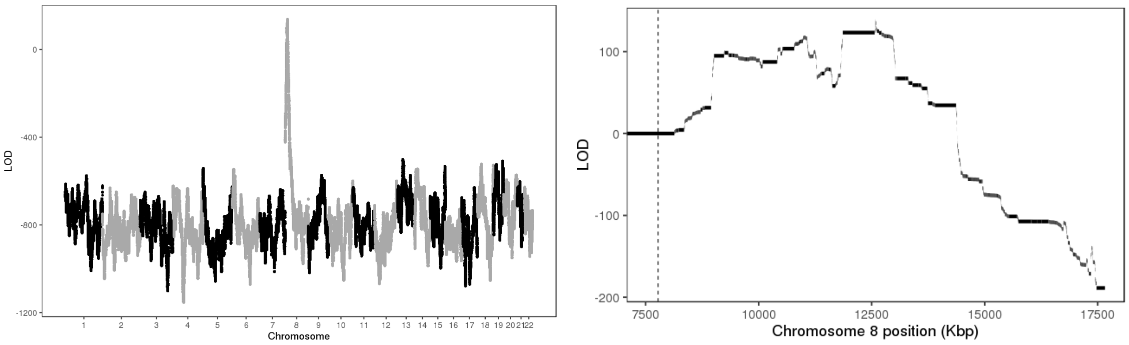


Figure 2: LOD scores of insertion locations across the genome (left) and around the 95% confidence interval (highlighted in gray, right).

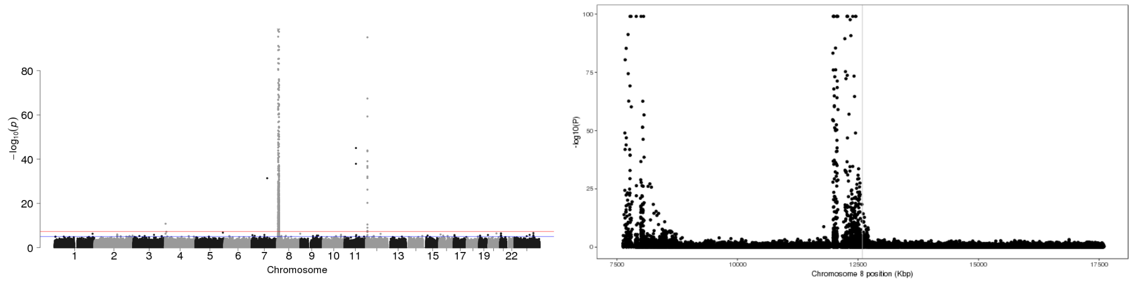


Figure 3: P-values of association between SNP genotypes and duplication copy number across the genome (left), and around the 95% confidence interval (highlighted in gray, right).

All coordinates refer to GRCh37.

### CNV_8_8077800_8087700

Insertion Location 95% Confidence Region:

8:11859851-12580568

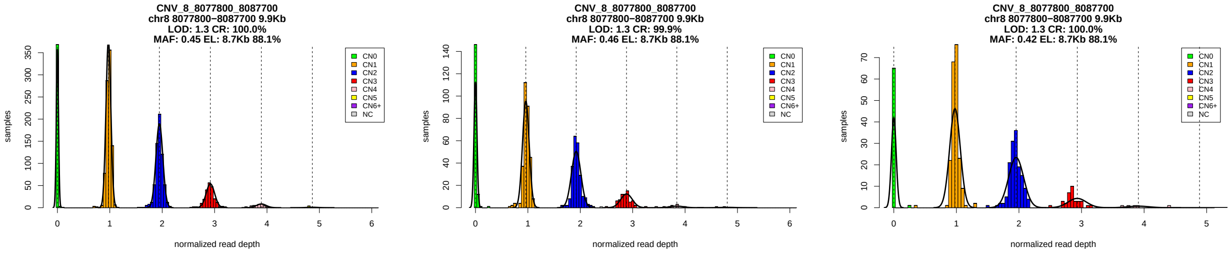


Figure 1: Copy number genotypes in the ASD, ID, and SCZ cohorts.

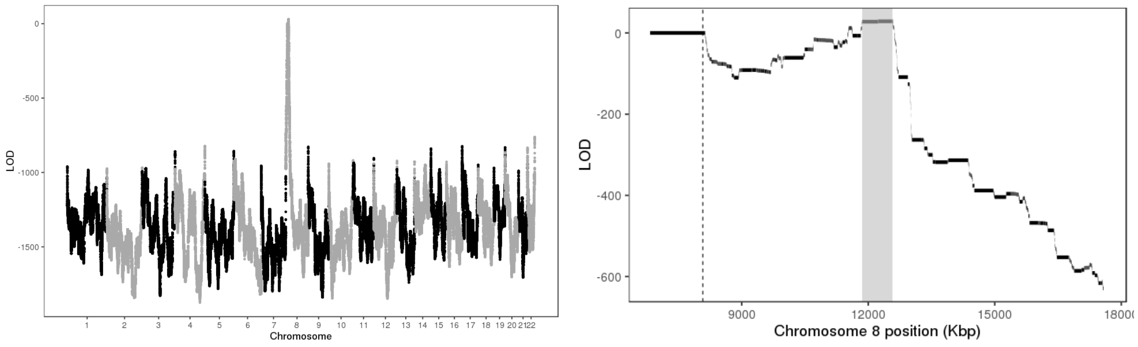


Figure 2: LOD scores of insertion locations across the genome (left) and around the 95% confidence interval (highlighted in gray, right).

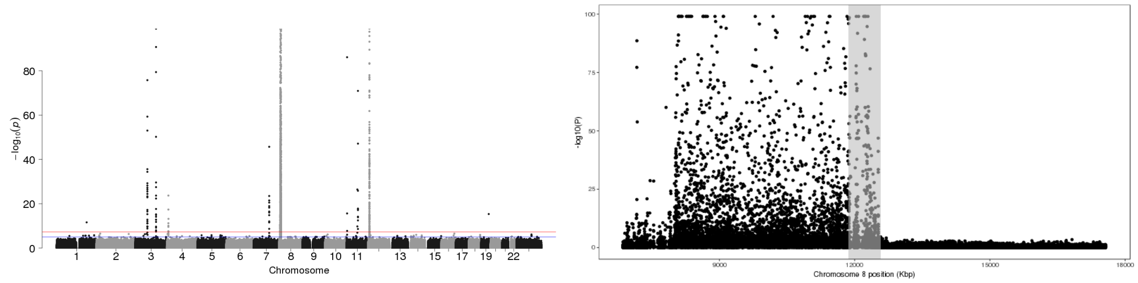


Figure 3: P-values of association between SNP genotypes and duplication copy number across the genome (left), and around the 95% confidence interval (highlighted in gray, right).

All coordinates refer to GRCh37.

### CNV_8_12235600_12248700

Insertion Location 95% Confidence Region:

8:12588849-12593045

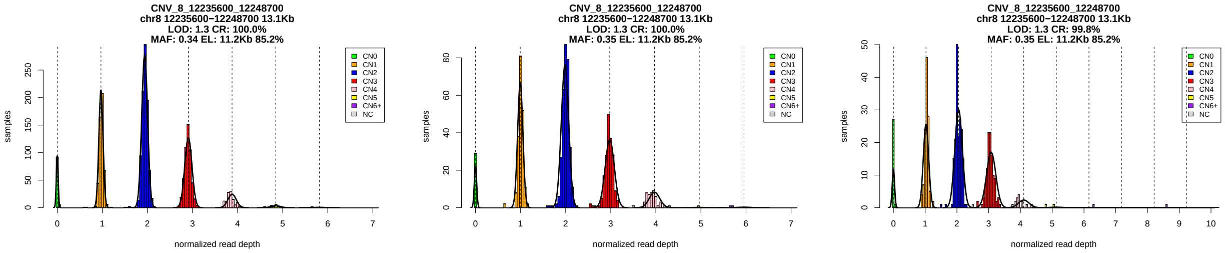


Figure 1: Copy number genotypes in the ASD, ID, and SCZ cohorts.

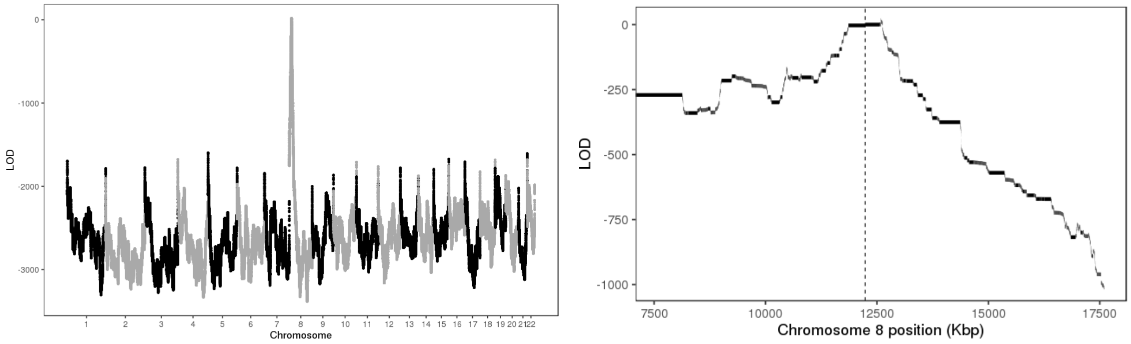


Figure 2: LOD scores of insertion locations across the genome (left) and around the 95% confidence interval (highlighted in gray, right).

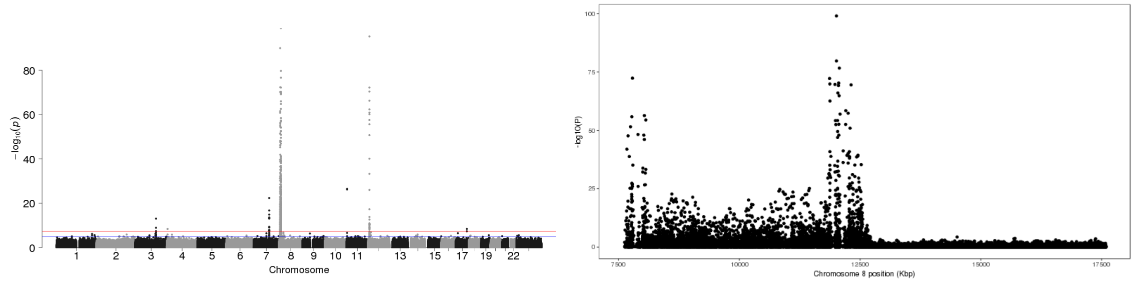


Figure 3: P-values of association between SNP genotypes and duplication copy number across the genome (left), and around the 95% confidence interval (highlighted in gray, right).

All coordinates refer to GRCh37.

### CNV_9_101476413_101477612

Insertion Location 95% Confidence Region:

22:32334686-32664241

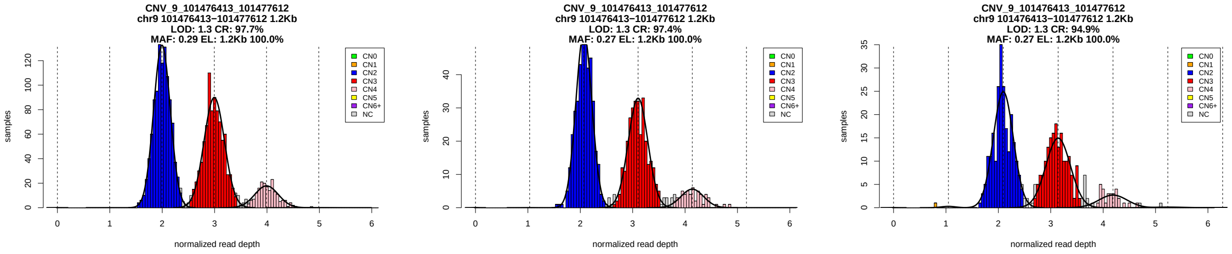


Figure 1: Copy number genotypes in the ASD, ID, and SCZ cohorts.

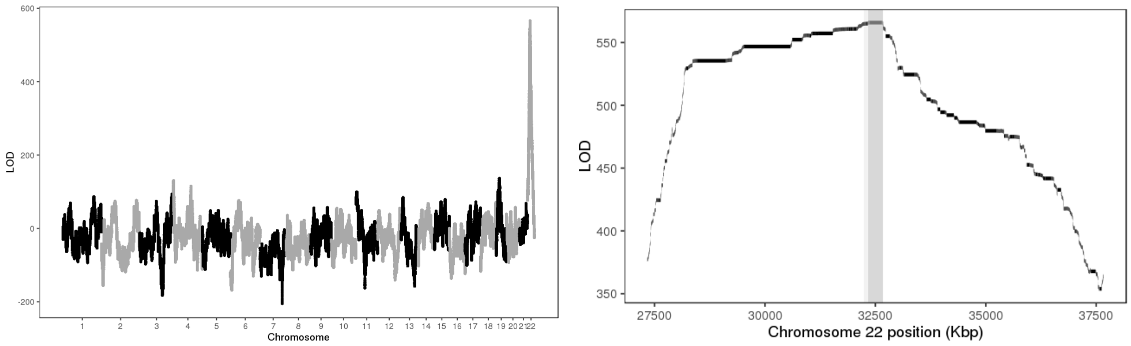


Figure 2: LOD scores of insertion locations across the genome (left) and around the 95% confidence interval (highlighted in gray, right).

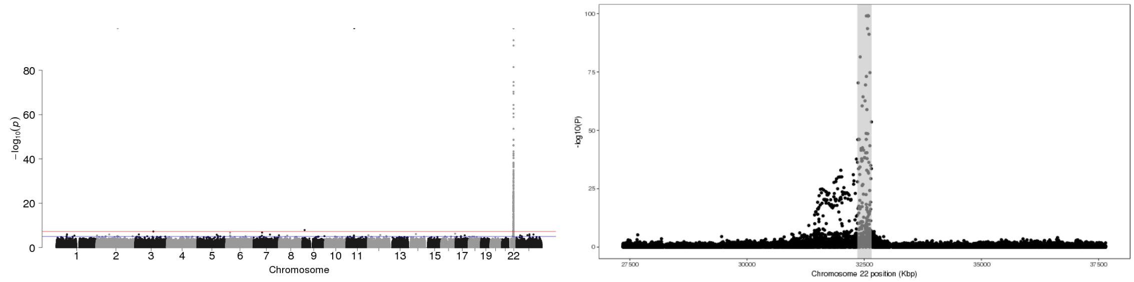


Figure 3: P-values of association between SNP genotypes and duplication copy number across the genome (left), and around the 95% confidence interval (highlighted in gray, right).

All coordinates refer to GRCh37.

### CNV_9_132186781_132206286

Insertion Location 95% Confidence Region:

22:23853706-23993051

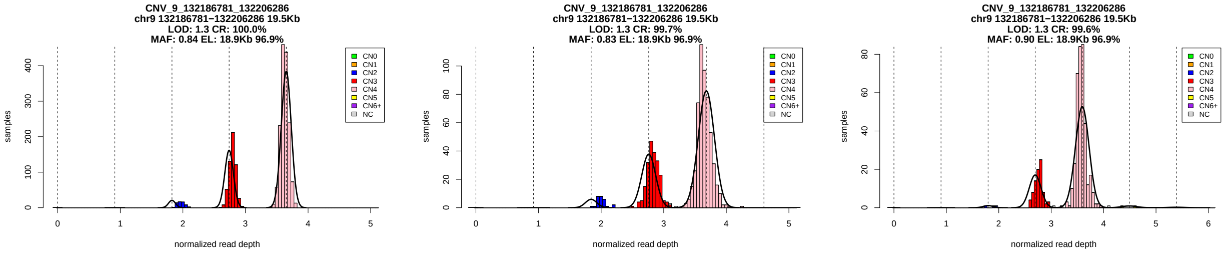


Figure 1: Copy number genotypes in the ASD, ID, and SCZ cohorts.

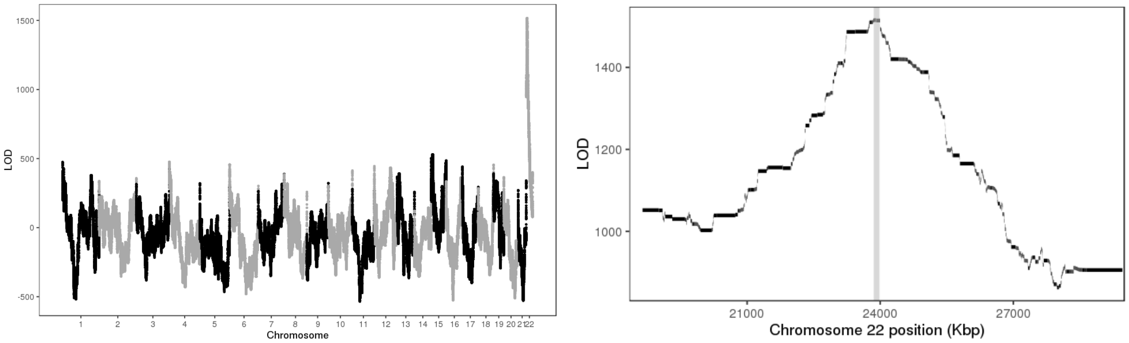


Figure 2: LOD scores of insertion locations across the genome (left) and around the 95% confidence interval (highlighted in gray, right).

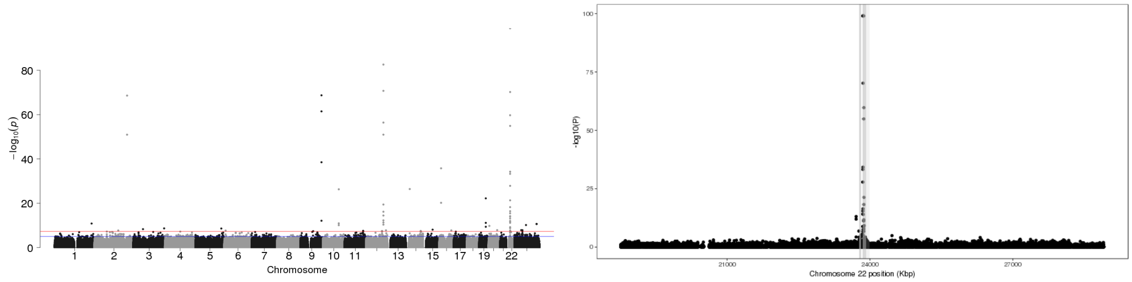


Figure 3: P-values of association between SNP genotypes and duplication copy number across the genome (left), and around the 95% confidence interval (highlighted in gray, right).

All coordinates refer to GRCh37.

### CNV_10_26913676_26915975

Insertion Location 95% Confidence Region:

10:27210569-27865573

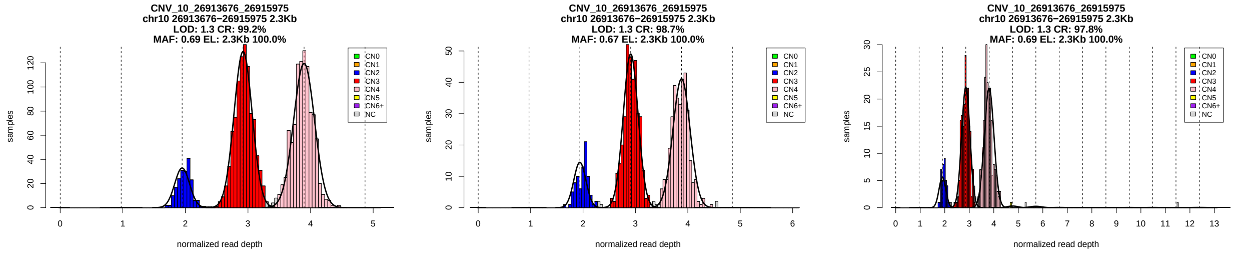


Figure 1: Copy number genotypes in the ASD, ID, and SCZ cohorts.

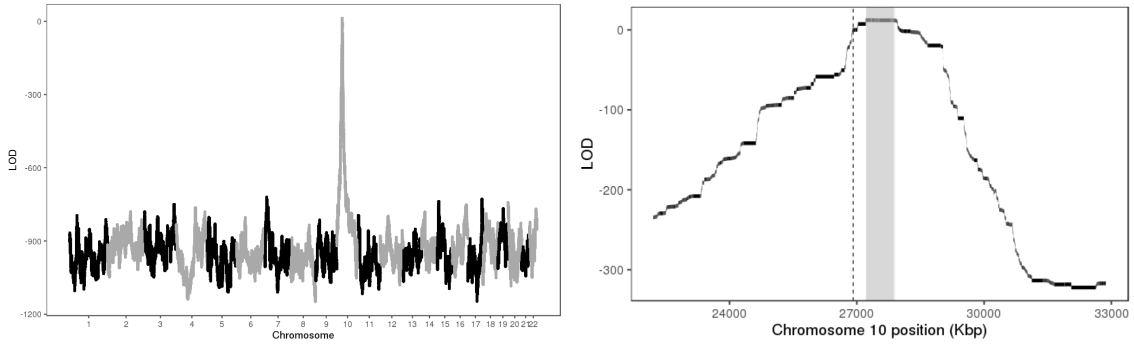


Figure 2: LOD scores of insertion locations across the genome (left) and around the 95% confidence interval (highlighted in gray, right).

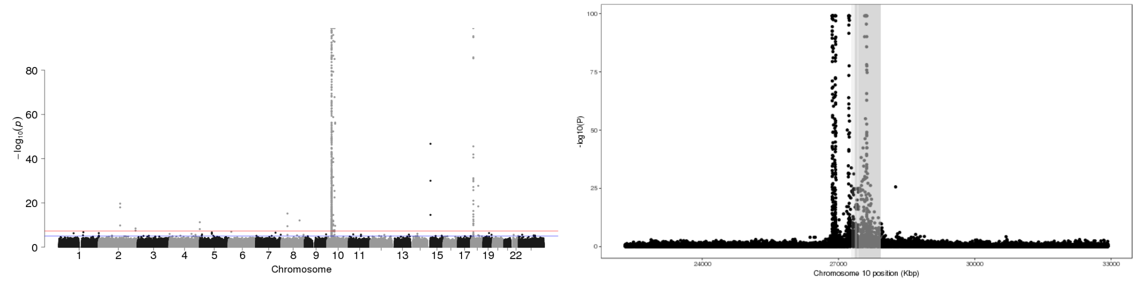


Figure 3: P-values of association between SNP genotypes and duplication copy number across the genome (left), and around the 95% confidence interval (highlighted in gray, right).

All coordinates refer to GRCh37.

### CNV_10_46219000_46222900

Insertion Location 95% Confidence Region:

10:50862335-52119297

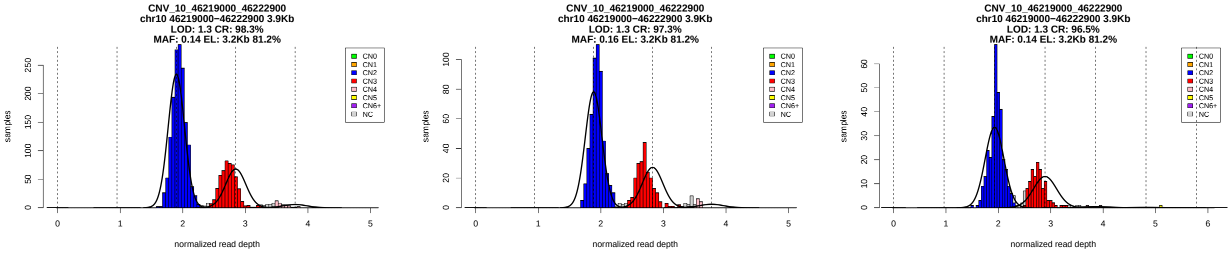


Figure 1: Copy number genotypes in the ASD, ID, and SCZ cohorts.

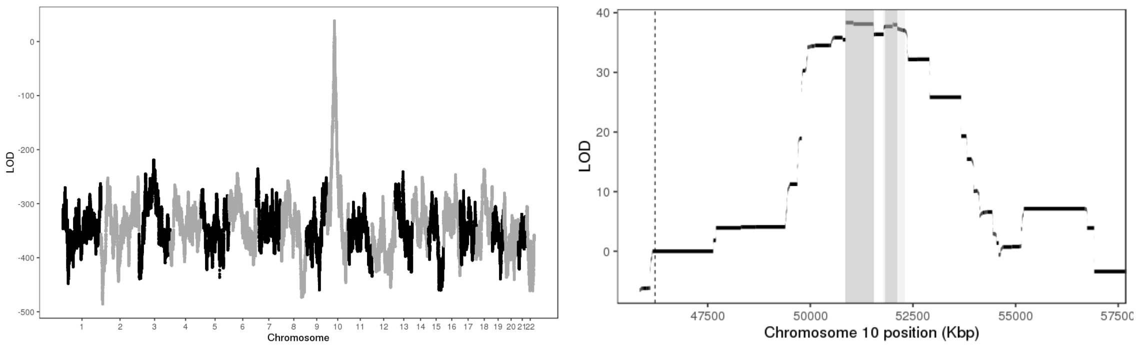


Figure 2: LOD scores of insertion locations across the genome (left) and around the 95% confidence interval (highlighted in gray, right).

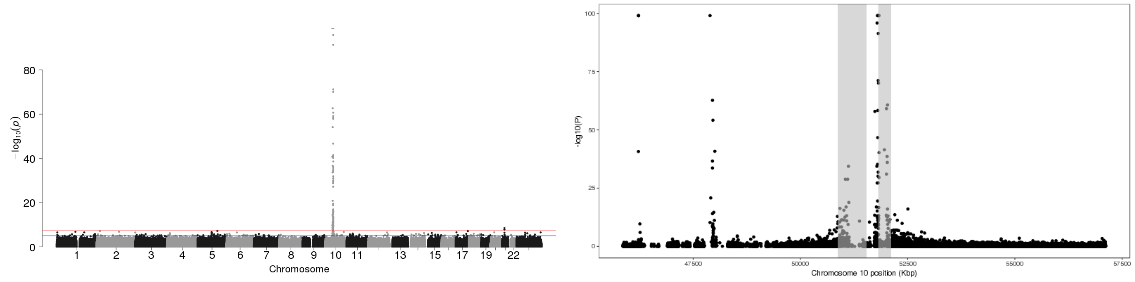


Figure 3: P-values of association between SNP genotypes and duplication copy number across the genome (left), and around the 95% confidence interval (highlighted in gray, right).

All coordinates refer to GRCh37.

# GS_SD_M2_10_46712812_46947015_10_47190000_47429169

Insertion Location 95% Confidence Region:

10:47703870-48303900

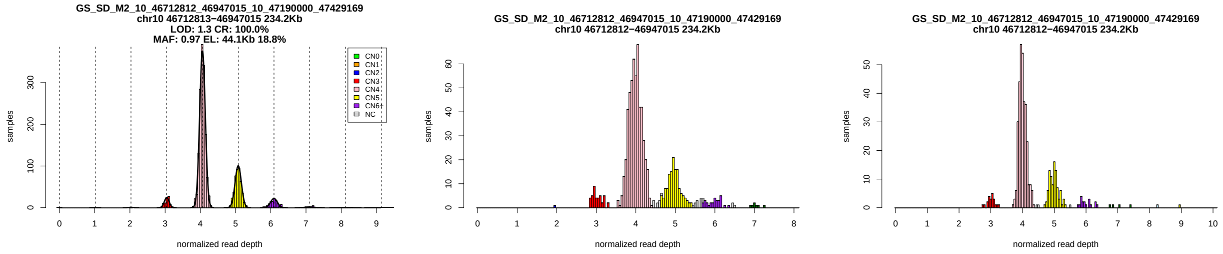


Figure 1: Copy number genotypes in the ASD, ID, and SCZ cohorts.

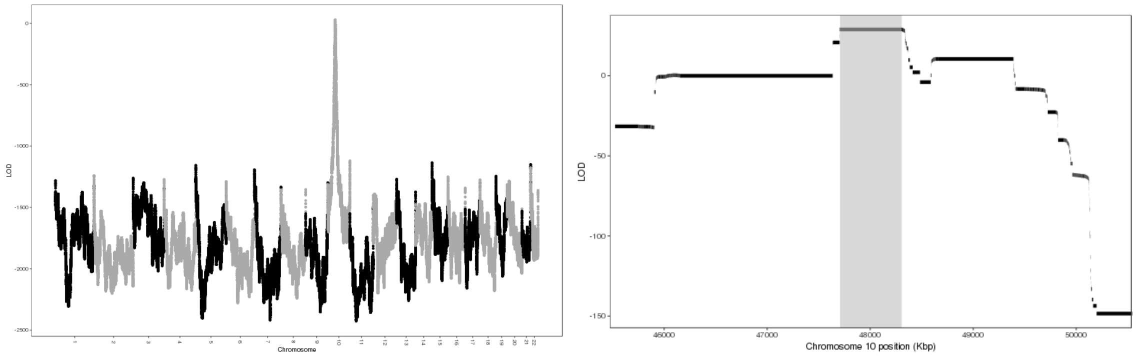


Figure 2: LOD scores of insertion locations across the genome (left) and around the 95% confidence interval (highlighted in gray, right).

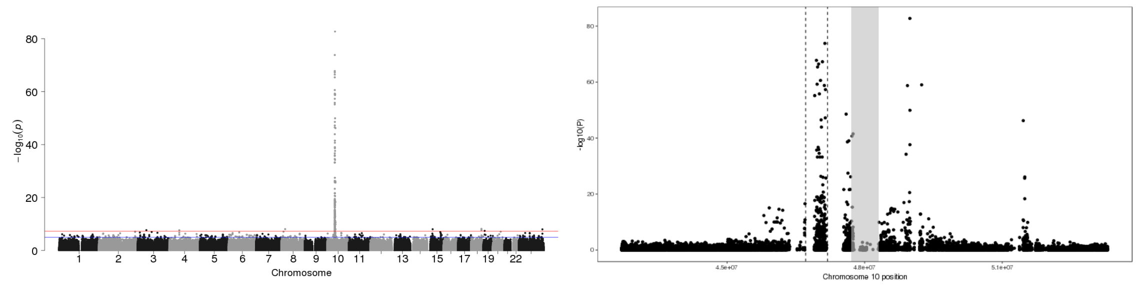


Figure 3: P-values of association between SNP genotypes and duplication copy number across the genome (left), and around the 95% confidence interval (highlighted in gray, right).

All coordinates refer to GRCh37.

### CNV_10_65432549_65434248

Insertion Location 95% Confidence Region:

1:162451182-163813482

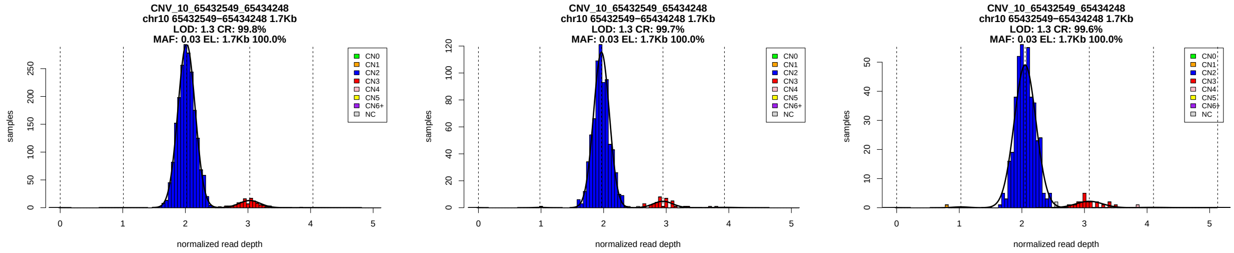

Figure 1: Copy number genotypes in the ASD, ID, and SCZ cohorts.

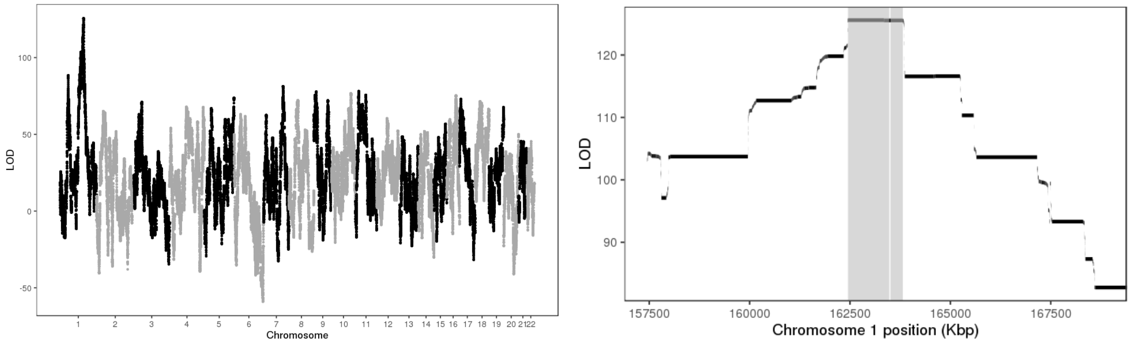

Figure 2: LOD scores of insertion locations across the genome (left) and around the 95% confidence interval (highlighted in gray, right).

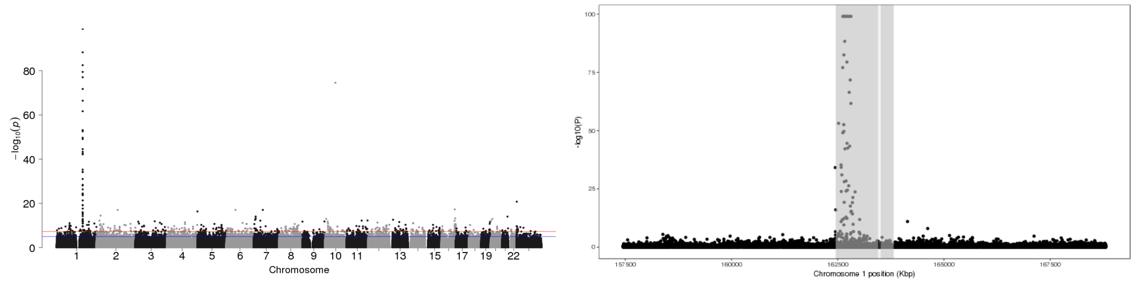

Figure 3: P-values of association between SNP genotypes and duplication copy number across the genome (left), and around the 95% confidence interval (highlighted in gray, right).

All coordinates refer to GRCh37.

### CNV_12_31266000_31407500

Insertion Location 95% Confidence Region:

12:8970652-11630778

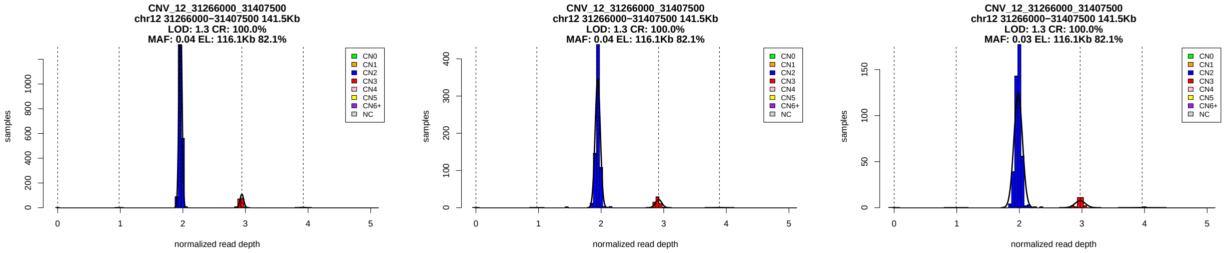


Figure 1: Copy number genotypes in the ASD, ID, and SCZ cohorts.

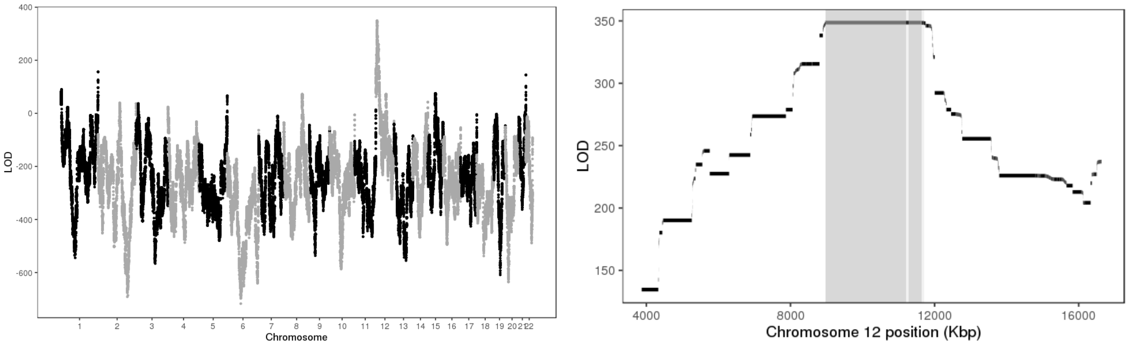


Figure 2: LOD scores of insertion locations across the genome (left) and around the 95% confidence interval (highlighted in gray, right).

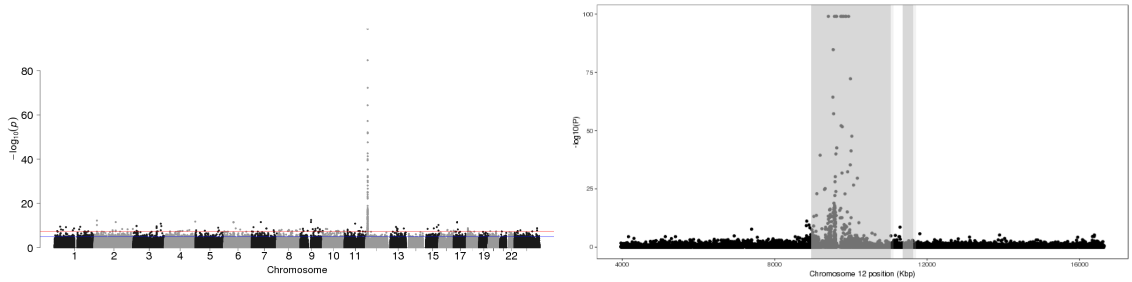


Figure 3: P-values of association between SNP genotypes and duplication copy number across the genome (left), and around the 95% confidence interval (highlighted in gray, right).

All coordinates refer to GRCh37.

### CNV_12_55782038_55784208

Insertion Location 95% Confidence Region:

2:177374056-180168421

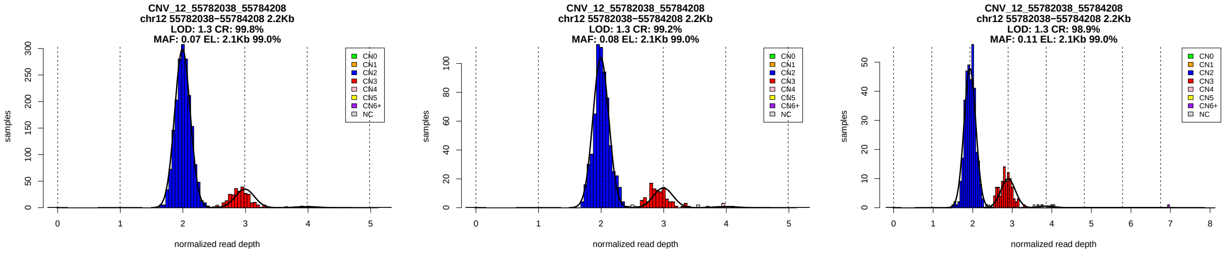


Figure 1: Copy number genotypes in the ASD, ID, and SCZ cohorts.

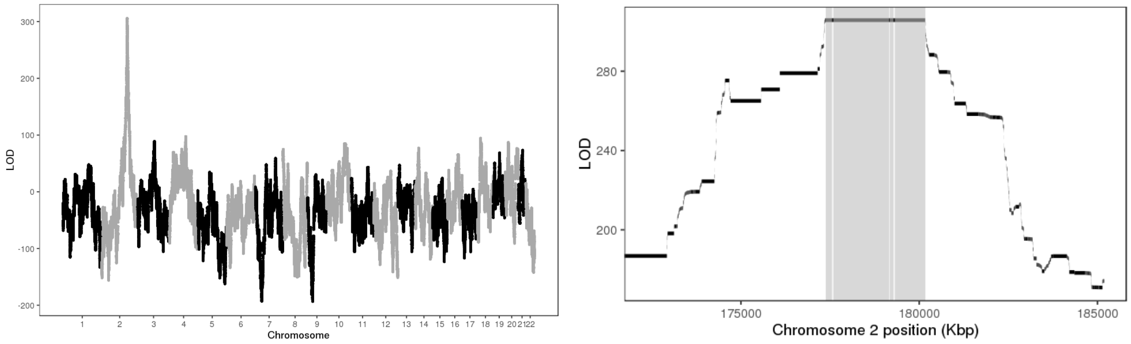


Figure 2: LOD scores of insertion locations across the genome (left) and around the 95% confidence interval (highlighted in gray, right).

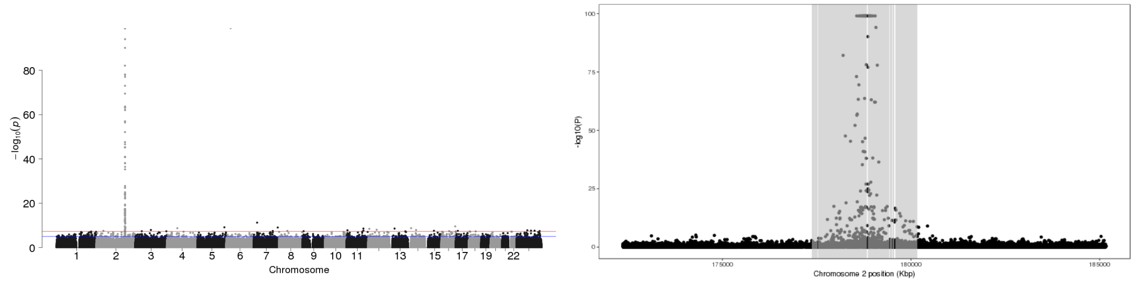


Figure 3: P-values of association between SNP genotypes and duplication copy number across the genome (left), and around the 95% confidence interval (highlighted in gray, right).

All coordinates refer to GRCh37.

### CNV_12_124495925_124499824

Insertion Location 95% Confidence Region:

2:3913045-3994446

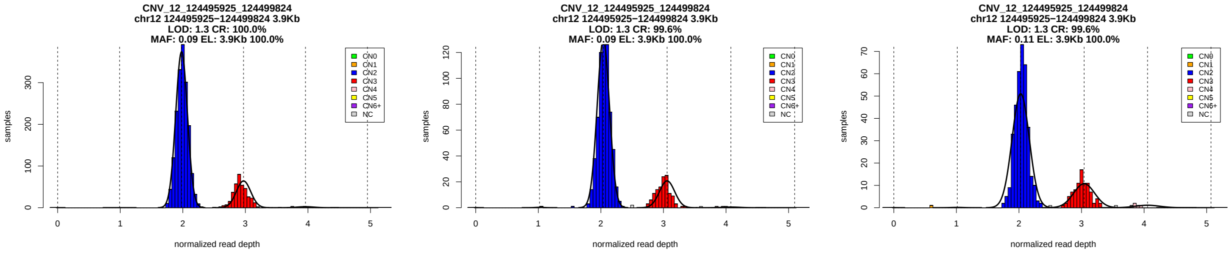


Figure 1: Copy number genotypes in the ASD, ID, and SCZ cohorts.

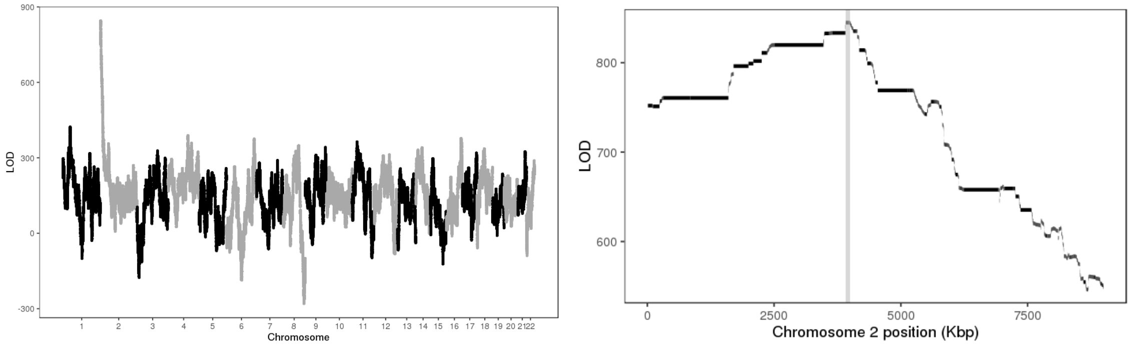


Figure 2: LOD scores of insertion locations across the genome (left) and around the 95% confidence interval (highlighted in gray, right).

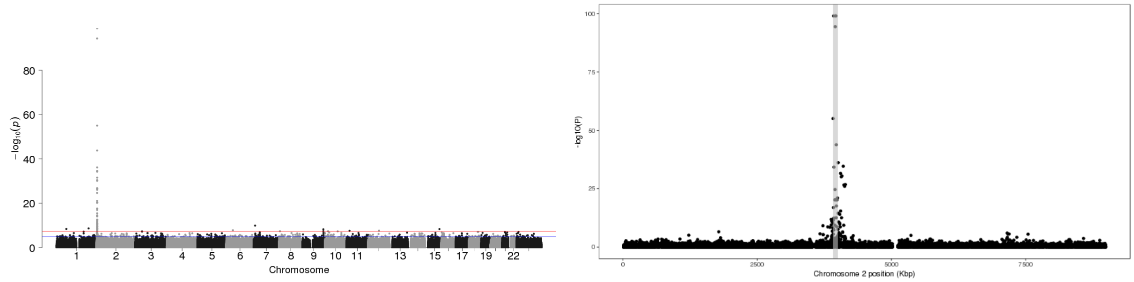


Figure 3: P-values of association between SNP genotypes and duplication copy number across the genome (left), and around the 95% confidence interval (highlighted in gray, right).

All coordinates refer to GRCh37.

### CNV_12_131792472_131796941

Insertion Location 95% Confidence Region:

12:132174274-132178598

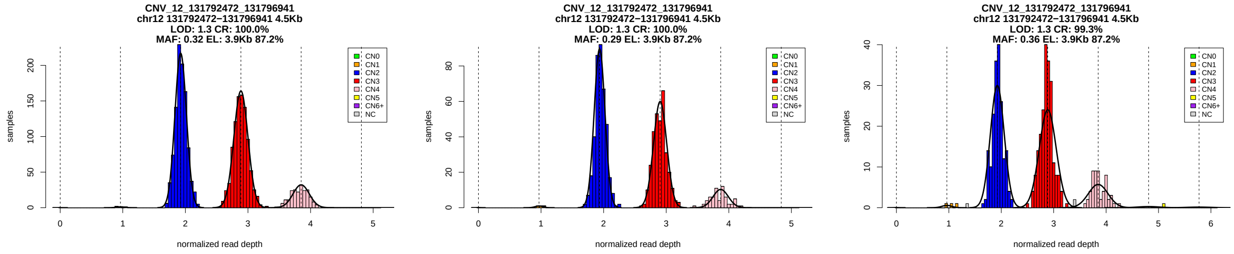

Figure 1: Copy number genotypes in the ASD, ID, and SCZ cohorts.

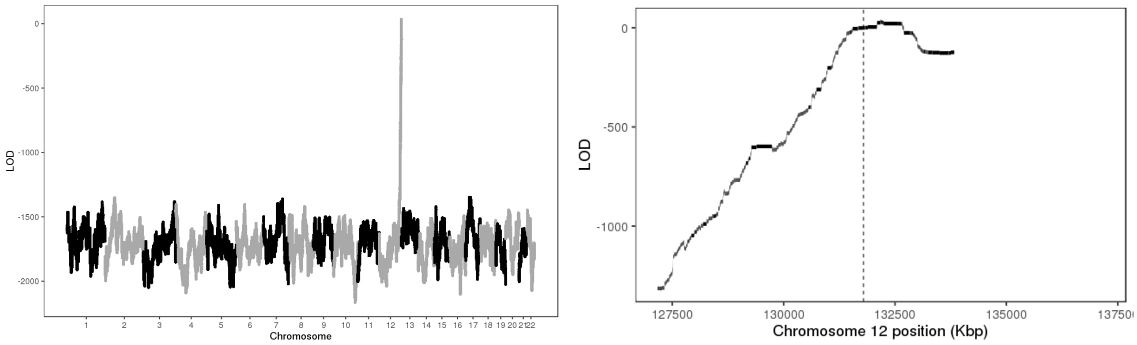

Figure 2: LOD scores of insertion locations across the genome (left) and around the 95% confidence interval (highlighted in gray, right).

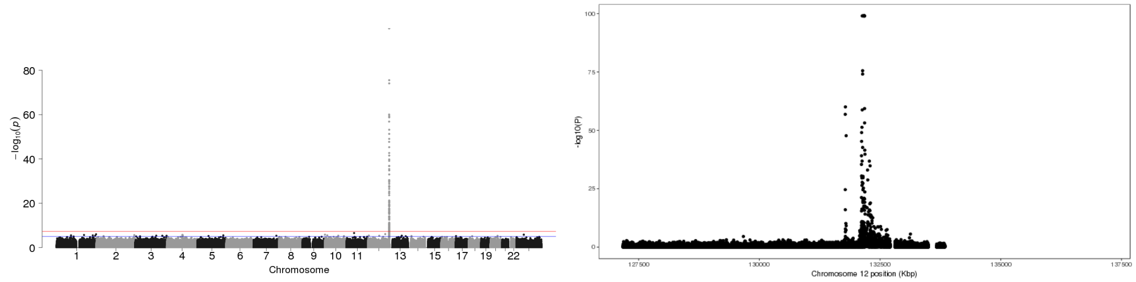

Figure 3: P-values of association between SNP genotypes and duplication copy number across the genome (left), and around the 95% confidence interval (highlighted in gray, right).

All coordinates refer to GRCh37.

### CNV_12_133085650_133088265

Insertion Location 95% Confidence Region:

6:27518819-30365740

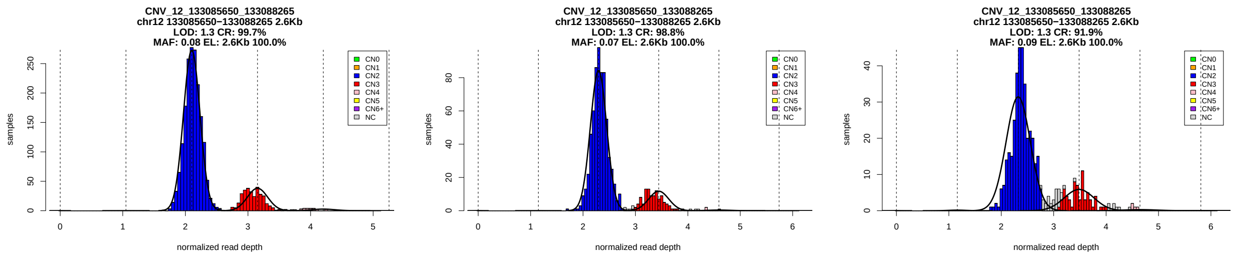


Figure 1: Copy number genotypes in the ASD, ID, and SCZ cohorts.

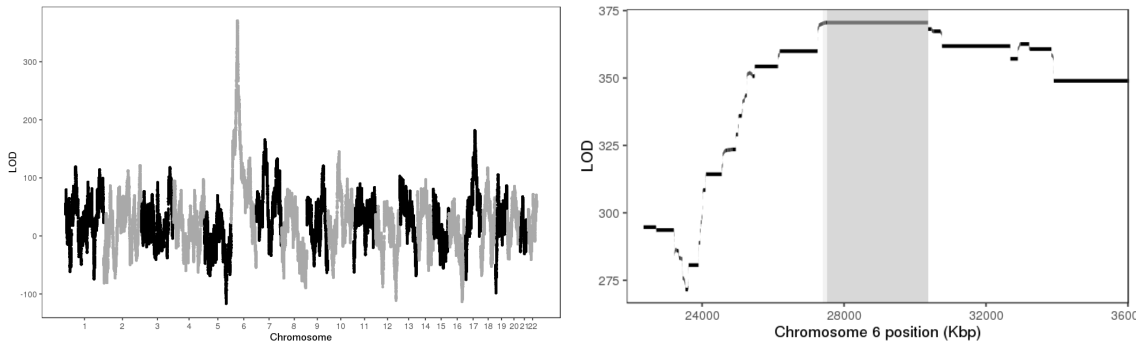


Figure 2: LOD scores of insertion locations across the genome (left) and around the 95% confidence interval (highlighted in gray, right).

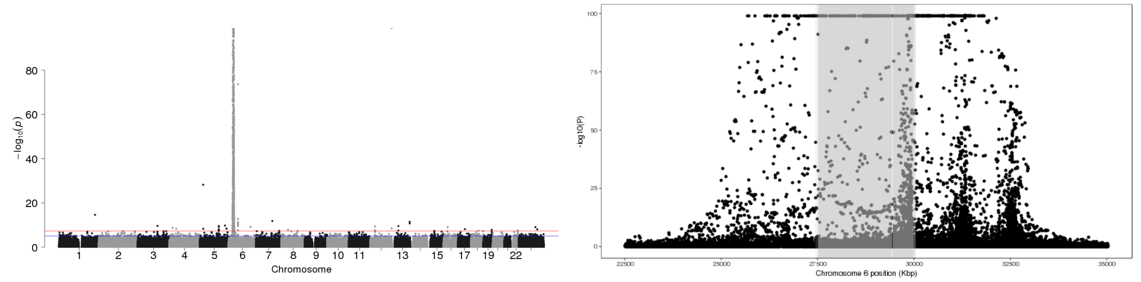


Figure 3: P-values of association between SNP genotypes and duplication copy number across the genome (left), and around the 95% confidence interval (highlighted in gray, right).

All coordinates refer to GRCh37.

### CNV_13_21727763_21729296

Insertion Location 95% Confidence Region:

11:108745546-109045098

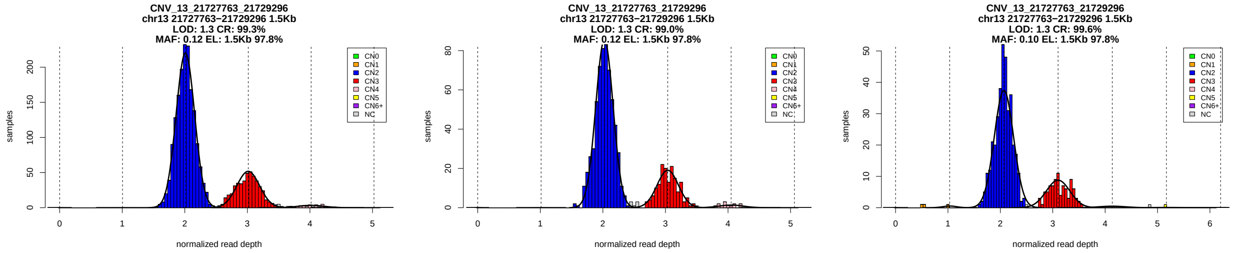


Figure 1: Copy number genotypes in the ASD, ID, and SCZ cohorts.

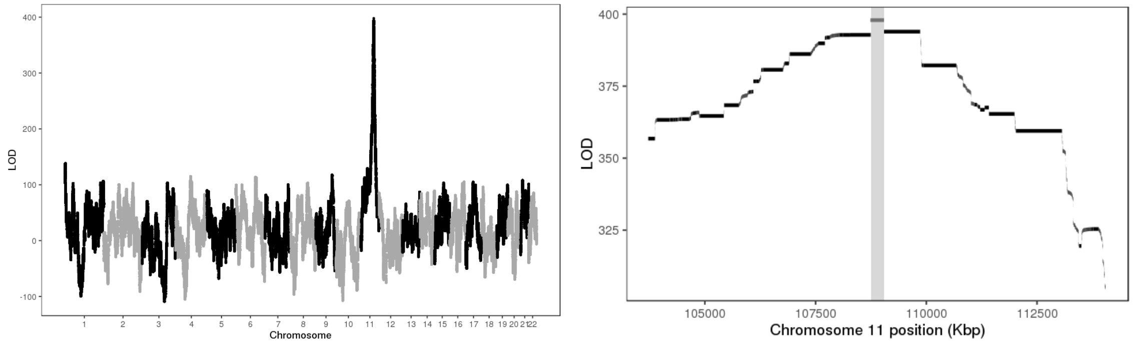


Figure 2: LOD scores of insertion locations across the genome (left) and around the 95% confidence interval (highlighted in gray, right).

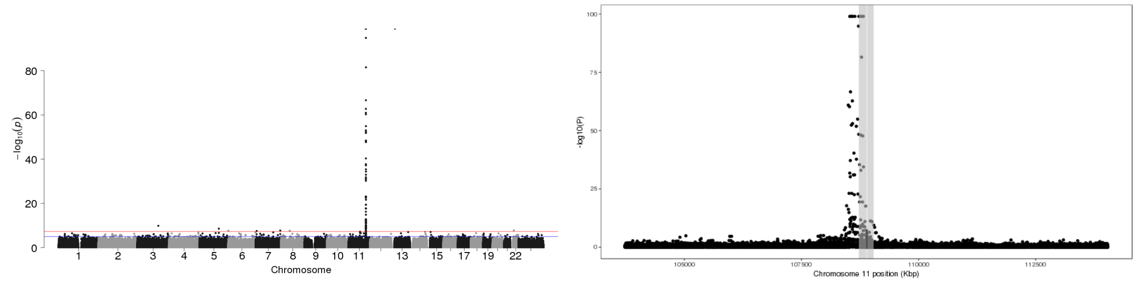


Figure 3: P-values of association between SNP genotypes and duplication copy number across the genome (left), and around the 95% confidence interval (highlighted in gray, right).

All coordinates refer to GRCh37.

### CNV_13_24899872_24901951

Insertion Location 95% Confidence Region:

13:23314312-23545996

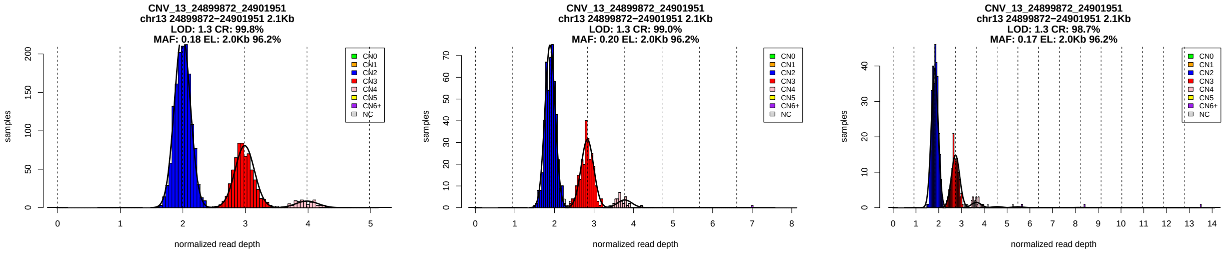


Figure 1: Copy number genotypes in the ASD, ID, and SCZ cohorts.

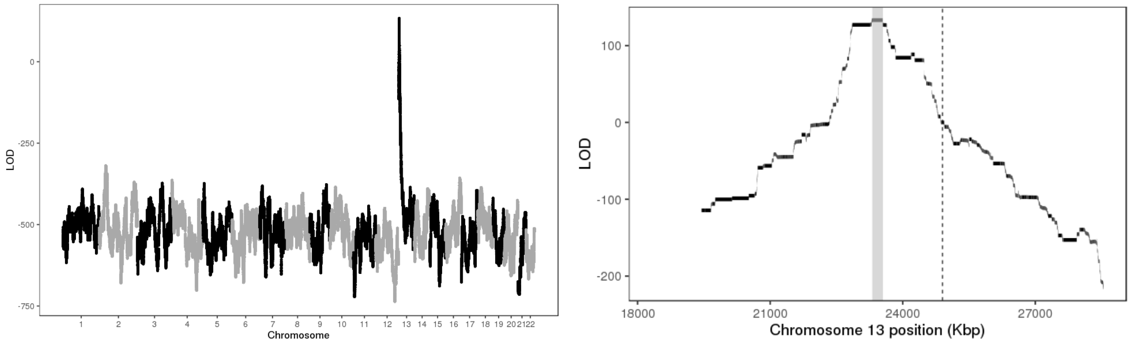


Figure 2: LOD scores of insertion locations across the genome (left) and around the 95% confidence interval (highlighted in gray, right).

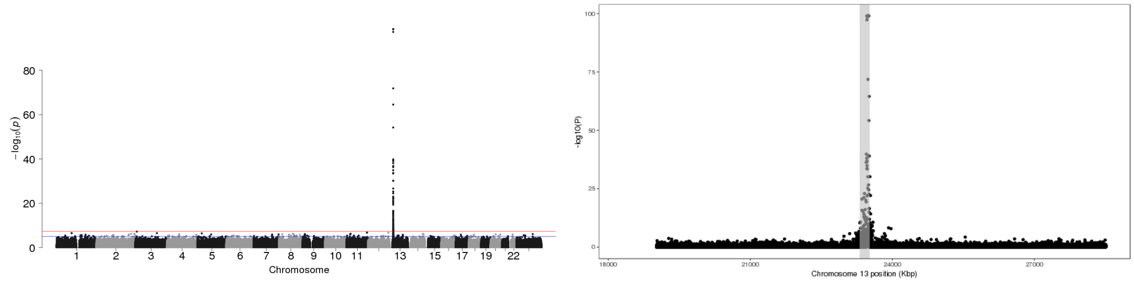


Figure 3: P-values of association between SNP genotypes and duplication copy number across the genome (left), and around the 95% confidence interval (highlighted in gray, right).

All coordinates refer to GRCh37.

### CNV_13_61461095_61462117

Insertion Location 95% Confidence Region:

5:19609838-23155734

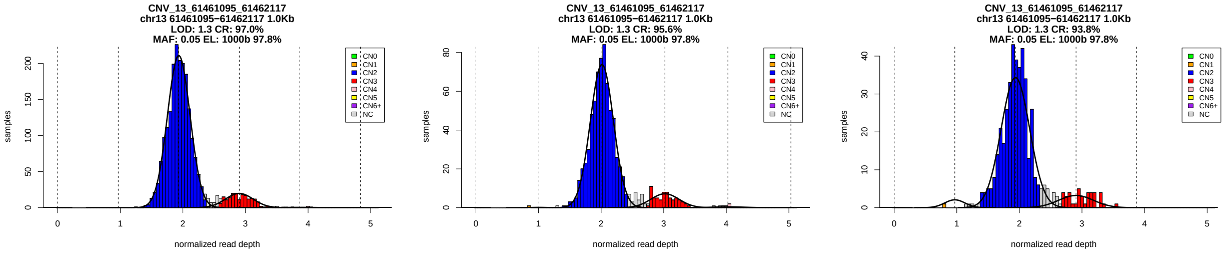


Figure 1: Copy number genotypes in the ASD, ID, and SCZ cohorts.

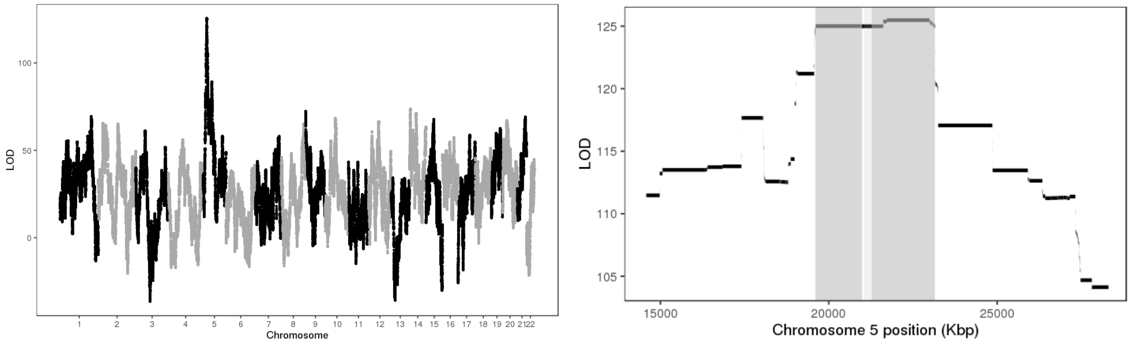


Figure 2: LOD scores of insertion locations across the genome (left) and around the 95% confidence interval (highlighted in gray, right).

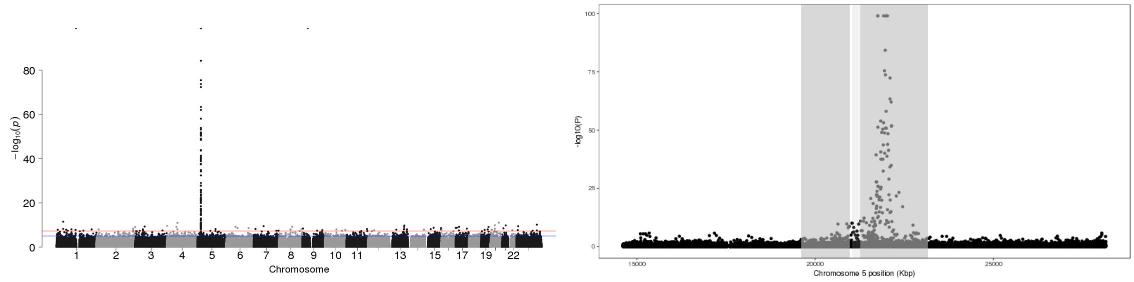


Figure 3: P-values of association between SNP genotypes and duplication copy number across the genome (left), and around the 95% confidence interval (highlighted in gray, right).

All coordinates refer to GRCh37.

### CNV_15_28700474_28710000

Insertion Location 95% Confidence Region:

15:20173314-23682893

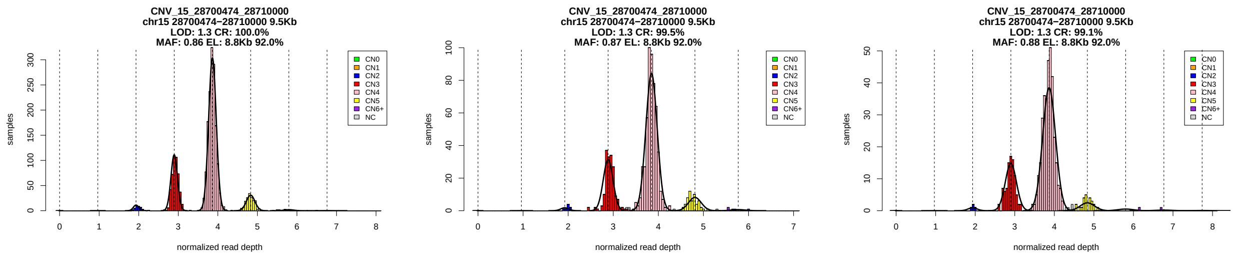


Figure 1: Copy number genotypes in the ASD, ID, and SCZ cohorts.

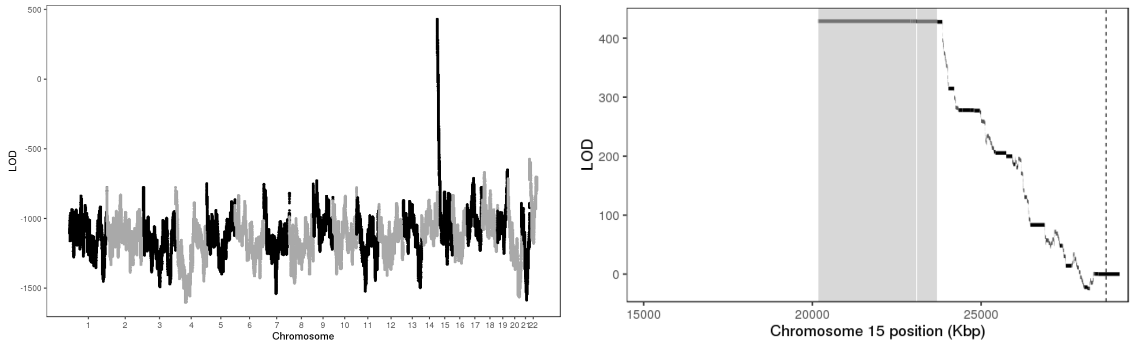


Figure 2: LOD scores of insertion locations across the genome (left) and around the 95% confidence interval (highlighted in gray, right).

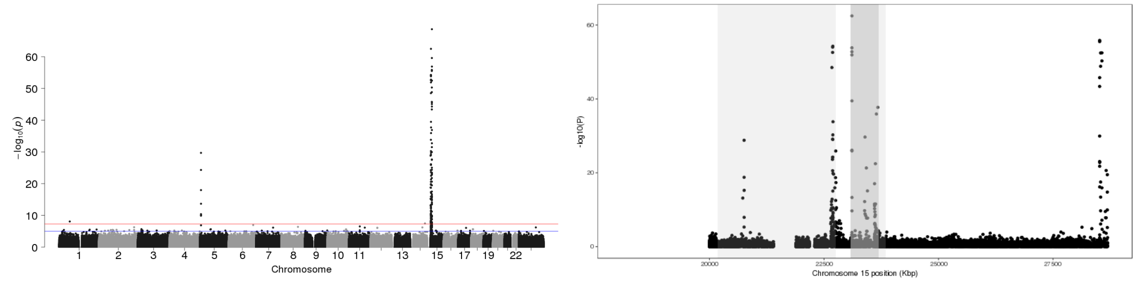


Figure 3: P-values of association between SNP genotypes and duplication copy number across the genome (left), and around the 95% confidence interval (highlighted in gray, right).

All coordinates refer to GRCh37.

### CNV_15_28934858_28936946

Insertion Location 95% Confidence Region:

15:20173314-23874362

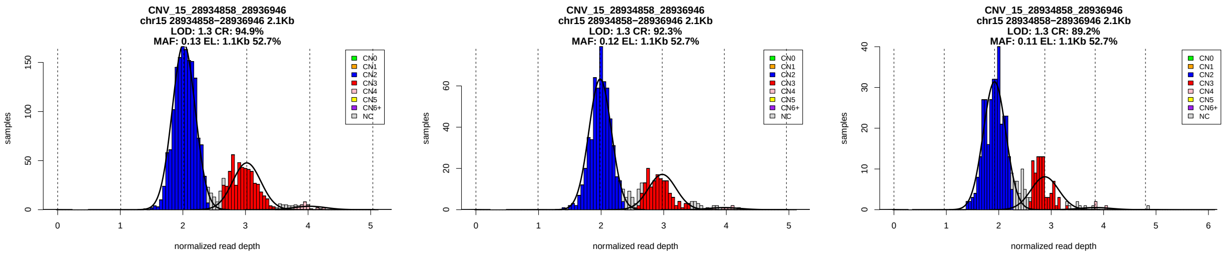


Figure 1: Copy number genotypes in the ASD, ID, and SCZ cohorts.

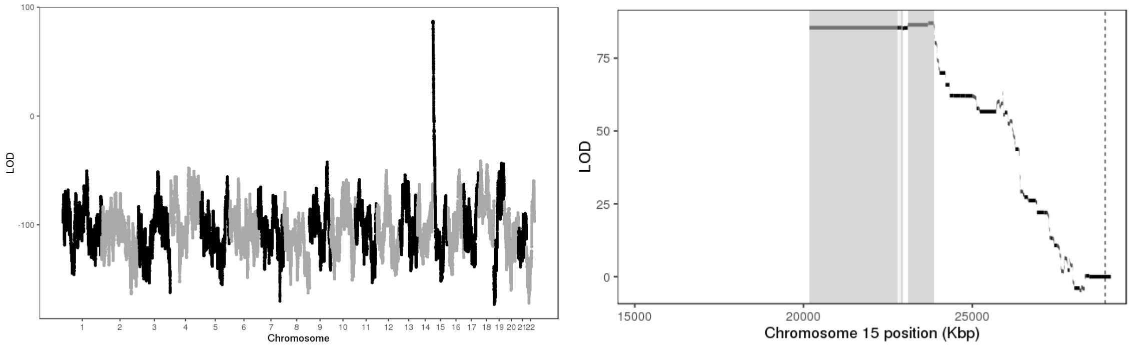


Figure 2: LOD scores of insertion locations across the genome (left) and around the 95% confidence interval (highlighted in gray, right).

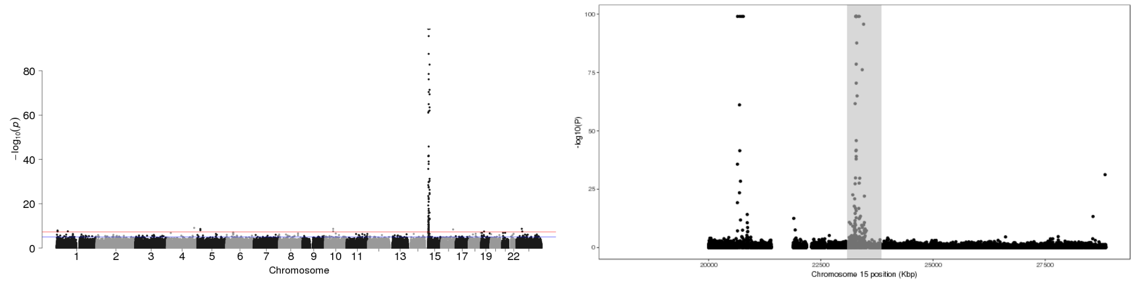


Figure 3: P-values of association between SNP genotypes and duplication copy number across the genome (left), and around the 95% confidence interval (highlighted in gray, right).

All coordinates refer to GRCh37.

### CNV_15_28937648_28943000

Insertion Location 95% Confidence Region:

15:20173314-23619572

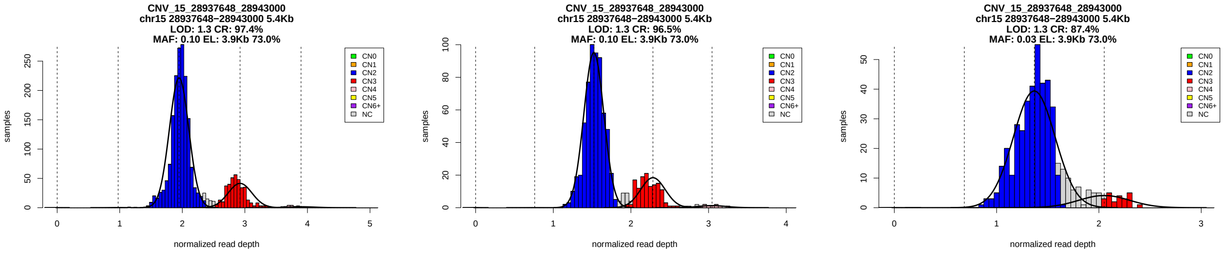


Figure 1: Copy number genotypes in the ASD, ID, and SCZ cohorts.

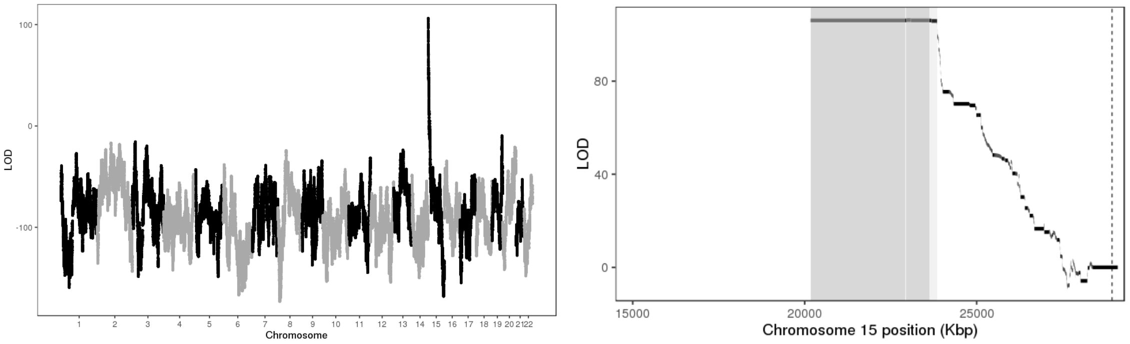


Figure 2: LOD scores of insertion locations across the genome (left) and around the 95% confidence interval (highlighted in gray, right).

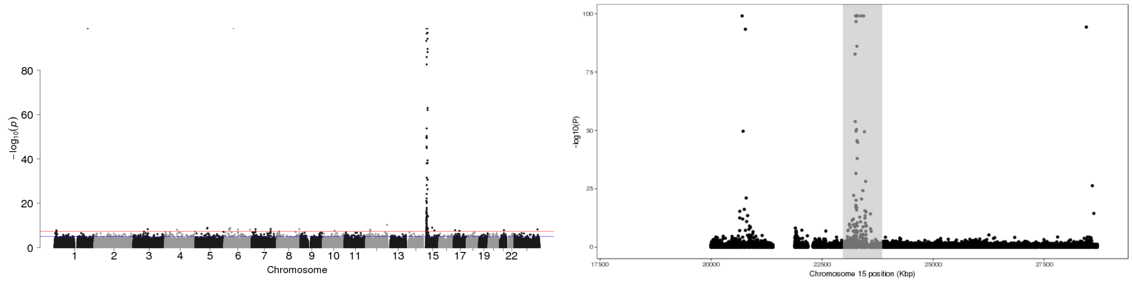


Figure 3: P-values of association between SNP genotypes and duplication copy number across the genome (left), and around the 95% confidence interval (highlighted in gray, right).

All coordinates refer to GRCh37.

### CNV_15_41870174_41871276

Insertion Location 95% Confidence Region:

13:43848721-44191220

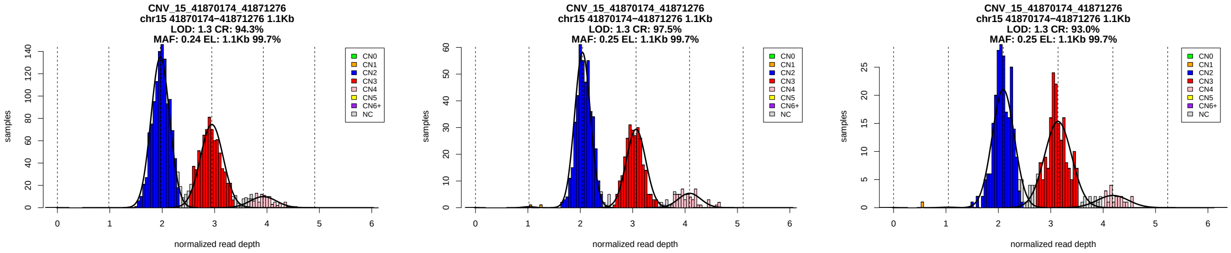


Figure 1: Copy number genotypes in the ASD, ID, and SCZ cohorts.

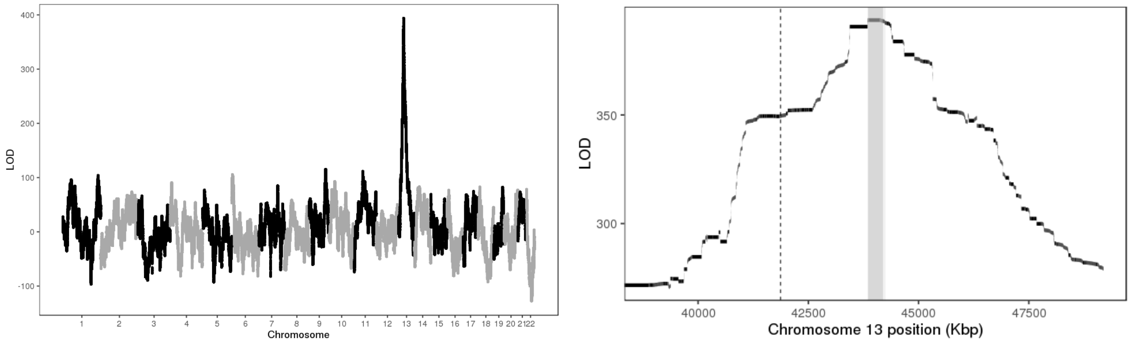


Figure 2: LOD scores of insertion locations across the genome (left) and around the 95% confidence interval (highlighted in gray, right).

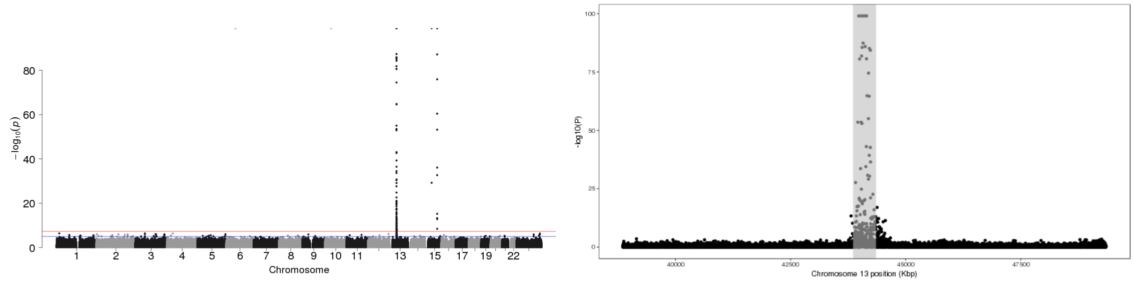


Figure 3: P-values of association between SNP genotypes and duplication copy number across the genome (left), and around the 95% confidence interval (highlighted in gray, right).

All coordinates refer to GRCh37.

### CNV_15_94886400_94888292

Insertion Location 95% Confidence Region:

12:72981831-73667370

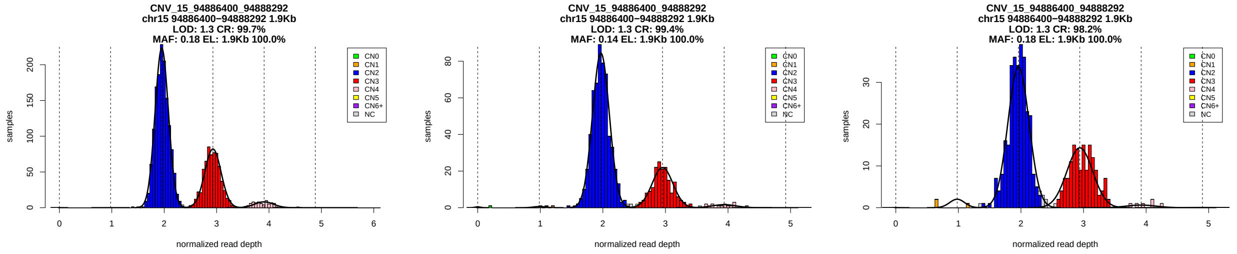


Figure 1: Copy number genotypes in the ASD, ID, and SCZ cohorts.

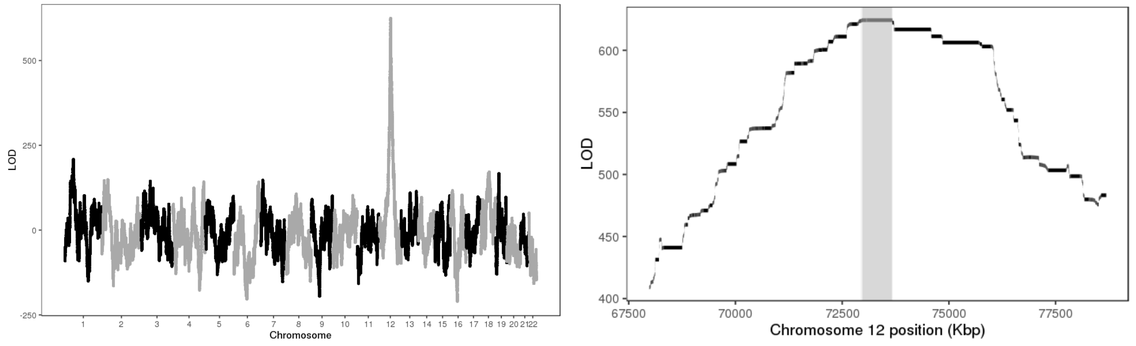


Figure 2: LOD scores of insertion locations across the genome (left) and around the 95% confidence interval (highlighted in gray, right).

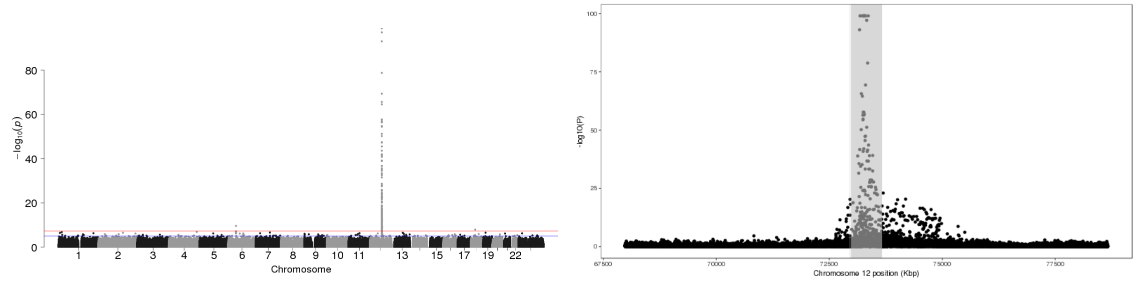


Figure 3: P-values of association between SNP genotypes and duplication copy number across the genome (left), and around the 95% confidence interval (highlighted in gray, right).

All coordinates refer to GRCh37.

### CNV_16_16559993_16563400

Insertion Location 95% Confidence Region:

16:18126929-18821191

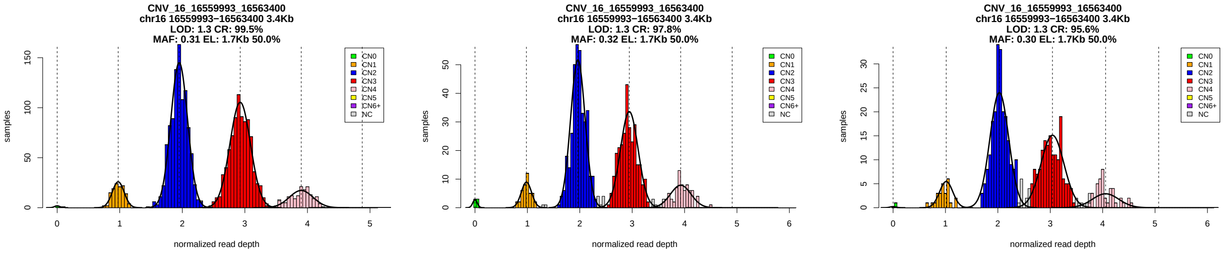


Figure 1: Copy number genotypes in the ASD, ID, and SCZ cohorts.

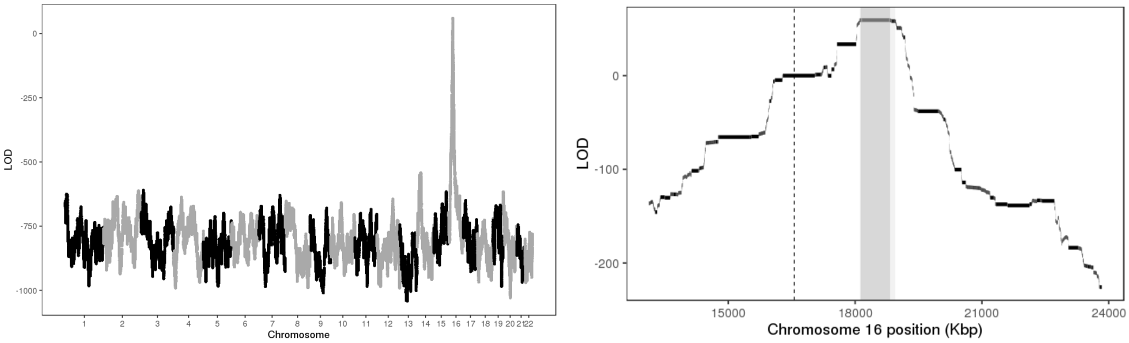


Figure 2: LOD scores of insertion locations across the genome (left) and around the 95% confidence interval (highlighted in gray, right).

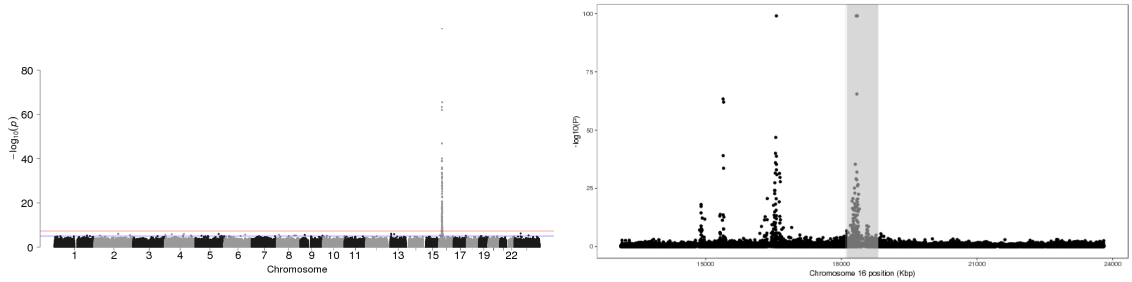


Figure 3: P-values of association between SNP genotypes and duplication copy number across the genome (left), and around the 95% confidence interval (highlighted in gray, right).

All coordinates refer to GRCh37.

### CNV_16_16629916_16637000

Insertion Location 95% Confidence Region:

16:18166840-18969357

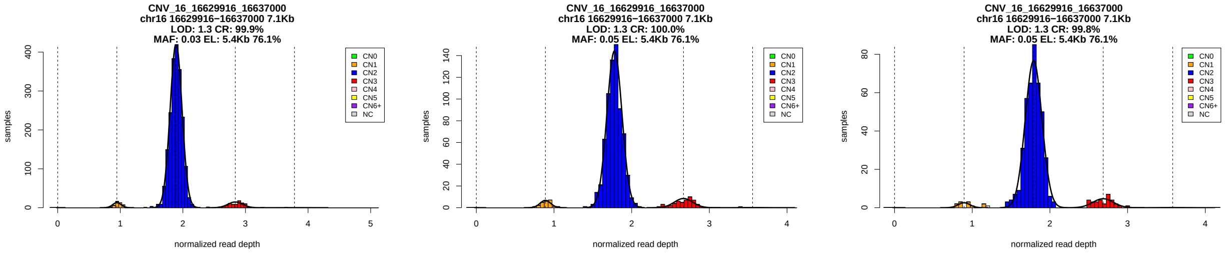


Figure 1: Copy number genotypes in the ASD, ID, and SCZ cohorts.

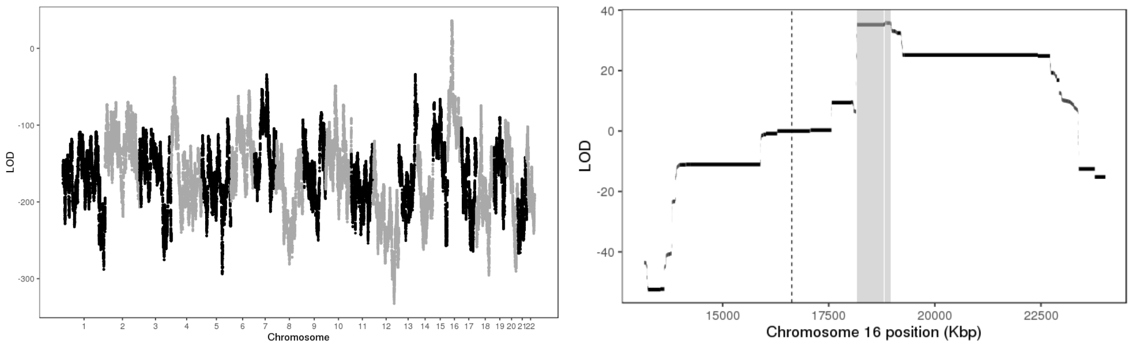


Figure 2: LOD scores of insertion locations across the genome (left) and around the 95% confidence interval (highlighted in gray, right).

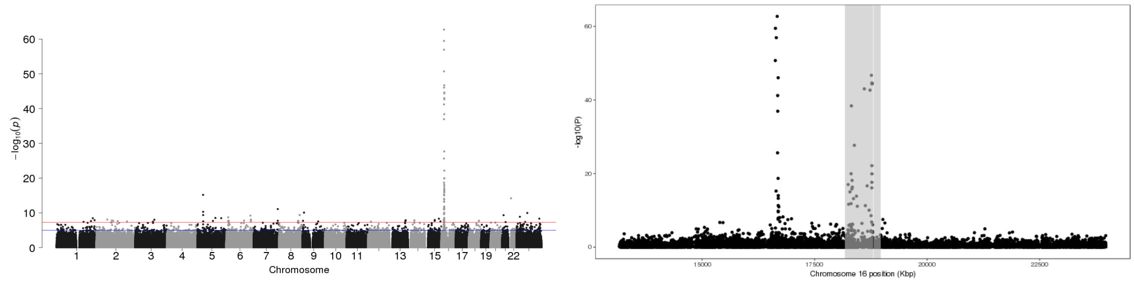


Figure 3: P-values of association between SNP genotypes and duplication copy number across the genome (left), and around the 95% confidence interval (highlighted in gray, right).

All coordinates refer to GRCh37.

### CNV_16_16725100_16730200

Insertion Location 95% Confidence Region:

16:18244090-19090721

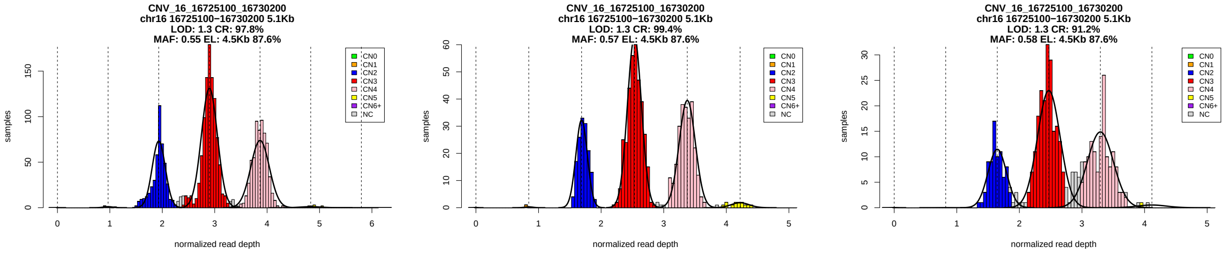


Figure 1: Copy number genotypes in the ASD, ID, and SCZ cohorts.

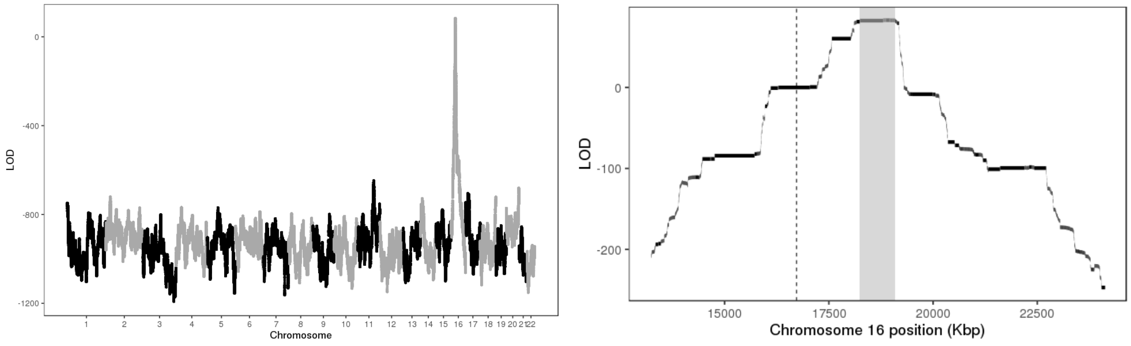


Figure 2: LOD scores of insertion locations across the genome (left) and around the 95% confidence interval (highlighted in gray, right).

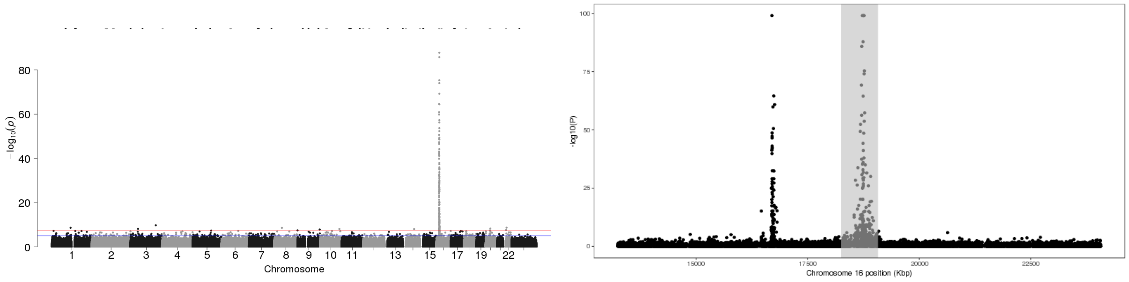


Figure 3: P-values of association between SNP genotypes and duplication copy number across the genome (left), and around the 95% confidence interval (highlighted in gray, right).

All coordinates refer to GRCh37.

### CNV_16_16745112_16746849

Insertion Location 95% Confidence Region:

16:18158085-19181300

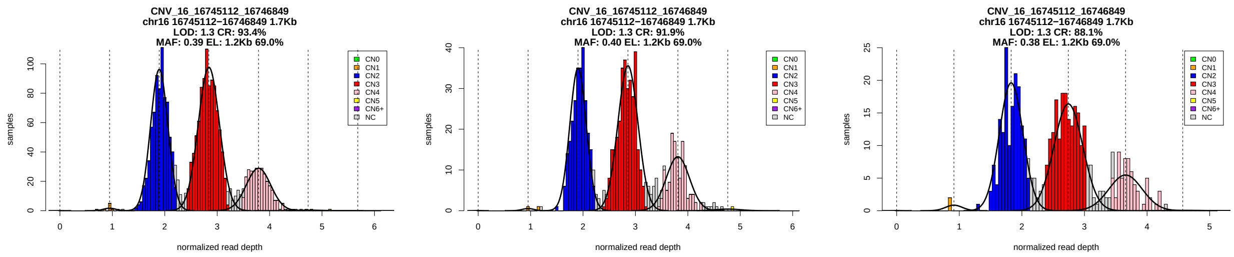


Figure 1: Copy number genotypes in the ASD, ID, and SCZ cohorts.

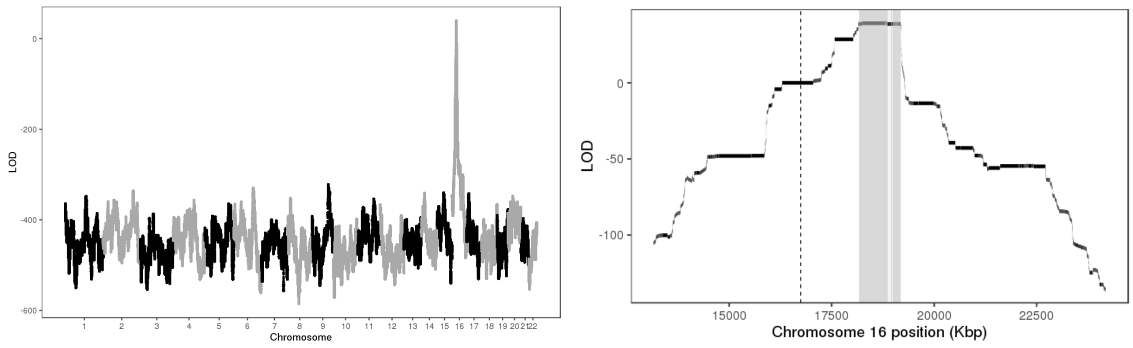


Figure 2: LOD scores of insertion locations across the genome (left) and around the 95% confidence interval (highlighted in gray, right).

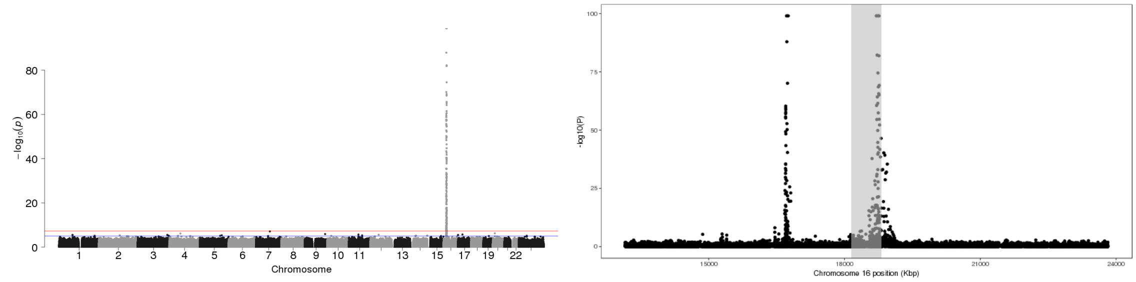


Figure 3: P-values of association between SNP genotypes and duplication copy number across the genome (left), and around the 95% confidence interval (highlighted in gray, right).

All coordinates refer to GRCh37.

### CNV_16_18615000_18635104

Insertion Location 95% Confidence Region:

16:16291971-17208426

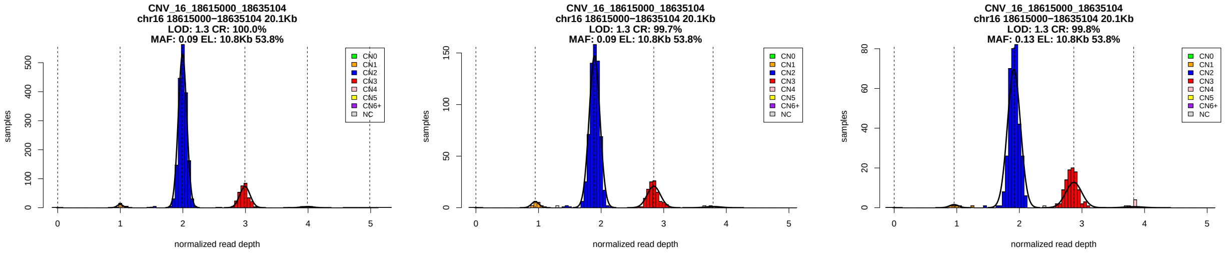


Figure 1: Copy number genotypes in the ASD, ID, and SCZ cohorts.

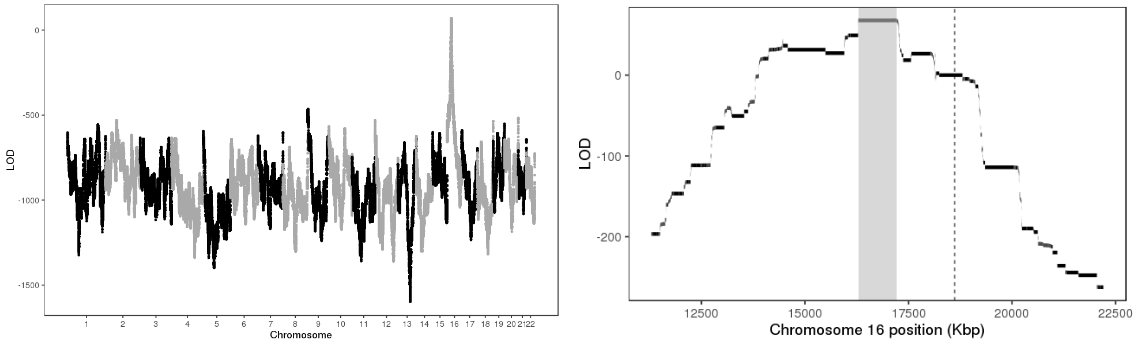


Figure 2: LOD scores of insertion locations across the genome (left) and around the 95% confidence interval (highlighted in gray, right).

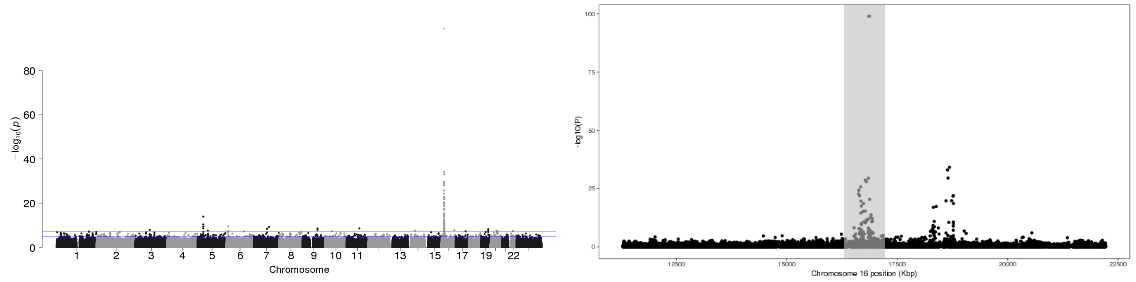


Figure 3: P-values of association between SNP genotypes and duplication copy number across the genome (left), and around the 95% confidence interval (highlighted in gray, right).

All coordinates refer to GRCh37.

### CNV_16_70155387_70200078

Insertion Location 95% Confidence Region:

16:74171216-74761843

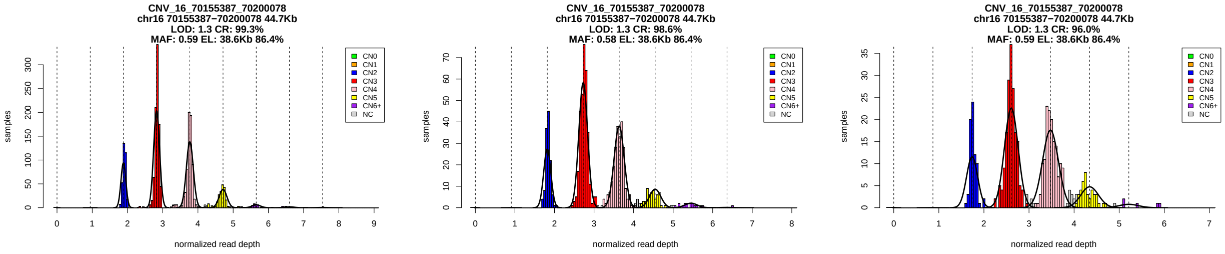


Figure 1: Copy number genotypes in the ASD, ID, and SCZ cohorts.

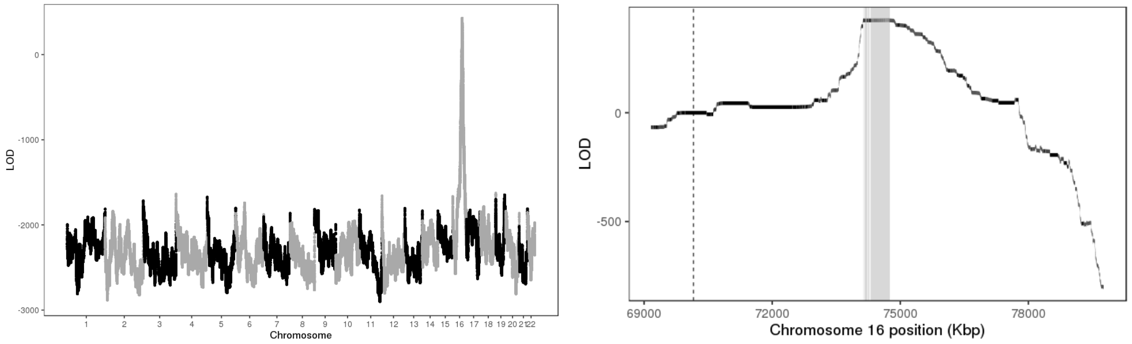


Figure 2: LOD scores of insertion locations across the genome (left) and around the 95% confidence interval (highlighted in gray, right).

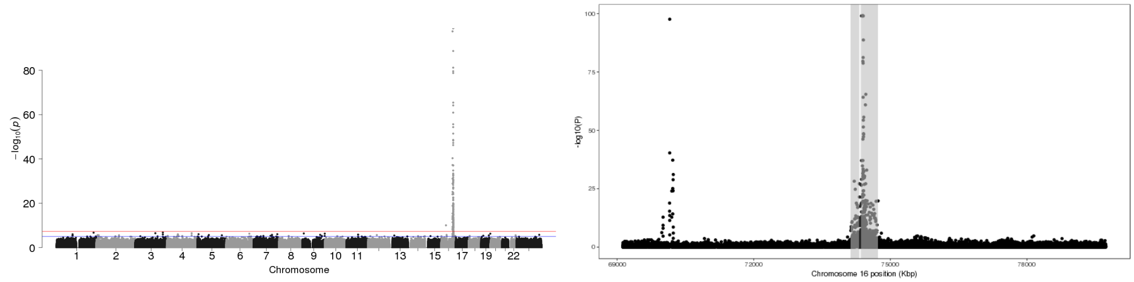


Figure 3: P-values of association between SNP genotypes and duplication copy number across the genome (left), and around the 95% confidence interval (highlighted in gray, right).

All coordinates refer to GRCh37.

### CNV_16_70977503_70980000

Insertion Location 95% Confidence Region:

1:145514053-150506263

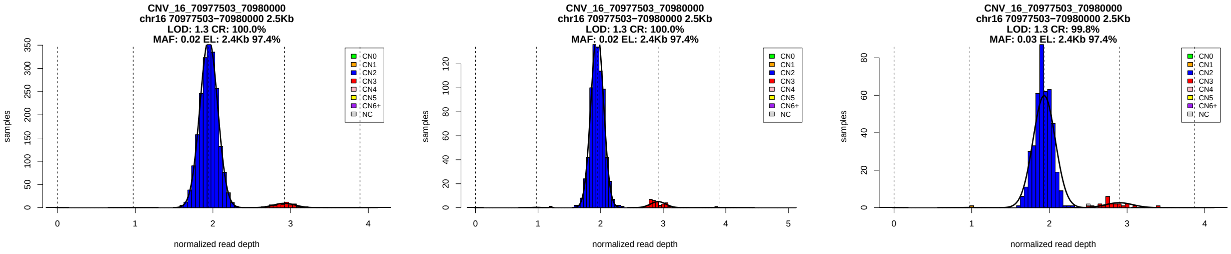


Figure 1: Copy number genotypes in the ASD, ID, and SCZ cohorts.

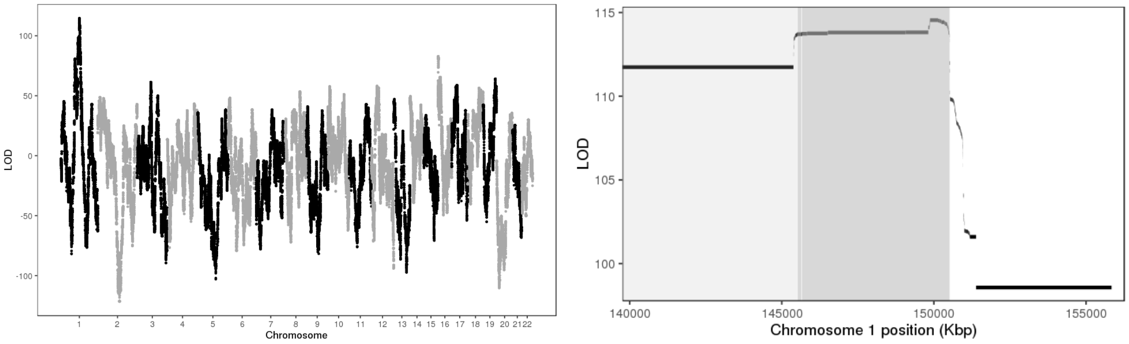


Figure 2: LOD scores of insertion locations across the genome (left) and around the 95% confidence interval (highlighted in gray, right).

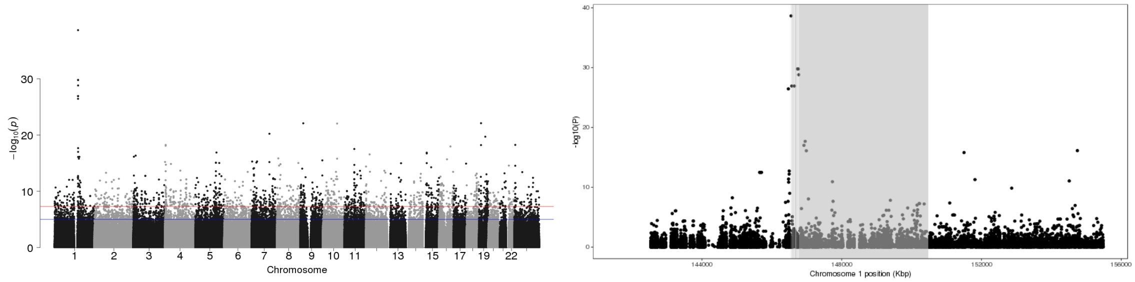


Figure 3: P-values of association between SNP genotypes and duplication copy number across the genome (left), and around the 95% confidence interval (highlighted in gray, right).

All coordinates refer to GRCh37.

### CNV_17_16656700_16709271

Insertion Location 95% Confidence Region:

17:17496869-18578832

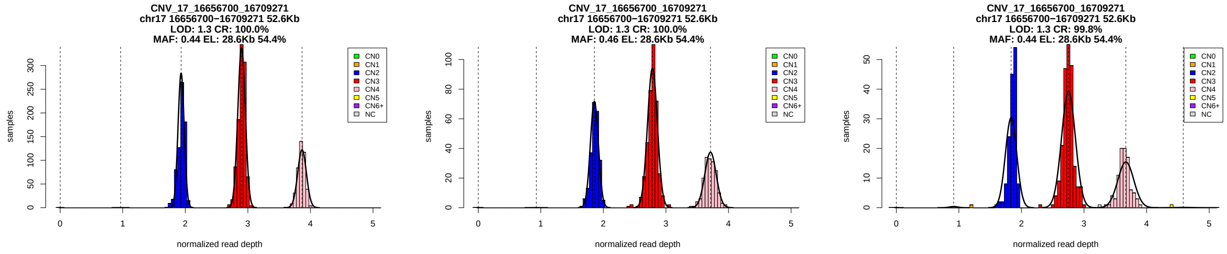


Figure 1: Copy number genotypes in the ASD, ID, and SCZ cohorts.

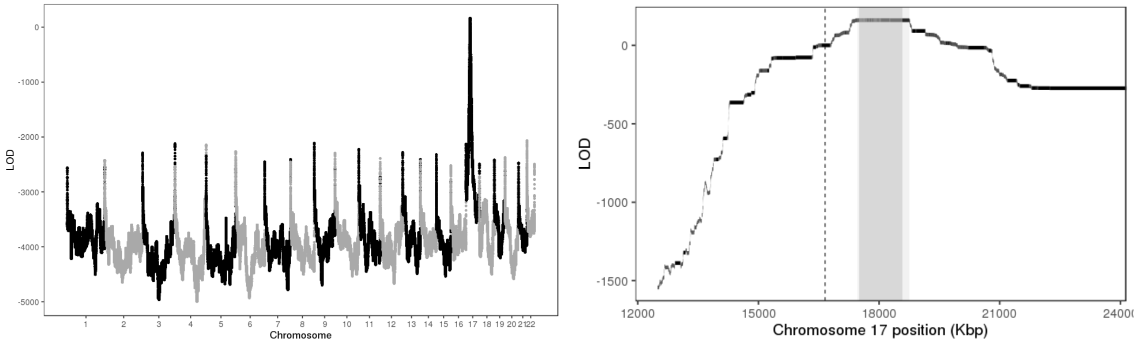


Figure 2: LOD scores of insertion locations across the genome (left) and around the 95% confidence interval (highlighted in gray, right).

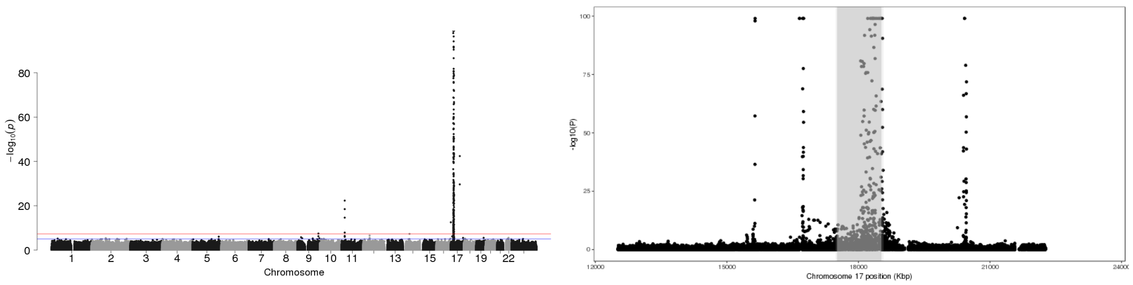


Figure 3: P-values of association between SNP genotypes and duplication copy number across the genome (left), and around the 95% confidence interval (highlighted in gray, right).

All coordinates refer to GRCh37.

### CNV_17_16710600_16725700

Insertion Location 95% Confidence Region:

17:17325046-17340512

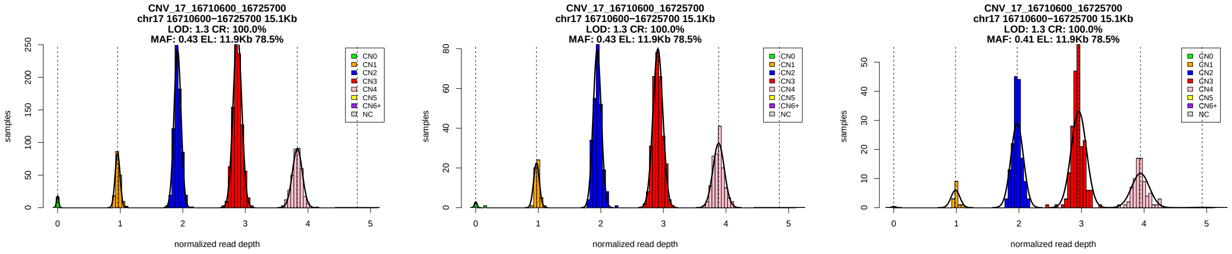


Figure 1: Copy number genotypes in the ASD, ID, and SCZ cohorts.

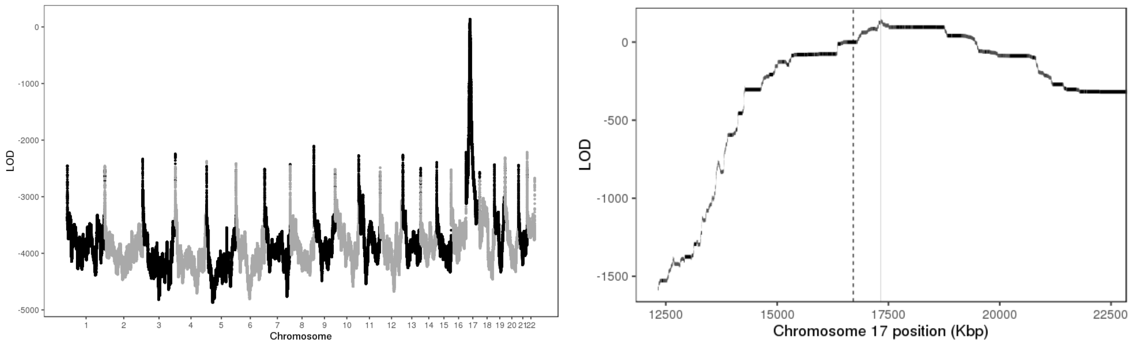


Figure 2: LOD scores of insertion locations across the genome (left) and around the 95% confidence interval (highlighted in gray, right).

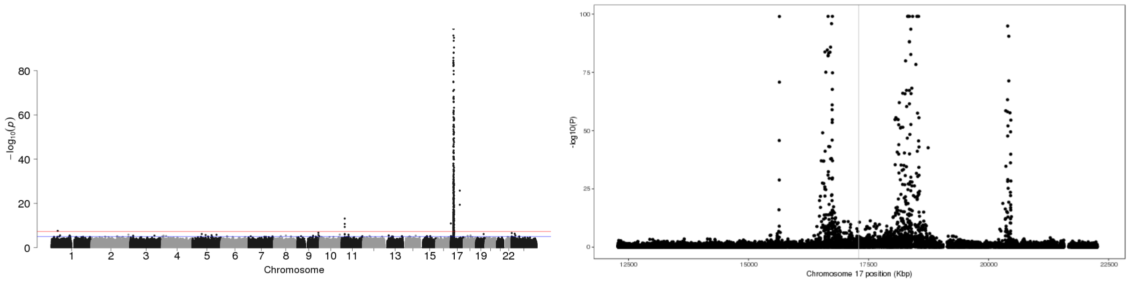


Figure 3: P-values of association between SNP genotypes and duplication copy number across the genome (left), and around the 95% confidence interval (highlighted in gray, right).

All coordinates refer to GRCh37.

### CNV_17_16744849_16747500

Insertion Location 95% Confidence Region:

17:17322196-18177661

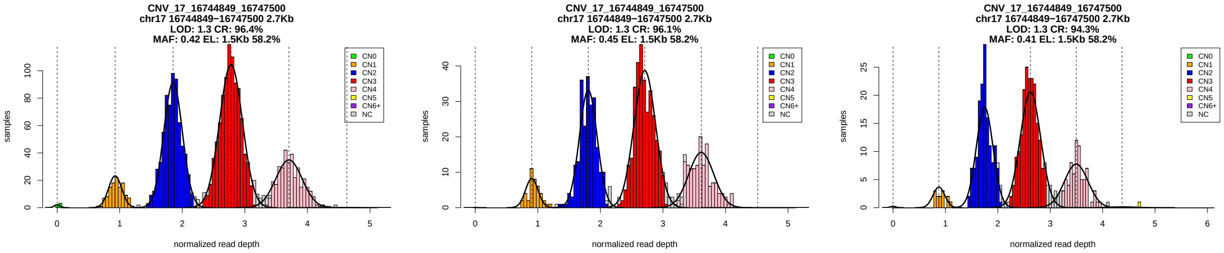


Figure 1: Copy number genotypes in the ASD, ID, and SCZ cohorts.

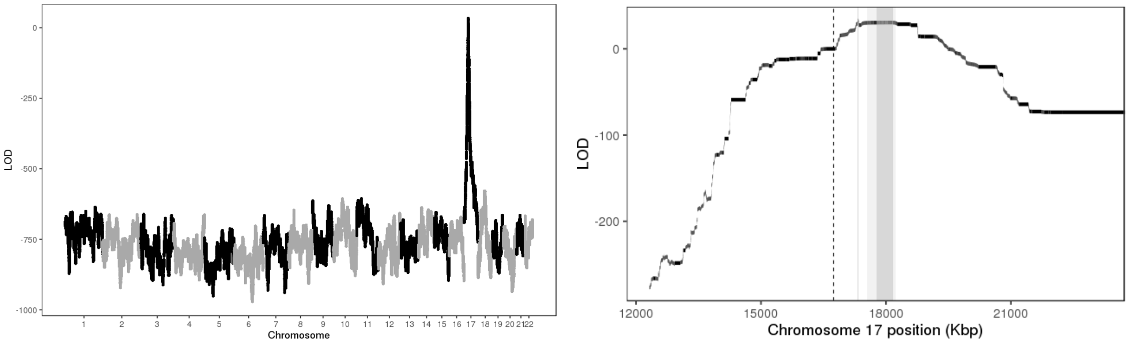


Figure 2: LOD scores of insertion locations across the genome (left) and around the 95% confidence interval (highlighted in gray, right).

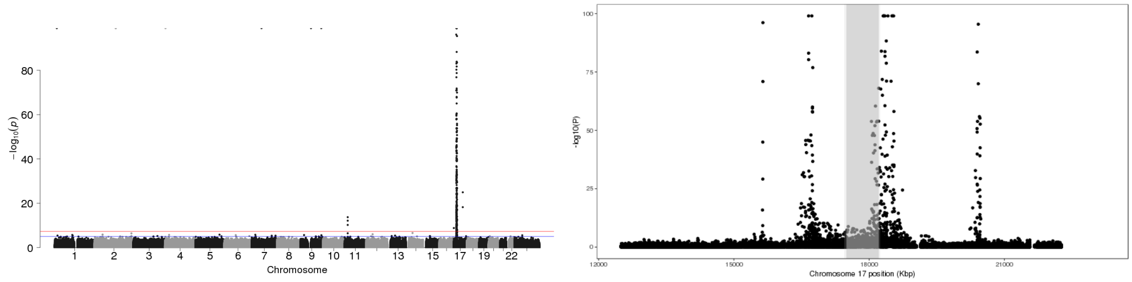


Figure 3: P-values of association between SNP genotypes and duplication copy number across the genome (left), and around the 95% confidence interval (highlighted in gray, right).

All coordinates refer to GRCh37.

### CNV_17_18624874_18633247

Insertion Location 95% Confidence Region:

17:15382339-16302201

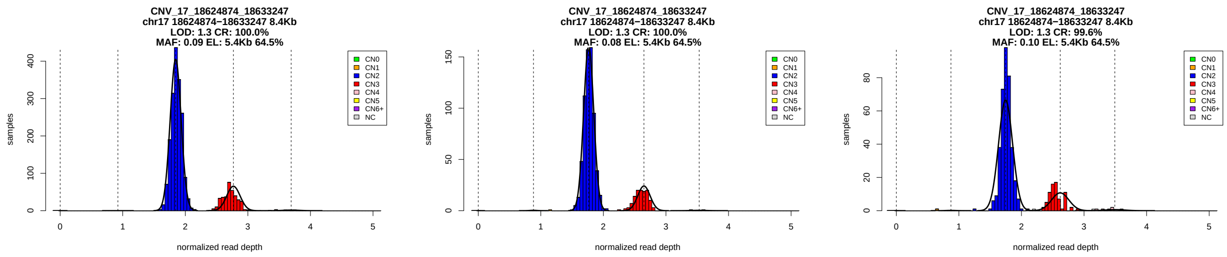


Figure 1: Copy number genotypes in the ASD, ID, and SCZ cohorts.

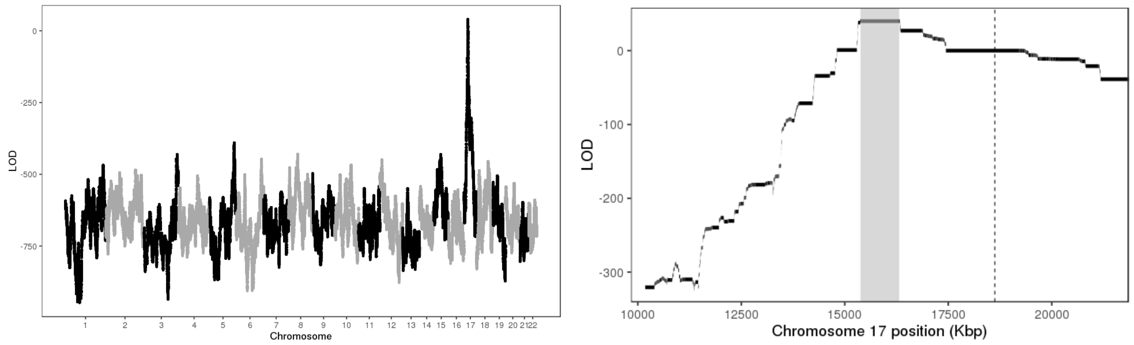


Figure 2: LOD scores of insertion locations across the genome (left) and around the 95% confidence interval (highlighted in gray, right).

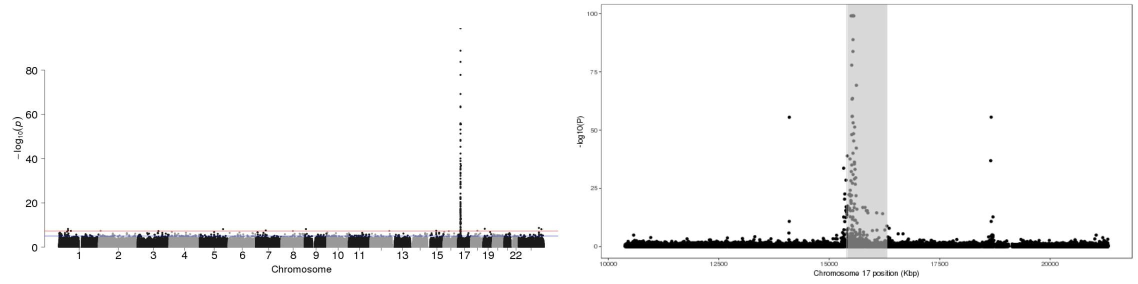


Figure 3: P-values of association between SNP genotypes and duplication copy number across the genome (left), and around the 95% confidence interval (highlighted in gray, right).

All coordinates refer to GRCh37.

### CNV_17_18687000_18691000

Insertion Location 95% Confidence Region:

17:15402638-16866164

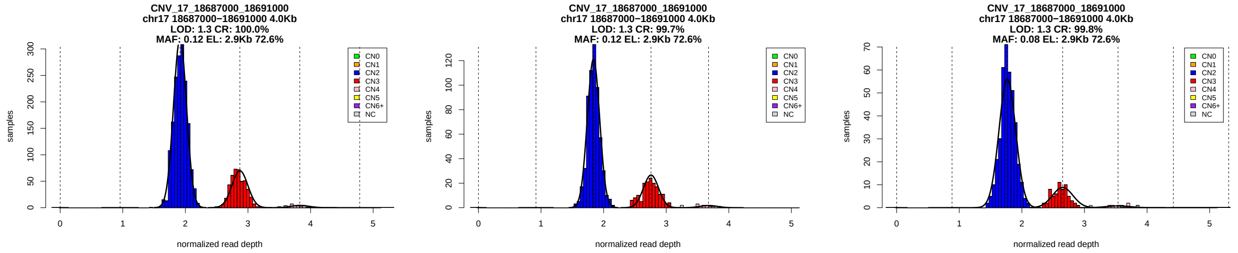


Figure 1: Copy number genotypes in the ASD, ID, and SCZ cohorts.

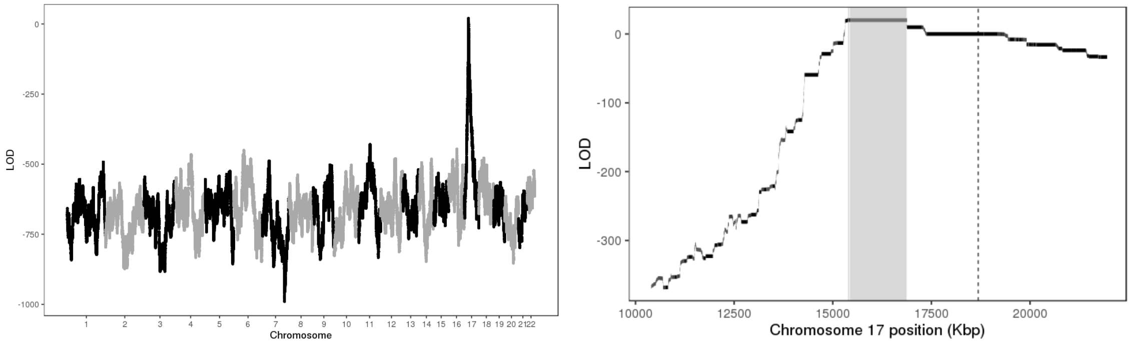


Figure 2: LOD scores of insertion locations across the genome (left) and around the 95% confidence interval (highlighted in gray, right).

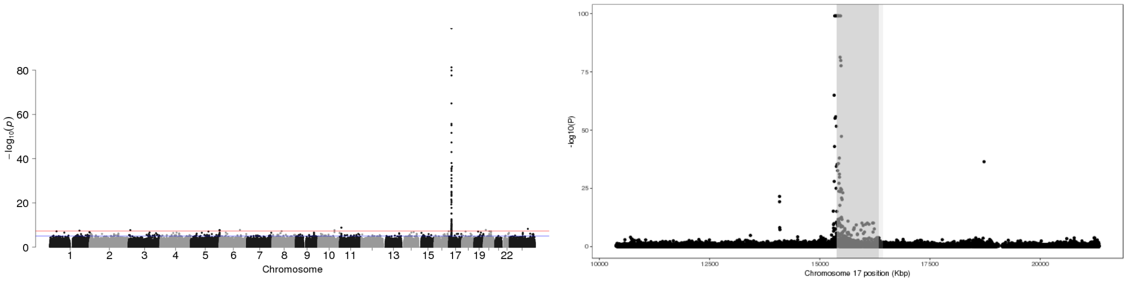


Figure 3: P-values of association between SNP genotypes and duplication copy number across the genome (left), and around the 95% confidence interval (highlighted in gray, right).

All coordinates refer to GRCh37.

### CNV_17_20443387_20446620

Insertion Location 95% Confidence Region:

17:17848286-18753870

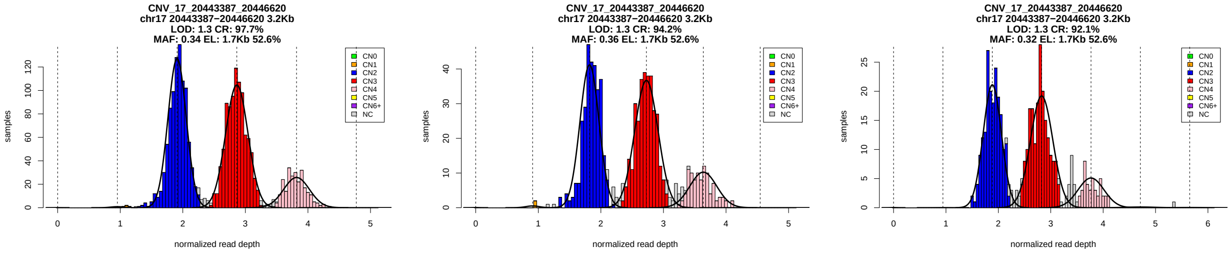


Figure 1: Copy number genotypes in the ASD, ID, and SCZ cohorts.

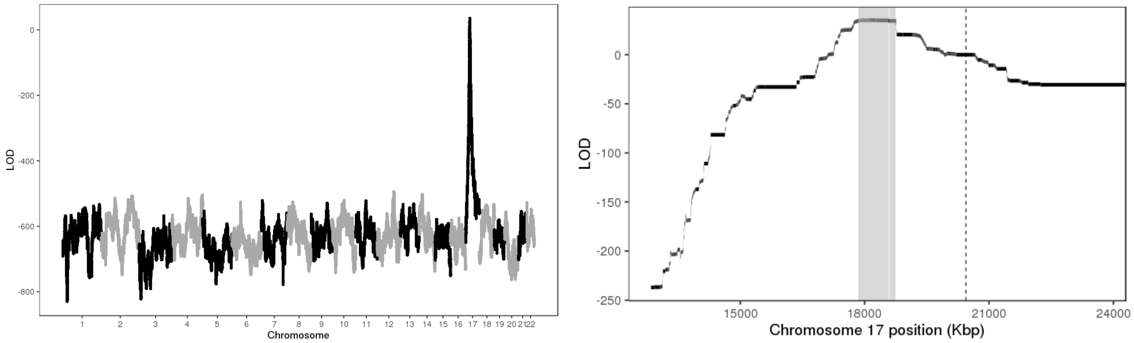


Figure 2: LOD scores of insertion locations across the genome (left) and around the 95% confidence interval (highlighted in gray, right).

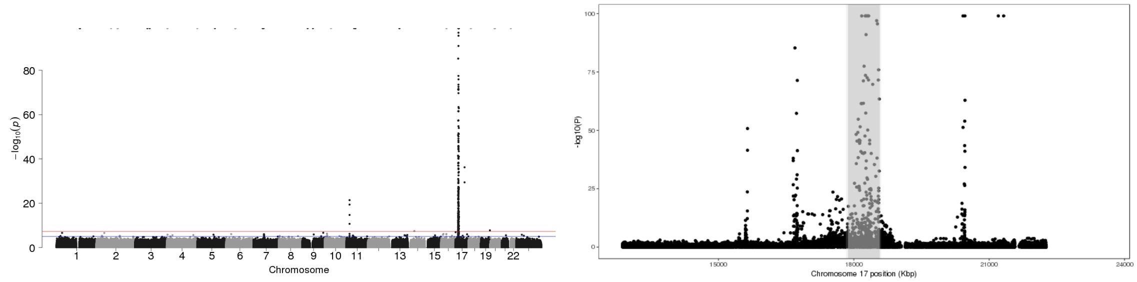


Figure 3: P-values of association between SNP genotypes and duplication copy number across the genome (left), and around the 95% confidence interval (highlighted in gray, right).

All coordinates refer to GRCh37.

### CNV_17_36387600_36399500

Insertion Location 95% Confidence Region:

17:34916879-35267969

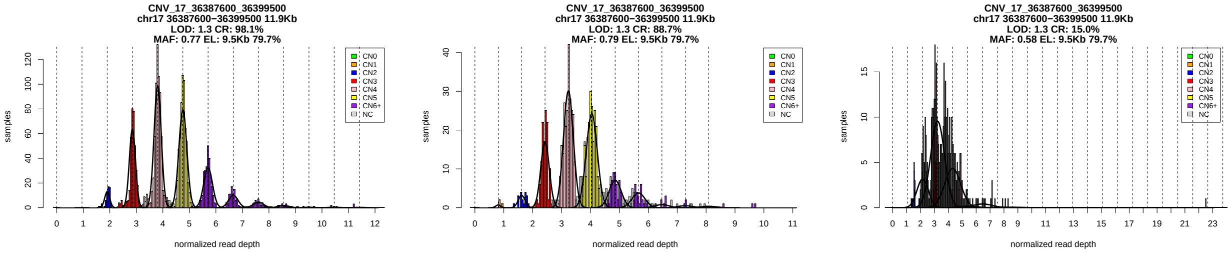


Figure 1: Copy number genotypes in the ASD, ID, and SCZ cohorts.

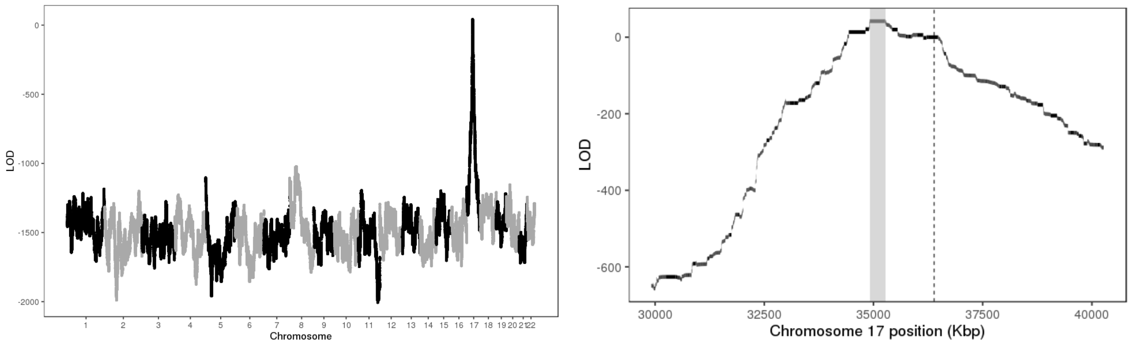


Figure 2: LOD scores of insertion locations across the genome (left) and around the 95% confidence interval (highlighted in gray, right).

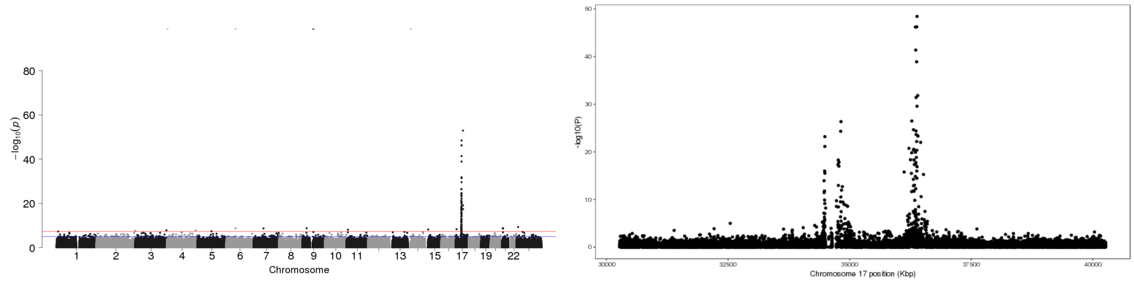


Figure 3: P-values of association between SNP genotypes and duplication copy number across the genome (left), and around the 95% confidence interval (highlighted in gray, right).

All coordinates refer to GRCh37.

### CNV_17_43655745_43662300

Insertion Location 95% Confidence Region:

17:44076063-44819519

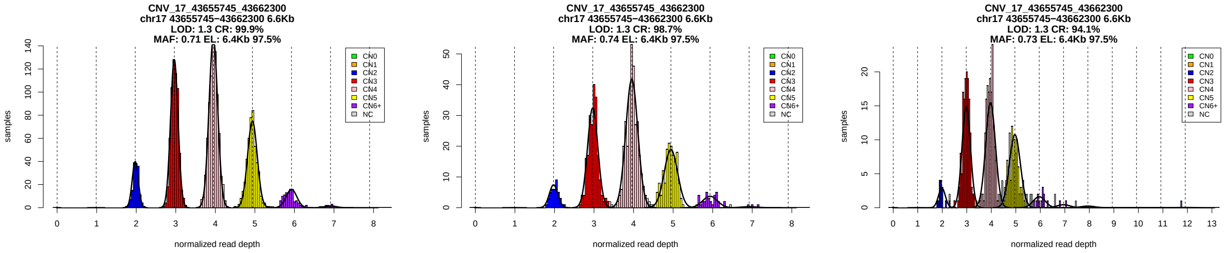


Figure 1: Copy number genotypes in the ASD, ID, and SCZ cohorts.

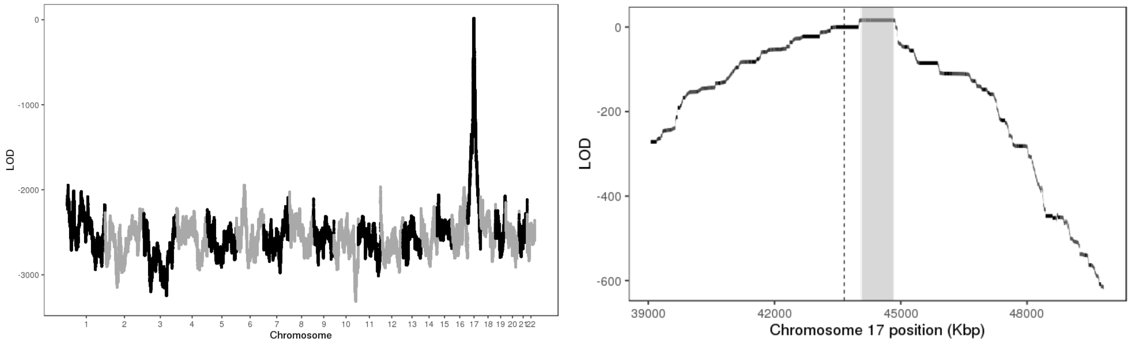


Figure 2: LOD scores of insertion locations across the genome (left) and around the 95% confidence interval (highlighted in gray, right).

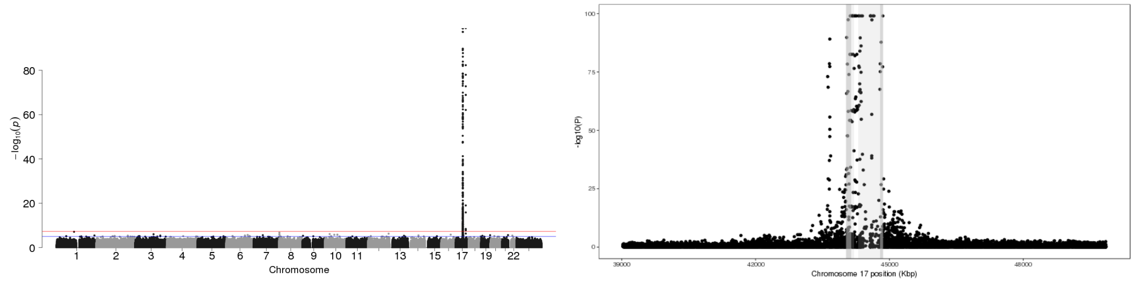


Figure 3: P-values of association between SNP genotypes and duplication copy number across the genome (left), and around the 95% confidence interval (highlighted in gray, right).

All coordinates refer to GRCh37.

### CNV_17_62915465_62918489

Insertion Location 95% Confidence Region:

17:44128876-44855683

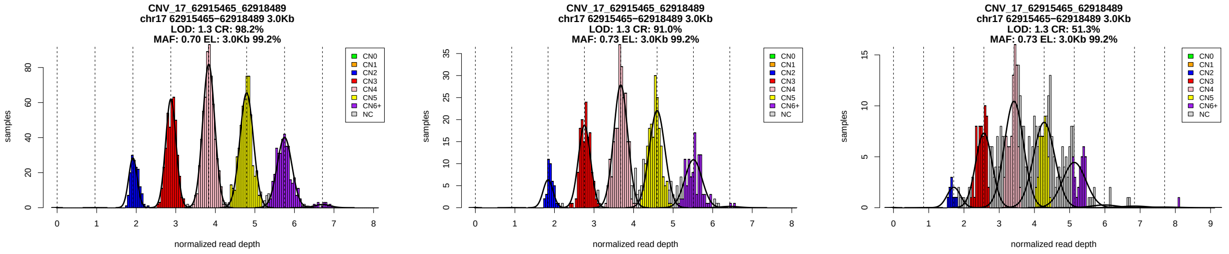


Figure 1: Copy number genotypes in the ASD, ID, and SCZ cohorts.

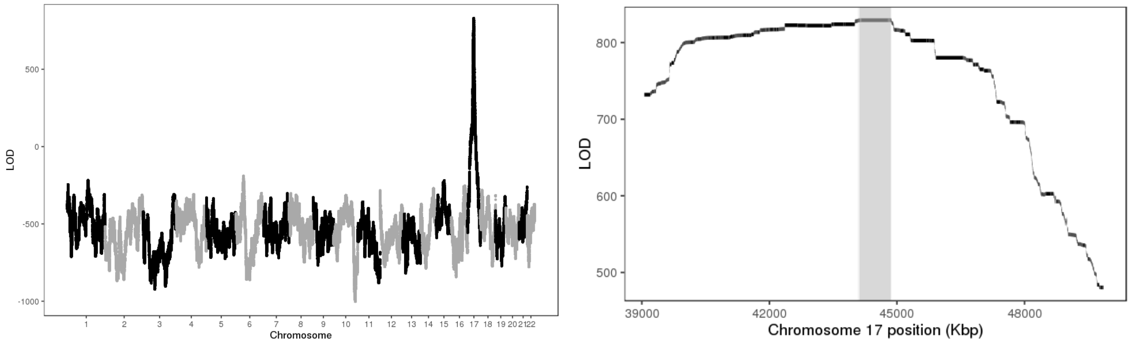


Figure 2: LOD scores of insertion locations across the genome (left) and around the 95% confidence interval (highlighted in gray, right).

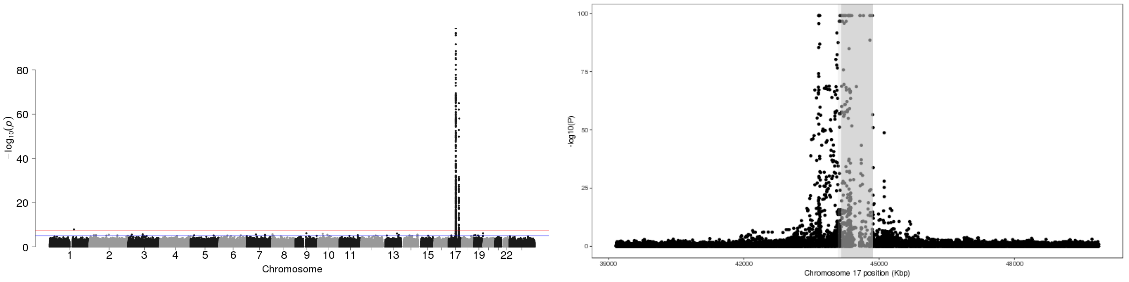


Figure 3: P-values of association between SNP genotypes and duplication copy number across the genome (left), and around the 95% confidence interval (highlighted in gray, right).

All coordinates refer to GRCh37.

### CNV_19_50583100_50594147

Insertion Location 95% Confidence Region:

19:50948550-51244631

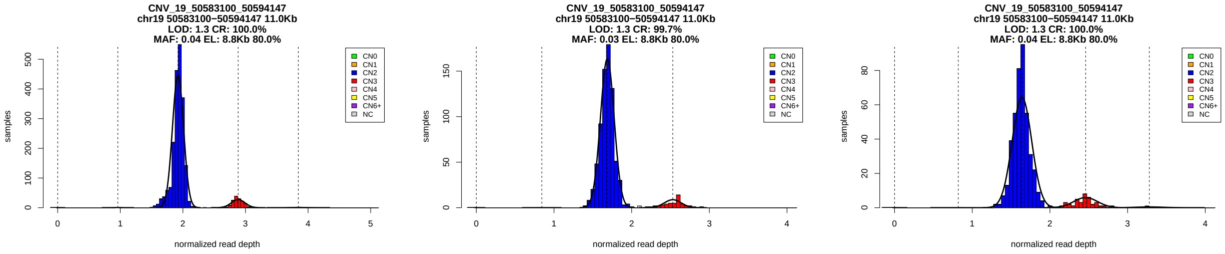


Figure 1: Copy number genotypes in the ASD, ID, and SCZ cohorts.

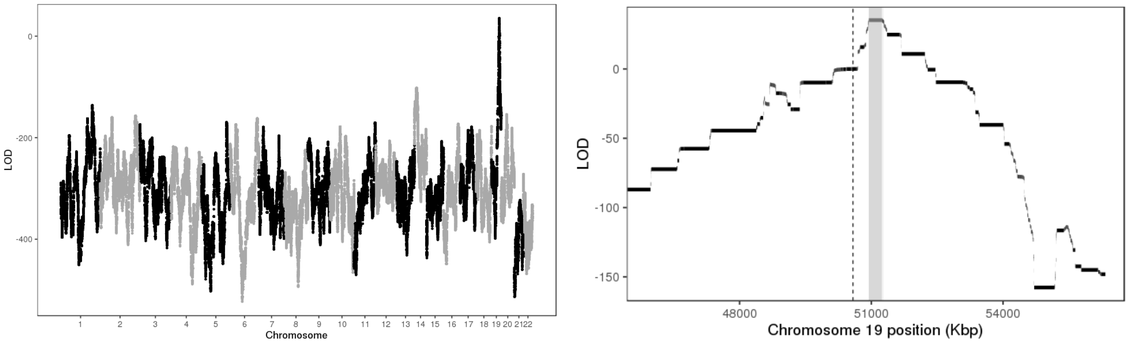


Figure 2: LOD scores of insertion locations across the genome (left) and around the 95% confidence interval (highlighted in gray, right).

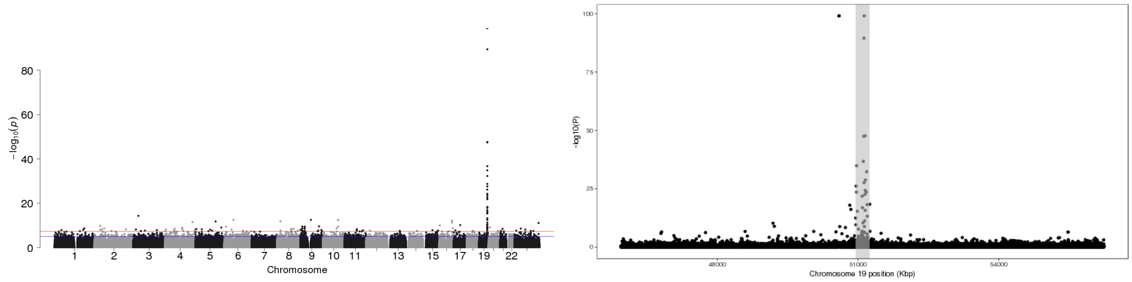


Figure 3: P-values of association between SNP genotypes and duplication copy number across the genome (left), and around the 95% confidence interval (highlighted in gray, right).

All coordinates refer to GRCh37.

### CNV_20_29638759_29641400

Insertion Location 95% Confidence Region:

4:12083991-14123784

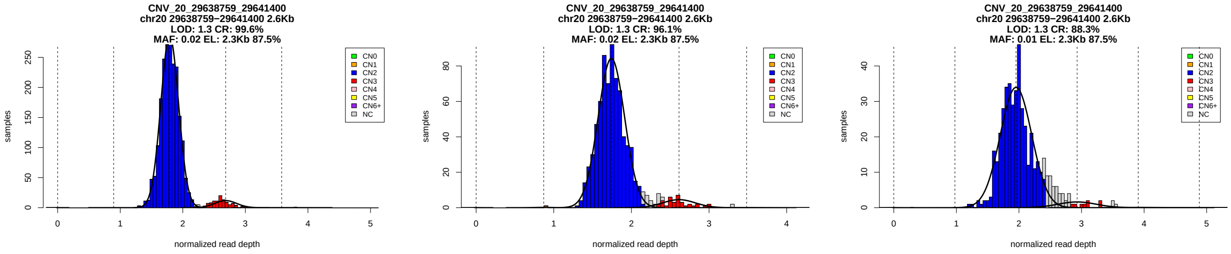


Figure 1: Copy number genotypes in the ASD, ID, and SCZ cohorts.

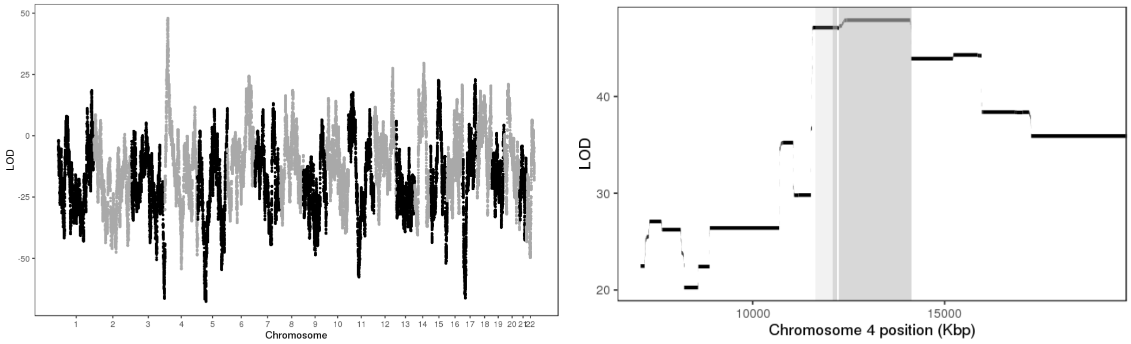


Figure 2: LOD scores of insertion locations across the genome (left) and around the 95% confidence interval (highlighted in gray, right).

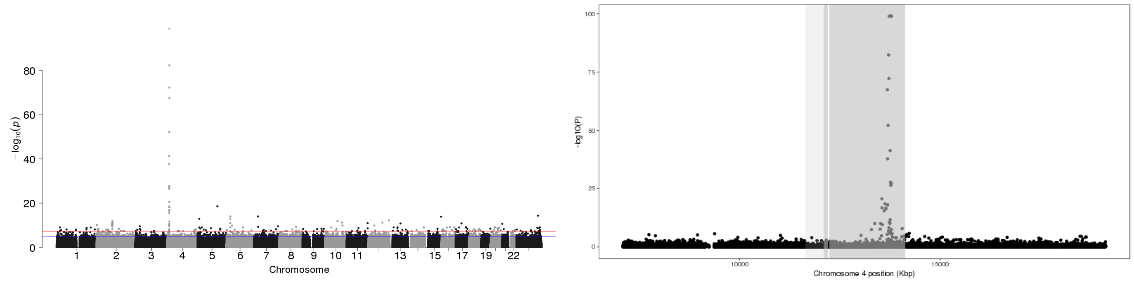


Figure 3: P-values of association between SNP genotypes and duplication copy number across the genome (left), and around the 95% confidence interval (highlighted in gray, right).

All coordinates refer to GRCh37.

### CNV_20_52475400_52484096

Insertion Location 95% Confidence Region:

16:34280305-47986157

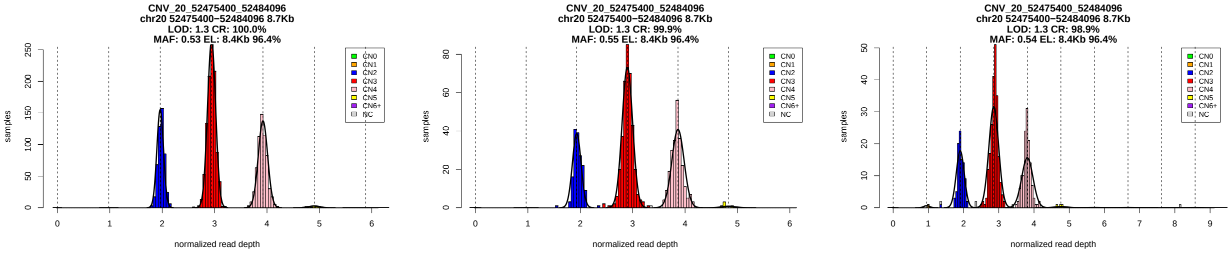


Figure 1: Copy number genotypes in the ASD, ID, and SCZ cohorts.

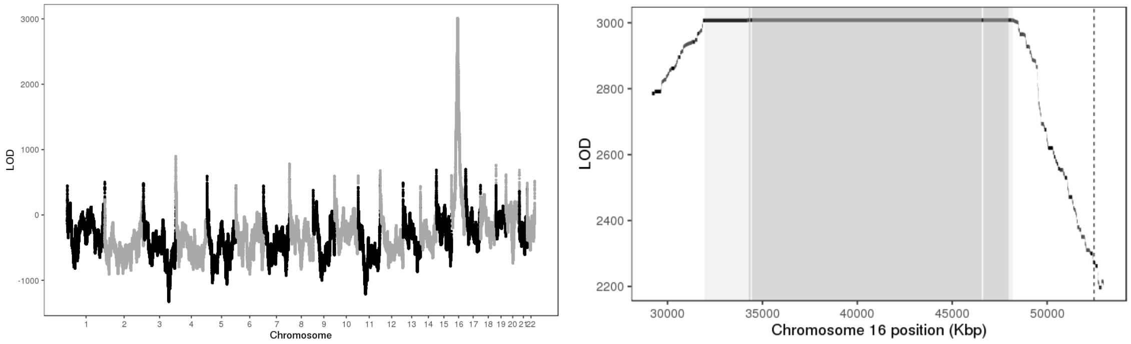


Figure 2: LOD scores of insertion locations across the genome (left) and around the 95% confidence interval (highlighted in gray, right).

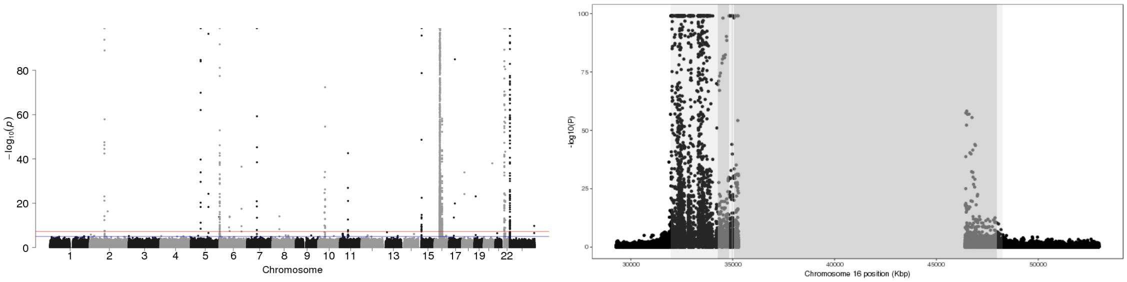


Figure 3: P-values of association between SNP genotypes and duplication copy number across the genome (left), and around the 95% confidence interval (highlighted in gray, right).

All coordinates refer to GRCh37.

### CNV_21_15436372_15437600

Insertion Location 95% Confidence Region:

13:19448715-20138252

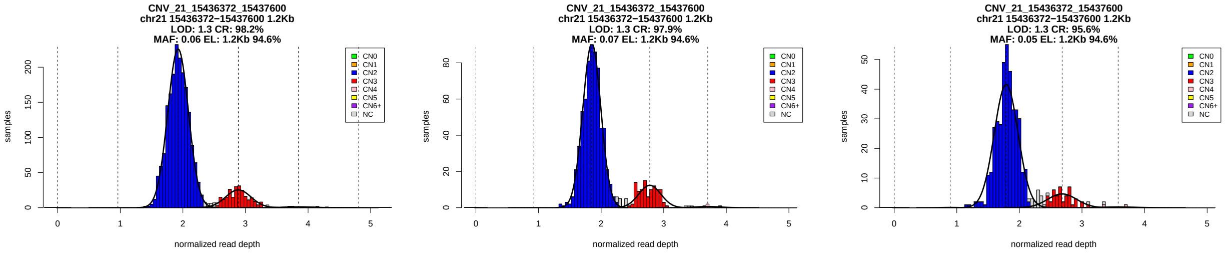


Figure 1: Copy number genotypes in the ASD, ID, and SCZ cohorts.

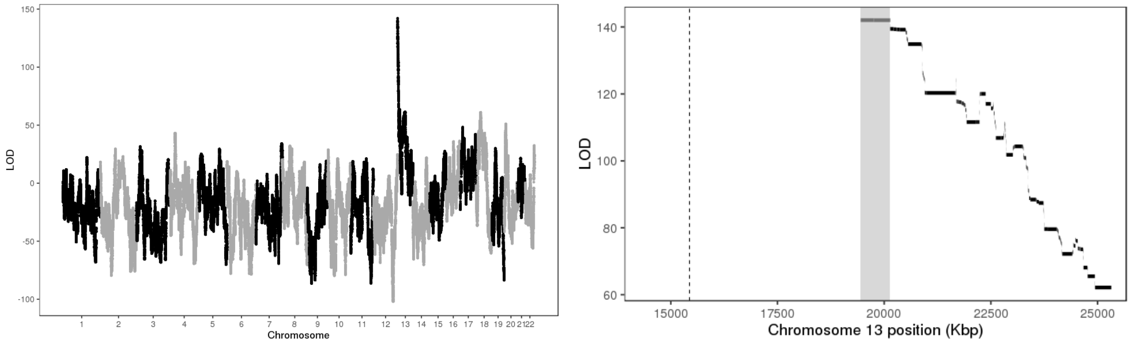


Figure 2: LOD scores of insertion locations across the genome (left) and around the 95% confidence interval (highlighted in gray, right).

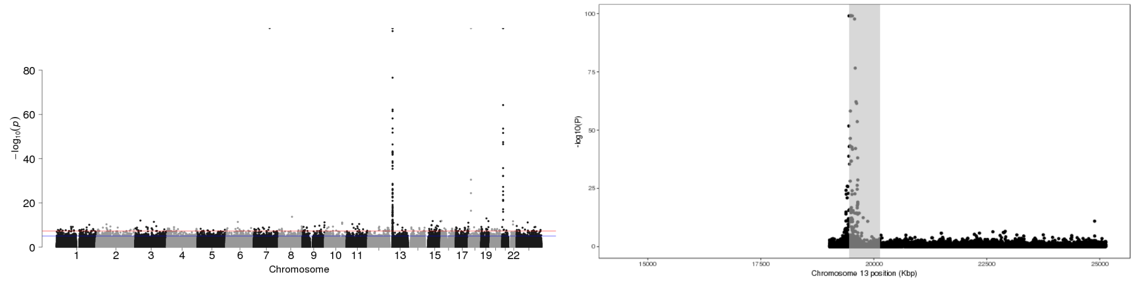


Figure 3: P-values of association between SNP genotypes and duplication copy number across the genome (left), and around the 95% confidence interval (highlighted in gray, right).

All coordinates refer to GRCh37.

# GS_SD_M2_22_20368704_20472334_22_21812477_21917116

Insertion Location 95% Confidence Region:

22:19000061-19091977

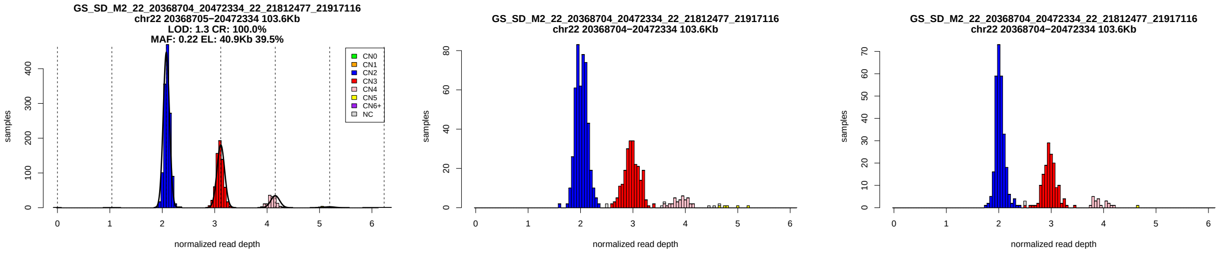


Figure 1: Copy number genotypes in the ASD, ID, and SCZ cohorts.

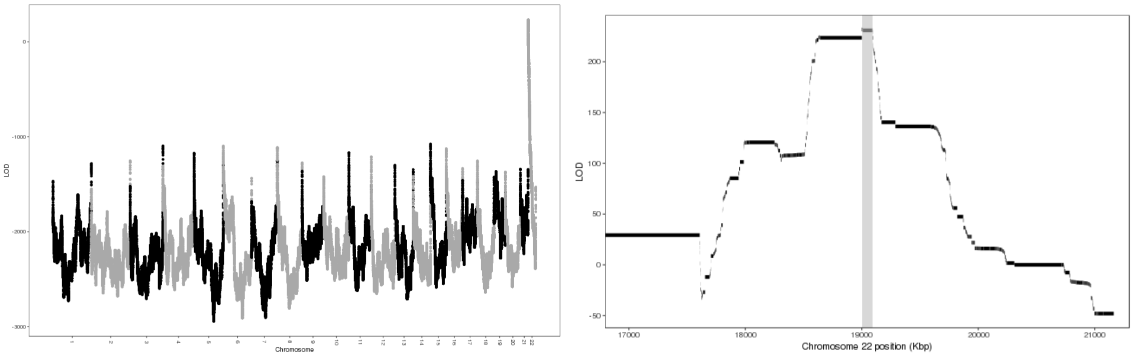


Figure 2: LOD scores of insertion locations across the genome (left) and around the 95% confidence interval (highlighted in gray, right).

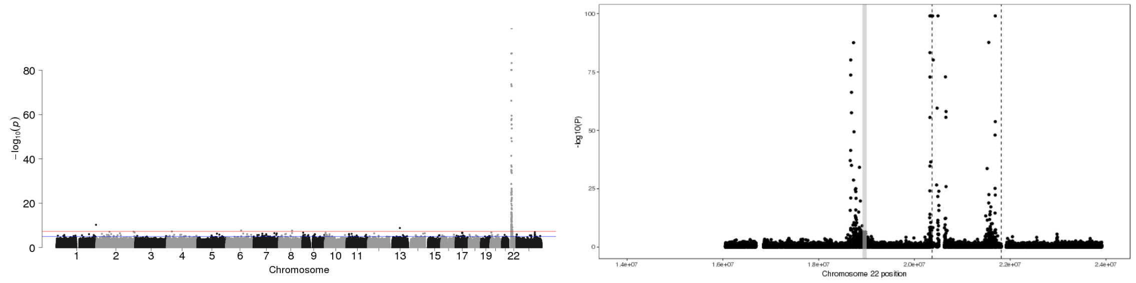


Figure 3: P-values of association between SNP genotypes and duplication copy number across the genome (left), and around the 95% confidence interval (highlighted in gray, right).

All coordinates refer to GRCh37.

# GS_SD_M2_22_20402108_20509428_22_21689945_21797378

Insertion Location 95% Confidence Region:

22:19000061-19091977

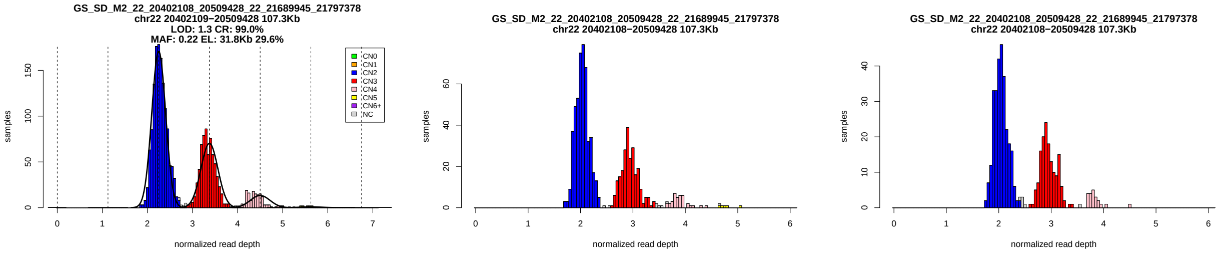


Figure 1: Copy number genotypes in the ASD, ID, and SCZ cohorts.

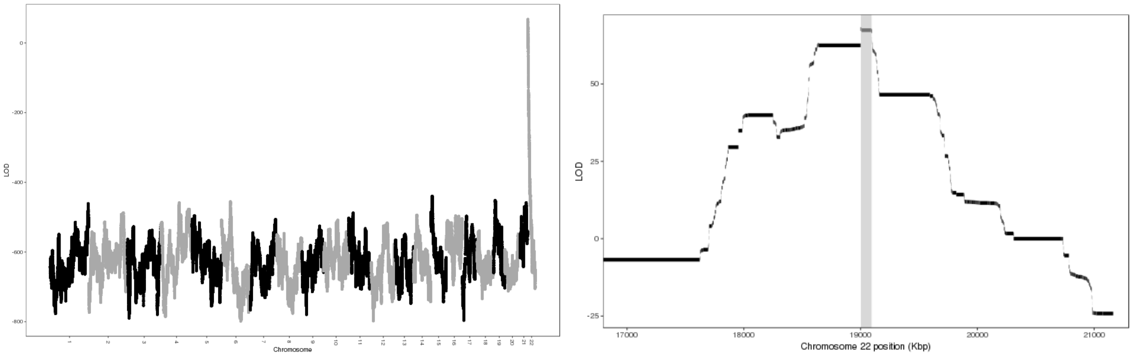


Figure 2: LOD scores of insertion locations across the genome (left) and around the 95% confidence interval (highlighted in gray, right).

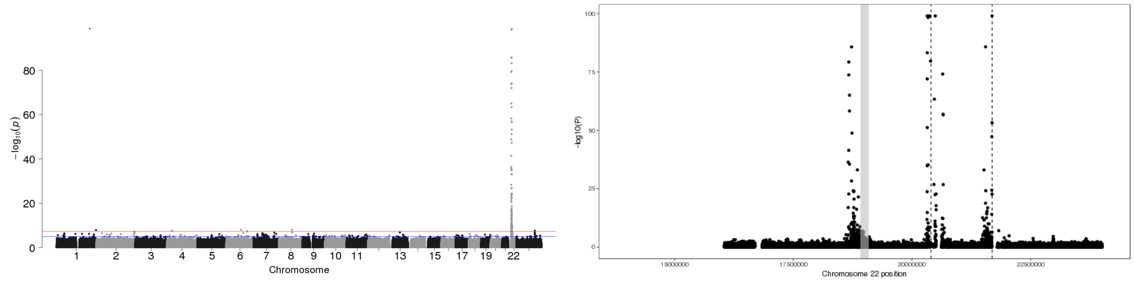


Figure 3: P-values of association between SNP genotypes and duplication copy number across the genome (left), and around the 95% confidence interval (highlighted in gray, right).

All coordinates refer to GRCh37.

### CNV_22_23771600_23774339

Insertion Location 95% Confidence Region:

7:1854348-2306708

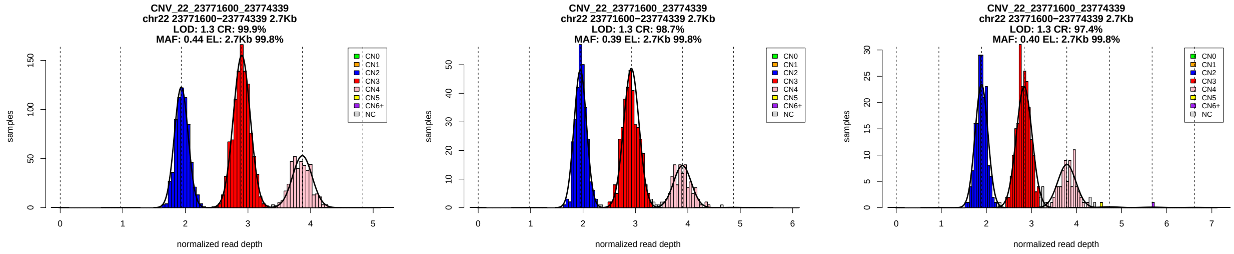


Figure 1: Copy number genotypes in the ASD, ID, and SCZ cohorts.

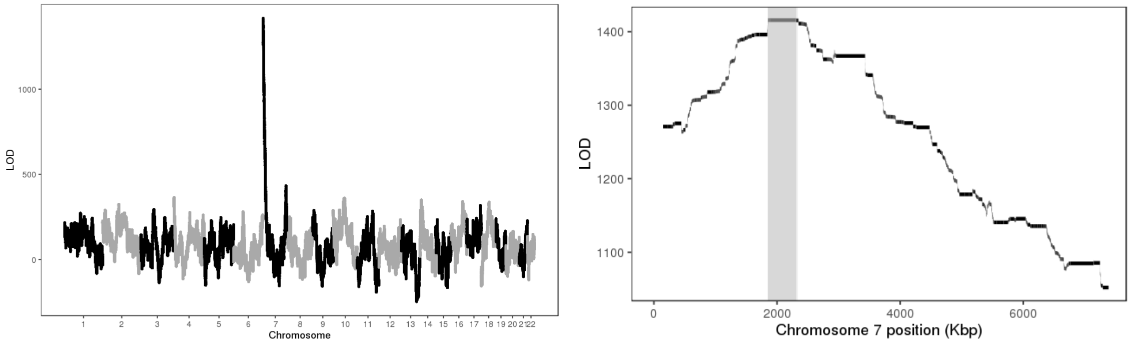


Figure 2: LOD scores of insertion locations across the genome (left) and around the 95% confidence interval (highlighted in gray, right).

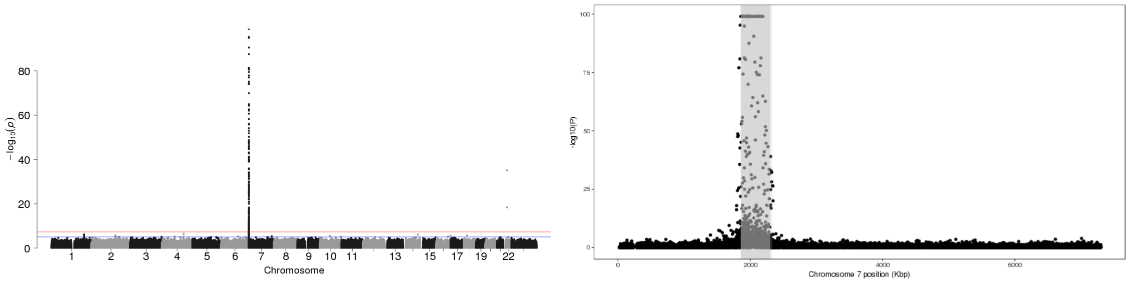


Figure 3: P-values of association between SNP genotypes and duplication copy number across the genome (left), and around the 95% confidence interval (highlighted in gray, right).

All coordinates refer to GRCh37.
