## Supplementary Figures and Tables for "Insights into dispersed duplications and complex structural mutations from whole genome sequencing 706 families"

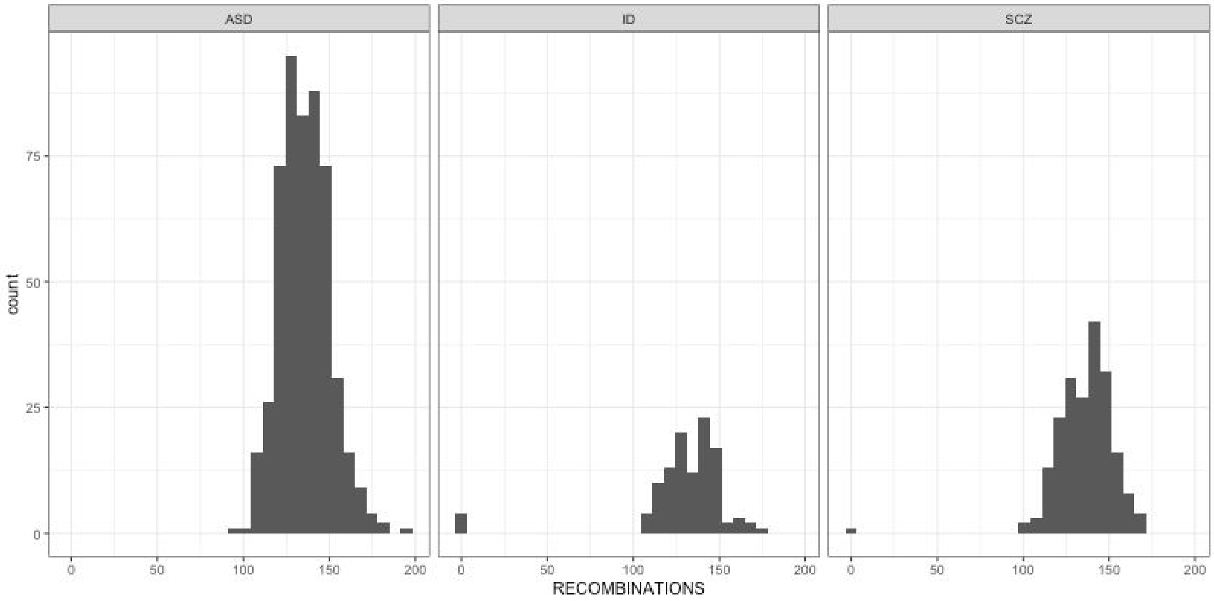


**S1 Fig.** Number of recombinations detected per sibling pair, broken down by cohort used in this study.

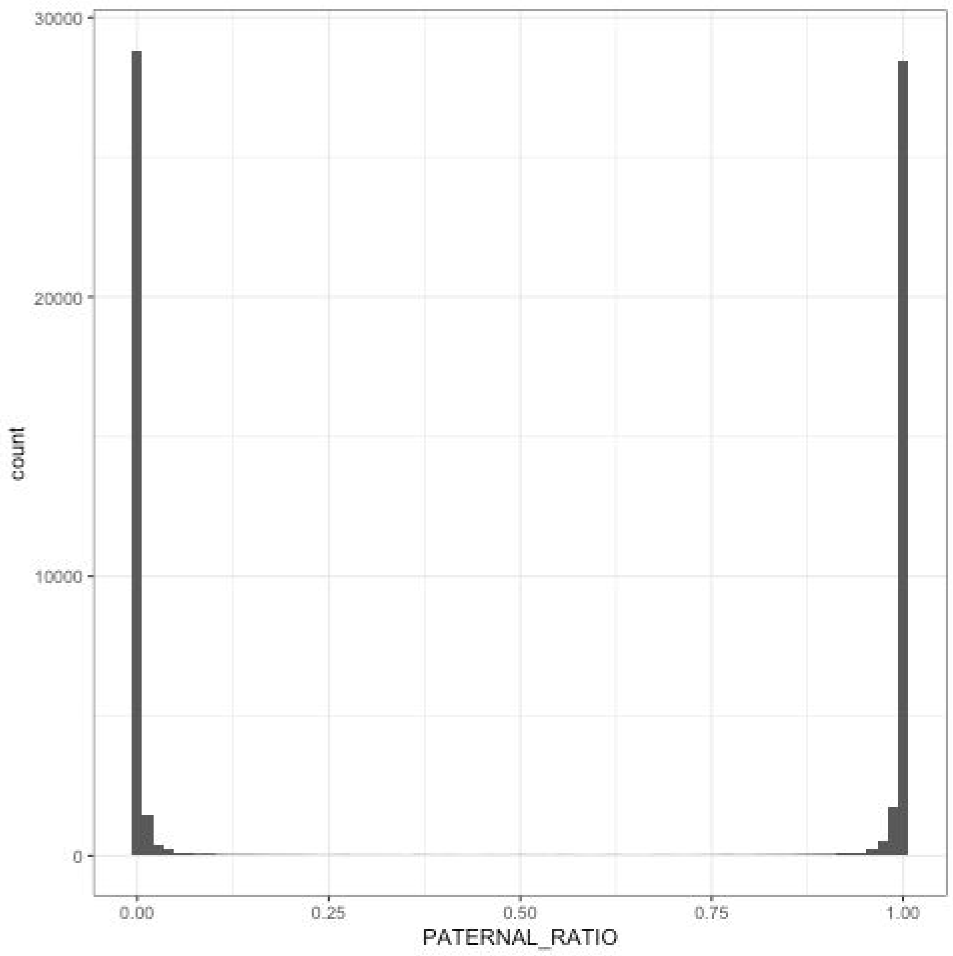


**S2 Fig.** Within IBD-1 blocks from all sibling pairs, a histogram of the ratio of SNP sites supporting a paternal origin of the shared haplotypes.

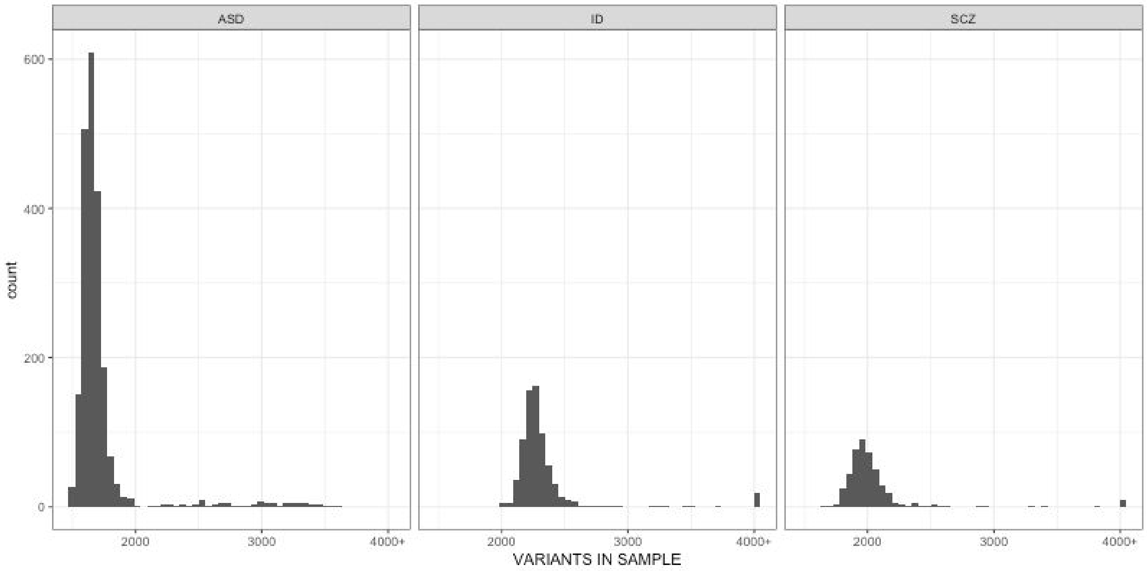


**S3 Fig.** Number of copy number variant sites called in each sample during the first round of CNV calling.

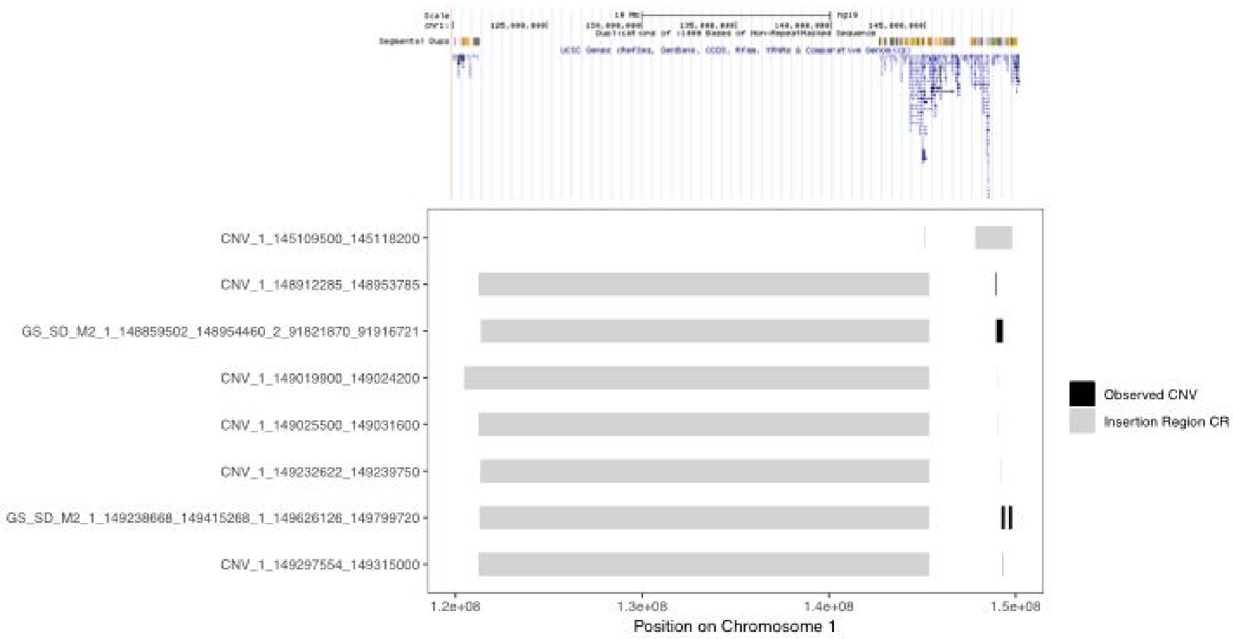


**S4 Fig.** A cluster of dispersed duplications detected in 1q21. The segment or segments at which variable copy number was detected are plotted in black; the 95% confidence regions for the locations of the dispersed segments are plotted in gray.

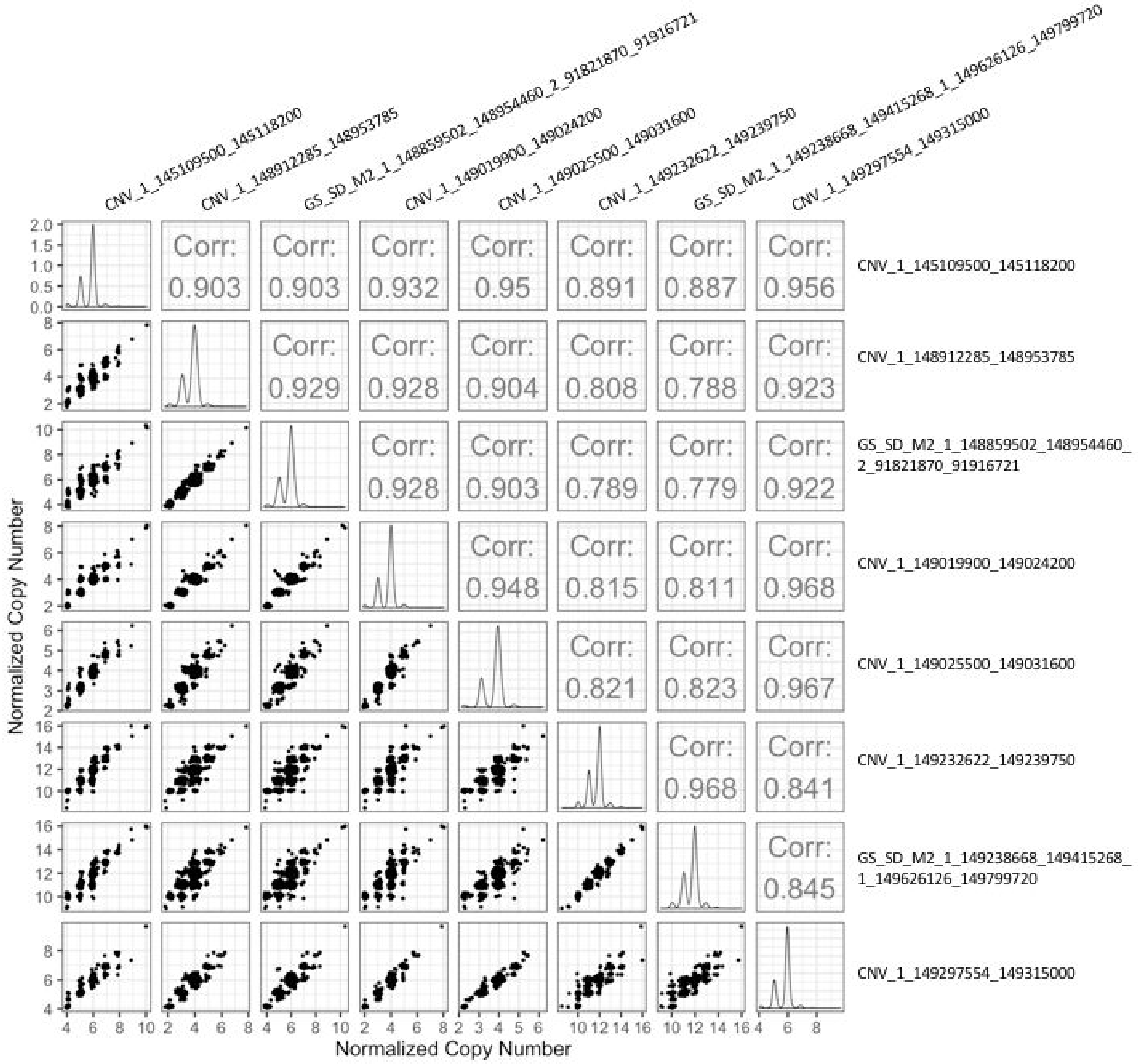


**S5 Fig.** Pairwise scatter plots of sample copy numbers in the cluster of CNV events detected in 1q21. In this case the variable segments exhibit high correlation, indicating that they are mostly participating in a single large copy number polymorphism.

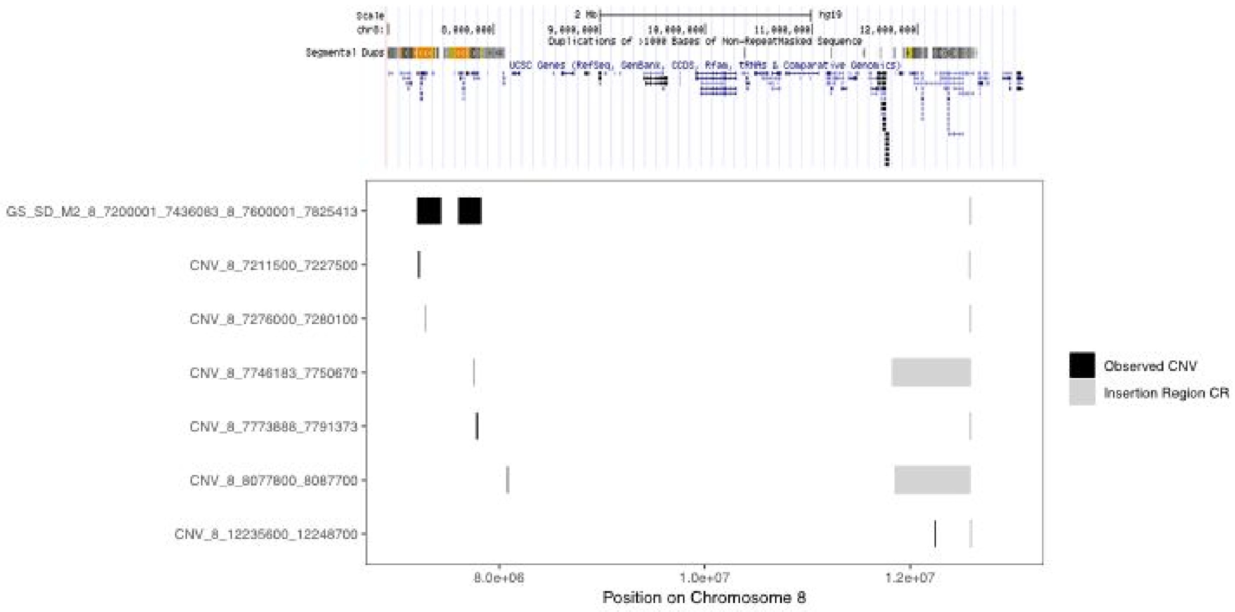


**S6 Fig.** A cluster of dispersed duplications observed at the edges of the polymorphic inversion in 8p23. The segment or segments at which variable copy number was detected are plotted in black; the 95% confidence regions for the locations of the dispersed segments are plotted in gray.

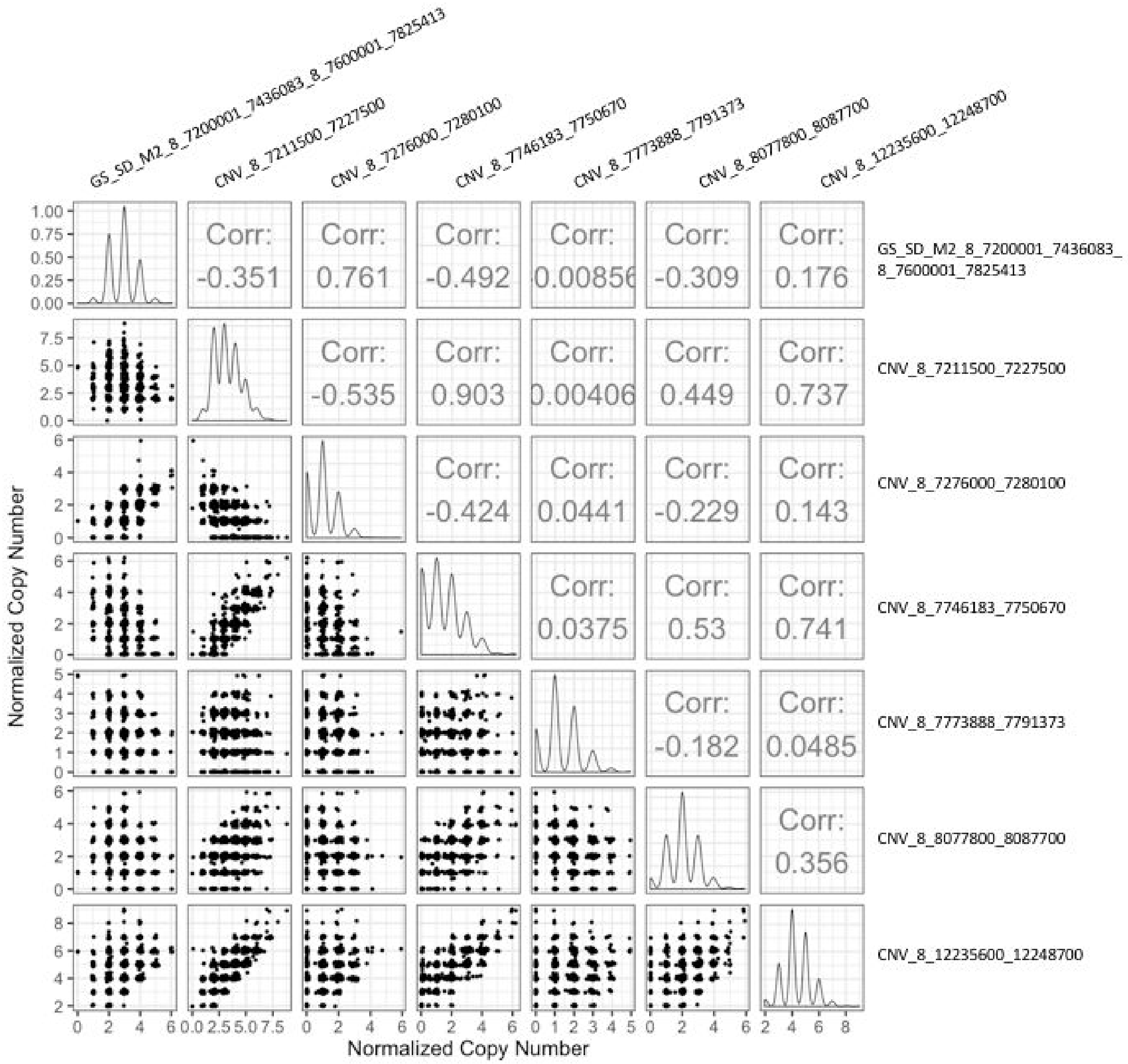


**S7 Fig.** Pairwise scatter plots of sample copy numbers in the cluster of CNV events detected in 8p23. The measured copy numbers of pairs of segments have low correlations, indicating that they are capturing different dispersed duplication polymorphism events.

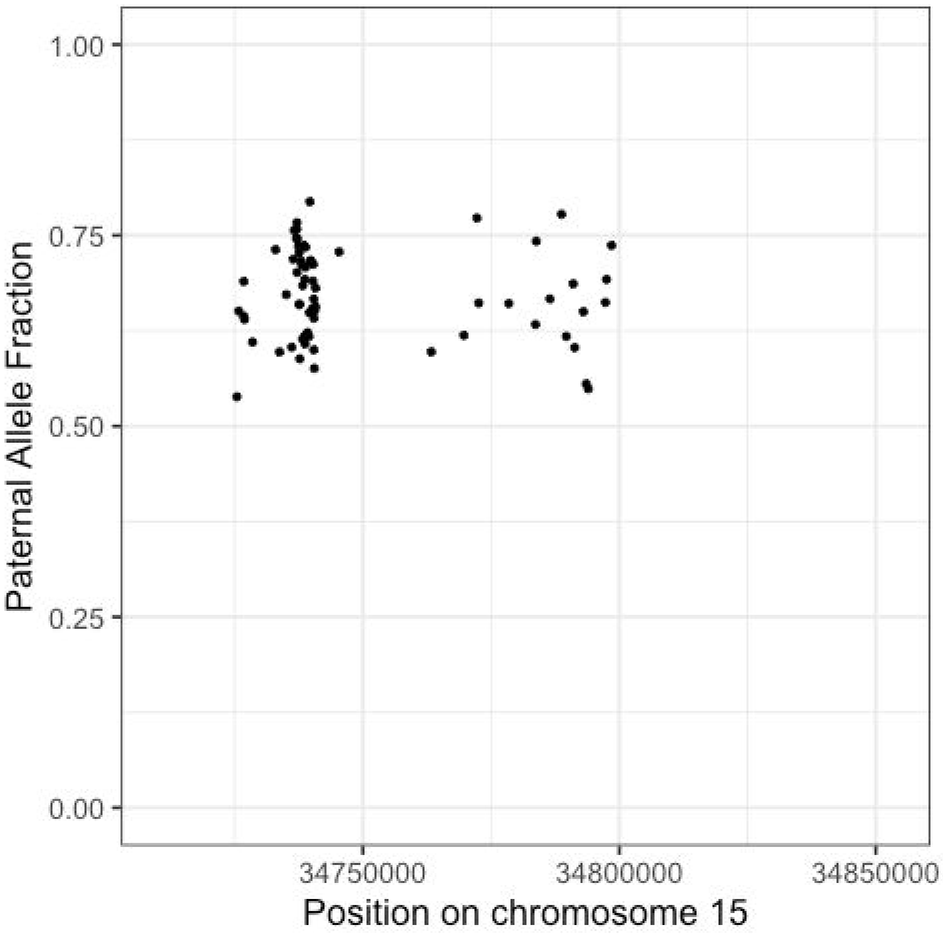


**S8 Fig.** Paralog specific variant analysis of one of two IBD discordancies at the *GOLGA8A/B* locus. In this family, the IBD state between siblings at this locus is zero, and the diploid copy numbers are father: 1; mother: 2; sibling 1: 3; sibling 2: 1. Plotted are allele fractions of SNP variants present only in the father's copy of the segment on the y axis, and position on chromosome 15 (within the CNV region) on the x axis. Although the father had only a single copy of the segment at this genomic region, the clustering of allele fractions at 66% in the sibling with copy number 3 indicates that two copies of the father's segment were transmitted to that child -- and therefore a *de novo* duplication of the segment appeared in the father.

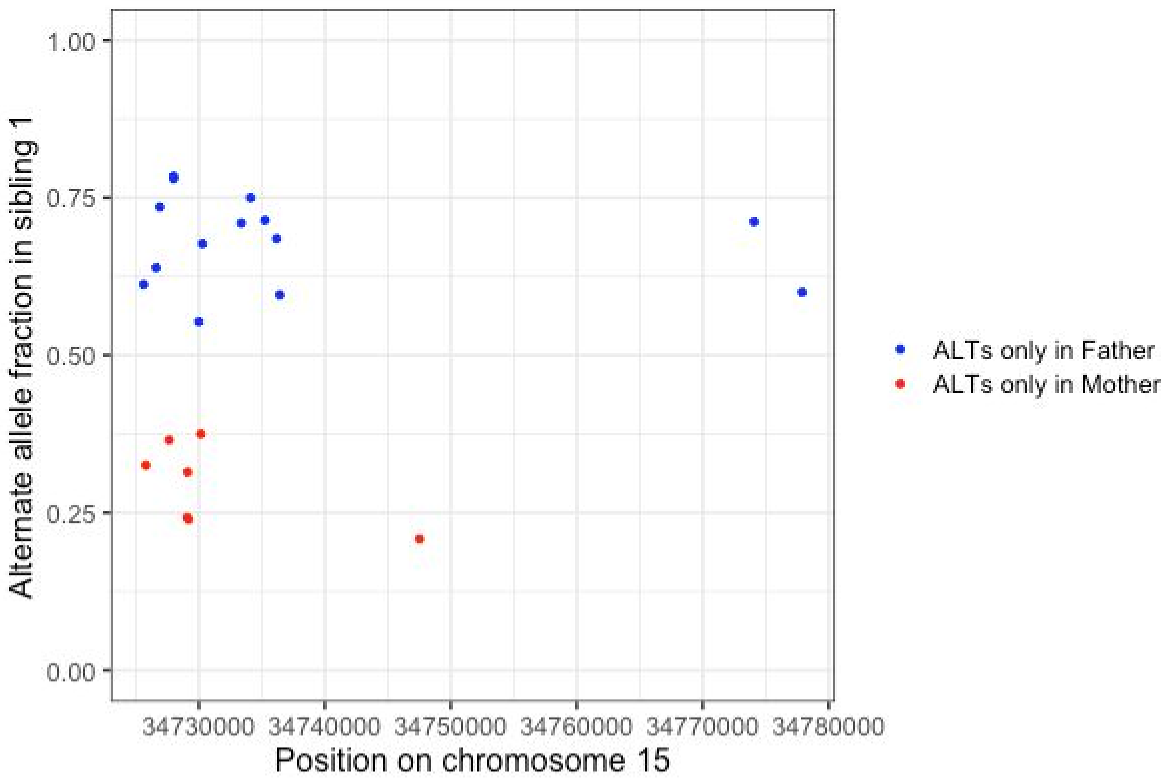


**S9 Fig.** Paralog specific variant analysis of the second of two IBD discordancies at the *GOLGA8A/B* locus (see **S8 Fig** for the other instance). In this family, the IBD state between siblings at this locus is zero, and the diploid copy numbers are father: 2; mother: 1; sibling 1: 3; sibling 2: 1. Plotted on the y-axis are the allele fractions of SNP alleles which were found in sibling 1 that were present in only one, but not both, of the parents, and were not found in sibling 2. Position on chromosome 15 (within the CNV region) is plotted on the x-axis. Variants which were present only in the father cluster at 66% allele fraction, while variants that were specific to the mother cluster at 33% allele fraction. This indicates that sibling 1 received one copy of the mother's segment at this locus, and two copies of one of the father's segments, which was duplicated *de novo*.

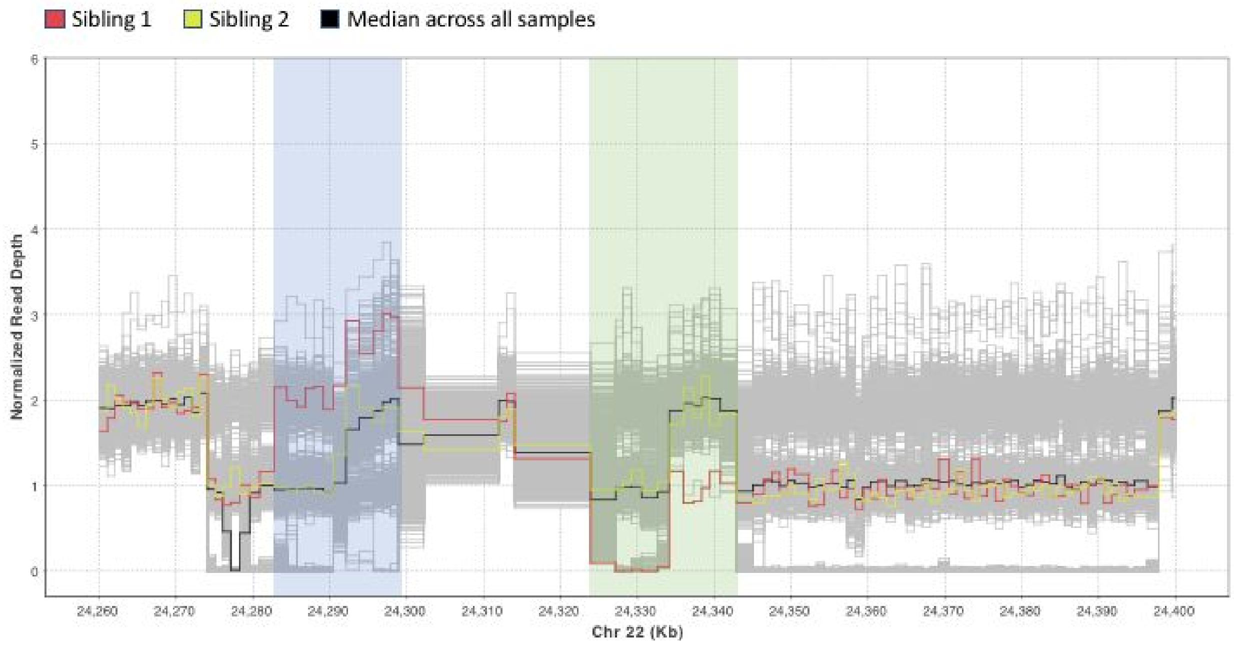


**S10 Fig.** A complex rearrangement at the *DDTL* / *GSTT2* locus. Normalized read depth profiles for the ASD cohort are shown across the locus, where each horizontal line represents the normalized read depth in a single sample in each binned genomic location. Profiles for the two siblings in which the mutation was detected are highlighted as follows: Sibling 1: Red; Sibling 2: Yellow. The siblings are IBD 2 at the region. Divergence of the two sibling's profiles shows that Sibling 1 has undergone a complex rearrangement involving a duplication of the sequence present at approximate coordinates 24,282,000-24,300,000 on chr22 (highlighted in blue) and a deletion of a segment at approximate coordinates 24,324,000-24,343,000 (highlighted in green).

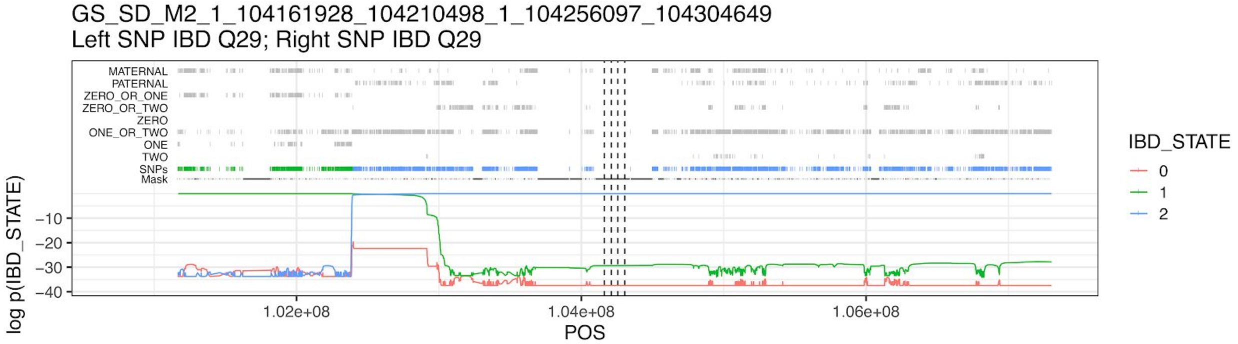


**S11 Fig**. A visualization of the IBD state as supported by SNP markers in the vicinity of a potential CNV mutation. The top tracks show the location of SNPs with various characteristics used by the IBD calculation algorithm to determine IBD state in the vicinity of a copy number variant -- in this case within segmental duplications at the *AMY* locus, indicated by dotted vertical lines. Top two tracks: MATERNAL and PATERNAL indicate whether the SNP supports inheritance of a maternal or paternal haplotype given that the IBD state is 1. The next six tracks highlight which SNPs could support which IBD states. For example, a SNP marked ONE_OR_TWO could support either IBD 1 or IBD 2 status. Below these all SNPs are shown, colored according to the IBD state with the maximum posterior probability, as well as the mask that was used to filter out SNPs for the purposes of IBD state calculation. At the bottom of the plot, the log10 posterior likelihoods of each IBD state at each SNP are shown. In this case, the quartet is in IBD 2 state at the center of the plot, with a transition to IBD 1-M 1.5 Mb upstream.

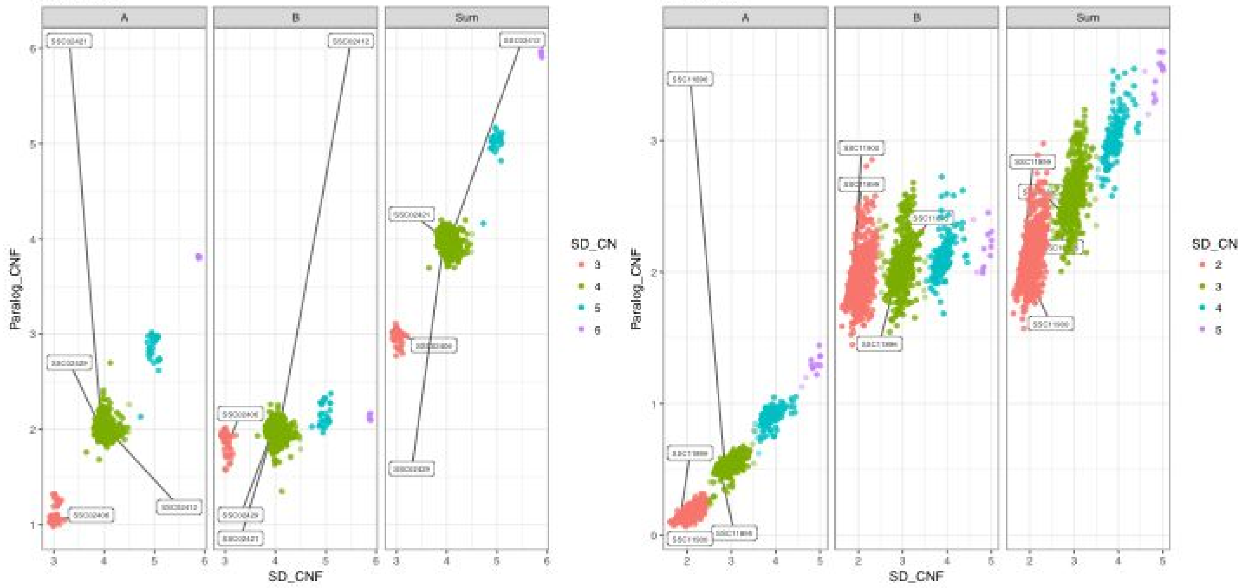


**S12 Fig.** Copy number variation at paralogs of segmental duplications. Data for two different pairs of annotated segmental duplications are shown. Within each pair, we plot the estimated copy number computed by measuring read depth in only the upstream copy of the SD (Paralog A), the downstream copy (Paralog B), and the sum of the two paralog copy number estimates, on the y-axis. On the x-axis we plot the estimate of the copy number for the pair of paralogs, based on read depth at pairs of loci that have identical sequence content in both paralogs. In each of these cases it is clear that Paralog A is varying in copy number across the cohort and not Paralog B, so we used only the IBD state for Paralog A in mutation analysis. For each of these examples, the copy number estimates for the four members of a quartet determined to have a mutation are highlighted.

### Supplementary Tables

**S4 Table 4.** A list of studies associating the copy number of the *GSTT1* and *GSTT2* genes with various phenotypes.

| **Author** | **Title** | **Pubmed ID** |
|---|---|---|
| Lavrov AV et al. | Copy number variation analysis in cytochromes and glutathione S-transferases may predict efficacy of tyrosine kinase inhibitors in chronic myeloid leukemia. | 28902850 |
| Ramírez B et al. | Copy number variation profiling in pharmacogenetics CYP-450 and GST genes in Colombian population. | 31324178 |
| Nørskov MS et al. | Copy number variation in glutathione-S-transferase T1 and M1 predicts incidence and 5-year survival from prostate and bladder cancer, and incidence of corpus uteri cancer in the general population. | 20514077 |
| Hersoug LG et al. | The relationship of glutathione-S-transferases copy number variation and indoor air pollution to symptoms and markers of respiratory disease. | 21651749 |
| Nørskov MS et al. | Copy number variation in glutathione S-transferases M1 and T1 and ischemic vascular disease: four studies and meta-analyses. | 21562205 |
| Butler MW et al. | Glutathione S-transferase copy number variation alters lung gene expression. | 21349909 |
| Karageorgi S et al. | GSTM1 and GSTT1 copy number variation in population-based studies of endometrial cancer risk. | 21558497 |
| Nørskov MS et al. | High-throughput genotyping of copy number variation in glutathione S-transferases M1 and T1 using real-time PCR in 20,687 individuals. | 19026998 |
| Bustamante M et al. | Influence of fetal glutathione S-transferase copy number variants on adverse reproductive outcomes. | 22676722 |
| Rodríguez-Santiago B et al. | Association of common copy number variants at the glutathione S-transferase genes and rare novel genomic changes with schizophrenia. | 19528963 |
| Christiansen L et al. | A longitudinal study of the effect of GSTT1 and GSTM1 | 16574194 |

| | | |
|---|---|---|
| | gene copy number on survival. | |
| Luo J et al. | Urinary polyphenols, glutathione S-transferases copy number variation, and breast cancer risk: results from the Shanghai women's health study. | 21557334 |
| Funke S et al. | Genetic polymorphisms in GST genes and survival of colorectal cancer patients treated with chemotherapy. | 20017670 |
| Orlewski J et al. | Effects of genetic polymorphisms of glutathione S-transferase genes (GSTM1, GSTT1, GSTP1) on the risk of diabetic nephropathy: a meta-analysis. | 26252359 |
| Bremer S et al. | Glutathione Transferase Gene Variants Influence Busulfan Pharmacokinetics and Outcome After Myeloablative Conditioning | 25565670 |
| Kap EJ et al. | Genetic variants in the glutathione S-transferase genes and survival in colorectal cancer patients after chemotherapy and differences according to treatment with oxaliplatin. | 24842074 |
| ThekkePurakkal AS et al. | Genetic variants in CYP and GST genes, smoking and risk for head and neck cancers: a gene-environment interaction hospital-based case-control study among Canadian Caucasians. | 30938417 |
| Cong LX et al. | Single nucleotide polymorphisms in glutathione S-transferase P1 and M1 genes and overall survival of patients with ovarian serous cystadenocarcinoma treated with chemotherapy. | 27073511 |
| Ibarrola-Villava M et al. | Role of glutathione S-transferases in melanoma susceptibility: association with GSTP1 rs1695 polymorphism. | 22251241 |
| Buchard A et al | Multiplex PCR detection of GSTM1, GSTT1, and GSTP1 gene variants: simultaneously detecting GSTM1 and GSTT1 gene copy number and the allelic status of the GSTP1 Ile105Val genetic variant. | 17916600 |
| Brasch-Andersen C et al. | Possible gene dosage effect of glutathione-S-transferases on atopic asthma: using real-time PCR for quantification of GSTM1 and GSTT1 gene copy numbers. | 15300848 |
| Borst L et al. | Gene dose effects of GSTM1, GSTT1 and GSTP1 polymorphisms on outcome in childhood acute lymphoblastic leukemia. | 22215096 |
| Yang F et al. | GSTT1 deletion is related to polycyclic aromatic | 24586676 |

| | | |
|---|---|---|
| | hydrocarbons-induced DNA damage and lymphoma progression. | |
| McLellan RA et al. | Characterization of a human glutathione S-transferase mu cluster containing a duplicated GSTM1 gene that causes ultrarapid enzyme activity. | 9415705 |
| Zhao Y et al. | Linkage disequilibrium between two high-frequency deletion polymorphisms: implications for association studies involving the glutathione-S transferase (GST) genes. | 19424424 |
| Timofeeva M et al. | A multiplex real-time PCR method for detection of GSTM1 and GSTT1 copy numbers. | 19161997 |
| Naseer MI et al. | Copy number variations in Saudi family with intellectual disability and epilepsy. | 27766957 |
| Zheng X et al. | Association of maternal CNVs in GSTT1/GSTT2 with smoking, preterm delivery, and low birth weight. | 24194744 |

**S5 Table.** Filtering process for identifying CNV mutations.

| | **CNV** | **Seg Dup** | **Total** |
|---|---|---|---|
| Total qCNVs | 93,984,078 | 1,677,802 | 95,661,880 |
| All family members have called copy numbers | 80,889,711 | 828,308 | 81,718,019 |
| All copy number calls at CNQ >= 40 | 73,625,889 | 330,694 | 73,956,583 |
| Not within 1MB of IBD state transition, IBD call confident | 69,026,896 | 313,010 | 69,339,906 |
| Not overlapping with dispersed duplication | 68,747,128 | 313,010 | 69,060,138 |
| Not within a 1MB gap in the SNP mask | 67,815,685 | 285,091 | 68,100,776 |
| Family does not contain a sample flagged for high variants in CNV discovery | 60,305,347 | 254,560 | 60,559,907 |
| Quartet not an outlier for discordant qCNV count | 57,187,080 | 242,282 | 57,429,362 |
| From a site with at least one discordant qCNV but not more than 1% of qCNVs discordant | 1,068,753 | 14,921 | 1,083,674 |
| Keep only discordant qCNVS | 2,484 | 38 | 2,522 |
| Deduplicate qCNVs and subset to ASD cohort | 571 | 28 | 599 |
| Manual curation: | | | |
| Filtered for CN genotypes | 374 | 5 | 379 |
| Filtered for bad IBD state or nearby recombination | 33 | 4 | 37 |
| Filtered for having multiple discordant qCNVs and suspect depth genotypes | 44 | 1 | 45 |
| Potential rare dispersed duplication | 72 | 1 | 73 |
| Potential somatic events | 5 | 0 | 5 |
| Redundant with other calls | 7 | 13 | 20 |
| One additional event detected in single end of a segmental duplication | 0 | 1 | 0 |
| **Final Mutation Calls** | **36** | **5** | **41** |

We began by considering the copy number state for every family in our input cohorts at every CNV site detected in our input CNV set. We call the grouping of the copy number state of a quartet and the IBD state at a CNV site a qCNV. Beginning with 95.6 million qCNVs, we applied a series of automatic filters and manual curation to identify a set of 41 mutation calls (see methods for details).
